# Multiple dimensions underlying the functional organisation of the language network

**DOI:** 10.1101/2021.02.05.429902

**Authors:** Victoria J. Hodgson, Matthew A. Lambon Ralph, Rebecca L. Jackson

## Abstract

Understanding the different neural networks that support human language is an ongoing challenge for cognitive neuroscience. Which divisions are capable of distinguishing the functional significance of regions across the language network? A key separation between semantic cognition and phonological processing was highlighted in early meta-analyses, yet these seminal works did not formally test this proposition. Moreover, organisation by domain is not the only possibility. Regions may be organised by the type of process performed, as in the separation between representation and control processes proposed within the Controlled Semantic Cognition framework. The importance of these factors was assessed in a series of activation likelihood estimation meta-analyses that investigated which regions of the language network are consistently recruited for semantic and phonological domains, and for representation and control processes. Whilst semantic and phonological processing consistently recruit many overlapping regions, they can be dissociated (by differential involvement of bilateral anterior temporal lobes and superior temporal gyri) only when using both formal analysis methods and sufficient data. Both semantic and phonological regions are further dissociable into control and representation regions, highlighting this as an additional, distinct dimension on which the language network is functionally organised. Furthermore, some of these control regions overlap with multiple-demand network regions critical for control beyond the language domain, suggesting the relative level of domain-specificity is also informative. Multiple, distinct dimensions are critical to understand the role of language regions. Here we present a proposal as to the core principles underpinning the functional organisation of the language network.

**SIGNIFICANCE STATEMENT:** Traditional theories of the organisation of the cortical language network separate areas based on the type of information processed, such as distinct regions representing the meaning of words or their sounds. Here, we re-analyse and update a seminal study, to directly compare these networks. This differentiates some language regions, yet a high degree of overlap is found, suggesting this division alone, does not provide a full mapping of the language network. Additional dimensions, reflecting different kinds of information, are demonstrated to underlie the functional organisation of the language network. First, representation and control processes engage distinct regions in each subdomain. Secondly, comparison outside of language highlights the importance of a ‘domain-specificity’ dimension, distinguishing subdomain-specific semantic control and domain-general control regions.

## INTRODUCTION

The complexity of language and its neural substrates have long drawn attention within cognitive neuroscience. Though it remains unclear how precisely language is organised in the brain, different facets of language can be dissociated from one another at the behavioural and neuropsychological levels; in particular, a key division between semantics and phonology ^1–4^. Yet questions remain as to whether these processes are supported by different neural networks, and to what degree, and in what manner, these networks interact. Moreover, semantics and phonology may each be supported by multiple interactive networks, underpinning dissociable processes at the cognitive level. Process-based divisions, such as the separation between representation and control processes previously specified within the domain of semantic cognition ^5–7^, may provide additional, and perhaps orthogonal, information regarding the functional role of regions of the language network. To investigate whether these domain- and process-based divisions delineate the functional roles of regions across the language network, we conducted a series of neuroimaging meta-analyses to establish the patterns and locations of consistent activation within and across three domains (semantic cognition, phonological processing and non-verbal working memory) and across varying levels of control demands (identifying regions specialised for representation versus control processes).

The neural correlates of language may differ based on the importance of semantic versus phonological demands. Phonological processing is the perception, analysis and use of language sounds to understand and produce spoken and written language ^8,9^. The production and perception of phonological information converge in the temporoparietal junction, involving posterior superior temporal lobe, inferior superior marginal gyrus (SMG), and left inferior frontal gyrus (IFG) ^10–15^. Semantic cognition refers to the storage, retrieval and manipulation of multimodal conceptual knowledge. A distributed network of regions are implicated in multimodal semantic cognition, including IFG, posterior temporal cortex, inferior parietal sulcus (IPS) and anterior temporal lobe (ATL). The ATL acts as a multimodal hub mediating between numerous modality-specific regions, or “spokes” distributed across sensory and association cortices ^5–7,16–19^. Neighbouring and potentially overlapping areas of the left posterior temporal lobe, inferior parietal lobe and IFG have been implicated in both the semantic and phonological networks; some of these apparent similarities, however, may be obscuring graded differences in specialisation for each language domain, and there is a need for a more direct comparison. For example, neuroimaging provides some evidence of relative functional specialisation for semantic cognition in more ventral and caudal, and phonology in more dorsal, rostral left IFG ^8,20^. Transcranial magnetic stimulation induced virtual lesions of dorsal left IFG impair performance on phonological tasks ^21,22^, while disrupting the ventral left IFG diminishes semantic performance ^23,24^. Left IFG may therefore constitute two specialised subregions for different aspects of language, or a single complex with graded functional specialisation ^25^. Similarly, the inferior parietal lobe (IPL) is not a homogenous region, with debate as to which subregions are implicated in semantic and phonological processing and beyond language ^26–30^.

The neural correlates of phonology and semantics were the subject of a seminal review of organisation across the language network by Vigneau et al. ^3,4^. The authors identified neuroimaging studies targeting semantics, phonology and sentence processing and mapped the peak group activations (see Figure 1). Both domains resulted in highly distributed networks of peaks principally centred on the left hemisphere. The distribution of peaks clustered upon bilateral precentral, superior temporal and inferior frontal gyri, and left posterior temporal lobe and supramarginal gyrus for phonology, whilst implicating bilateral IFG, posterior inferior and superior temporal gyri and left angular gyrus (AG), planum temporale and ATL in semantics. Based on visual examination, Vigneau et al. ^3,4^ concluded the distribution of peaks reflected distinct networks for phonology and semantic cognition. As this was published prior to the widespread use of formal neuroimaging meta-analyses ^31^, it was not possible to determine which regions were significantly consistently involved, or to directly contrast the areas implicated in semantic and phonological processing. Despite these unavoidable limitations, the paper continues to be highly cited as evidence of a strong dissociation between the neural correlates of semantics and phonology. Accordingly, in this investigation we apply formal activation likelihood estimation (ALE) analyses and also update the neuroimaging database, to determine the regions involved in phonological and semantic cognition and directly contrast the likelihood of activation in these two subdomains of language.

**Figure 1.**
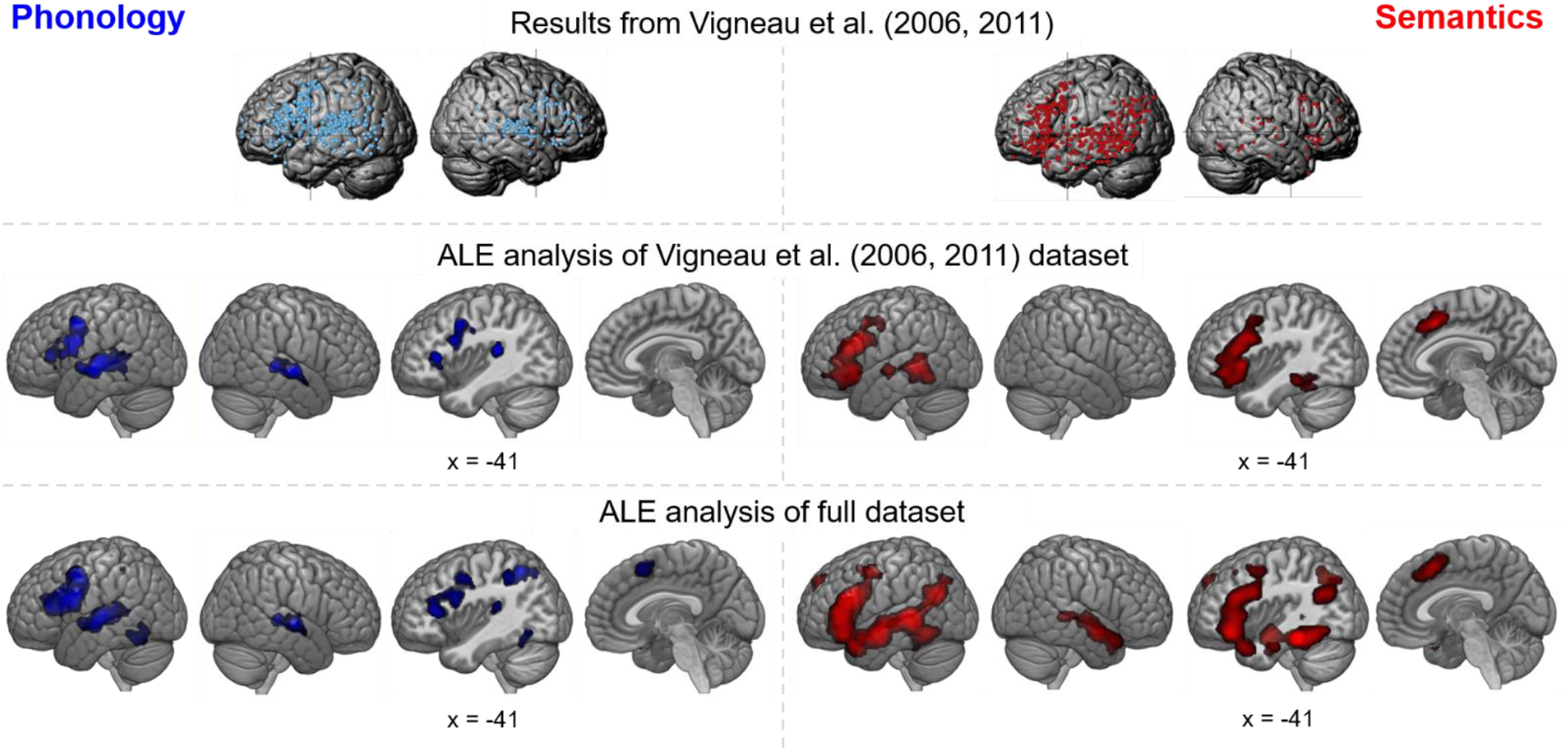
Results for the phonology and semantics domains are shown in blue (left column) and red (right column) respectively. Top row: original results by Vigneau et al. (2006, 2011); figures are reproduced from Vigneau et al. (2011). Middle row: formal ALE meta-analysis of the recreated Vigneau et al. (2006, 2011) datasets, with 615 foci from 44 experiments for phonology, and 788 foci from 70 experiments for semantics. Bottom row: meta-analysis of full datasets, with 870 foci from 64 experiments for phonology, and 3606 foci from 272 experiments for semantics.

In the years since Vigneau et al.’s ^3,4^ reports, an additional process-based distinction has gained increasing recognition, both within the language domain and in cognition more broadly. Alongside the representation of information relevant for each domain, executive control processes are critical for the context-appropriate access and use of this information ^5,32–34^. The crucial nature of the division between control and representation processes has been highlighted for semantic cognition, where the controlled semantic cognition (CSC) framework proposes relative functional specialisation of regions for representation versus control, resulting in distinct neural architectures ^5,35,36^. Semantic control refers to the contextual selection and manipulation of semantic information necessary for task-appropriate behaviour, particularly when dominant features must be supressed, or subordinate meanings must be accessed ^7,32,37^. A subset of the regions involved in general semantic processing are specifically implicated in semantic control: the IFG, left posterior lateral temporal lobe (specifically, posterior middle and inferior temporal gyri) and bilateral dorsomedial prefrontal cortex (dmPFC) ^7,29,38^.

Like semantic cognition, phonology is supported by control processes, though these have not been emphasised to the same extent, nor have their neural correlates been clearly isolated to date. However, some functional neuroimaging work has found activation in left IFG, precentral gyrus, SMG and dmPFC for controlled, effortful phonological tasks (e.g., tasks in which participants must regularise the pronunciation of irregular words) ^29,39,40^. To determine whether control versus representation is a critical organisational principle of the language network, this investigation used meta-analyses to isolate regions critical for phonological and semantic control. Control regions could be particular to a subdomain of language, specific to the broader domain of language, or domain general. Therefore, considering an additional ‘domain-specificity’ factor could help further elucidate the organisation of the language network. This may be tested by assessing whether the same regions are implicated in semantic control, phonological control and additional analyses targeting domain-general control. The multiple demand network (MDN) is a set of brain regions activated across a broad range of executive tasks, indicating support for a wide range of tasks across domains ^33^, and includes the posterior inferior frontal sulcus, intraparietal sulcus, anterior insula, pre-supplementary area and anterior cingulate cortex ^33,41,42^. This may overlap regions implicated in phonology (such as inferior parietal cortex ^26,43^) or semantics. Although largely distinct, the semantic control areas identified in Jackson’s ^38^ meta-analysis of semantic control may overlap Assem et al.’s ^44^ extended multiple demand network in a portion of the IFG and left posterior inferior temporal cortex.

There were three steps to the present study. Firstly, formal ALE analyses compared the regions implicated in the semantic and phonological subdomains, testing the supposition that these subdomains rely on distinct neural correlates. To determine the relative impact of changing method of analysis versus dataset, this was first performed using Vigneau et al.’s ^3,4^ dataset, and then with updated semantic and phonological datasets. Next, ALE analyses of phonological control and semantic control were compared to the full results for phonological and semantic cognition, to determine the importance of control versus representation processes in these language subdomains. Finally, the specificity of the identified control regions, was considered through comparison to two activation maps of domain-general control: an ALE analysis of the *n*-back working memory task (a formal analysis allowing careful control including the elimination of verbal stimuli); and the multiple demand network map from Fedorenko et al. ^45^ (used as a standard measure of the MDN, reflecting a larger breadth of executive tasks). Control regions could be involved in one particular language subdomain, language in general or all cognition,

## RESULTS

Meta-analyses were employed to ask several questions about the neural substrates of language; 1) which brain regions are consistently activated across studies employing semantic and phonology tasks, 2) how distinct are the networks for semantic and phonological processing, 3) does control versus representation provide an additional informative way to separate the functions of regions in the language network and 4) how domain-specific or domain-general are the control areas implicated in semantic and phonological tasks? In considering the division between the neural correlates of semantic cognition and phonology, we first consider whether a formal ALE meta-analysis would reveal a division between semantics and phonology using the datasets employed by Vigneau, before employing updated datasets to assess whether a division can be identified with the use of both modern methods and up-to-date data.

### Separating the Language Network by Phonological and Semantic Subdomain

The formal meta-analysis of the Vigneau et al. ^3,4^ phonology dataset, revealed three clusters (Table 1, Figure 1). One cluster encompasses mid and posterior superior temporal gyrus (STG), extending into the Sylvian fissure and posterior middle temporal gyrus (MTG). The right hemisphere analogue of this cluster was less extensive, comprising most of the middle to posterior STG with limited expansion into the superior temporal sulcus (STS). An additional left hemisphere cluster lay within the left precentral and inferior frontal gyri (including pars opercularis and triangularis). The formal ALE analysis of Vigneau et al.’s semantic dataset was almost entirely left-lateralised (Figure 1, peaks provided in Table 2). A large cluster covered left IFG (pars opercularis, triangularis and orbitalis), extending partially into middle frontal and precentral gyri. Another was centred in the posterior MTG, extending dorsally into the STG and ventrally/medially into the inferior temporal gyrus (ITG) and fusiform. The final cluster was located toward the midline, in the left dmPFC, pre-supplementary motor area and paracingulate gyrus.

**Table 1:** Phonology activation likelihood using Vigneau dataset

| Cluster | Region of Activation | Maximum ALE Value | Z Score | Peak MNI Coordinate |  |  |
| --- | --- | --- | --- | --- | --- | --- |
|  |  |  |  | x | y | z |
| 1 | Left superior temporal gyrus/middle temporal gyrus | 0.039 | 6.976 | -62 | -14 | -4 |
|  |  | 0.038 | 6.889 | -60 | -22 | 2 |
|  |  | 0.025 | 5.038 | -58 | -4 | -4 |
|  |  | 0.024 | 4.980 | -42 | -32 | 16 |
|  |  | 0.024 | 4.926 | -64 | -34 | 4 |
|  |  | 0.022 | 4.596 | -54 | -42 | 6 |
|  |  | 0.018 | 4.081 | -52 | -44 | -2 |
|  |  | 0.018 | 3.916 | -66 | -42 | 12 |
|  |  | 0.016 | 3.687 | -62 | -50 | 10 |
|  |  | 0.014 | 3.372 | -52 | -46 | -14 |
|  |  | 0.014 | 3.232 | -48 | -22 | 4 |
| 2 | Left precentral gyrus/inferior frontal gyrus | 0.038 | 6.902 | -50 | 6 | 22 |
|  |  | 0.034 | 6.352 | -48 | 4 | 44 |
|  |  | 0.026 | 5.231 | -44 | 30 | 10 |
|  |  | 0.022 | 4.574 | -50 | 16 | 12 |
|  |  | 0.016 | 3.633 | -36 | 24 | -2 |
| 3 | Right superior temporal gyrus/middle temporal gyrus | 0.035 | 6.492 | 62 | -10 | -4 |
|  |  | 0.029 | 5.702 | 62 | -30 | 2 |
|  |  | 0.017 | 3.877 | 52 | -26 | 8 |
|  |  | 0.016 | 3.713 | 52 | -18 | 6 |

**Table 2:** Semantic activation likelihood using Vigneau dataset

| Cluster | Region of Activation | Maximum ALE Value | Z Score | Peak MNI Coordinate |  |  |
| --- | --- | --- | --- | --- | --- | --- |
|  |  |  |  | x | y | z |
| 1 | Left inferior frontal gyrus/middle frontal gyrus | 0.051 | 7.767 | -46 | 18 | 24 |
|  |  | 0.049 | 7.520 | -44 | 24 | 18 |
|  |  | 0.041 | 6.622 | -44 | 22 | -2 |
|  |  | 0.034 | 5.747 | -38 | 36 | -6 |
|  |  | 0.034 | 5.709 | -44 | 22 | -10 |
|  |  | 0.030 | 5.316 | -36 | 28 | -14 |
|  |  | 0.027 | 4.926 | -42 | 40 | 6 |
|  |  | 0.026 | 4.702 | -50 | -6 | 44 |
|  |  | 0.025 | 4.552 | -38 | 8 | 46 |
|  |  | 0.018 | 3.510 | -48 | 34 | -22 |
| 2 | Left middle temporal gyrus/superior temporal gyrus/fusiform gyrus | 0.043 | 6.844 | -60 | -44 | -2 |
|  |  | 0.034 | 5.720 | -46 | -36 | -14 |
|  |  | 0.032 | 5.523 | -54 | -44 | 8 |
|  |  | 0.030 | 5.288 | -54 | -52 | -10 |
|  |  | 0.030 | 5.285 | -58 | -14 | -4 |
|  |  | 0.023 | 4.249 | -36 | -48 | -20 |
| 3 | Right dorsomedial prefrontal cortex | 0.044 | 6.953 | -2 | 16 | 44 |
|  |  | 0.019 | 3.369 | 12 | 26 | 32 |
|  |  | 0.018 | 3.498 | -14 | 30 | 46 |

Directly contrasting Vigneau et al.’s semantic and phonology datasets (Table 6, Figure 2) revealed two small clusters for the semantics > phonology analysis, located in the left fusiform gyrus and left pars triangularis, and four small clusters for phonology > semantics, located in each middle STG, left precentral gyrus, and left inferior precentral sulcus. None of these clusters were larger than 400mm^3^. Whilst these clusters may reflect true differences between semantics and phonology, these small differences do not provide strong evidence of distinct networks for phonological and semantic cognition.

**Figure 2.**
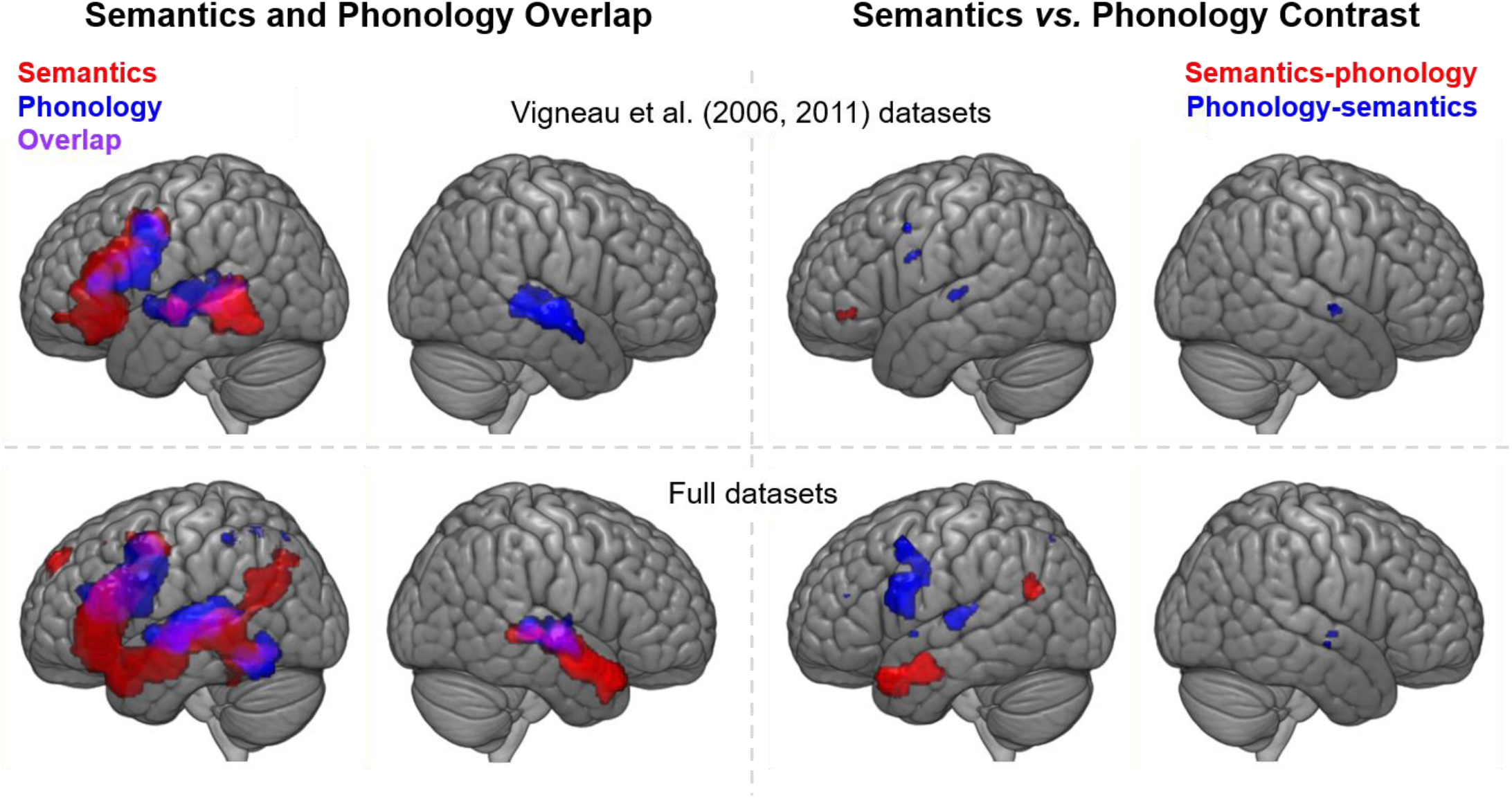
Top row: a comparison of the formal ALE analyses conducted for Vigneau et al.’s semantics (red) and phonology (blue) datasets. Bottom row: a comparison of the formal ALE analyses conducted on the full datasets. Left column: the semantic and phonological activation maps are shown overlaid. Overlap may be seen in violet. Right column: formal ALE contrasts (phonology > semantics in blue, semantics > phonology in red)

In comparison, the analyses of the larger, updated phonology and semantic databases generated more extensive ALE maps. For phonology (Figure 1, peaks given in Table 3), there are six clusters in total. Three of these are analogous, although more extensive, to the clusters identified with the Vigneau dataset: a large cluster encompassing left IFG (pars opercularis and some of pars triangularis) and a large swathe of precentral gyrus, and one cluster encompassing middle and posterior STG and extending ventrally into the STS in each hemisphere (albeit with a greater extent on the left). In addition, a cluster is located in the left SMG and superior parietal lobule, recruiting a small amount of the AG. Further clusters were located across the left fusiform and inferior temporal gyri and near the midline in the left dmPFC. The full semantic activation map, shown in Figure 1 (peaks given in Table 4), is considerably more extensive than the Vigneau result. A single, large cluster extends across the length of left temporal lobe, a large portion of lateral frontal cortex, insula and ventral aspects of the parietal lobe. The frontal contribution includes the inferior frontal, precentral and middle frontal gyri. In the temporal lobe, the cluster spans from the temporal pole to the planum temporale, recruiting STG, MTG and posterior inferior temporal, fusiform and parahippocampal gyri. This single cluster encompasses and extends beyond the two lateral left hemisphere clusters revealed in the Vigneau analysis. The second semantic cluster is located in the dmPFC, analogous to, yet more extensive than, the dmPFC cluster for Vigneau et al.’s data. Finally, the full semantic ALE analysis shows right hemisphere activation not revealed using the Vigneau dataset, encompassing the STG across the length of the temporal lobe, with some involvement of MTG.

**Table 3:**
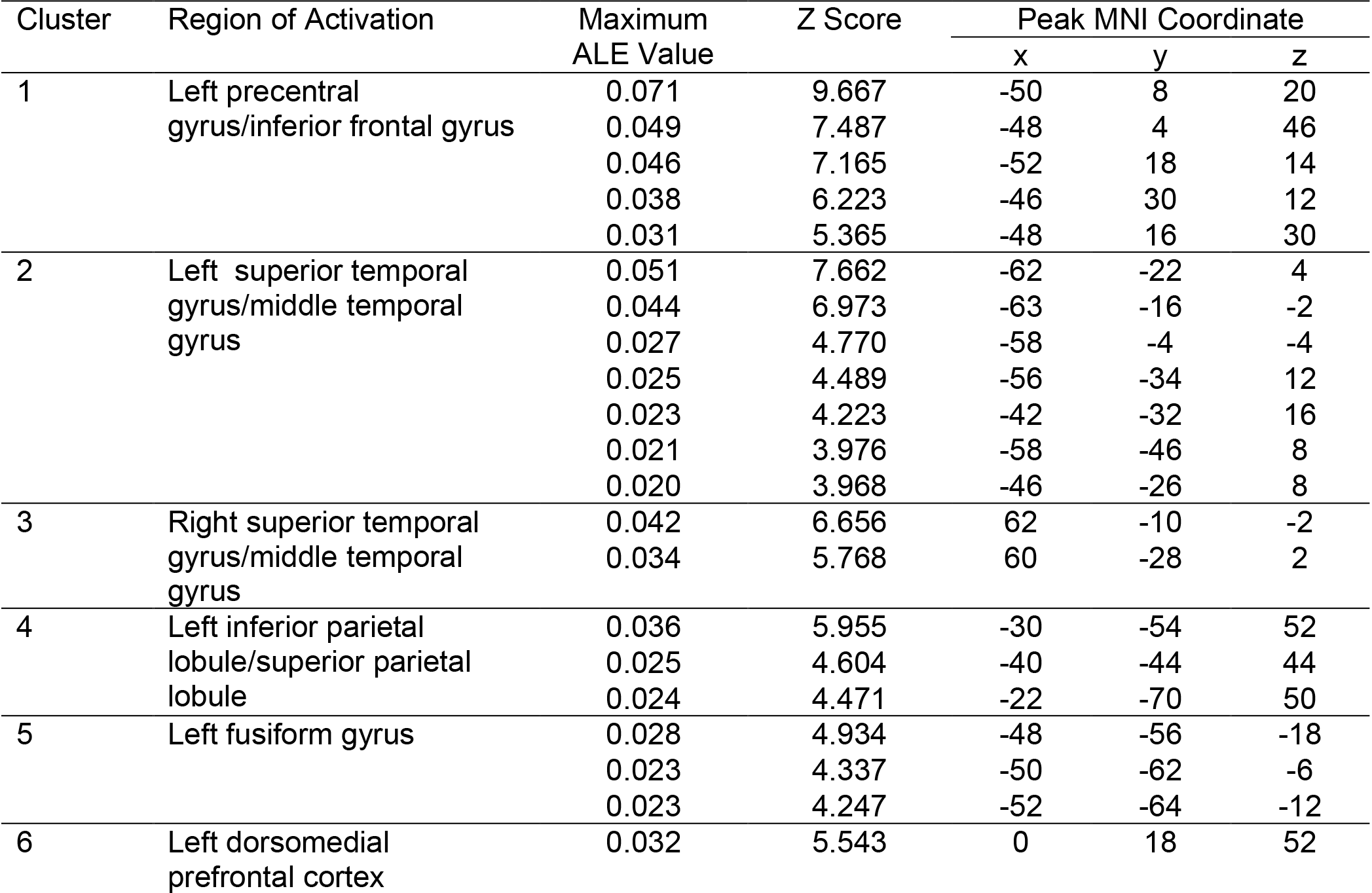
Phonology activation likelihood

**Table 4:**
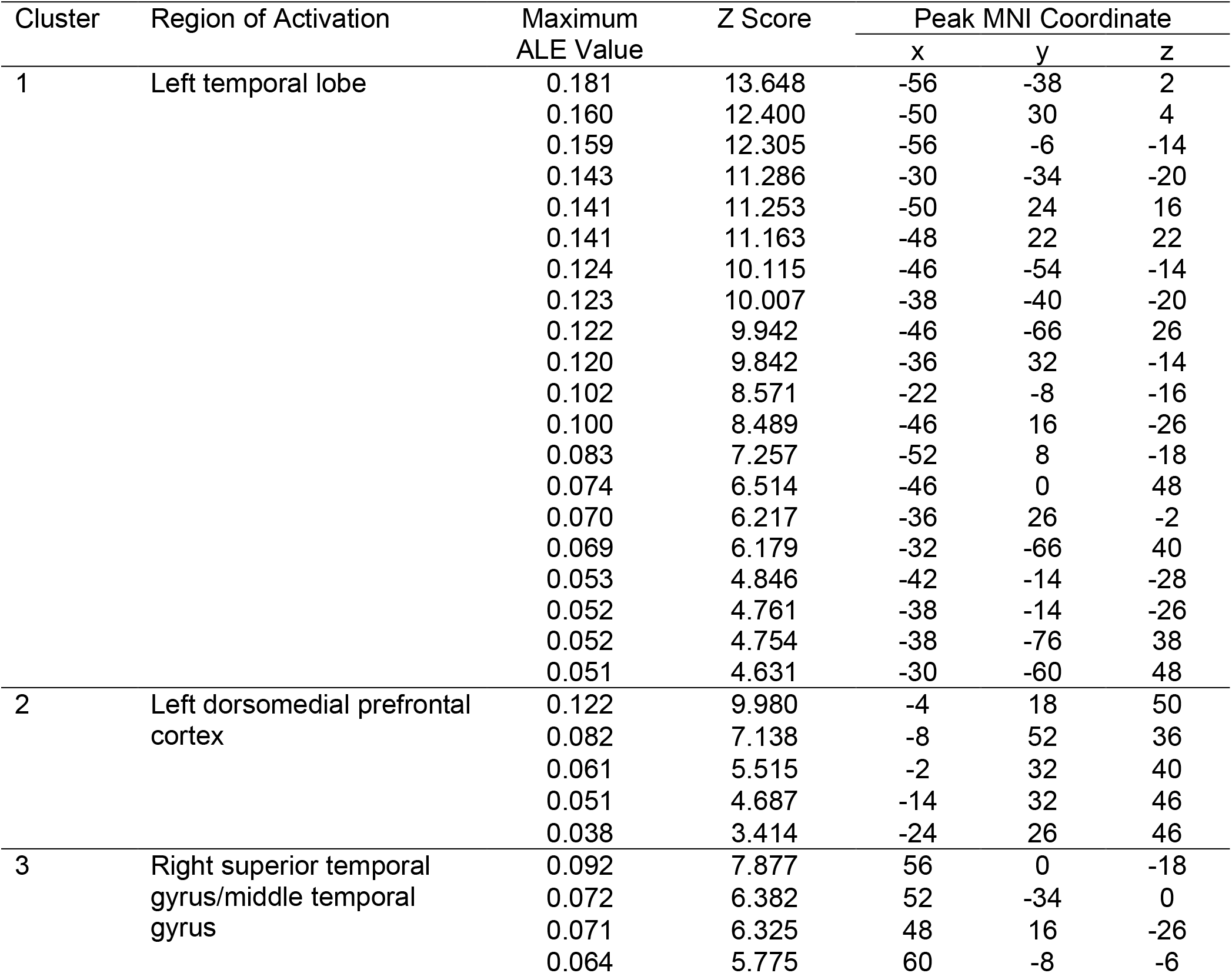
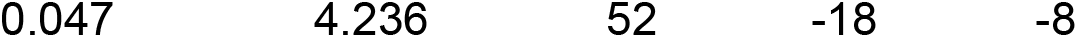
Semantic activation likelihood

Contrasting the full phonology and semantic activation likelihood maps revealed significant differences between the networks supporting the two domains (see Figure 2 & Table 5). Greater involvement in phonology was found for a large cluster within the frontal lobe, principally in the precentral gyrus, with some involvement of pars opercularis. Further differences were located in posterior STG and SMG, together comprising the majority of the parietal cluster implicated in phonology. Additional differences were identified in left and right STG, the left Sylvian fissure, right STS, left middle frontal gyrus and right dmPFC. Thus, the superior aspects of left temporal lobe, the left precentral gyrus, and left superior parietal lobule are consistently recruited for phonology more than semantic cognition. The semantics > phonology map is comprised of three clusters, with the largest located in the left parahippocampal and fusiform gyri, regions that were not implicated in phonology. Further clusters span the left ATL and planum temporale. Thus, the left ATL, fusiform gyrus, and planum temporale are consistently recruited for semantic cognition to a greater extent than phonological processing. Additional regions (pars triangularis and orbitalis, right ATL) were present for the semantic but not phonological subdomain, but these apparent differences did not reach statistical significance.

**Table 5:** Formal contrast phonology vs. semantics

| Cluster | Region of Activation | Z Score | Peak MNI Coordinate |  |  |
| --- | --- | --- | --- | --- | --- |
|  |  |  | x | y | z |
| *Phonology > semantics* |  |  |  |  |  |
| 1 | Left precentral gyrus | 3.891 | -55 | 5 | 26 |
|  |  | 3.719 | -62 | 13 | 23 |
|  |  | 3.540 | -48 | -7 | 30 |
| 2 | Left posterior superior temporal gyrus | 3.891 | -56 | -24 | 9 |
| 3 | Left precuneus | 0 | -24 | -66 | 51 |
|  |  | 3.540 | -20 | -61 | 52 |
| 4 | Left supramarginal gyrus | 3.719 | -42 | -42 | 40 |
|  |  | 3.291 | -34 | -50 | 38 |
| 5 | Right superior temporal gyrus | 3.719 | 54 | -6 | 0 |
|  |  | 3.540 | 56 | -10 | 0 |
| 6 | Left middle superior temporal gyrus | 3.719 | -55 | 2 | -1 |
| 7 | Left middle frontal gyrus | 3.540 | -44 | 40 | 20 |
| 8 | Right superior temporal sulcus | 3.195 | 70 | -12 | -6 |
| 9 | Right dorsomedial prefrontal cortex | 3.291 | 4 | 38 | 41 |
| *Semantics > phonology* |  |  |  |  |  |
| 1 | Left fusiform/parahippocampal gyri | 3.891 | -32 | -35 | -18 |
|  |  | 3.719 | -26 | -20 | -14 |
|  |  | 3.432 | -16 | -14 | -16 |
|  |  | 3.353 | -20 | -13 | -16 |
|  |  | 3.291 | -17 | -8 | -18 |
| 2 | Left posterior superior temporal sulcus/planum temporale | 3.891 | -51 | 1 | -23 |
|  |  | 3.719 | -46 | 16 | -22 |
| 3 | Left anterior middle temporal gyrus/superior temporal gyrus | 3.891 | -43 | -60 | 20 |
|  |  | 3.719 | -48 | -64 | 24 |

**Table 6:** Formal contrast phonology vs. semantics Vigneau dataset

| Cluster | Region of Activation | Z Score | Peak MNI Coordinate |  |  |
| --- | --- | --- | --- | --- | --- |
|  |  |  | x | y | z |
| *Phonology > semantics* |  |  |  |  |  |
| 1 | Left STG | 3.239 | -63 | -24 | 7 |
|  |  | 3.432 | -60 | -24 | 6 |
| 2 | Right STG | 3.353 | 63 | -7 | -3 |
| 3 | Left precentral gyrus | 3.291 | -60 | 2 | 27 |
| 4 | Left precentral sulcus | 3.719 | -50 | 4 | 42 |
| *Semantics > phonology* |  |  |  |  |  |
| 1 | Left IFG (pars triangularis) | 3.432 | -45 | 38 | -5 |
|  |  | 3.239 | -41 | 43 | -6 |
| 2 | Left fusiform/parahippocampal gyri | 3.432 | -38 | -48 | -16 |

The neural correlates of semantics and phonology are dissociable despite considerable overlap between the networks recruited. However, substantial separation was only possible with both formal ALE meta-analyses and the additional data present in the full dataset; application of formal analyses alone was not sufficient. This may be due to the increased power of a larger dataset, or various methodological improvements in more recent studies, such as increased sample sizes, multi-banding^46,47^ or the development of fMRI techniques to reduce signal loss in regions such as the anterior temporal lobe ^48–50^.

### Separating the Language Network by Representation vs. Control Processes

To determine whether control versus representation is an informative principle of organisation of the language network, ALE analyses of semantic and phonological control were conducted. Semantic control analyses were performed as in Jackson ^38^ identifying the same five clusters (Figure 3 & Table 7). The largest and most significant of these encompasses the entirety of the left IFG, and extends somewhat into the precentral gyrus and orbitofrontal cortex. The second cluster is located in left posterior temporal cortex, namely the posterior MTG/ITG. A third cluster in bilateral dmPFC overlaps with the dmPFC cluster for general semantics, though does not extend as rostrally. In the right hemisphere, two smaller clusters are located in the IFG, one in pars triangularis, and one more ventrally in pars orbitalis and the insula. Control versus representation demands split the semantic network; the IFG, left dmPFC and left posterior temporal cortex form the semantic control network, whereas the remaining regions (bilateral temporal lobe and inferior parietal cortex) likely reflect semantic representation processes.

**Figure 3.**
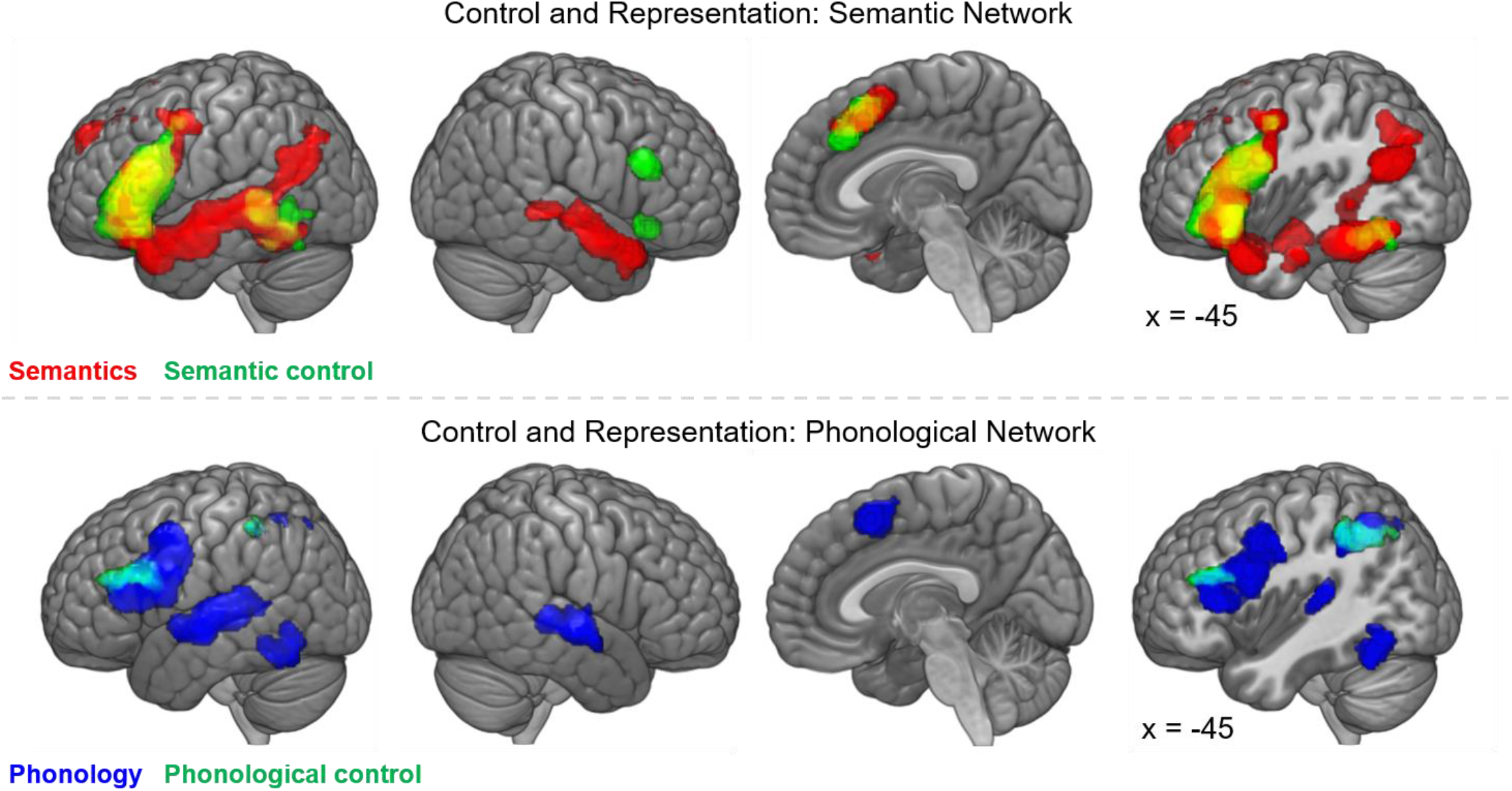
Top row: activation map for semantics domain (red) overlaid with semantic control (green, overlap in yellow). Bottom row: activation map for full phonological domain (blue) overlaid with phonological control, represented by hard > easy phonology formal ALE contrast (green, overlap in cyan).

**Table 7:** Semantic control activation likelihood

| Cluster | Region of Activation | Maximum ALE Value | Z Score | Peak MNI Coordinate |  |  |
| --- | --- | --- | --- | --- | --- | --- |
|  |  |  |  | x | y | z |
| 1 | Left inferior frontal gyrus | 0.093 | 10.679 | -48 | 22 | 20 |
|  |  | 0.060 | 7.809 | -46 | 24 | -2 |
|  |  | 0.055 | 7.402 | -50 | 30 | 0 |
|  |  | 0.037 | 5.495 | -34 | 26 | -6 |
|  |  | 0.036 | 5.378 | -46 | 40 | -10 |
|  |  | 0.034 | 5.134 | -48 | 34 | -12 |
|  |  | 0.029 | 4.565 | -30 | 24 | -16 |
|  |  | 0.024 | 3.909 | -44 | 2 | 48 |
|  |  | 0.020 | 3.349 | -38 | 28 | -22 |
| 2 | Left middle temporal gyrus/inferior temporal gyrus/fusiform | 0.039 | 5.725 | -54 | -42 | 4 |
|  |  | 0.037 | 5.500 | -46 | -48 | -16 |
|  |  | 0.037 | 5.478 | -46 | -56 | -12 |
|  |  | 0.036 | 5.386 | -56 | -46 | -4 |
|  |  | 0.021 | 3.514 | -50 | -68 | -2 |
| 3 | Left dorsomedial prefrontal cortex | 0.058 | 7.697 | -2 | 20 | 52 |
|  |  | 0.034 | 5.225 | 2 | 28 | 36 |
|  |  | 0.025 | 4.047 | -4 | 8 | 58 |
| 4 | Right insula/inferior frontal gyrus | 0.046 | 6.502 | 32 | 24 | -6 |
|  |  | 0.019 | 3.294 | 30 | 18 | -18 |
| 5 | Right middle frontal gyrus | 0.044 | 6.217 | 50 | 24 | 26 |

Phonological control, represented by the formal contrast of hard > easy phonology, is displayed in Figure 3 (also see Table 8). This implicated the inferior parietal lobule (with a cluster spanning supramarginal and angular gyri and extending medially) and the IFG (specifically pars opercularis). This does not identify all the regions implicated in both semantic control and phonology, which may be hypothesised to be control regions. However, at a less stringent threshold (voxel-level p-value of .01), two additional clusters are revealed, one in posterior fusiform gyrus, and one in the dmPFC spanning the midline (see Supplementary Figure 1), indicating that these regions may also reflect control demands in the phonology domain. Like the semantic network, the regions implicated in phonology may be divided on the basis of performing control versus representation processes. Specifically, the bilateral STG may underpin phonological representation, whilst IFG, inferior parietal cortex and posterior ITG may all contribute to phonological control.

**Table 8:** Phonological control (hard > easy phonology contrast)

| Cluster | Region of Activation | Z Score | Peak MNI Coordinate |  |  |
| --- | --- | --- | --- | --- | --- |
|  |  |  | x | y | z |
| 0.001 |  |  |  |  |  |
| 1 | Left inferior parietal lobule | 3.891 | -37 | -48 | 45 |
|  |  | 3.719 | -34 | -53 | 46 |
|  |  | 3.540 | -25 | -68 | 44 |
|  |  | 3.291 | -24 | -72 | 46 |
| 2 | Left inferior frontal gyrus/middle frontal gyrus | 3.891 | -54 | 20 | 25 |
|  |  | 3.719 | -53 | 30 | 22 |
|  |  | 3.432 | -46 | 18 | 22 |
| 0.01 |  |  |  |  |  |
| 1 | Left inferior frontal gyrus/middle frontal gyrus | 3.891 | -54 | 20 | 25 |
|  |  | 3.719 | -48 | 31 | 25 |
|  |  | 3.891 | -36 | -49 | 45 |
| 2 | Left inferior parietal lobule | 3.719 | -26 | -69 | 43 |
|  |  | 3.540 | -28 | -68 | 46 |
|  |  | 3.432 | -26 | -60 | 48 |
|  |  | 3.291 | -21 | -63 | 46 |
| 3 | Bilateral dorsomedial prefrontal cortex | 2.911 | 2 | 24 | 54 |
|  |  | 2.834 | 6 | 20 | 52 |
|  |  | 2.794 | 5 | 17 | 50 |
| 4 | Left fusiform gyrus | 2.697 | -44 | -60 | -14 |

### Separating the Language Network by Level of Domain-Specificity

Do these control regions respond selectively to specific subdomains of language, subserve language in general, or underpin all cognitive domains? Ventral IFG and posterior MTG were implicated in semantic control only, whilst dorsolateral prefrontal cortex, dorsal IFG (pars opercularis and triangularis), dmPFC and posterior ITG were implicated in both semantic and phonological control. These regions may reflect language-general or domain-general control regions. To help to distinguish these two possibilities, we examined the overlap with two measures of domain-general control: an ALE analysis of the working memory *n*-back task and a mask of the multiple demand network (from Fedorenko et al., 2013). A formal analysis of nonverbal working memory allows rigorous inclusion criterion without any effects of semantic or phonological stimuli, whereas an *a priori* MDN map may provide a more complete picture of domain-general control encompassing different executive functions.

The *n*-back working memory ALE analysis yielded a distributed bilateral network for domain-general control (Figure 4 & Table 9), highly similar to the mask from Fedorenko et al., ^45^ although lacking temporal and occipital involvement. There are three clusters in bilateral dorsolateral prefrontal cortex, extending into precentral and inferior frontal gyri, and two in left and right insula, with some involvement of pars orbitalis. Further clusters are located in the inferior parietal lobule, extending dorsally and medially into the superior parietal lobule, and the dmPFC, right cerebellum and left thalamus. Both these results and the *a priori* MDN map overlap substantially with the regions implicated in both phonological and semantic control, but not the regions implicated in semantic control alone. Thus, the language network may include some regions implicated specifically in semantic control, or the manipulation of meaningful representations (i.e., ventral IFG and posterior MTG), and some regions responsible for domain-general control (i.e., dmPFC, dorsal IFG, posterior ITG and inferior parietal cortex).

**Figure 4.**
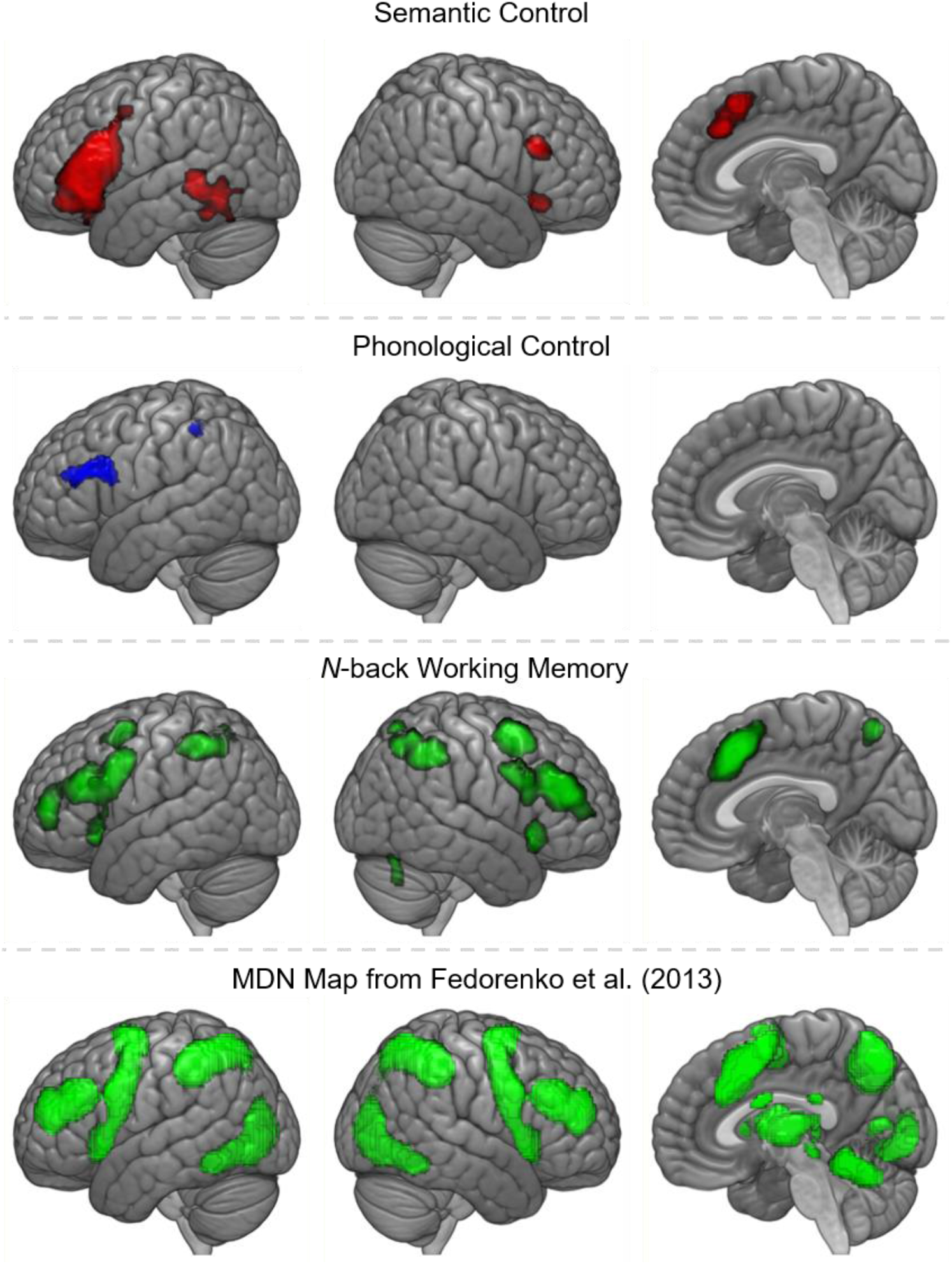
Top row: activation map of semantic control (red). Second row: map of phonological control regions (blue), represented by hard > easy phonology contrast. Third row: activation map for the *n*-back working memory domain. Bottom row: map of the multiple demand network from Fedorenko et al. (2013).

**Table 9:** N-back working memory activation likelihood

| Cluster | Region of Activation | Maximum<br>ALE Value | Z Score | Peak MNI Coordinate |  |  |
| --- | --- | --- | --- | --- | --- | --- |
|  |  |  |  | x | y | z |
| 1 | Left middle frontal<br>gyrus/precentral gyrus | 0.080 | 9.437 | -44 | 8 | 32 |
|  |  | 0.059 | 7.583 | -44 | 30 | 24 |
|  |  | 0.051 | 6.778 | -36 | 52 | 10 |
|  |  | 0.049 | 6.632 | -30 | 2 | 56 |
|  |  | 0.027 | 4.113 | -56 | 20 | 24 |
|  |  | 0.023 | 3.599 | -50 | 16 | 6 |
| 2 | Right middle frontal<br>gyrus/precentral gyrus | 0.078 | 9.253 | 44 | 42 | 24 |
|  |  | 0.071 | 8.623 | 46 | 34 | 26 |
|  |  | 0.044 | 6.068 | 52 | 12 | 30 |
| 3 | Right inferior parietal lobule | 0.107 | 11.62 | 44 | -46 | 44 |
|  |  | 0.064 | 8.003 | 34 | -58 | 48 |
|  |  | 0.044 | 6.111 | 16 | -66 | 58 |
| 4 | Bilateral dorsomedial<br>prefrontal cortex | 0.089 | 10.227 | 0 | 16 | 48 |
| 5 |  | 0.077 | 9.199 | -36 | -48 | 44 |
|  |  | 0.071 | 8.654 | -42 | -44 | 46 |
|  |  | 0.026 | 3.990 | -12 | -68 | 56 |
|  |  | 0.024 | 3.738 | -4 | -60 | 54 |
| 6 | Left inferior parietal lobule | 0.071 | 8.644 | 28 | 8 | 58 |
| 7 | Left insular/inferior frontal<br>gyrus | 0.092 | 10.423 | -32 | 22 | 0 |
| 8 | Left thalamus | 0.042 | 5.877 | -14 | 0 | 10 |
|  |  | 0.040 | 5.633 | -12 | -6 | 10 |
| 9 | Right insula/inferior frontal<br>gyrus | 0.090 | 10.290 | 34 | 24 | -2 |
| 10 | Right posterior cerebellum | 0.037 | 5.314 | 32 | -64 | -32 |
|  |  | 0.032 | 4.800 | 34 | -68 | -20 |

## DISCUSSION

A multi-dimensional approach is necessary to describe the underlying functional organisation of the language network. Formal analyses confirmed prior hypotheses that distinguishing semantic and phonological subdomains provides one key organisational principle. Yet this domain-based separation alone is not sufficient. Distinguishing the processes of representation and control provides additional, distinct indications of the functional roles of regions throughout the language network. Consistent with prior assessments, the semantic network comprises the bilateral ATL for representation, alongside control regions in the left IFG, bilateral dmPFC and left posterior lateral temporal lobe ^7,17,19,38^. The AG was also implicated in semantic representation, although there is ongoing debate as to whether this region truly contributes to semantic cognition overall, underpins a particular aspect of semantic cognition, or is identified solely based on confounding factors, such as difficulty-related deactivations ^26–28,51^. Phonological representation is supported by bilateral STG and left precentral gyrus, with control dependent on the left supramarginal and superior angular gyri, dorsal IFG and posterior lateral temporal cortex, as well as the dmPFC. This is highly consistent with prior assessments of the phonological network ^8,10–13,26^, yet provides additional evidence regarding the nature of the processing in these regions. Whilst the semantic and phonological representation regions are dissociable, there are both shared (left dorsal IFG and posterior inferior temporal cortex, bilateral dmPFC) and distinct (left ventral IFG and posterior MTG for semantic control only) control regions. These shared regions are all implicated in domain-general executive functions, forming part of the multiple demand network ^33,42,44^. Semantic control has its own distinctive neural correlates, yet effortful tasks in both semantic and phonological domains recruit additional domain-general frontal and posterior inferior temporal control regions. This highlights the importance of a third factor; the domain-specificity of the regions recruited for language tasks. The language network is multi-faceted and only by considering the interactions between subdomain, process and domain-specificity, can we understand the function of an individual region. The contribution of multiple dimensions to the function of each region is demonstrated in a proposed schematic for the organisation of the language network, presented in Figure 5, based on the present meta-analyses in the context of the broader literature. The remainder of the Discussion will first describe important methodological considerations for future work before delineating the key implications for existing theories of organisation within the language network.

**Figure 5.**
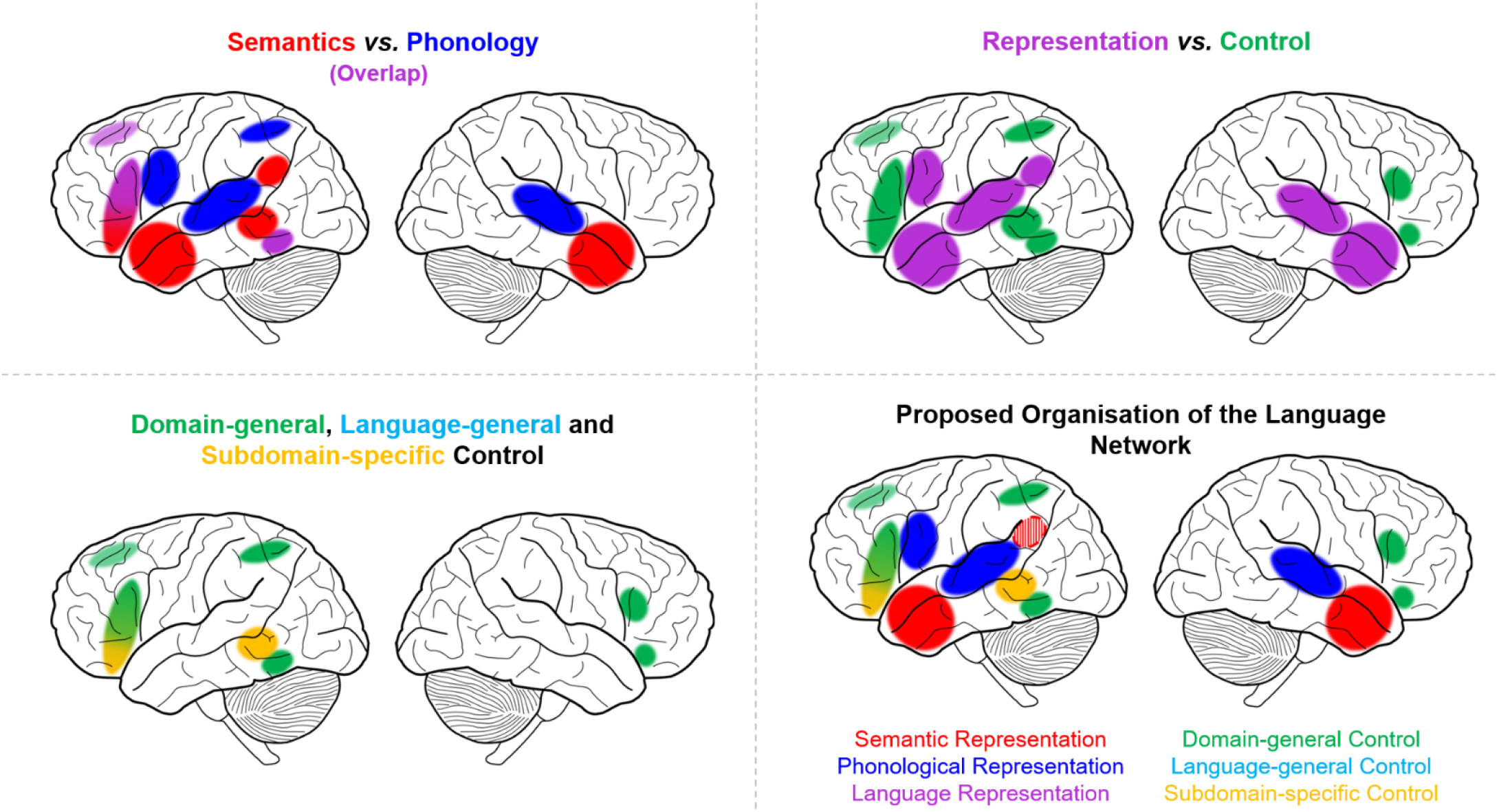
Schematic diagrams of the core organisational principles of the language network, based on the meta-analytic results. Top left: regions implicated in the semantic (red) and phonological (blue) subdomains with overlap shown in purple. Top right: regions implicated for semantic/phonological representation (purple) and control (green). Bottom left: domain-specificity of regions implicated in semantic and phonological control. Domain-general regions are shown in green and subdomain-specific regions in yellow (all of which reflect semantic control regions). Language-general regions would be displayed in blue, yet no regions were found to be shared between semantics and phonology without a more domain-general role. Bottom right: a proposed multi-dimensional organisation of the language network. The angular gyrus is shaded as its role in semantic representation is highly debated ^26–28^; see Discussion for more detail.

Activation likelihood estimation is a powerful meta-analytic tool. Visual analysis of the distribution of peaks is clearly insufficient to determine the regions involved in a task or compare two distributed networks. However, it is the difference between the two ALE analyses that may be more surprising. Despite claims of distinct distributions, the Vigneau et al. ^3,4^ datasets are not adequate to formally identify a strong separation between semantic and phonological processing. These datasets included 44 contrasts and 70 contrasts for semantics and phonology respectively, far higher than the recommended 17-20 contrasts minimum for a neuroimaging meta-analysis ^31,52^. Although both within and between domain results varied by dataset, researchers should be particularly cautious when using datasets of this size to perform contrasts between meta-analysis results, which may require substantially more data. Separating phonology and semantics benefited from the increase in power brought about by the addition of a decade and a half of additional functional neuroimaging research. Of course, the simple amount of data may not be the critical issue, but the increase in data quality across time. More recent papers generally reported larger samples of participants and made use of a wide range of improved fMRI techniques.

Many models of the language network focus on the separation of subdomains as the single organising principle ^53–56^. For example, the dual-stream model of language, which proposes a largely bilateral ventral stream for lexico-semantic access and a left-lateralised dorsal stream for mapping sound to meaning, broadly divides the network into phonological and semantic streams ^11,57–59^. The identification of differences in ATL and precentral gyrus aligns well with this dorsal-ventral division, as does the pattern of activation across hemispheres. Of course, such models do not attempt to capture the relationship between representation and control, or the overlap between language and domain-general regions. Yet it is the large amount of overlap between semantic and phonological processing that may be surprising from a dual-stream perspective, or on the basis of any unidimensional, process-based account. Divisions between other subdomains (e.g., syntactic processing, processing of sequences) may extend the current findings ^60,61^, yet the evidence for a strong behavioural, cognitive or neural separation between these processes are not as clear ^62^. Instead, a full consideration of function may require consideration of multiple dimensions focused on different kinds of information, including process and involvement of these regions outside language tasks. This may be understood in terms of the primary systems hypothesis; language processes are performed by combining the necessary domain-specific representation regions with the appropriate control regions ^63,64^.

Language and executive function are often considered behaviourally and neurally independent, due to the ‘special’ nature of language ^65,66^. Yet areas typically associated with language may have more domain-general roles and multiple lines of evidence suggest interaction between these processes. For instance, after cerebrovascular accident, executive function is an important predictor of aphasia therapy outcome ^67^ and there is increased reliance on the dmPFC for speech ^68,69^. A large body of neuropsychological ^34,36^, neuroimaging ^29,38^ and neurostimulation ^70,71^ work now supports the importance of control processes within the semantic subdomain (see ^5,7^ for a more detailed review). Our results indicate the need to examine the same distinction within the phonology subdomain. Only the superior temporal gyri and precentral gyrus were specifically implicated in phonological processing. All other regions implicated in phonology were also involved in domain-general control, including dorsal IFG, inferior parietal lobe, dmPFC and posterior ITG. Whilst language researchers may not expect temporal regions to be implicated in control (typically thought to rely on frontoparietal regions), the posterior ITG has received increasing recognition in the executive function literature, and is not simply related to the presentation of visual stimuli ^44,45,72^. This is hard to align with models of phonological processing that associate these regions with phonology-specific processes, such as orthographic-phonologic mapping or speech segmentation ^8,73^. It should be noted that posterior ITG could reflect language-specific control as it was not implicated specifically in nonverbal working memory but found only in MDN assessments which included some verbal stimuli. However, this region has been implicated in non-verbal executive function, such as task switching in both patients and neuroimaging ^74–76^; therefore, it may be more likely that the pITG is instead responsible for a domain-general function distinct from working memory, perhaps related to task-shifting or attention. What particular control processes might be critical for phonological processing? The term ‘phonological control’ has previously been associated with phonological working memory in the form of the articulatory loop (Baddeley et al., 1984; Baddeley & Hitch, 1994; Clark & Wagner, 2003). However, working memory alone cannot explain the full distribution of the current results. For instance, posterior ITG was not implicated in working memory. Moreover, regions across the multiple demand network are found to have a role across tasks requiring different forms of executive control ^42,44^. Thus, phonological control may require a range of executive functions as with semantic control, perhaps reflecting similar elements, such as selection between possible words, inhibition of alternatives and attention shifting.

Unlike phonological control, not all of the areas implicated in semantic control are responsible for domain-general control. Ventral inferior frontal and posterior middle temporal regions were not identified as part of the MDN, in keeping with prior research ^33,44^. Instead, these regions appear specialised for the control of meaningful stimuli. Indeed, a specific impairment of semantic control results from damage to these particular regions in semantic aphasia ^34,80^. Why, if language as a whole is not ‘special’, might semantic cognition recruit unique control regions? These regions are critical for the context-dependent access and manipulation of all meaningful items, including pictures, objects, faces and environmental sounds, as well as language ^5^. The control of such meaningful multimodal stimuli may heavily rely on particular control processes, such as competition selection and suppression ^7,32,37,81,82^. Alternatively, the process may be equivalent across control regions, yet the connections of these particular areas may be conducive to the application of control to meaningful stimuli, for example, due to the nature of their connections with the anterior temporal lobe hub. Indeed, it is not clear whether these distinctions ought to be viewed as distinct regions for semantic and domain-general control or graded changes within a larger complex ^38,44^. Semantic control regions lie adjacent to regions implicated in more domain-general control, including ventral versus dorsal IFG and posterior middle versus inferior temporal gyri. Further research should consider to what extent these reflect graded changes in function versus a sharp shift between distinct functional regions.

## METHODS

Activation likelihood estimation (ALE) meta-analyses were conducted for the semantic and phonology domains, independently for Vigneau et al.’s data and the full datasets, which were compared to one another using formal contrast analyses. Subsequently, formal ALE analyses were conducted for semantic control, phonological control, and the *n*-back working memory task.

### Inclusion and Exclusion Criteria

All meta-analyses included only peer-reviewed English language articles, describing task-based fMRI and PET studies that reported peak coordinates of a univariate contrast in standard space (Talairach or MNI) and focused on a young healthy adult sample (below 40 years old). Contrasts were excluded if they focused on patients, clinical trials or individual differences (e.g., age, gender, native language). Finally, any contrasts that overlapped with the other domains – e.g., phonological working memory tasks – were excluded, to allow for meaningful comparisons to be made between domains. Within each paper, wherever multiple task contrasts were reported for the same participant sample, all the peak activation coordinates were analysed as a single contrast, following the recommendation from Mueller et al. ^31^.

For each domain, studies were sourced from one or more existing published meta-analyses, providing appropriate inclusion and exclusion criteria in keeping with the accepted definition of that particular domain. In addition, each contrast had to meet the general inclusion criteria provided above. For semantics and phonology, formal analyses were conducted employing the data from Vigneau et al. ^3,4^ including both the left and right hemisphere peaks. For these datasets, the original inclusion and exclusion criteria were kept; in total, this included 44 experiments for phonology, and 70 experiments for semantics.

The studies for the semantics domain (Supplementary Table 1) were taken from a meta-analysis by Jackson ^38^, which included 272 verbal and nonverbal contrasts that specifically compared a semantic condition with a non-semantic (or less semantic) condition. This included contrasts that compared semantic > less semantic tasks, semantic > non-semantic tasks and meaningful/known > non-meaningful/unknown stimuli. Comparison of high > low familiarity or imageability were excluded, as were studies that used rest or fixation as a baseline, as it has been shown that subtraction of low-level baselines is likely to remove semantic activations due to the high level of semantic processing during rest ^83^. To determine which semantic regions were involved in control, an additional semantic control assessment was included from Jackson ^38^. This included 96 experiments contrasting more controlled/harder semantics over less controlled/easier semantics, and included tasks that manipulated variables such as homonym ambiguity, interference from competitors and strength of semantic association (Supplementary Table 3). As Jackson ^38^ employed the same inclusion and exclusion criteria as the present study, all contrasts were included.

For the phonological domain, studies were sourced from Vigneau et al. ^3,4^ and Humphreys and Lambon Ralph ^26^ that reported peaks for phonology > non-phonological or less phonological tasks. Vigneau et al. ^3,4^ included 44 studies (86 contrasts) across a wide range of phonological tasks, including repetition, listening, reading or attending to syllables, letters, pseudo-words and words, judging rhyme, and phonological working memory. These data were supplemented with contrasts from 27 additional papers from Humphreys and Lambon Ralph ^26^, primarily contrasts that compared phonological tasks > semantic or orthographical tasks and reported peaks in the parietal lobe. Six contrasts which explicitly investigated working memory were excluded, in order to eliminate conceptual overlap with the *n*-back working memory ALE analysis. The final dataset included 64 experiments with a total of 870 foci (Supplementary Table 2). In order to assess phonological control, the dataset for the phonological domain was divided into hard versus easy tasks, for subsequent formal ALE contrast. Tasks that were passive or simply required a straightforward stimulus-response mapping, e.g., repetition or listening, were classed as easier, while those that involved decision-making, such as judgement of rhyme or syllable number, were classified as harder. In total, 20 experiments with 247 foci in total were included for easy phonology, and 44 experiments with 623 foci in total for hard phonology.

The studies for the *n*-back domain were sourced from Wang et al. ^84^. Wang et al. ^84^ included 96 published fMRI studies of healthy adults completing verbal, nonverbal, spatial and nonspatial variants of the n-back working memory task, with an *n* between 0 and 3. For the purposes of the present study, only those contrasts that reported a higher > lower *n*-value and met the general inclusion criteria were included. A total of 11 contrasts that contained meaningful (e.g., words or faces) or phonological stimuli were also excluded due to the potential overlap with semantic and phonological processing. The final dataset for this domain included 66 experiments with a total of 1216 foci (Supplementary Table 4).

### Activation Likelihood Estimation (ALE) Analyses

Meta-analyses were performed using activation likelihood estimation (ALE) performed in GingerALE version 3.0.2 (https://brainmap.org/ale/; ^85–88^). All analyses were performed in MNI152 space; before running the analyses, all peaks given in Talairach space were converted to MNI space within GingerALE. ALE is a meta-analytic technique that maps the statistically significant convergence of activation probabilities between experiments considered to reflect similar processes. This is achieved by modelling all foci for each experiment as Gaussian probability distributions, with a full width at half maximum (FWHM) for each Gaussian determined by the sample size of the study (i.e., larger samples result in less uncertainty of the peak’s location) to produce a modelled activation map for each experiment included in the analysis. These maps are then merged and ALE scores computed on a voxel-by-voxel basis, with each ALE score representing the probability of an activation being present at that given voxel. For all meta-analyses, ALE scores were thresholded with a voxel-wise p-value of .001. Cluster-level family-wise error correction at a p-value of .001, with 10000 permutations, was then applied to determine minimum significant cluster size and remove non-significant clusters. Formal ALE meta-analyses were conducted for the semantics, phonology and *n*-back domains, and for the Vigneau semantics and phonology datasets. The activation maps for the semantics and phonology domains were also directly contrasted using GingerALE, creating pairwise thresholded conjunction and subtraction ALE images using a p-value of .001 with 10000 permutations and a minimum cluster volume of 20mm^3^.

## Acknowledgements

This work was supported by a Biotechnology and Biological Sciences Research Council studentship to V.J.H, a British Academy Postdoctoral Fellowship awarded to R.L.J. (no. pf170068), a programme grant to M.A.L.R. from the Medical Research Council (grant no. MR/R023883/1), an Advanced Grant from the European Research Council to M.A.L.R. (GAP: 670428) and Medical Research Council intramural funding (no. MC_UU_00005/18).

## REFERENCES

1. Démonet, J. F. et al. THE ANATOMY OF PHONOLOGICAL AND SEMANTIC PROCESSING IN NORMAL SUBJECTS. Brain 115, 1753–1768 (1992).

2. Geschwind, N. The Organization of Language and the Brain. Science 170, 940–944 (1970).

3. Vigneau, M. et al. Meta-analyzing left hemisphere language areas: Phonology, semantics, and sentence processing. NeuroImage 30, 1414–1432 (2006).

4. Vigneau, M. et al. What is right-hemisphere contribution to phonological, lexico-semantic, and sentence processing?: Insights from a meta-analysis. NeuroImage 54, 577–593 (2011).

5. Lambon Ralph, M. A. L., Jefferies, E., Patterson, K. & Rogers, T. T. The neural and computational bases of semantic cognition. Nat. Rev. Neurosci. 18, 42–55 (2017).

6. Rogers, T. T. et al. Structure and Deterioration of Semantic Memory: A Neuropsychological and Computational Investigation. Psychol. Rev. 111, 205–235 (2004).

7. Jefferies, E. The neural basis of semantic cognition: Converging evidence from neuropsychology, neuroimaging and TMS. Cortex 49, 611–625 (2013).

8. Poldrack, R. A. et al. Functional Specialization for Semantic and Phonological Processing in the Left Inferior Prefrontal Cortex. NeuroImage 10, 15–35 (1999).

9. Wagner, R. K. & Torgesen, J. K. The nature of phonological processing and its causal role in the acquisition of reading skills. Psychol. Bull. 101, 192–212 (1987).

10. Buchsbaum, B. R., Hickok, G. & Humphries, C. Role of left posterior superior temporal gyrus in phonological processing for speech perception and production. Cogn. Sci. 25, 663–678 (2001).

11. Hickok, G. & Poeppel, D. The cortical organization of speech processing. Nat. Rev. Neurosci. 8, 393–402 (2007).

12. Buchsbaum, B. R. et al. Conduction aphasia, sensory-motor integration, and phonological short-term memory – An aggregate analysis of lesion and fMRI data. Brain Lang. 119, 119–128 (2011).

13. Graves, W. W., Grabowski, T. J., Mehta, S. & Gupta, P. The Left Posterior Superior Temporal Gyrus Participates Specifically in Accessing Lexical Phonology. J. Cogn. Neurosci. 20, 1698–1710 (2008).

14. Démonet, J. F., Fiez, J. A., Paulesu, E., Petersen, S. E. & Zatorre, R. J. PET Studies of Phonological Processing: A Critical Reply to Poeppel. Brain Lang. 55, 352–379 (1996).

15. Zatorre, R. J., Meyer, E., Gjedde, A. & Evans, A. C. PET Studies of Phonetic Processing of Speech: Review, Replication, and Reanalysis. Cereb. Cortex 6, 21–30 (1996).

16. Patterson, K., Nestor, P. J. & Rogers, T. T. Where do you know what you know? The representation of semantic knowledge in the human brain. Nat. Rev. Neurosci. 8, 976–987 (2007).

17. Patterson, K. & Lambon Ralph, M. A. Chapter 61 - The Hub-and-Spoke Hypothesis of Semantic Memory. in Neurobiology of Language (eds. Hickok, G. & Small, S. L.) 765–775 (Academic Press, 2016). doi:10.1016/B978-0-12-407794-2.00061-4.

18. Binder, J. R. & Desai, R. H. The neurobiology of semantic memory. Trends Cogn. Sci. 15, 527–536 (2011).

19. Binder, J. R., Desai, R. H., Graves, W. W. & Conant, L. L. Where Is the Semantic System? A Critical Review and Meta-Analysis of 120 Functional Neuroimaging Studies. Cereb. Cortex 19, 2767–2796 (2009).

20. Fiez, J. A. Phonology, semantics, and the role of the left inferior prefrontal cortex. Hum. Brain Mapp. 5, 79–83 (1997).

21. Aziz-Zadeh, L., Cattaneo, L., Rochat, M. & Rizzolatti, G. Covert Speech Arrest Induced by rTMS over Both Motor and Nonmotor Left Hemisphere Frontal Sites. J. Cogn. Neurosci. 17, 928–938 (2005).

22. Nixon, P., Lazarova, J., Hodinott-Hill, I., Gough, P. & Passingham, R. The Inferior Frontal Gyrus and Phonological Processing: An Investigation using rTMS. J. Cogn. Neurosci. 16, 289–300 (2004).

23. Devlin, J. T., Matthews, P. M. & Rushworth, M. F. S. Semantic Processing in the Left Inferior Prefrontal Cortex: A Combined Functional Magnetic Resonance Imaging and Transcranial Magnetic Stimulation Study. J. Cogn. Neurosci. 15, 71–84 (2003).

24. Köhler, S., Paus, T., Buckner, R. L. & Milner, B. Effects of Left Inferior Prefrontal Stimulation on Episodic Memory Formation: A Two-Stage fMRI—rTMS Study. J. Cogn. Neurosci. 16, 178–188 (2004).

25. Devlin, J. T. & Watkins, K. E. Stimulating language: insights from TMS. Brain 130, 610–622 (2007).

26. Humphreys, G. F. & Lambon Ralph, M. A. Fusion and Fission of Cognitive Functions in the Human Parietal Cortex. Cereb. Cortex 25, 3547–3560 (2015).

27. Cabeza, R., Ciaramelli, E. & Moscovitch, M. Cognitive contributions of the ventral parietal cortex: an integrative theoretical account. Trends Cogn. Sci. 16, 338–352 (2012).

28. Seghier, M. L. The Angular Gyrus: Multiple Functions and Multiple Subdivisions. The Neuroscientist 19, 43–61 (2013).

29. Noonan, K. A., Jefferies, E., Visser, M. & Lambon Ralph, M. A. Going beyond Inferior Prefrontal Involvement in Semantic Control: Evidence for the Additional Contribution of Dorsal Angular Gyrus and Posterior Middle Temporal Cortex. J. Cogn. Neurosci. 25, 1824–1850 (2013).

30. Bzdok, D. et al. Left inferior parietal lobe engagement in social cognition and language. Neurosci. Biobehav. Rev. 68, 319–334 (2016).

31. Müller, V. I. et al. Ten simple rules for neuroimaging meta-analysis. Neurosci. Biobehav. Rev. 84, 151–161 (2018).

32. Badre, D., Poldrack, R. A., Paré-Blagoev, E. J., Insler, R. Z. & Wagner, A. D. Dissociable Controlled Retrieval and Generalized Selection Mechanisms in Ventrolateral Prefrontal Cortex. Neuron 47, 907–918 (2005).

33. Duncan, J. The multiple-demand (MD) system of the primate brain: mental programs for intelligent behaviour. Trends Cogn. Sci. 14, 172–179 (2010).

34. Jefferies, E. & Lambon Ralph, M. A. Semantic impairment in stroke aphasia versus semantic dementia: a case-series comparison. Brain 129, 2132–2147 (2006).

35. Jackson, R. L., Bajada, C. J., Lambon Ralph, M. A. & Cloutman, L. L. The Graded Change in Connectivity across the Ventromedial Prefrontal Cortex Reveals Distinct Subregions. Cereb. Cortex (2019) doi:10.1093/cercor/bhz079.

36. Rogers, T. T., Patterson, K., Jefferies, E. & Lambon Ralph, M. A. Disorders of representation and control in semantic cognition: Effects of familiarity, typicality, and specificity. Neuropsychologia 76, 220–239 (2015).

37. Davey, J. et al. Exploring the role of the posterior middle temporal gyrus in semantic cognition: Integration of anterior temporal lobe with executive processes. NeuroImage 137, 165–177 (2016).

38. Jackson, R. L. The neural correlates of semantic control revisited. NeuroImage 224, 117444 (2020).

39. Gold, B. T., Balota, D. A., Kirchhoff, B. A. & Buckner, R. L. Common and Dissociable Activation Patterns Associated with Controlled Semantic and Phonological Processing: Evidence from fMRI Adaptation. Cereb. Cortex 15, 1438–1450 (2005).

40. Gold, B. T. & Buckner, R. L. Common Prefrontal Regions Coactivate with Dissociable Posterior Regions during Controlled Semantic and Phonological Tasks. Neuron 35, 803–812 (2002).

41. Assem, M., Glasser, M. F., Essen, D. C. V. & Duncan, J. A Domain-general Cognitive Core defined in Multimodally Parcellated Human Cortex. bioRxiv 517599 (2019) doi:10.1101/517599.

42. Camilleri, J. A. et al. Definition and characterization of an extended multiple-demand network. NeuroImage 165, 138–147 (2018).

43. Pollack, C. & Ashby, N. C. Where arithmetic and phonology meet: The meta-analytic convergence of arithmetic and phonological processing in the brain. Dev. Cogn. Neurosci. 30, 251–264 (2018).

44. Assem, M., Glasser, M. F., Van Essen, D. C. & Duncan, J. A Domain-General Cognitive Core Defined in Multimodally Parcellated Human Cortex. Cereb. Cortex 30, 4361–4380 (2020).

45. Fedorenko, E., Duncan, J. & Kanwisher, N. Broad domain generality in focal regions of frontal and parietal cortex. Proc. Natl. Acad. Sci. 110, 16616–16621 (2013).

46. Barth, M., Breuer, F., Koopmans, P. J., Norris, D. G. & Poser, B. A. Simultaneous multislice (SMS) imaging techniques. Magn. Reson. Med. 75, 63–81 (2016).

47. Feinberg, D. A. & Setsompop, K. Ultra-fast MRI of the human brain with simultaneous multi-slice imaging. J. Magn. Reson. 229, 90–100 (2013).

48. Visser, M., Embleton, K. V., Jefferies, E., Parker, G. J. & Ralph, M. A. L. The inferior, anterior temporal lobes and semantic memory clarified: Novel evidence from distortion-corrected fMRI. Neuropsychologia 48, 1689–1696 (2010).

49. Halai, A. D., Welbourne, S. R., Embleton, K. & Parkes, L. M. A comparison of dual gradient-echo and spin-echo fMRI of the inferior temporal lobe. Hum. Brain Mapp. 35, 4118–4128 (2014).

50. Poser, B. A., Versluis, M. J., Hoogduin, J. M. & Norris, D. G. BOLD contrast sensitivity enhancement and artifact reduction with multiecho EPI: Parallel-acquired inhomogeneity-desensitized fMRI. Magn. Reson. Med. 55, 1227–1235 (2006).

51. Humphreys, G., Ralph, M. L. & Simons, J. A Unifying Account of Angular Gyrus Contributions to Episodic and Semantic Cognition. https://psyarxiv.com/r2deu/ (2020) doi:10.31234/osf.io/r2deu.

52. Eickhoff, S. B. et al. Behavior, sensitivity, and power of activation likelihood estimation characterized by massive empirical simulation. NeuroImage 137, 70–85 (2016).

53. Catani, M., Jones, D. K. & Ffytche, D. H. Perisylvian language networks of the human brain. Ann. Neurol. 57, 8–16 (2005).

54. Friederici, A. D. Pathways to language: fiber tracts in the human brain. Trends Cogn. Sci. 13, 175–181 (2009).

55. Price, C. J. The anatomy of language: contributions from functional neuroimaging. J. Anat. 197, 335–359 (2000).

56. Price, C. J. A review and synthesis of the first 20years of PET and fMRI studies of heard speech, spoken language and reading. NeuroImage 62, 816–847 (2012).

57. Hickok, G. & Poeppel, D. Dorsal and ventral streams: a framework for understanding aspects of the functional anatomy of language. Cognition 92, 67–99 (2004).

58. Saur, D. et al. Ventral and dorsal pathways for language. Proc. Natl. Acad. Sci. 105, 18035–18040 (2008).

59. Ueno, T. & Lambon Ralph, M. A. The roles of the “ventral” semantic and “dorsal” pathways in conduite d’approche: a neuroanatomically-constrained computational modeling investigation. Front. Hum. Neurosci. 7, (2013).

60. Bornkessel-Schlesewsky, I., Schlesewsky, M., Small, S. L. & Rauschecker, J. P. Neurobiological roots of language in primate audition: common computational properties. Trends Cogn. Sci. 19, 142–150 (2015).

61. Matchin, W. & Hickok, G. The Cortical Organization of Syntax. Cereb. Cortex 30, 1481–1498 (2020).

62. Friederici, A. D. Towards a neural basis of auditory sentence processing. Trends Cogn. Sci. 6, 78–84 (2002).

63. Patterson, K. & Lambon Ralph, M. A. Selective disorders of reading? Curr. Opin. Neurobiol. 9, 235–239 (1999).

64. Woollams, A. M. Connectionist neuropsychology: uncovering ultimate causes of acquired dyslexia. Philos. Trans. R. Soc. B Biol. Sci. (2014) doi:10.1098/rstb.2012.0398.

65. Fedorenko, E., Duncan, J. & Kanwisher, N. Language-Selective and Domain-General Regions Lie Side by Side within Broca’s Area. Curr. Biol. 22, 2059–2062 (2012).

66. Fedorenko, E. & Blank, I. A. Broca’s Area Is Not a Natural Kind. Trends Cogn. Sci. 24, 270–284 (2020).

67. Lambon Ralph, M. A., Snell, C., Fillingham, J. K., Conroy, P. & Sage, K. Predicting the outcome of anomia therapy for people with aphasia post CVA: Both language and cognitive status are key predictors. Neuropsychol. Rehabil. 20, 289–305 (2010).

68. Geranmayeh, F., Chau, T. W., Wise, R. J. S., Leech, R. & Hampshire, A. Domain-general subregions of the medial prefrontal cortex contribute to recovery of language after stroke. Brain 140, 1947–1958 (2017).

69. Sliwinska, M. W. et al. Stimulating Multiple-Demand Cortex Enhances Vocabulary Learning. J. Neurosci. 37, 7606–7618 (2017).

70. Whitney, C., Kirk, M., O’Sullivan, J., Lambon Ralph, M. A. & Jefferies, E. Executive Semantic Processing Is Underpinned by a Large-scale Neural Network: Revealing the Contribution of Left Prefrontal, Posterior Temporal, and Parietal Cortex to Controlled Retrieval and Selection Using TMS. J. Cogn. Neurosci. 24, 133–147 (2011).

71. Whitney, C., Kirk, M., O’Sullivan, J., Lambon Ralph, M. A. & Jefferies, E. The Neural Organization of Semantic Control: TMS Evidence for a Distributed Network in Left Inferior Frontal and Posterior Middle Temporal Gyrus. Cereb. Cortex 21, 1066–1075 (2011).

72. Woolgar, A. & Zopf, R. Multisensory coding in the multiple-demand regions: vibrotactile task information is coded in frontoparietal cortex. J. Neurophysiol. 118, 703–716 (2017).

73. Burton, M. W. The role of inferior frontal cortex in phonological processing. Cogn. Sci. 25, 695–709 (2001).

74. Buchsbaum, B. R., Greer, S., Chang, W.-L. & Berman, K. F. Meta-analysis of neuroimaging studies of the Wisconsin Card-Sorting task and component processes. Hum. Brain Mapp. 25, 35–45 (2005).

75. Kim, C., Cilles, S. E., Johnson, N. F. & Gold, B. T. Domain general and domain preferential brain regions associated with different types of task switching: A Meta-Analysis. Hum. Brain Mapp. 33, 130–142 (2012).

76. Schumacher, R., Halai, A. D. & Lambon Ralph, M. A. Assessing and mapping language, attention and executive multidimensional deficits in stroke aphasia. Brain 142, 3202–3216 (2019).

77. Baddeley, A. D., Lewis, V. & Vallar, G. Exploring the Articulatory Loop. Q. J. Exp. Psychol. Sect. A 36, 233–252 (1984).

78. Baddeley, A. D. & Hitch, G. J. Developments in the concept of working memory. Neuropsychology 8, 485–493 (1994).

79. Clark, D. & Wagner, A. D. Assembling and encoding word representations: fMRI subsequent memory effects implicate a role for phonological control. Neuropsychologia 41, 304–317 (2003).

80. Berthier, M. L. Unexpected brain-language relationships in aphasia: Evidence from transcortical sensory aphasia associated with frontal lobe lesions. Aphasiology 15, 99–130 (2001).

81. Corbett, F., Jefferies, E. & Lambon Ralph, M. A. Deregulated Semantic Cognition Follows Prefrontal and Temporo-parietal Damage: Evidence from the Impact of Task Constraint on Nonverbal Object Use. J. Cogn. Neurosci. 23, 1125–1135 (2010).

82. Corbett, F., Jefferies, E., Ehsan, S. & Lambon Ralph, M. A. Different impairments of semantic cognition in semantic dementia and semantic aphasia: evidence from the non-verbal domain. Brain 132, 2593–2608 (2009).

83. Visser, M., Jefferies, E. & Lambon Ralph, M. A. Semantic Processing in the Anterior Temporal Lobes: A Meta-analysis of the Functional Neuroimaging Literature. J. Cogn. Neurosci. 22, 1083–1094 (2009).

84. Wang, H. et al. A coordinate-based meta-analysis of the n-back working memory paradigm using activation likelihood estimation. Brain Cogn. 132, 1–12 (2019).

85. Eickhoff, S. B. et al. Coordinate-based activation likelihood estimation meta-analysis of neuroimaging data: A random-effects approach based on empirical estimates of spatial uncertainty. Hum. Brain Mapp. 30, 2907–2926 (2009).

86. Eickhoff, S. B., Bzdok, D., Laird, A. R., Kurth, F. & Fox, P. T. Activation likelihood estimation meta-analysis revisited. NeuroImage 59, 2349–2361 (2012).

87. Eickhoff, S. B., Laird, A. R., Fox, P. M., Lancaster, J. L. & Fox, P. T. Implementation errors in the GingerALE Software: Description and recommendations. Hum. Brain Mapp. 38, 7–11 (2017).

88. Turkeltaub, P. E., Eden, G. F., Jones, K. M. & Zeffiro, T. A. Meta-Analysis of the Functional Neuroanatomy of Single-Word Reading: Method and Validation. NeuroImage 16, 765–780 (2002).

