## Supplemental Information for "Multiple dimensions underlying the functional organisation of the language network"

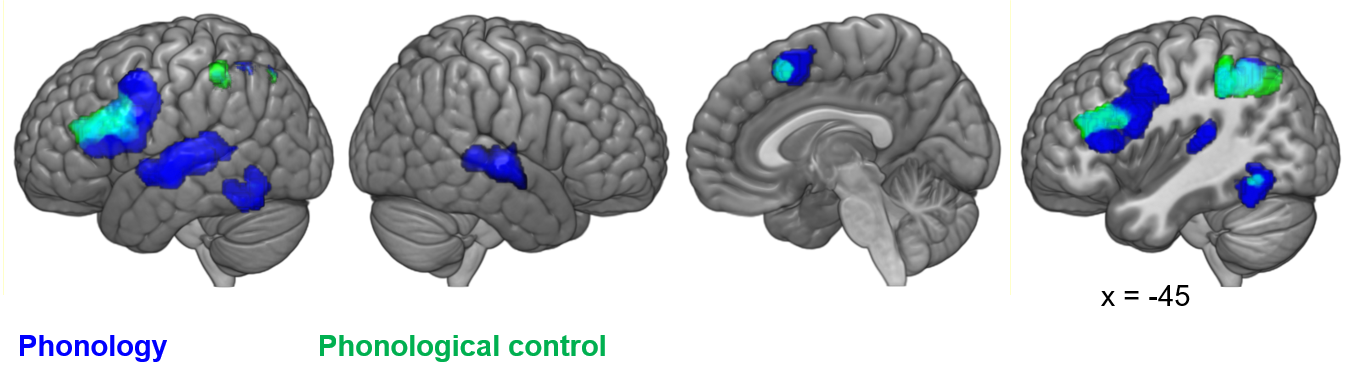

Supplementary Figure 1. Activation map for full phonological domain (blue) overlaid with phonological control, represented by hard > easy phonology formal ALE contrast (green, overlap in cyan) at a less stringent voxel-level threshold (p-value of .01).

Supplementary Table 1. Data included in the semantics meta-analysis.

| **Author(s)** | **Year** | **DOI** | **Contrast included** | **N** | **MNI coordinates** | | |
| --- | --- | --- | --- | --- | --- | --- | --- |
|  |  |  |  |  | **X** | **Y** | **Z** |
| Booth et al. | 2006 | 10.1016/j.brainres.2005.11.097 | Semantic > phonological judgement | 13 | -39 | -57 | 18 |
|  |  |  |  |  | -24 | -3 | 48 |
|  |  |  |  |  | -36 | 42 | -21 |
| Demonet et al. | 1992 | [10.1093/brain/115.6.1753](https://doi.org/10.1093/brain/115.6.1753) | Words > phonemes | 9 | -48 | -55 | 42 |
|  |  |  |  |  | -50 | -55 | 38 |
|  |  |  |  |  | -50 | -55 | 33 |
|  |  |  |  |  | -20 | 38 | 41 |
|  |  |  |  |  | -20 | 37 | 37 |
|  |  |  |  |  | -18 | 39 | 32 |
|  |  |  |  |  | -1 | -60 | 28 |
|  |  |  |  |  | -1 | -60 | 24 |
|  |  |  |  |  | -42 | -41 | -18 |
| Devlin et al. | 2003 | [10.1162/089892903321107837](https://doi.org/10.1162/089892903321107837) | Semantic > phonological judgement | 12 | -10 | 52 | -8 |
|  |  |  |  |  | -14 | 44 | -8 |
|  |  |  |  |  | -20 | 30 | 48 |
|  |  |  |  |  | -42 | -66 | 28 |
|  |  |  |  |  | -4 | -56 | 28 |
| Fletcher et al. | 1995 | [10.1016/0010-0277(95)00692-R](https://doi.org/10.1016/0010-0277(95)00692-R) | Social story > sentences | 6 | -12 | 50 | 35 |
|  |  |  |  |  | -47 | 21 | -24 |
|  |  |  |  |  | 48 | 19 | -25 |
|  |  |  |  |  | 6 | -65 | 15 |
|  |  |  |  |  | -46 | -58 | 24 |
| Fletcher et al. | 1995 | [10.1016/0010-0277(95)00692-R](https://doi.org/10.1016/0010-0277(95)00692-R) | Physical story > sentences | 6 | -42 | 17 | -19 |
|  |  |  |  |  | 48 | 17 | -25 |
|  |  |  |  |  | -1 | -64 | 11 |
|  |  |  |  |  | -44 | -65 | 21 |
| Gitelman et al. | 2005 | 10.1016/j.neuroimage.2005.03.014 | Semantic > control task | 14 | -15 | -36 | 66 |
|  |  |  |  |  | -12 | 30 | 42 |
|  |  |  |  |  | -24 | 12 | 39 |
|  |  |  |  |  | -24 | 21 | 36 |
|  |  |  |  |  | -51 | 27 | -3 |
|  |  |  |  |  | -48 | 15 | 9 |
|  |  |  |  |  | -54 | 27 | 24 |
|  |  |  |  |  | -9 | 15 | 60 |
|  |  |  |  |  | -39 | -54 | 24 |
|  |  |  |  |  | -48 | 12 | -15 |
|  |  |  |  |  | -42 | 0 | -42 |
|  |  |  |  |  | -57 | -45 | 3 |
|  |  |  |  |  | 48 | -15 | -18 |
|  |  |  |  |  | 51 | -9 | -9 |
|  |  |  |  |  | -51 | -15 | -9 |
|  |  |  |  |  | 24 | -72 | -3 |
|  |  |  |  |  | 15 | -87 | 21 |
|  |  |  |  |  | -15 | -45 | -36 |
|  |  |  |  |  | 6 | -54 | -33 |
|  |  |  |  |  | 27 | -78 | -42 |
|  |  |  |  |  | 15 | -81 | -36 |
| Gourovitch et al. | 2000 | [10.1037/0894-4105.14.3.353](https://psycnet.apa.org/doi/10.1037/0894-4105.14.3.353) | Semantic generation > phonology | 18 | -1 | 69 | 2 |
|  |  |  |  |  | -25 | -15 | -21 |
|  |  |  |  |  | -1 | 35 | -17 |
|  |  |  |  |  | -8 | 35 | -17 |
|  |  |  |  |  | -57 | -24 | -24 |
|  |  |  |  |  | 47 | -65 | 15 |
| Homae et al. | 2003 | [10.1016/S1053-8119(03)00272-6](https://doi.org/10.1016/S1053-8119(03)00272-6) | Sentences > jumbled words | 10 | -51 | 27 | -6 |
|  |  |  |  |  | -45 | 15 | 48 |
|  |  |  |  |  | -63 | -36 | -6 |
|  |  |  |  |  | 63 | -36 | -3 |
|  |  |  |  |  | -57 | -54 | 27 |
| Homae et al. | 2003 | [10.1016/S1053-8119(03)00272-6](https://doi.org/10.1016/S1053-8119(03)00272-6) | Sentences > jumbled words | 10 | -45 | 24 | -9 |
|  |  |  |  |  | -57 | 24 | 12 |
|  |  |  |  |  | -42 | 24 | 48 |
|  |  |  |  |  | -60 | -54 | 12 |
|  |  |  |  |  | 66 | -36 | -3 |
|  |  |  |  |  | -48 | -54 | 30 |
| Kotz et al. | 2002 | [10.1006/nimg.2002.1316](https://doi.org/10.1006/nimg.2002.1316) | Words > pseudowords | 13 | -24 | 15 | -5 |
|  |  |  |  |  | -33 | 9 | 5 |
|  |  |  |  |  | -14 | 14 | 0 |
|  |  |  |  |  | -46 | -67 | 23 |
|  |  |  |  |  | -56 | -44 | 37 |
|  |  |  |  |  | -37 | -61 | 38 |
|  |  |  |  |  | 54 | -50 | 9 |
| Kuperberg et al. | 2000 | 10.1162/089892900562138 | Sentences > words | 4 | -42 | -44 | -7 |
|  |  |  |  |  | -33 | -45 | 10 |
|  |  |  |  |  | 51 | -25 | 1 |
|  |  |  |  |  | -45 | -53 | 24 |
|  |  |  |  |  | 39 | 13 | -9 |
|  |  |  |  |  | -20 | -82 | -4 |
|  |  |  |  |  | -11 | -53 | 23 |
|  |  |  |  |  | 22 | 18 | -21 |
|  |  |  |  |  | -2 | -71 | -5 |
|  |  |  |  |  | -12 | -73 | -12 |
|  |  |  |  |  | -6 | 19 | -9 |
| Mechelli et al. | 2007 | 10.1002/hbm.20272 | Semantic > phonology related | 20 | -66 | -38 | -8 |
|  |  |  |  |  | -56 | -24 | -10 |
|  |  |  |  |  | -32 | -72 | 44 |
|  |  |  |  |  | -58 | -52 | 40 |
|  |  |  |  |  | 2 | 30 | 40 |
|  |  |  |  |  | -6 | 18 | 44 |
|  |  |  |  |  | -46 | 24 | -14 |
|  |  |  |  |  | -52 | 38 | -6 |
| Mummery et al. | 1998 | 10.1162/089892998563059 | Semantic > phonological decision | 10 | -48 | -72 | 35 |
|  |  |  |  |  | -59 | -25 | -8 |
|  |  |  |  |  | -31 | -32 | -19 |
|  |  |  |  |  | -47 | -24 | -19 |
|  |  |  |  |  | -5 | 66 | 18 |
|  |  |  |  |  | -14 | 58 | -15 |
|  |  |  |  |  | -34 | 19 | -24 |
| Nichelli et al. | 1995 | 10.1097/00001756-199511270-00010 | Semantic > orthographic decision | 9 | -31 | -58 | 47 |
|  |  |  |  |  | -22 | 25 | 43 |
|  |  |  |  |  | 9 | 56 | -19 |
| Price et al. | 1997 | 10.1162/jocn.1997.9.6.727. | Semantic > phonological decision | 6 | -34 | -5 | -26 |
|  |  |  |  |  | -52 | -69 | 21 |
|  |  |  |  |  | -3 | 9 | 4 |
| Raposo et al. | 2006 | [10.1016/j.neuropsychologia.2006.05.017](https://doi.org/10.1016/j.neuropsychologia.2006.05.017) | Unrelated > repeated stimuli | 15 | 10 | -51 | -20 |
|  |  |  |  |  | 23 | -72 | -23 |
|  |  |  |  |  | 29 | -64 | -26 |
|  |  |  |  |  | 29 | -24 | 8 |
|  |  |  |  |  | 40 | -39 | 3 |
|  |  |  |  |  | 34 | -57 | 14 |
|  |  |  |  |  | -42 | 5 | 27 |
|  |  |  |  |  | -51 | 24 | 12 |
|  |  |  |  |  | -53 | 15 | 17 |
|  |  |  |  |  | -3 | -32 | -4 |
|  |  |  |  |  | -10 | -43 | -11 |
|  |  |  |  |  | -33 | -38 | -2 |
| Roskies et al. | 2001 | [10.1162/08989290152541485](https://doi.org/10.1162/08989290152541485) | Semantic > phonological decision | 20 | 0 | -27 | -22 |
|  |  |  |  |  | -11 | 69 | -9 |
|  |  |  |  |  | -13 | 55 | 24 |
|  |  |  |  |  | -23 | 26 | 36 |
|  |  |  |  |  | -26 | 3 | 29 |
|  |  |  |  |  | -26 | -21 | -2 |
|  |  |  |  |  | -37 | 26 | -31 |
|  |  |  |  |  | -39 | 25 | -20 |
|  |  |  |  |  | -41 | -58 | 35 |
|  |  |  |  |  | -43 | 44 | -17 |
|  |  |  |  |  | -52 | -7 | -19 |
|  |  |  |  |  | -5 | 43 | -31 |
|  |  |  |  |  | -5 | 28 | -32 |
|  |  |  |  |  | -54 | 23 | -8 |
|  |  |  |  |  | -53 | 16 | 46 |
|  |  |  |  |  | -6 | -58 | 17 |
|  |  |  |  |  | 13 | 43 | 11 |
|  |  |  |  |  | 17 | -91 | -25 |
|  |  |  |  |  | 23 | 20 | 42 |
|  |  |  |  |  | 41 | -25 | -25 |
|  |  |  |  |  | 58 | 4 | -15 |
|  |  |  |  |  | 9 | -72 | -26 |
| Scott et al. | 2003 | [10.1016/S1053-8119(03)00083-1](https://doi.org/10.1016/S1053-8119(03)00083-1) | Semantic > phonological decision | 9 | -3 | 48 | 42 |
|  |  |  |  |  | -49 | 34 | -28 |
|  |  |  |  |  | -39 | 15 | -38 |
|  |  |  |  |  | -59 | -4 | -23 |
|  |  |  |  |  | -36 | -24 | -25 |
|  |  |  |  |  | -63 | -43 | -3 |
|  |  |  |  |  | -53 | -66 | 27 |
| Seghier et al. | 2010 | 10.1523/JNEUROSCI.3377-10.2010 | Semantic > perceptual decision | 94 | -30 | -66 | 42 |
|  |  |  |  |  | -48 | -68 | 28 |
|  |  |  |  |  | -48 | -68 | 20 |
|  |  |  |  |  | -34 | -64 | 24 |
| Stringaris et al. | 2007 | [10.1016/j.bandl.2005.08.001](https://doi.org/10.1016/j.bandl.2005.08.001) | Literal > meaningless sentences | 11 | -54 | -7 | -11 |
|  |  |  |  |  | -22 | -76 | 20 |
|  |  |  |  |  | 52 | 20 | 15 |
|  |  |  |  |  | 41 | -1 | 42 |
| Sugiura et al. | 2006 | [10.1016/j.neuroimage.2006.01.002](https://doi.org/10.1016/j.neuroimage.2006.01.002) | Personal > famous names | 24 | -50 | -70 | 41 |
|  |  |  |  |  | 54 | -72 | 37 |
|  |  |  |  |  | -9 | -66 | 36 |
|  |  |  |  |  | 12 | -64 | 33 |
| Sugiura et al. | 2006 | [10.1016/j.neuroimage.2006.01.002](https://doi.org/10.1016/j.neuroimage.2006.01.002) | Famous > unfamiliar names | 24 | -47 | 14 | -50 |
|  |  |  |  |  | 50 | 15 | -52 |
|  |  |  |  |  | -56 | 0 | -42 |
|  |  |  |  |  | -65 | -44 | 37 |
| Sugiura et al. | 2006 | [10.1016/j.neuroimage.2006.01.002](https://doi.org/10.1016/j.neuroimage.2006.01.002) | Personal > unfamiliar names | 24 | -34 | 11 | -43 |
|  |  |  |  |  | 37 | 15 | -42 |
|  |  |  |  |  | -60 | -6 | -39 |
|  |  |  |  |  | 61 | 1 | -39 |
|  |  |  |  |  | -59 | -49 | 35 |
|  |  |  |  |  | -41 | -81 | 38 |
|  |  |  |  |  | 56 | -76 | 36 |
|  |  |  |  |  | -5 | -68 | 29 |
|  |  |  |  |  | 10 | -60 | 30 |
| Wirth et al. | 2011 | [10.1016/j.neuroimage.2010.10.039](https://doi.org/10.1016/j.neuroimage.2010.10.039) | Semantic > phonological decision | 21 | -48 | 17 | -28 |
|  |  |  |  |  | -53 | -20 | -11 |
|  |  |  |  |  | -57 | -68 | 21 |
|  |  |  |  |  | 27 | -86 | -32 |
|  |  |  |  |  | -5 | 38 | 53 |
|  |  |  |  |  | -2 | 38 | -11 |
|  |  |  |  |  | -2 | -56 | 12 |
| Woodard et al. | 2007 | 10.1162/jocn.2007.19.7.1113 | Familiar > unfamiliar names | 15 | -1 | -51 | 22 |
|  |  |  |  |  | 2 | 49 | -8 |
|  |  |  |  |  | -17 | 36 | 38 |
|  |  |  |  |  | -49 | 24 | -5 |
|  |  |  |  |  | 27 | 26 | 38 |
|  |  |  |  |  | -57 | -13 | -17 |
|  |  |  |  |  | -46 | -69 | 29 |
|  |  |  |  |  | -60 | -42 | -7 |
|  |  |  |  |  | 53 | -62 | 15 |
|  |  |  |  |  | 27 | -21 | -16 |
|  |  |  |  |  | 18 | 9 | 15 |
|  |  |  |  |  | -10 | -48 | 52 |
| Sharp, Scott & Wise | 2004 | 10.1093/cercor/bhg086 | Semantic > phonological decision | 9 | -38 | -18 | -32 |
|  |  |  |  |  | 36 | -24 | -20 |
|  |  |  |  |  | 28 | 12 | -36 |
|  |  |  |  |  | -32 | 24 | -16 |
|  |  |  |  |  | -4 | 52 | 40 |
|  |  |  |  |  | -8 | 62 | 24 |
|  |  |  |  |  | 16 | 54 | 26 |
|  |  |  |  |  | -40 | -76 | 30 |
|  |  |  |  |  | 24 | -82 | -34 |
| Gurd et al. | 2002 | [doi.org/10.1093/brain/awf093](https://doi.org/10.1093/brain/awf093) | Category > rote fluency | 11 | -3 | 25 | 38 |
|  |  |  |  |  | 10 | 26 | 31 |
|  |  |  |  |  | -33 | 45 | 23 |
|  |  |  |  |  | 38 | 53 | 19 |
|  |  |  |  |  | -38 | 24 | -17 |
|  |  |  |  |  | 42 | 28 | -19 |
|  |  |  |  |  | -37 | 8 | 27 |
|  |  |  |  |  | 9 | -88 | -43 |
| Thioux et al. | 2005 | 10.1016/j.cogbrainres.2005.02.009 | Animals > numbers | 6 | -47 | -23 | -37 |
|  |  |  |  |  | 21 | -85 | -10 |
| von Kriegstein et al. | 2003 | [10.1016/S0926-6410(03)00079-X](https://doi.org/10.1016/S0926-6410(03)00079-X) | Sentence > speech envelope | 14 | -67 | -18 | -5 |
|  |  |  |  |  | -64 | -41 | -3 |
|  |  |  |  |  | 69 | -6 | -22 |
|  |  |  |  |  | -28 | -63 | -1 |
|  |  |  |  |  | 27 | -53 | 0 |
|  |  |  |  |  | 34 | -56 | -6 |
|  |  |  |  |  | -41 | -71 | 10 |
|  |  |  |  |  | 40 | -94 | 4 |
|  |  |  |  |  | -64 | -10 | -19 |
| Xiao et al. | 2005 | 10.1002/hbm.20105 | Words > pseudowords | 14 | -50 | -59 | 22 |
|  |  |  |  |  | 55 | -47 | 14 |
|  |  |  |  |  | -10 | -41 | 31 |
| Cappa et al. | 1998 | [10.1006/nimg.1998.0368](https://doi.org/10.1006/nimg.1998.0368) | Words > pseudowords | 13 | -46 | 27 | 16 |
|  |  |  |  |  | -44 | -73 | 30 |
|  |  |  |  |  | -11 | 28 | 47 |
|  |  |  |  |  | -14 | 40 | 37 |
|  |  |  |  |  | 1 | -59 | 15 |
|  |  |  |  |  | 47 | -76 | 34 |
| Craik et al. | 1999 | [10.1111/1467-9280.00102](https://doi.org/10.1111%2F1467-9280.00102) | Trait desirabillity > phonological decision | 8 | -8 | 56 | -15 |
|  |  |  |  |  | -5 | 52 | 31 |
|  |  |  |  |  | -38 | 39 | -12 |
|  |  |  |  |  | -44 | -67 | 25 |
| Craik et al. | 1999 | [10.1111/1467-9280.00102](https://doi.org/10.1111%2F1467-9280.00102) | Other trait matching judgement > phonological decision | 8 | -6 | 56 | -15 |
|  |  |  |  |  | -5 | 52 | 35 |
|  |  |  |  |  | -3 | 22 | -16 |
|  |  |  |  |  | -40 | 10 | -23 |
|  |  |  |  |  | -5 | -54 | 19 |
|  |  |  |  |  | -46 | -69 | 25 |
| Craik et al. | 1999 | [10.1111/1467-9280.00102](https://doi.org/10.1111%2F1467-9280.00102) | Self trait matching judgement > phonological decision | 8 | -3 | 54 | 30 |
|  |  |  |  |  | -34 | 26 | -16 |
| Baumgaertner et al. | 2007 | 10.1.1.892.5481 | Sentences > reversed sentences | 19 | -42 | -45 | -18 |
|  |  |  |  |  | -3 | 9 | 54 |
|  |  |  |  |  | -54 | 36 | 9 |
|  |  |  |  |  | -45 | 6 | 33 |
| Baumgaertner et al. | 2007 | 10.1.1.892.5481 | Videos > scrambled videos | 19 | 48 | -69 | -3 |
|  |  |  |  |  | -48 | -72 | 0 |
|  |  |  |  |  | 48 | 15 | 21 |
|  |  |  |  |  | -21 | 3 | 57 |
|  |  |  |  |  | -48 | 9 | 24 |
|  |  |  |  |  | -36 | 33 | -15 |
|  |  |  |  |  | 39 | 36 | -12 |
|  |  |  |  |  | -6 | -48 | 6 |
|  |  |  |  |  | 33 | 0 | 54 |
|  |  |  |  |  | 6 | 21 | 51 |
|  |  |  |  |  | -51 | 36 | 6 |
| Dapretto et al. | 1999 | [10.1016/s0896-6273(00)80855-7](https://doi.org/10.1016/s0896-6273(00)80855-7) | Semantic > syntax | 8 | -51 | 22 | -10 |
| Davis et al. | 2004 | 10.1016/S0093-934X(03)00471-1 | Words > letter strings | 11 | -63 | -42 | -3 |
|  |  |  |  |  | -45 | -42 | -12 |
|  |  |  |  |  | -6 | 27 | -42 |
|  |  |  |  |  | -48 | 30 | -15 |
|  |  |  |  |  | -33 | 27 | -12 |
|  |  |  |  |  | -48 | 24 | 18 |
|  |  |  |  |  | -18 | -9 | -15 |
|  |  |  |  |  | 9 | 27 | 42 |
|  |  |  |  |  | 33 | -12 | 60 |
| Devlin et al. | 2002 | [10.1016/s0028-3932(01)00066-5](https://doi.org/10.1016/s0028-3932(01)00066-5) | Semantic > letter detection | 12 | -28 | 2 | -26 |
|  |  |  |  |  | -34 | 16 | -20 |
|  |  |  |  |  | -62 | -20 | -6 |
|  |  |  |  |  | 42 | 12 | -26 |
|  |  |  |  |  | 32 | 6 | -50 |
|  |  |  |  |  | 44 | -8 | -40 |
| Devlin et al. | 2002 | [10.1016/s0028-3932(01)00066-5](https://doi.org/10.1016/s0028-3932(01)00066-5) | Semantic > letter categorisation | 8 | -40 | -12 | -34 |
|  |  |  |  |  | -36 | -32 | -20 |
|  |  |  |  |  | -28 | 12 | -30 |
|  |  |  |  |  | -48 | 36 | -4 |
|  |  |  |  |  | -44 | 18 | 26 |
|  |  |  |  |  | -6 | 34 | 46 |
|  |  |  |  |  | 18 | -84 | -32 |
| Devlin et al. | 2002 | [10.1016/s0028-3932(01)00066-5](https://doi.org/10.1016/s0028-3932(01)00066-5) | Semantic > letter categorisation | 8 | -58 | -38 | -2 |
|  |  |  |  |  | -64 | -44 | -2 |
|  |  |  |  |  | -46 | 30 | -8 |
|  |  |  |  |  | -48 | 36 | 4 |
|  |  |  |  |  | -48 | 24 | 24 |
|  |  |  |  |  | 48 | 40 | 12 |
|  |  |  |  |  | 48 | 46 | -8 |
|  |  |  |  |  | 46 | 30 | 28 |
|  |  |  |  |  | 34 | 22 | -8 |
|  |  |  |  |  | -2 | 40 | 40 |
|  |  |  |  |  | -6 | 18 | 50 |
|  |  |  |  |  | -4 | 26 | 46 |
|  |  |  |  |  | -20 | 26 | 46 |
|  |  |  |  |  | -44 | 40 | 26 |
|  |  |  |  |  | -38 | -62 | 40 |
|  |  |  |  |  | -44 | -74 | 40 |
|  |  |  |  |  | -46 | -62 | 24 |
|  |  |  |  |  | 38 | -62 | 40 |
|  |  |  |  |  | -6 | -56 | 22 |
|  |  |  |  |  | -6 | -82 | -28 |
|  |  |  |  |  | 6 | -78 | -20 |
|  |  |  |  |  | -36 | -44 | -32 |
|  |  |  |  |  | -38 | -50 | -24 |
|  |  |  |  |  | -52 | -66 | -20 |
|  |  |  |  |  | -4 | -22 | 8 |
|  |  |  |  |  | -18 | -6 | 14 |
|  |  |  |  |  | 8 | -82 | -28 |
| Ebisch et al. | 2007 | [10.1093/cercor/bhm001](https://doi.org/10.1093/cercor/bhm001) | Real > control items | 17 | -33 | -61 | 36 |
|  |  |  |  |  | -52 | 28 | 23 |
|  |  |  |  |  | -31 | 11 | 55 |
|  |  |  |  |  | -5 | -67 | 30 |
|  |  |  |  |  | -35 | -36 | -13 |
| Baumgaertener et al. | 2002 | [10.1006/nimg.2002.1134](https://doi.org/10.1006/nimg.2002.1134) | Word > pseudoword in sentence | 9 | -42 | -66 | 33 |
|  |  |  |  |  | -63 | -42 | 0 |
|  |  |  |  |  | -60 | -48 | -12 |
|  |  |  |  |  | -30 | 24 | 48 |
|  |  |  |  |  | -54 | -3 | -30 |
| Giraud et al. | 2001 | 10.1162/08989290152541421 | Meaningful > meaningless words & sounds | 12 | -50 | 16 | 35 |
|  |  |  |  |  | -57 | -9 | -21 |
|  |  |  |  |  | -72 | -45 | -15 |
| Grossman et al. | 2002a | [10.1006/nimg.2001.1028](https://doi.org/10.1006/nimg.2001.1028) | Nouns > pseudowords | 16 | -55 | -72 | -6 |
|  |  |  |  |  | -21 | 54 | 4 |
|  |  |  |  |  | -12 | -71 | 7 |
| Grossman et al., 2002b | 2002b | 0.1002/hbm.10017 | Verbs > pseudowords | 16 | -68 | -49 | 15 |
|  |  |  |  |  | -4 | 43 | -22 |
|  |  |  |  |  | 1 | -96 | 9 |
| Hagoort et al. | 1999 | [10.1162/089892999563490](https://doi.org/10.1162/089892999563490) | Words > pseudowords | 10 | -18 | -50 | 1 |
|  |  |  |  |  | 67 | -35 | 18 |
|  |  |  |  |  | 42 | -22 | 6 |
|  |  |  |  |  | -53 | -36 | -14 |
|  |  |  |  |  | 60 | -48 | 7 |
|  |  |  |  |  | 63 | -28 | -14 |
|  |  |  |  |  | 14 | -16 | 39 |
|  |  |  |  |  | -1 | -13 | 54 |
| Henke et al. | 1999 | [10.1162/089892999563490](https://doi.org/10.1162/089892999563490) | Semantic > orthographic decision | 12 | -55 | 23 | 3 |
|  |  |  |  |  | 42 | -84 | -51 |
|  |  |  |  |  | 72 | -7 | -32 |
|  |  |  |  |  | 46 | 8 | -33 |
|  |  |  |  |  | 58 | 28 | 45 |
|  |  |  |  |  | -10 | 20 | 7 |
|  |  |  |  |  | -27 | -93 | -53 |
| Herbster et al. | 1997 | [10.1002/(SICI)1097-0193](https://doi.org/10.1002/(SICI)1097-0193(1997)5:2%3C84::AID-HBM2%3E3.0.CO;2-I) | Word > pseudoword | 10 | -38 | -33 | -28 |
| Binder et al. | 1999 | [10.1162/089892999563265](https://doi.org/10.1162/089892999563265) | Semantic > phonological decision | 30 | -47 | -71 | 38 |
|  |  |  |  |  | -7 | -56 | 18 |
|  |  |  |  |  | -17 | 35 | 46 |
|  |  |  |  |  | -27 | -38 | -18 |
| Joubert et al. | 2004 | [10.1016/S0093-934X(03)00403-6](https://doi.org/10.1016/S0093-934X(03)00403-6) | Low frequency words > nonwords | 10 | -53 | -35 | 15 |
|  |  |  |  |  | -49 | -23 | 12 |
|  |  |  |  |  | 65 | -22 | 18 |
|  |  |  |  |  | 61 | 3 | -3 |
|  |  |  |  |  | 8 | -75 | 24 |
|  |  |  |  |  | -50 | -67 | -32 |
| Kuchinke et al. | 2005 | [10.1016/j.neuroimage.2005.06.050](https://doi.org/10.1016/j.neuroimage.2005.06.050) | Word > pseudoword | 20 | -36 | 27 | 45 |
|  |  |  |  |  | -15 | 57 | 9 |
|  |  |  |  |  | -48 | -69 | 30 |
|  |  |  |  |  | -63 | -21 | -24 |
|  |  |  |  |  | -63 | -54 | -9 |
|  |  |  |  |  | -39 | -69 | 42 |
|  |  |  |  |  | -21 | 39 | 42 |
|  |  |  |  |  | -39 | -78 | 36 |
|  |  |  |  |  | -12 | 57 | 39 |
|  |  |  |  |  | -6 | -30 | 33 |
|  |  |  |  |  | -6 | 27 | -18 |
|  |  |  |  |  | 54 | -63 | 18 |
|  |  |  |  |  | 42 | -69 | 33 |
|  |  |  |  |  | 12 | -57 | 27 |
| Binder et al. | 2003 | [10.1162/089892903321593108](https://doi.org/10.1162/089892903321593108) | Word > pseudoword | 24 | -25 | 15 | 55 |
|  |  |  |  |  | -16 | 30 | 48 |
|  |  |  |  |  | -7 | 49 | 27 |
|  |  |  |  |  | -35 | 9 | 47 |
|  |  |  |  |  | -31 | 21 | 34 |
|  |  |  |  |  | 0 | 46 | 13 |
|  |  |  |  |  | -1 | 47 | 2 |
|  |  |  |  |  | -9 | 40 | -5 |
|  |  |  |  |  | 1 | 28 | -12 |
|  |  |  |  |  | -38 | 38 | -5 |
|  |  |  |  |  | -50 | -64 | 26 |
|  |  |  |  |  | -38 | -74 | 40 |
|  |  |  |  |  | -41 | -76 | 27 |
|  |  |  |  |  | -51 | -48 | 43 |
|  |  |  |  |  | -50 | -65 | 14 |
|  |  |  |  |  | -38 | -62 | 52 |
|  |  |  |  |  | -59 | -47 | -5 |
|  |  |  |  |  | -59 | -49 | -15 |
|  |  |  |  |  | -1 | -38 | 35 |
|  |  |  |  |  | -9 | -60 | 12 |
|  |  |  |  |  | -5 | -58 | 41 |
|  |  |  |  |  | -4 | -54 | 27 |
|  |  |  |  |  | 1 | -25 | 39 |
|  |  |  |  |  | -7 | -68 | 56 |
|  |  |  |  |  | 7 | -93 | 4 |
|  |  |  |  |  | -23 | -10 | -14 |
|  |  |  |  |  | -43 | -1 | 24 |
|  |  |  |  |  | -55 | 1 | 18 |
|  |  |  |  |  | -51 | -8 | 43 |
|  |  |  |  |  | 44 | 5 | 33 |
|  |  |  |  |  | 44 | 20 | 29 |
|  |  |  |  |  | -1 | 3 | 51 |
|  |  |  |  |  | 0 | 13 | 45 |
|  |  |  |  |  | 27 | 49 | 19 |
|  |  |  |  |  | -40 | 17 | 2 |
| Laine et al. | 2002 | [10.1016/S0926-6410(01)00095-7](https://doi.org/10.1016/S0926-6410(01)00095-7) | Animal generation > reading | 7 | 34 | -16 | -32 |
|  |  |  | Tool generation > reading |  | 16 | 44 | 24 |
|  |  |  |  |  | -24 | 54 | -12 |
|  |  |  |  |  | 18 | 60 | -4 |
|  |  |  |  |  | 6 | -70 | 8 |
| Meyer et al. | 2002 | 10.1002/hbm.10042 | Word > pseudoword sentence | 14 | -48 | -56 | 30 |
|  |  |  |  |  | 42 | -64 | 30 |
| Noppeney et al. | 2003 | [10.1016/S0093-934X(02)00525-4](https://doi.org/10.1016/S0093-934X(02)00525-4) | Normal > reversed words | 9 | -57 | -54 | -18 |
|  |  |  |  |  | -29 | -41 | -25 |
|  |  |  |  |  | -49 | 28 | 7 |
|  |  |  |  |  | 1 | -62 | 8 |
|  |  |  |  |  | 14 | -97 | -29 |
|  |  |  |  |  | 5 | -18 | -30 |
|  |  |  |  |  | -8 | 73 | 2 |
| Orfanidou et al. | 2006 | 10.1162/jocn.2006.18.8.1237 | Words > pseudowords | 13 | -38 | -36 | -16 |
|  |  |  |  |  | -38 | -70 | 24 |
|  |  |  |  |  | -44 | -80 | 28 |
|  |  |  |  |  | -58 | -42 | 46 |
|  |  |  |  |  | -32 | -82 | 36 |
|  |  |  |  |  | -54 | -54 | 30 |
|  |  |  |  |  | -30 | -76 | 46 |
|  |  |  |  |  | -36 | -68 | 48 |
|  |  |  |  |  | -34 | 58 | 16 |
|  |  |  |  |  | 16 | -66 | 30 |
|  |  |  |  |  | 14 | -66 | 40 |
|  |  |  |  |  | 8 | -32 | 46 |
|  |  |  |  |  | -4 | -14 | 40 |
|  |  |  |  |  | -8 | -64 | 34 |
|  |  |  |  |  | 12 | -56 | 38 |
|  |  |  |  |  | -12 | -50 | 28 |
|  |  |  |  |  | 42 | -72 | 20 |
|  |  |  |  |  | 34 | -82 | 36 |
|  |  |  |  |  | -22 | 26 | -16 |
|  |  |  |  |  | -18 | 12 | -16 |
|  |  |  |  |  | 14 | 12 | -18 |
|  |  |  |  |  | -36 | 42 | -12 |
|  |  |  |  |  | 54 | -50 | 30 |
|  |  |  |  |  | 52 | -52 | 20 |
|  |  |  |  |  | -56 | 0 | -20 |
|  |  |  |  |  | -16 | 44 | 10 |
|  |  |  |  |  | -6 | 38 | 32 |
|  |  |  |  |  | -12 | 50 | 30 |
|  |  |  |  |  | 18 | 20 | 54 |
|  |  |  |  |  | 56 | -8 | 10 |
|  |  |  |  |  | 54 | -2 | 16 |
|  |  |  |  |  | 50 | -8 | -14 |
|  |  |  |  |  | 42 | -16 | -8 |
| Perani et al. | 1999 | [10.1016/S0028-3932(98)00073-6](https://doi.org/10.1016/S0028-3932(98)00073-6) | Living objects > shapes | 11 | -18 | -104 | -25 |
|  |  |  |  |  | -49 | 34 | -25 |
|  |  |  |  |  | 27 | -100 | -35 |
| Perani et al. | 1999 | [10.1016/S0028-3932(98)00073-6](https://doi.org/10.1016/S0028-3932(98)00073-6) | Nonliving objects > shapes | 11 | -38 | 42 | -8 |
|  |  |  |  |  | -66 | -41 | 10 |
|  |  |  |  |  | -59 | -49 | -8 |
| Perani et al. | 1999 | [10.1016/S0028-3932(98)00073-6](https://doi.org/10.1016/S0028-3932(98)00073-6) | Living words > pseudowords | 8 | 4 | -7 | 45 |
|  |  |  |  |  | 25 | -79 | 3 |
|  |  |  |  |  | 22 | -32 | -38 |
| Perani et al. | 1999 | [10.1016/S0028-3932(98)00073-6](https://doi.org/10.1016/S0028-3932(98)00073-6) | Nonliving words > pseudowords | 8 | -7 | -80 | -1 |
|  |  |  |  |  | 8 | -88 | 0 |
|  |  |  |  |  | 4 | -96 | 9 |
|  |  |  |  |  | 4 | -9 | 41 |
|  |  |  |  |  | -12 | 16 | -10 |
|  |  |  |  |  | -16 | -61 | 15 |
|  |  |  |  |  | -24 | -37 | 71 |
|  |  |  |  |  | -31 | -14 | 11 |
|  |  |  |  |  | 53 | -10 | 9 |
|  |  |  |  |  | 10 | -80 | -1 |
|  |  |  |  |  | 38 | -24 | 38 |
| Pilgrim et al. | 2002 | [10.1006/nimg.2002.1105](https://doi.org/10.1006/nimg.2002.1105) | Words > letter strings | 14 | -50 | 30 | -14 |
|  |  |  |  |  | -50 | 34 | -6 |
|  |  |  |  |  | -44 | 36 | -10 |
|  |  |  |  |  | 26 | 22 | -4 |
|  |  |  |  |  | 36 | 30 | -18 |
|  |  |  |  |  | 22 | 22 | 2 |
|  |  |  |  |  | -2 | 24 | 48 |
|  |  |  |  |  | 4 | 28 | 44 |
|  |  |  |  |  | -2 | 38 | 42 |
|  |  |  |  |  | -36 | -30 | -22 |
|  |  |  |  |  | -40 | -40 | -26 |
|  |  |  |  |  | -48 | -50 | -18 |
|  |  |  |  |  | -48 | -44 | 0 |
|  |  |  |  |  | -62 | -40 | 4 |
|  |  |  |  |  | -40 | -46 | -4 |
|  |  |  |  |  | -12 | 14 | 2 |
| Rissman et al. | 2003 | [10.1162/089892903322598120](https://doi.org/10.1162/089892903322598120) | Words > pseudowords | 15 | -65 | -46 | -7 |
|  |  |  |  |  | -1 | 53 | -4 |
|  |  |  |  |  | -1 | -55 | 48 |
|  |  |  |  |  | -35 | -65 | 41 |
|  |  |  |  |  | 52 | -62 | 34 |
| Robertson et al. | 2000 | [10.1111/1467-9280.00251](https://dx.doi.org/10.1111%2F1467-9280.00251) | Indefinite article sentence > letter strings | 8 | -70 | -39 | 0 |
|  |  |  |  |  | -42 | -57 | -27 |
|  |  |  |  |  | 14 | -75 | 7 |
|  |  |  |  |  | -41 | 13 | 53 |
|  |  |  |  |  | -60 | 29 | -6 |
|  |  |  |  |  | 59 | -28 | -3 |
|  |  |  |  |  | -21 | -3 | -26 |
|  |  |  |  |  | -5 | 9 | 12 |
| Robertson et al. | 2000 | [10.1111/1467-9280.00251](https://dx.doi.org/10.1111%2F1467-9280.00251) | Definite article sentence > letter strings | 8 | -66 | -43 | 5 |
|  |  |  |  |  | -42 | -59 | -25 |
|  |  |  |  |  | 59 | -9 | -27 |
|  |  |  |  |  | -51 | 27 | 16 |
|  |  |  |  |  | -3 | -97 | 1 |
|  |  |  |  |  | 42 | -50 | -29 |
|  |  |  |  |  | 58 | -44 | 14 |
|  |  |  |  |  | -37 | 20 | 61 |
| Cai et al. | 2007 | 10.1097/WNR.0b013e32810f2de7 | Words > pseudowords | 15 | -36 | 36 | -25 |
|  |  |  |  |  | -48 | 29 | 18 |
| Crinion et al. | 2003 | [10.1093/brain/awg104](https://doi.org/10.1093/brain/awg104) | Stories > reversed stories | 17 | -44 | 26 | -16 |
|  |  |  |  |  | -52 | 8 | -20 |
|  |  |  |  |  | -52 | 10 | -18 |
|  |  |  |  |  | -44 | 20 | -24 |
|  |  |  |  |  | -46 | 16 | -16 |
|  |  |  |  |  | -58 | -6 | -12 |
|  |  |  |  |  | -58 | -42 | 2 |
|  |  |  |  |  | -56 | -52 | 20 |
|  |  |  |  |  | -56 | -58 | 18 |
|  |  |  |  |  | -42 | -16 | -34 |
|  |  |  |  |  | -42 | -16 | -38 |
|  |  |  |  |  | 50 | 14 | -24 |
|  |  |  |  |  | 52 | 6 | -32 |
|  |  |  |  |  | 56 | -8 | -18 |
|  |  |  |  |  | 60 | -4 | -18 |
|  |  |  |  |  | 60 | -36 | 0 |
|  |  |  |  |  | 64 | -54 | 2 |
| Damasio et al. | 2001 | 10.1006/nimg.2001.077 | Actions without implement > control task | 10 | -36 | 30 | -11 |
|  |  |  |  |  | -46 | 33 | 7 |
|  |  |  |  |  | -36 | 12 | 34 |
|  |  |  |  |  | -31 | 19 | -33 |
|  |  |  |  |  | -55 | -53 | -13 |
|  |  |  |  |  | -44 | -34 | -16 |
|  |  |  |  |  | 58 | -35 | 43 |
|  |  |  |  |  | -39 | -81 | 38 |
|  |  |  |  |  | -58 | -24 | 30 |
|  |  |  |  |  | 51 | -81 | 23 |
|  |  |  |  |  | -49 | -78 | 24 |
|  |  |  |  |  | 57 | 1 | 3 |
|  |  |  |  |  | 61 | -1 | 38 |
|  |  |  |  |  | 72 | -29 | 13 |
|  |  |  |  |  | -64 | -19 | 14 |
|  |  |  |  |  | -20 | -81 | 42 |
|  |  |  |  |  | 35 | -90 | -7 |
|  |  |  |  |  | -34 | -87 | -7 |
|  |  |  |  |  | 4 | -82 | 15 |
|  |  |  |  |  | 13 | -61 | 12 |
|  |  |  |  |  | -42 | 0 | 1 |
|  |  |  |  |  | 34 | -4 | 1 |
|  |  |  |  |  | 13 | -29 | -37 |
| Damasio et al. | 2001 | 10.1006/nimg.2001.077 | Actions with implement > control task | 10 | -52 | 36 | 13 |
|  |  |  |  |  | -36 | 31 | -11 |
|  |  |  |  |  | -46 | 14 | 27 |
|  |  |  |  |  | -25 | 11 | 37 |
|  |  |  |  |  | -54 | -51 | -12 |
|  |  |  |  |  | -42 | -32 | -17 |
|  |  |  |  |  | 58 | -40 | 35 |
|  |  |  |  |  | 63 | -46 | 31 |
|  |  |  |  |  | -36 | -82 | 38 |
|  |  |  |  |  | 39 | 40 | 30 |
|  |  |  |  |  | 49 | 2 | 8 |
|  |  |  |  |  | 58 | 3 | 38 |
|  |  |  |  |  | 59 | -19 | 10 |
|  |  |  |  |  | 72 | -30 | 11 |
|  |  |  |  |  | -57 | -11 | 3 |
|  |  |  |  |  | 34 | -79 | 43 |
|  |  |  |  |  | -21 | -83 | 40 |
|  |  |  |  |  | 37 | -88 | -10 |
|  |  |  |  |  | 22 | -95 | -6 |
|  |  |  |  |  | -36 | -89 | -10 |
|  |  |  |  |  | 9 | -78 | 13 |
| Engelien et al. | 2006 | 10.1007/s00702-005-0342-0 | Meaningful > meaningless sounds | 6 | -66 | -20 | 6 |
|  |  |  |  |  | -58 | 24 | 12 |
|  |  |  |  |  | -12 | -26 | -16 |
|  |  |  |  |  | 74 | 24 | 8 |
|  |  |  |  |  | 26 | 40 | -28 |
| Farias et al. | 2005 | N/A | Responsive naming > reversed speech | 10 | -53 | 6 | -15 |
|  |  |  |  |  | -51 | -12 | -9 |
|  |  |  |  |  | -53 | -42 | -1 |
|  |  |  |  |  | -29 | -1 | -41 |
| Farias et al. | 2005 | N/A | Confrontation naming > viewing lines | 10 | -53 | -44 | -1 |
|  |  |  |  |  | -63 | -29 | -16 |
| Rogers et al. | 2006 | [10.3758/cabn.6.3.201](https://doi.org/10.3758/cabn.6.3.201) | Semantics > baseline | 12 | -42 | -74 | -16 |
|  |  |  |  |  | -32 | -90 | 4 |
|  |  |  |  |  | -38 | -56 | -18 |
|  |  |  |  |  | -28 | -38 | -16 |
|  |  |  |  |  | 48 | -78 | -12 |
|  |  |  |  |  | 38 | -88 | 4 |
|  |  |  |  |  | 38 | -60 | -18 |
|  |  |  |  |  | 26 | -28 | -20 |
| Rogers et al. | 2006 | [10.3758/cabn.6.3.201](https://doi.org/10.3758/cabn.6.3.201) | Specific-level judgement > baseline | 12 | -54 | 6 | -26 |
|  |  |  |  |  | 48 | 22 | -28 |
| Stowe et al. | 1999 | [10.1111/1469-8986.3660786](https://doi.org/10.1111/1469-8986.3660786) | Sentences > word lists | 12 | -53 | -3 | -21 |
|  |  |  |  |  | -12 | 43 | 27 |
|  |  |  |  |  | -55 | 20 | 8 |
|  |  |  |  |  | -49 | 13 | 8 |
|  |  |  |  |  | -29 | 6 | -5 |
|  |  |  |  |  | 27 | -3 | -4 |
|  |  |  |  |  | -29 | -24 | -24 |
|  |  |  |  |  | -23 | -28 | -19 |
| Tieleman et al. | 2005 | [10.1016/j.neuroimage.2005.02.017](https://doi.org/10.1016/j.neuroimage.2005.02.017) | Self-paced semantic > perceptual decision | 22 | -53 | 30 | 5 |
|  |  |  |  |  | -38 | 33 | -27 |
|  |  |  |  |  | -55 | 23 | 18 |
|  |  |  |  |  | -46 | 21 | 23 |
|  |  |  |  |  | 0 | 26 | 50 |
|  |  |  |  |  | 4 | 41 | 52 |
|  |  |  |  |  | -1 | 41 | 52 |
|  |  |  |  |  | -1 | 30 | 38 |
|  |  |  |  |  | 37 | 31 | -22 |
|  |  |  |  |  | 35 | 36 | -26 |
|  |  |  |  |  | -51 | 9 | -20 |
|  |  |  |  |  | -38 | 24 | 9 |
|  |  |  |  |  | -53 | 10 | 10 |
|  |  |  |  |  | -53 | -42 | 3 |
|  |  |  |  |  | 46 | 16 | -27 |
|  |  |  |  |  | -27 | -17 | -20 |
|  |  |  |  |  | -25 | -25 | -15 |
|  |  |  |  |  | -27 | -32 | -20 |
|  |  |  |  |  | -19 | -14 | -21 |
|  |  |  |  |  | -38 | -47 | -17 |
|  |  |  |  |  | 20 | -15 | -17 |
|  |  |  |  |  | -9 | -92 | 16 |
|  |  |  |  |  | -7 | -81 | 8 |
|  |  |  |  |  | -38 | -47 | -18 |
|  |  |  |  |  | -3 | -71 | 20 |
|  |  |  |  |  | 14 | -90 | 13 |
|  |  |  |  |  | 15 | -90 | 20 |
|  |  |  |  |  | 17 | -68 | 14 |
|  |  |  |  |  | -7 | -62 | 8 |
| Tieleman et al. | 2005 | [10.1016/j.neuroimage.2005.02.017](https://doi.org/10.1016/j.neuroimage.2005.02.017) | Fixed-paced semantic > perceptual decision | 22 | -48 | 21 | 15 |
|  |  |  |  |  | -40 | 25 | -15 |
|  |  |  |  |  | -55 | 24 | 20 |
|  |  |  |  |  | -50 | 6 | 43 |
|  |  |  |  |  | 0 | 21 | 58 |
|  |  |  |  |  | 6 | 27 | 40 |
|  |  |  |  |  | -7 | 24 | 48 |
|  |  |  |  |  | 38 | 30 | 39 |
|  |  |  |  |  | 51 | 24 | -15 |
|  |  |  |  |  | -49 | 18 | -20 |
|  |  |  |  |  | -40 | 15 | 8 |
|  |  |  |  |  | -57 | -37 | 2 |
|  |  |  |  |  | -11 | -95 | 11 |
|  |  |  |  |  | -22 | -93 | -3 |
|  |  |  |  |  | -20 | -94 | -2 |
|  |  |  |  |  | -42 | -52 | -13 |
|  |  |  |  |  | 5 | -52 | -14 |
|  |  |  |  |  | 15 | -92 | 23 |
|  |  |  |  |  | 10 | -81 | -25 |
|  |  |  |  |  | 38 | -59 | -31 |
| Bright et al. | 2004 | [10.1016/j.bandl.2004.01.010](https://doi.org/10.1016/j.bandl.2004.01.010) | Semantics > baseline | 38 | -40 | 32 | -21 |
|  |  |  |  |  | -51 | 30 | 2 |
|  |  |  |  |  | -32 | 19 | -42 |
|  |  |  |  |  | -34 | -41 | -22 |
|  |  |  |  |  | -36 | -17 | -36 |
|  |  |  |  |  | -32 | -27 | -31 |
|  |  |  |  |  | -14 | 49 | 47 |
|  |  |  |  |  | -7 | 57 | 39 |
|  |  |  |  |  | 31 | -95 | -32 |
|  |  |  |  |  | 25 | -89 | -34 |
|  |  |  |  |  | 35 | -89 | -37 |
|  |  |  |  |  | 40 | 26 | -21 |
| Bright et al. | 2004 | [10.1016/j.bandl.2004.01.010](https://doi.org/10.1016/j.bandl.2004.01.010) | Words & pictures > baseline | 38 | -36 | 32 | -18 |
|  |  |  |  |  | -32 | 18 | -35 |
|  |  |  |  |  | -34 | -12 | -36 |
|  |  |  |  |  | -32 | -35 | -25 |
| Bright et al. | 2004 | [10.1016/j.bandl.2004.01.010](https://doi.org/10.1016/j.bandl.2004.01.010) | Words > baseline | 38 | -32 | 10 | -30 |
|  |  |  |  |  | -23 | -37 | -23 |
|  |  |  |  |  | 42 | 16 | -38 |
| Bright et al. | 2004 | [10.1016/j.bandl.2004.01.010](https://doi.org/10.1016/j.bandl.2004.01.010) | Pictures > baseline | 38 | -36 | -54 | -21 |
|  |  |  |  |  | -16 | -83 | -7 |
|  |  |  |  |  | -40 | -83 | -11 |
|  |  |  |  |  | 47 | -83 | -13 |
|  |  |  |  |  | 44 | -56 | -22 |
|  |  |  |  |  | 48 | -84 | -31 |
| D'Arcy et al. | 2007 | 10.1016/j.neures.2006.09.018 | Basic-level living objects > baseline | 10 | 34 | -68 | 24 |
|  |  |  |  |  | -40 | -34 | -22 |
|  |  |  |  |  | -42 | 14 | -38 |
|  |  |  |  |  | 42 | 12 | -36 |
| D'Arcy et al. | 2007 | 10.1016/j.neures.2006.09.018 | Basic-level non-living objects > baseline | 10 | -42 | -60 | -12 |
|  |  |  |  |  | 44 | -70 | -8 |
| D'Arcy et al. | 2007 | 10.1016/j.neures.2006.09.018 | Superordinate-level living objects >baseline | 10 | -42 | -68 | 10 |
|  |  |  |  |  | 46 | -70 | 12 |
| D'Arcy et al. | 2007 | 10.1016/j.neures.2006.09.018 | Superordinate-level non-living objects >baseline | 10 | 40 | -48 | -22 |
|  |  |  |  |  | -42 | -55 | -14 |
| Devlin et al. | 2000 | 10.1006/nimg.2000.059 | Semantics > baseline | 16 | -52 | 34 | 2 |
|  |  |  |  |  | -48 | 18 | 30 |
|  |  |  |  |  | -42 | -14 | -28 |
|  |  |  |  |  | -36 | -32 | -16 |
|  |  |  |  |  | -28 | 10 | -24 |
|  |  |  |  |  | 14 | -78 | -32 |
| Devlin et al. | 2000 | 10.1006/nimg.2000.059 | Semantics > baseline | 16 | -36 | 30 | -20 |
|  |  |  |  |  | -52 | 36 | 4 |
|  |  |  |  |  | -46 | 24 | -8 |
|  |  |  |  |  | 52 | 42 | 16 |
|  |  |  |  |  | 46 | 40 | 10 |
|  |  |  |  |  | -6 | 20 | 50 |
|  |  |  |  |  | 10 | -84 | -32 |
| Foki et al. | 2008 | [10.1016/j.neuroimage.2007.10.020](https://doi.org/10.1016/j.neuroimage.2007.10.020) | Semantic judgement > tongue movements | 23 | -46 | -12 | 38 |
|  |  |  |  |  | -42 | 8 | 27 |
|  |  |  |  |  | -49 | 31 | -5 |
|  |  |  |  |  | -5 | 7 | 58 |
|  |  |  |  |  | -24 | -52 | 44 |
|  |  |  |  |  | -29 | -92 | -15 |
|  |  |  |  |  | -51 | -38 | 9 |
|  |  |  |  |  | -66 | -18 | -6 |
|  |  |  |  |  | 51 | -8 | 33 |
|  |  |  |  |  | 68 | 3 | 5 |
|  |  |  |  |  | 24 | -69 | 62 |
|  |  |  |  |  | 27 | -103 | -2 |
|  |  |  |  |  | 61 | 15 | -27 |
|  |  |  |  |  | 5 | -65 | -30 |
| Gerlach et al. | 1999 | [10.1093/brain/122.11.2159](https://doi.org/10.1093/brain/122.11.2159) | Object decision > pattern discrimination | 15 | 42 | -66 | -12 |
|  |  |  |  |  | 46 | -49 | -21 |
|  |  |  |  |  | 38 | -90 | 6 |
|  |  |  |  |  | 40 | -30 | -27 |
|  |  |  |  |  | 33 | -2 | -45 |
|  |  |  |  |  | 34 | -66 | 55 |
|  |  |  |  |  | -38 | -61 | -9 |
|  |  |  |  |  | -44 | -79 | -12 |
|  |  |  |  |  | -38 | -43 | -20 |
|  |  |  |  |  | -38 | -29 | -28 |
|  |  |  |  |  | -38 | -51 | -17 |
|  |  |  |  |  | -47 | -54 | -19 |
|  |  |  |  |  | -29 | -95 | -1 |
|  |  |  |  |  | -55 | -76 | -3 |
|  |  |  |  |  | -57 | -62 | -18 |
|  |  |  |  |  | -37 | -1 | -33 |
|  |  |  |  |  | 49 | 15 | 22 |
|  |  |  |  |  | 55 | 29 | 10 |
| Giraud et al. | 2004 | [10.1093/cercor/bhg124](https://doi.org/10.1093/cercor/bhg124) | Speech > speech envelope | 8 | 60 | 2 | -18 |
| Giraud et al. | 2004 | [10.1093/cercor/bhg124](https://doi.org/10.1093/cercor/bhg124) | Speech comprehension success | 8 | 56 | 4 | -16 |
|  |  |  |  |  | -58 | -6 | -14 |
| Herbster et al. | 1997 | [10.1002/(SICI)1097-0193(1997)5](https://psycnet.apa.org/doi/10.1002/(SICI)1097-0193(1997)5:2%3C84::AID-HBM2%3E3.0.CO;2-I) | Read > say 'hiya' | 10 | -40 | -44 | -26 |
|  |  |  |  |  | -64 | -18 | -2 |
|  |  |  |  |  | -49 | 1 | -8 |
|  |  |  |  |  | -12 | 3 | -9 |
|  |  |  |  |  | -7 | -70 | -7 |
|  |  |  |  |  | 42 | -80 | -24 |
| Ikuta et al. | 2006 | 10.1016/j.bandl.2005.10.006 | Sentences > word lists | 34 | -12 | -97 | 3 |
|  |  |  |  |  | 17 | -90 | 6 |
|  |  |  |  |  | -52 | 2 | 52 |
|  |  |  |  |  | -53 | -46 | 1 |
|  |  |  |  |  | -66 | -48 | 17 |
|  |  |  |  |  | -47 | 1 | -53 |
|  |  |  |  |  | 29 | -94 | -36 |
|  |  |  |  |  | -60 | 27 | -4 |
| Kang et al. | 2006 | [10.1016/j.neuroimage.2006.03.016](https://doi.org/10.1016/j.neuroimage.2006.03.016) | Speech > white noise | 17 | -65 | -21 | -4 |
|  |  |  |  |  | -32 | -4 | -37 |
|  |  |  |  |  | -43 | 20 | -34 |
|  |  |  |  |  | 78 | -12 | -1 |
|  |  |  |  |  | 60 | 10 | -24 |
| Kang et al. | 2006 | [10.1016/j.neuroimage.2006.03.016](https://doi.org/10.1016/j.neuroimage.2006.03.016) | Speech > white noise | 17 | -65 | -10 | -4 |
|  |  |  |  |  | -56 | 11 | -23 |
|  |  |  |  |  | 69 | -12 | 0 |
|  |  |  |  |  | 44 | 11 | -31 |
| Kang et al. | 2006 | [10.1016/j.neuroimage.2006.03.016](https://doi.org/10.1016/j.neuroimage.2006.03.016) | Speech > baseline | 17 | -44 | 18 | 22 |
|  |  |  |  |  | -7 | 30 | 35 |
|  |  |  |  |  | -49 | 27 | -18 |
| Scott et al. | 2000 | [10.1093/brain/123.12.2400](https://doi.org/10.1093/brain/123.12.2400) | Intelligible > unintelligible speech | 8 | -54 | 6 | -16 |
|  |  |  |  |  | -66 | -12 | -12 |
| Scott et al. | 2004 | 10.1121/1.1639336 | Speech > noise | 7 | -64 | -20 | 2 |
|  |  |  |  |  | -58 | -8 | 2 |
|  |  |  |  |  | -68 | -30 | 10 |
|  |  |  |  |  | 64 | -18 | 2 |
|  |  |  |  |  | 70 | -26 | 4 |
|  |  |  |  |  | 66 | -8 | 0 |
| Scott et al. | 2004 | 10.1121/1.1639336 | Speech > signal correlated noise | 7 | -52 | -14 | 2 |
|  |  |  |  |  | -56 | -32 | 6 |
|  |  |  |  |  | -66 | -16 | 0 |
|  |  |  |  |  | 66 | -24 | 2 |
|  |  |  |  |  | -60 | 8 | -2 |
| Scott et al. | 2004 | 10.1121/1.1639336 | Intelligibility parameter | 7 | -58 | 0 | 0 |
| Thierry et al (2006) | 2006 | [10.1162/jocn.2006.18.6.1018](https://doi.org/10.1162/jocn.2006.18.6.1018) | Auditory real > scrambled words | 12 | -59 | -43 | 3 |
|  |  |  |  |  | -64 | -37 | 0 |
|  |  |  |  |  | -60 | 10 | -23 |
|  |  |  |  |  | -57 | 14 | 20 |
|  |  |  |  |  | 25 | -93 | -27 |
| Thierry et al (2006) | 2006 | [10.1162/jocn.2006.18.6.1018](https://doi.org/10.1162/jocn.2006.18.6.1018) | Auditory real > scrambled sounds | 12 | -61 | -39 | 7 |
|  |  |  |  |  | -62 | -40 | -11 |
|  |  |  |  |  | -60 | 6 | -22 |
|  |  |  |  |  | -53 | 19 | 21 |
|  |  |  |  |  | 18 | -96 | -36 |
| Thierry et al (2006) | 2006 | [10.1162/jocn.2006.18.6.1018](https://doi.org/10.1162/jocn.2006.18.6.1018) | Visual real > distorted sounds | 12 | -64 | -38 | 9 |
|  |  |  |  |  | -64 | -40 | -4 |
|  |  |  |  |  | -62 | 6 | -22 |
|  |  |  |  |  | -53 | 18 | 19 |
|  |  |  |  |  | 27 | -83 | -33 |
| Thierry et al (2006) | 2006 | [10.1162/jocn.2006.18.6.1018](https://doi.org/10.1162/jocn.2006.18.6.1018) | Visual real > distorted videos | 12 | -57 | -41 | 5 |
|  |  |  |  |  | -59 | -40 | -9 |
|  |  |  |  |  | -62 | 10 | -27 |
|  |  |  |  |  | -61 | 19 | 22 |
|  |  |  |  |  | 25 | -93 | -34 |
| Tyler et al (2003) | 2003 | [10.1016/S1053-8119(02)00047-2](https://doi.org/10.1016/S1053-8119(02)00047-2) | Tools > baseline | 12 | -46 | 12 | 32 |
|  |  |  |  |  | -50 | 20 | 12 |
|  |  |  |  |  | -48 | 28 | -10 |
|  |  |  |  |  | -30 | -34 | -26 |
|  |  |  |  |  | -20 | -14 | -14 |
|  |  |  |  |  | -42 | -42 | -26 |
|  |  |  |  |  | 4 | 24 | 36 |
|  |  |  |  |  | 0 | 14 | 48 |
|  |  |  |  |  | -6 | 28 | 54 |
|  |  |  |  |  | 0 | -80 | 8 |
|  |  |  |  |  | 16 | -82 | 18 |
|  |  |  |  |  | 10 | -58 | -2 |
| Tyler et al. | 2003 | [10.1016/S1053-8119(02)00047-2](https://doi.org/10.1016/S1053-8119(02)00047-2) | Animals > baseline | 12 | -24 | -12 | -18 |
|  |  |  |  |  | -42 | -46 | -28 |
|  |  |  |  |  | -48 | -40 | -12 |
|  |  |  |  |  | -28 | 24 | -8 |
|  |  |  |  |  | -30 | 28 | -18 |
|  |  |  |  |  | -44 | 18 | 4 |
|  |  |  |  |  | -8 | -20 | 20 |
|  |  |  |  |  | -6 | 4 | 24 |
|  |  |  |  |  | -4 | -28 | 16 |
|  |  |  |  |  | 10 | -72 | -34 |
|  |  |  |  |  | 8 | -64 | -26 |
|  |  |  |  |  | 34 | 28 | -10 |
|  |  |  |  |  | 42 | 18 | 6 |
|  |  |  |  |  | 18 | 26 | 24 |
|  |  |  |  |  | 16 | -26 | -22 |
|  |  |  |  |  | 10 | -20 | -20 |
|  |  |  |  |  | 38 | -28 | -22 |
|  |  |  |  |  | 4 | 20 | 44 |
| Tyler et al. | 2003 | [10.1016/S1053-8119(02)00047-2](https://doi.org/10.1016/S1053-8119(02)00047-2) | Tool action words > baseline | 12 | -50 | 22 | 10 |
|  |  |  |  |  | -32 | 22 | -6 |
|  |  |  |  |  | -46 | 32 | -10 |
|  |  |  |  |  | 6 | 22 | 34 |
|  |  |  |  |  | -4 | 14 | 46 |
|  |  |  |  |  | -4 | 28 | 44 |
|  |  |  |  |  | -44 | -40 | -24 |
|  |  |  |  |  | -20 | -12 | -24 |
|  |  |  |  |  | -18 | -10 | -14 |
|  |  |  |  |  | -24 | -58 | 2 |
|  |  |  |  |  | -10 | -78 | 6 |
|  |  |  |  |  | -14 | -66 | 2 |
|  |  |  |  |  | 36 | 24 | -8 |
|  |  |  |  |  | 44 | 0 | -14 |
|  |  |  |  |  | 46 | 20 | 6 |
|  |  |  |  |  | 10 | -76 | -34 |
|  |  |  |  |  | 6 | -60 | -22 |
|  |  |  |  |  | 10 | -50 | -16 |
| Tyler et al. | 2003 | [10.1016/S1053-8119(02)00047-2](https://doi.org/10.1016/S1053-8119(02)00047-2) | Biological action > baseline | 12 | -46 | 30 | -12 |
|  |  |  |  |  | -32 | 32 | -18 |
|  |  |  |  |  | -50 | 18 | -6 |
|  |  |  |  |  | -14 | -36 | -2 |
|  |  |  |  |  | -8 | -30 | -2 |
|  |  |  |  |  | -16 | -48 | 0 |
|  |  |  |  |  | -26 | -16 | -12 |
|  |  |  |  |  | -34 | -16 | -26 |
|  |  |  |  |  | -26 | -22 | -18 |
|  |  |  |  |  | -54 | -30 | -6 |
|  |  |  |  |  | -44 | -42 | -16 |
|  |  |  |  |  | -52 | -46 | -8 |
|  |  |  |  |  | -6 | -68 | -34 |
|  |  |  |  |  | 6 | -36 | -26 |
|  |  |  |  |  | 6 | -62 | -24 |
|  |  |  |  |  | 2 | 18 | 40 |
|  |  |  |  |  | -4 | 32 | 40 |
| Vandenberghe et al. | 1996 | [10.1038/383254a0](https://doi.org/10.1038/383254a0) | Semantics > physical size judgement | 6 | -44 | 27 | 16 |
|  |  |  |  |  | -17 | 32 | -21 |
|  |  |  |  |  | -45 | -2 | -35 |
|  |  |  |  |  | -62 | -40 | -4 |
|  |  |  |  |  | -47 | -12 | -34 |
|  |  |  |  |  | -49 | -50 | -21 |
|  |  |  |  |  | -42 | -71 | 30 |
|  |  |  |  |  | -31 | -75 | 48 |
|  |  |  |  |  | -19 | -17 | -16 |
|  |  |  |  |  | 16 | -89 | -27 |
|  |  |  |  |  | 42 | -84 | -51 |
| Vandenberghe et al. | 2002 | 10.1162/08989290260045800 | Sentences > letter strings | 10 | -49 | 3 | -27 |
|  |  |  |  |  | -47 | 5 | -36 |
|  |  |  |  |  | -66 | -56 | 7 |
|  |  |  |  |  | -55 | -55 | 15 |
|  |  |  |  |  | -70 | -17 | -11 |
|  |  |  |  |  | -51 | 25 | -2 |
| Grabowski | 1998 | [10.1006/nimg.1998.0324](https://doi.org/10.1006/nimg.1998.0324) | Naming animals > baseline | 9 | -43 | 25 | -3 |
| Grabowski | 1998 | [10.1006/nimg.1998.0324](https://doi.org/10.1006/nimg.1998.0324) | Naming tools > baseline | 9 | -46 | 27 | 3 |
|  |  |  |  |  | -29 | 59 | 8 |
| Sergent et al. | 1992 | [doi.org/10.1093/brain/115.1.15](https://doi.org/10.1093/brain/115.1.15) | Identity > gender | 7 | 40 | 20 | -43 |
|  |  |  |  |  | -38 | 8 | -35 |
|  |  |  |  |  | -24 | -5 | -40 |
|  |  |  |  |  | 28 | -7 | -40 |
|  |  |  |  |  | 27 | -16 | -26 |
|  |  |  |  |  | -3 | 26 | -26 |
|  |  |  |  |  | -39 | -64 | -11 |
|  |  |  |  |  | 41 | -58 | -12 |
|  |  |  |  |  | -55 | -9 | -13 |
|  |  |  |  |  | 24 | -62 | 3 |
| Sergent et al. | 1992 | [doi.org/10.1093/brain/115.1.15](https://doi.org/10.1093/brain/115.1.15) | Identity > gender | 7 | -58 | -42 | -18 |
|  |  |  |  |  | -39 | -62 | -13 |
|  |  |  |  |  | -56 | -9 | -15 |
|  |  |  |  |  | -42 | -80 | -3 |
|  |  |  |  |  | -32 | -56 | 69 |
|  |  |  |  |  | 58 | -19 | 32 |
|  |  |  |  |  | -56 | -16 | 18 |
|  |  |  |  |  | -1 | 22 | -28 |
| Gorno-Tempini et al. | 1998 | [10.1093/brain/121.11.2103](https://doi.org/10.1093/brain/121.11.2103) | Famous faces > controls | 6 | 28 | 6 | -24 |
|  |  |  |  |  | -40 | 6 | -26 |
|  |  |  |  |  | -4 | 44 | -14 |
|  |  |  |  |  | -2 | 60 | 4 |
|  |  |  |  |  | -2 | -62 | 30 |
|  |  |  |  |  | -8 | -56 | 14 |
| Gorno-Tempini et al. | 1998 | [10.1093/brain/121.11.2103](https://doi.org/10.1093/brain/121.11.2103) | Famous names > controls | 6 | 24 | 4 | -26 |
|  |  |  |  |  | -42 | -2 | -24 |
|  |  |  |  |  | -2 | 44 | -14 |
|  |  |  |  |  | -2 | 56 | 6 |
|  |  |  |  |  | -16 | -62 | 30 |
| Gorno-Tempini et al. | 1998 | [10.1093/brain/121.11.2103](https://doi.org/10.1093/brain/121.11.2103) | Famous > unfamiliar names | 6 | -36 | 12 | -32 |
|  |  |  |  |  | -66 | -48 | 6 |
|  |  |  |  |  | -56 | -62 | -18 |
|  |  |  |  |  | 40 | 6 | -28 |
|  |  |  |  |  | -46 | -16 | -22 |
|  |  |  |  |  | -8 | -50 | -16 |
|  |  |  |  |  | 4 | -48 | 22 |
| Nakamura et al. | 2000 | [doi.org/10.1093/brain/123.9.1903](https://doi.org/10.1093/brain/123.9.1903) | Faces > baseline | 7 | 34 | 23 | -27 |
|  |  |  |  |  | 12 | -78 | -14 |
|  |  |  |  |  | -12 | -78 | -18 |
|  |  |  |  |  | 15 | -92 | 0 |
|  |  |  |  |  | 34 | -62 | -19 |
|  |  |  |  |  | -37 | -81 | -19 |
|  |  |  |  |  | 29 | -6 | -29 |
| Leveroni et al. | 2000 | [10.1523/jneurosci.20-02-00878.2000](https://doi.org/10.1523/jneurosci.20-02-00878.2000) | Familiar > unfamiliar faces | 11 | 42 | 9 | -38 |
|  |  |  |  |  | -16 | 36 | 44 |
|  |  |  |  |  | 12 | 58 | 32 |
|  |  |  |  |  | -1 | 54 | -5 |
|  |  |  |  |  | 57 | -5 | -23 |
|  |  |  |  |  | -55 | -16 | -16 |
|  |  |  |  |  | -52 | -63 | 27 |
|  |  |  |  |  | -55 | -44 | 5 |
|  |  |  |  |  | 49 | -26 | 16 |
|  |  |  |  |  | 64 | -50 | 18 |
|  |  |  |  |  | 27 | -13 | -23 |
|  |  |  |  |  | -1 | -52 | 33 |
|  |  |  |  |  | 10 | -46 | 8 |
| Sugiura et al. | 2001 | 10.1006/nimg.2001.0747 | Identity discrimination > control | 5 | -43 | 23 | -30 |
|  |  |  |  |  | 48 | 23 | -26 |
|  |  |  |  |  | 60 | 1 | -16 |
|  |  |  |  |  | -45 | -74 | -18 |
|  |  |  |  |  | 48 | -76 | -14 |
|  |  |  |  |  | -42 | -42 | -22 |
|  |  |  |  |  | 35 | -34 | -22 |
|  |  |  |  |  | -33 | -8 | -36 |
|  |  |  |  |  | -20 | -2 | -25 |
|  |  |  |  |  | 26 | -1 | -21 |
|  |  |  |  |  | -10 | 1 | -10 |
|  |  |  |  |  | -10 | 35 | -8 |
|  |  |  |  |  | -6 | 50 | -11 |
|  |  |  |  |  | 8 | 70 | 12 |
|  |  |  |  |  | -2 | 59 | 28 |
|  |  |  |  |  | 4 | -53 | 19 |
|  |  |  |  |  | -25 | -88 | -26 |
|  |  |  |  |  | 17 | -86 | -37 |
|  |  |  |  |  | -55 | -1 | -25 |
| Nakamura et al. | 2001 | [10.1016/S0028-3932(01)00037-9](https://doi.org/10.1016/S0028-3932(01)00037-9) | Familiar voice > baseline | 9 | 35 | 12 | -35 |
|  |  |  |  |  | -1 | 65 | -11 |
|  |  |  |  |  | 0 | -61 | 35 |
|  |  |  |  |  | 29 | 4 | -30 |
|  |  |  |  |  | 49 | 22 | 28 |
|  |  |  |  |  | 46 | 49 | -8 |
|  |  |  |  |  | 46 | -12 | -9 |
| Grabowski et al. | 2001 | 10.1002/hbm.1033.abs | Naming > orientation judgement | 10 | -39 | 14 | -28 |
|  |  |  |  |  | 0 | 49 | 16 |
|  |  |  |  |  | -39 | 25 | -5 |
|  |  |  |  |  | 6 | -61 | 19 |
|  |  |  |  |  | -21 | -30 | -25 |
|  |  |  |  |  | 13 | -45 | -23 |
|  |  |  |  |  | -28 | 1 | 34 |
|  |  |  |  |  | -1 | 22 | 44 |
|  |  |  |  |  | -48 | -73 | 36 |
|  |  |  |  |  | 54 | 19 | -20 |
|  |  |  |  |  | 49 | 18 | -22 |
| Damasio et al. | 2004 | [10.1016/j.cognition.2002.07.001](https://doi.org/10.1016/j.cognition.2002.07.001) | People > scrambled pictures | 68 | -43 | 6 | -31 |
|  |  |  |  |  | -36 | 26 | -8 |
|  |  |  |  |  | -2 | 44 | 24 |
|  |  |  |  |  | -13 | 47 | -2 |
|  |  |  |  |  | -10 | -41 | 22 |
|  |  |  |  |  | 0 | -55 | 21 |
|  |  |  |  |  | -2 | -34 | -19 |
| Damasio et al. | 2004 | [10.1016/j.cognition.2002.07.001](https://doi.org/10.1016/j.cognition.2002.07.001) | Animals > scrambled pictures | 68 | -50 | -52 | -15 |
|  |  |  |  |  | -29 | -33 | -21 |
|  |  |  |  |  | 30 | -38 | -20 |
|  |  |  |  |  | -37 | 28 | 17 |
|  |  |  |  |  | -37 | 32 | -7 |
|  |  |  |  |  | -43 | -89 | 20 |
|  |  |  |  |  | -6 | -66 | -2 |
|  |  |  |  |  | 11 | -72 | 5 |
| Damasio et al. | 2004 | [10.1016/j.cognition.2002.07.001](https://doi.org/10.1016/j.cognition.2002.07.001) | Tools > scrambled pictures | 68 | -54 | -53 | -12 |
|  |  |  |  |  | -23 | -16 | -30 |
|  |  |  |  |  | -27 | -37 | -15 |
|  |  |  |  |  | -37 | 30 | -7 |
|  |  |  |  |  | -28 | 15 | 29 |
|  |  |  |  |  | -2 | 40 | 27 |
|  |  |  |  |  | -10 | 30 | 31 |
|  |  |  |  |  | -23 | -81 | 34 |
|  |  |  |  |  | -19 | -73 | 16 |
|  |  |  |  |  | -4 | -67 | 14 |
| Rothstein et al. | 2005 | [10.1038/nn1370](https://doi.org/10.1038/nn1370) | Familiarity & identity change effect | 20 | -45 | 0 | -39 |
|  |  |  |  |  | 63 | -6 | -33 |
|  |  |  |  |  | 24 | -12 | -27 |
|  |  |  |  |  | -6 | 18 | 69 |
| Elfgren et al. | 2006 | [10.1016/j.neuroimage.2005.09.060](https://doi.org/10.1016/j.neuroimage.2005.09.060) | Familiar > unfamiliar faces | 15 | -13 | 21 | 61 |
|  |  |  |  |  | -49 | 35 | -3 |
|  |  |  |  |  | -51 | 26 | 5 |
|  |  |  |  |  | 12 | 11 | 12 |
|  |  |  |  |  | -12 | 41 | 8 |
|  |  |  |  |  | -18 | 47 | 35 |
|  |  |  |  |  | -46 | -52 | -13 |
|  |  |  |  |  | 4 | -32 | 38 |
|  |  |  |  |  | -16 | -41 | 39 |
|  |  |  |  |  | 63 | 35 | 0 |
|  |  |  |  |  | 65 | 31 | -2 |
|  |  |  |  |  | 52 | 16 | -38 |
|  |  |  |  |  | 56 | 16 | -27 |
|  |  |  |  |  | -64 | -43 | 41 |
|  |  |  |  |  | 55 | -39 | 3 |
|  |  |  |  |  | -14 | -63 | -3 |
|  |  |  |  |  | 53 | 52 | -5 |
|  |  |  |  |  | -56 | -38 | 0 |
|  |  |  |  |  | 44 | -70 | -34 |
|  |  |  |  |  | 44 | -60 | -12 |
|  |  |  |  |  | -35 | -16 | -26 |
| Sugiura et al. | 2008 | [10.1162/jocn.2008.21150](https://doi.org/10.1162/jocn.2008.21150) | Familiar > unfamiliar names | 25 | -44 | 10 | -32 |
|  |  |  |  |  | 42 | 18 | -30 |
|  |  |  |  |  | -32 | 12 | -22 |
|  |  |  |  |  | -62 | -42 | -2 |
|  |  |  |  |  | 0 | -30 | 36 |
| Nielson et al. | 2010 | [10.1016/j.bandc.2010.01.006](https://dx.doi.org/10.1016%2Fj.bandc.2010.01.006) | Familiar > unfamiliar people | 17 | -44 | 14 | -29 |
|  |  |  |  |  | -10 | 42 | 43 |
|  |  |  |  |  | 0 | 62 | 6 |
|  |  |  |  |  | -33 | 13 | 52 |
|  |  |  |  |  | -38 | 2 | -2 |
|  |  |  |  |  | -51 | -63 | 28 |
|  |  |  |  |  | -1 | -51 | 29 |
|  |  |  |  |  | 53 | -72 | 35 |
|  |  |  |  |  | -57 | -43 | -6 |
|  |  |  |  |  | -55 | -17 | -13 |
|  |  |  |  |  | -47 | -3 | -38 |
|  |  |  |  |  | -26 | -31 | -13 |
|  |  |  |  |  | 34 | -27 | -15 |
|  |  |  |  |  | 65 | -5 | -15 |
|  |  |  |  |  | 69 | -52 | 6 |
| Brambati et al. | 2010 | [10.1016/j.neuroimage.2010.06.045](https://doi.org/10.1016/j.neuroimage.2010.06.045) | Occupation judgement > baseline | 12 | 64 | 0 | -18 |
|  |  |  |  |  | 44 | -80 | -10 |
|  |  |  |  |  | -34 | -78 | -10 |
|  |  |  |  |  | 38 | -78 | 2 |
|  |  |  |  |  | -36 | -88 | -2 |
|  |  |  |  |  | 28 | -90 | 26 |
|  |  |  |  |  | -14 | -102 | 14 |
|  |  |  |  |  | 40 | -54 | -20 |
|  |  |  |  |  | 36 | -51 | -21 |
|  |  |  |  |  | -36 | -50 | -22 |
|  |  |  |  |  | 12 | -88 | -6 |
|  |  |  |  |  | -20 | -76 | -12 |
|  |  |  |  |  | 44 | 18 | 28 |
|  |  |  |  |  | -42 | 22 | 20 |
|  |  |  |  |  | -62 | -8 | -10 |
|  |  |  |  |  | 18 | -4 | -18 |
|  |  |  |  |  | 52 | -70 | 0 |
|  |  |  |  |  | -58 | -38 | -4 |
|  |  |  |  |  | -40 | 4 | 34 |
|  |  |  |  |  | -2 | 18 | 50 |
| Barense et al. | 2011 | [10.1162/jocn_a_00010](https://doi.org/10.1162/jocn_a_00010) | Familiar > unfamiliar faces | 18 | -36 | 18 | -27 |
|  |  |  |  |  | -27 | 3 | -21 |
|  |  |  |  |  | 63 | 3 | -18 |
|  |  |  |  |  | -33 | -12 | -27 |
|  |  |  |  |  | -30 | -6 | -30 |
|  |  |  |  |  | -21 | -9 | -18 |
|  |  |  |  |  | -30 | -6 | -18 |
|  |  |  |  |  | 27 | -15 | -18 |
| Gesierich et al. | 2012 | 10.1093/cercor/bhr286 | Familiar > unfamiliar faces | 21 | -3 | -54 | 12 |
|  |  |  |  |  | -12 | 6 | 6 |
|  |  |  |  |  | -9 | -6 | 6 |
|  |  |  |  |  | -3 | -36 | 30 |
|  |  |  |  |  | -39 | 27 | 6 |
|  |  |  |  |  | 18 | 21 | -3 |
|  |  |  |  |  | -36 | 33 | -12 |
|  |  |  |  |  | -39 | 15 | -33 |
|  |  |  |  |  | -33 | -72 | 39 |
|  |  |  |  |  | -60 | -6 | -18 |
|  |  |  |  |  | 60 | -3 | -15 |
|  |  |  |  |  | -3 | 60 | -9 |
|  |  |  |  |  | 45 | -66 | 30 |
|  |  |  |  |  | -21 | 57 | 0 |
|  |  |  |  |  | -54 | -39 | -6 |
|  |  |  |  |  | 18 | -45 | -9 |
|  |  |  |  |  | 36 | -12 | -18 |
|  |  |  |  |  | 48 | 6 | -27 |
| Gesierich et al. | 2012 | [10.1093/cercor/bhr286](https://doi.org/10.1093/cercor/bhr286) | Familiar > scrambled items | 21 | 42 | -51 | -24 |
|  |  |  |  |  | 42 | -78 | -12 |
|  |  |  |  |  | 45 | -53 | 15 |
|  |  |  |  |  | -3 | -54 | 15 |
|  |  |  |  |  | -21 | -6 | -12 |
|  |  |  |  |  | -6 | -9 | 3 |
|  |  |  |  |  | 24 | -6 | -15 |
|  |  |  |  |  | 33 | -12 | -18 |
|  |  |  |  |  | -12 | 9 | 6 |
|  |  |  |  |  | -30 | -15 | -15 |
|  |  |  |  |  | 12 | 12 | 6 |
|  |  |  |  |  | -45 | 24 | 21 |
|  |  |  |  |  | -42 | -72 | -18 |
|  |  |  |  |  | -42 | -81 | -15 |
|  |  |  |  |  | -36 | -75 | 42 |
|  |  |  |  |  | 33 | 33 | -12 |
|  |  |  |  |  | -39 | 12 | -33 |
|  |  |  |  |  | 54 | -9 | -21 |
|  |  |  |  |  | -57 | -6 | -18 |
|  |  |  |  |  | 6 | 42 | -18 |
|  |  |  |  |  | 36 | 12 | -33 |
|  |  |  |  |  | 45 | 24 | 21 |
| Gesierich et al. | 2012 | [10.1093/cercor/bhr286](https://doi.org/10.1093/cercor/bhr286) | Familiar > unfamiliar items | 12 | 60 | 0 | -15 |
|  |  |  |  |  | -42 | 15 | -33 |
|  |  |  |  |  | -6 | -66 | 27 |
|  |  |  |  |  | -6 | -57 | 12 |
|  |  |  |  |  | -3 | -39 | 30 |
|  |  |  |  |  | -42 | 24 | 24 |
|  |  |  |  |  | -9 | -18 | 27 |
|  |  |  |  |  | -33 | -72 | 42 |
|  |  |  |  |  | 51 | -66 | 27 |
|  |  |  |  |  | -57 | -6 | -18 |
|  |  |  |  |  | -24 | 54 | 3 |
|  |  |  |  |  | 15 | 15 | -3 |
|  |  |  |  |  | -3 | 54 | -12 |
|  |  |  |  |  | -39 | 27 | 3 |
|  |  |  |  |  | -39 | 33 | -12 |
|  |  |  |  |  | -6 | 9 | 3 |
| Ross et al. | 2012 | 10.1093/cercor/bhr274 | Famous > unknown faces & landmarks | 16 | -40 | 21 | -28 |
|  |  |  |  |  | -4 | 59 | 19 |
|  |  |  |  |  | -2 | 60 | -5 |
|  |  |  |  |  | -5 | 42 | -11 |
|  |  |  |  |  | -3 | 7 | -5 |
|  |  |  |  |  | -22 | -7 | -17 |
|  |  |  |  |  | -61 | -11 | -8 |
|  |  |  |  |  | -36 | -35 | -23 |
|  |  |  |  |  | 5 | 49 | -6 |
|  |  |  |  |  | 5 | 50 | -7 |
| Yen | 2019 | 10.1016/j.neuroimage.2019.01.040 | Semantic > perceptual decision | 16 | -45 | 23 | 9 |
|  |  |  |  |  | -56 | -40 | 2 |
|  |  |  |  |  | -7 | 36 | 45 |
|  |  |  |  |  | 23 | -78 | -32 |
|  |  |  |  |  | -20 | -12 | -14 |
|  |  |  |  |  | 46 | 32 | -7 |
|  |  |  |  |  | 29 | -48 | 13 |
|  |  |  |  |  | -9 | -52 | 9 |
| Yen | 2019 | 10.1016/j.neuroimage.2019.01.040 | Semantic > phonological judgement | 16 | -54 | -45 | 7 |
|  |  |  |  |  | -9 | -51 | 19 |
|  |  |  |  |  | -12 | 48 | 36 |
|  |  |  |  |  | -49 | 29 | -4 |
|  |  |  |  |  | 0 | -86 | 22 |
|  |  |  |  |  | 25 | -81 | -33 |
|  |  |  |  |  | -44 | 11 | -31 |
| Yen | 2019 | 10.1016/j.neuroimage.2019.01.040 | Semantic > phonological judgement | 16 | -10 | 43 | 36 |
|  |  |  |  |  | -50 | 15 | -8 |
|  |  |  |  |  | -52 | -57 | 16 |
|  |  |  |  |  | -4 | -54 | 16 |
|  |  |  |  |  | 0 | -86 | 20 |
|  |  |  |  |  | 25 | -80 | -32 |
|  |  |  |  |  | -27 | -36 | -16 |
|  |  |  |  |  | 41 | 33 | -14 |
| Wilson | 2018 | 10.1002/hbm.24077 | Semantic > perceptual decision | 14 | -45 | 26 | 9 |
|  |  |  |  |  | -43 | -28 | -6 |
|  |  |  |  |  | 25 | -76 | -33 |
|  |  |  |  |  | -7 | 48 | 43 |
|  |  |  |  |  | 24 | -10 | -17 |
| Dreyer & Pulvermueller | 2018 | 10.1016/j.cortex.2017.10.021 | Nouns > hashmarks | 28 | -42 | 32 | -2 |
|  |  |  |  |  | -54 | -8 | 44 |
|  |  |  |  |  | -52 | 14 | -10 |
|  |  |  |  |  | -56 | -18 | 18 |
|  |  |  |  |  | -30 | -36 | 0 |
|  |  |  |  |  | -22 | 0 | -4 |
|  |  |  |  |  | -42 | -52 | -14 |
|  |  |  |  |  | -32 | -18 | -10 |
|  |  |  |  |  | -24 | -10 | -10 |
|  |  |  |  |  | -6 | -16 | 70 |
|  |  |  |  |  | 0 | -10 | 64 |
|  |  |  |  |  | -60 | -38 | 16 |
| Bulut, Hung, Tzeng & Wu | 2017 | 10.1371/journal.pone.0188526 | Sentences > word lists | 20 | 24 | -8 | -6 |
|  |  |  |  |  | -24 | -8 | -14 |
|  |  |  |  |  | -16 | 56 | 36 |
|  |  |  |  |  | 68 | -32 | 26 |
|  |  |  |  |  | -42 | 22 | -32 |
|  |  |  |  |  | 48 | -2 | 12 |
|  |  |  |  |  | -60 | -42 | 28 |
| Bulut, Hung, Tzeng & Wu | 2017 | 10.1371/journal.pone.0188526 | Sentences > characters | 20 | -48 | 18 | -24 |
|  |  |  |  |  | -16 | 56 | 36 |
|  |  |  |  |  | -24 | -6 | -18 |
|  |  |  |  |  | 22 | -4 | -18 |
| Schell, Zaccarella & Friederici | 2017 | 10.1016/j.cortex.2017.09.002 | Phrases > words | 21 | -42 | 29 | 19 |
|  |  |  |  |  | -45 | 32 | 7 |
| Perrone-Bertolotti, Kauffmann, Pichat, Vidal & Baciu | 2017 | 10.3389/fnhum.2017.00325 | Words > unreadable font | 24 | -6 | -3 | 67 |
|  |  |  |  |  | -42 | 3 | 25 |
|  |  |  |  |  | -48 | 24 | 18 |
|  |  |  |  |  | -57 | -12 | 49 |
|  |  |  |  |  | -57 | -42 | 11 |
|  |  |  |  |  | -27 | -9 | -7 |
|  |  |  |  |  | 24 | -75 | -46 |
| Liuzzi, Bruffaerts, Peeters, Adamczuk, Keuleers, De Deyne, Storms, Dupont & Vandenberghe | 2017 | 10.1016/j.neuroimage.2017.02.032 | Semantic judgement > detection | 18 | -3 | -55 | 7 |
|  |  |  |  |  | -12 | -58 | -11 |
|  |  |  |  |  | -9 | -46 | -2 |
|  |  |  |  |  | -30 | 35 | -14 |
|  |  |  |  |  | 33 | 35 | -14 |
|  |  |  |  |  | 30 | 41 | -8 |
|  |  |  |  |  | -39 | 26 | 13 |
|  |  |  |  |  | -45 | 35 | 13 |
|  |  |  |  |  | 0 | 59 | -8 |
|  |  |  |  |  | -51 | -55 | -14 |
|  |  |  |  |  | -48 | -43 | -17 |
|  |  |  |  |  | -24 | -22 | -17 |
|  |  |  |  |  | -36 | -13 | -23 |
|  |  |  |  |  | -33 | -34 | -20 |
| Matchin, Hammerly & Lau | 2017 | 10.1016/j.cortex.2016.12.010 | Sentences > word lists | 16 | -50 | 1 | -16 |
|  |  |  |  |  | -51 | 33 | 1 |
|  |  |  |  |  | -53 | 29 | 9 |
|  |  |  |  |  | -47 | 35 | -5 |
|  |  |  |  |  | -34 | 10 | -34 |
|  |  |  |  |  | 24 | -81 | -35 |
| Matchin, Hammerly & Lau | 2017 | 10.1016/j.cortex.2016.12.010 | Sentences > phrases | 16 | -50 | 31 | 7 |
|  |  |  |  |  | -54 | -41 | 2 |
|  |  |  |  |  | 25 | -72 | -36 |
|  |  |  |  |  | -26 | -4 | -15 |
| Matchin, Hammerly & Lau | 2017 | 10.1016/j.cortex.2016.12.010 | Real > pseudoword lists | 16 | -51 | -37 | 4 |
| Matchin, Hammerly & Lau | 2017 | 10.1016/j.cortex.2016.12.010 | Real > pseudoword phrases | 16 | -49 | -37 | 1 |
| Matchin, Hammerly & Lau | 2017 | 10.1016/j.cortex.2016.12.010 | Real > pseudoword sentences | 16 | -51 | -45 | 7 |
|  |  |  |  |  | -47 | 18 | -26 |
|  |  |  |  |  | -50 | 6 | -14 |
|  |  |  |  |  | -53 | -10 | -11 |
|  |  |  |  |  | -52 | 25 | -21 |
|  |  |  |  |  | -53 | -39 | 2 |
|  |  |  |  |  | -35 | -51 | 10 |
|  |  |  |  |  | -51 | -45 | 7 |
|  |  |  |  |  | 24 | -75 | -32 |
|  |  |  |  |  | -29 | -12 | -13 |
| Zhuang & Devereux | 2017 | 10.1080/23273798.2016.1241886 | Phrases > words | 16 | -34 | 34 | -2 |
|  |  |  |  |  | -50 | 12 | 12 |
|  |  |  |  |  | -40 | 22 | 20 |
|  |  |  |  |  | -58 | -20 | -2 |
|  |  |  |  |  | -48 | -40 | 2 |
|  |  |  |  |  | -44 | -46 | 10 |
| Redcay, Velnoskey & Rowe | 2016 | 10.1002/hbm.23251 | Meaningful > meaningless stimuli | 24 | 43 | -50 | 14 |
|  |  |  |  |  | -60 | -47 | 26 |
|  |  |  |  |  | -48 | 16 | -32 |
|  |  |  |  |  | -6 | -56 | 41 |
|  |  |  |  |  | -8 | 66 | 12 |
|  |  |  |  |  | 14 | -53 | 37 |
|  |  |  |  |  | 8 | 59 | 5 |
| Redcay, Velnoskey & Rowe | 2016 | 10.1002/hbm.23251 | Communicative > non-communicative gesture | 24 | -56 | -29 | 6 |
|  |  |  |  |  | 56 | -22 | 1 |
|  |  |  |  |  | -50 | -49 | 13 |
|  |  |  |  |  | -53 | 27 | 21 |
|  |  |  |  |  | 55 | 11 | -16 |
| Redcay, Velnoskey & Rowe | 2016 | 10.1002/hbm.23251 | Real > pseudoword sentences | 24 | 58 | -53 | 6 |
|  |  |  |  |  | -38 | -71 | 35 |
|  |  |  |  |  | -12 | -42 | 34 |
|  |  |  |  |  | -49 | -9 | -23 |
|  |  |  |  |  | -9 | 65 | 12 |
|  |  |  |  |  | -67 | -38 | -1 |
|  |  |  |  |  | 14 | -54 | 37 |
|  |  |  |  |  | -3 | -99 | -3 |
|  |  |  |  |  | 10 | 59 | -6 |
|  |  |  |  |  | -28 | -30 | -16 |
|  |  |  |  |  | 49 | -19 | -8 |
|  |  |  |  |  | 14 | -94 | 10 |
|  |  |  |  |  | -24 | 37 | 35 |
|  |  |  |  |  | -29 | 31 | 40 |
|  |  |  |  |  | 11 | -31 | 42 |
| Sheldon, McAndrews, Pruessner & Moscovitch | 2016 | 10.1016/j.neuropsychologia.2016.06.028 | Semantic fluency > perceptual task | 15 | -30 | -42 | -12 |
|  |  |  |  |  | 18 | -56 | 22 |
|  |  |  |  |  | 30 | -40 | -12 |
|  |  |  |  |  | -36 | -78 | 38 |
|  |  |  |  |  | 46 | -76 | 26 |
|  |  |  |  |  | -26 | 24 | 46 |
|  |  |  |  |  | 30 | 20 | 52 |
| Sheldon, McAndrews, Pruessner & Moscovitch | 2016 | 10.1016/j.neuropsychologia.2016.06.028 | Semantic > autobiographical fluency | 15 | 36 | -56 | -26 |
|  |  |  |  |  | -18 | -10 | -14 |
|  |  |  |  |  | 20 | -66 | 16 |
|  |  |  |  |  | -6 | 20 | 16 |
| Haberling, Corballis &Corballis | 2016 | 10.1016/j.cortex.2016.06.003 | Meaningful pantomimes > unknown sign language | 91 | -21 | -94 | 1 |
|  |  |  |  |  | 21 | -94 | -2 |
|  |  |  |  |  | -6 | -55 | 7 |
|  |  |  |  |  | 51 | -58 | 7 |
|  |  |  |  |  | 45 | -79 | -8 |
|  |  |  |  |  | -18 | -85 | 34 |
|  |  |  |  |  | -30 | -34 | -17 |
|  |  |  |  |  | 39 | -34 | -17 |
|  |  |  |  |  | -9 | -55 | 16 |
|  |  |  |  |  | -6 | -70 | 1 |
|  |  |  |  |  | 27 | -16 | -14 |
|  |  |  |  |  | 30 | -22 | -17 |
| Haberling, Corballis &Corballis | 2016 | 10.1016/j.cortex.2016.06.003 | Meaningful pantomimes > dog videos | 91 | -48 | 32 | 7 |
|  |  |  |  |  | 54 | 35 | 4 |
|  |  |  |  |  | -27 | -10 | 55 |
|  |  |  |  |  | 33 | -7 | 55 |
|  |  |  |  |  | 36 | -43 | 61 |
|  |  |  |  |  | -24 | -37 | 70 |
|  |  |  |  |  | -6 | -4 | 67 |
|  |  |  |  |  | 42 | -4 | 58 |
|  |  |  |  |  | -24 | -49 | 67 |
|  |  |  |  |  | -36 | -43 | 52 |
|  |  |  |  |  | 60 | -28 | 25 |
|  |  |  |  |  | -57 | -31 | 31 |
|  |  |  |  |  | -54 | -1 | -14 |
|  |  |  |  |  | 54 | -61 | 1 |
|  |  |  |  |  | -51 | -64 | 4 |
|  |  |  |  |  | -45 | -43 | -17 |
|  |  |  |  |  | 54 | 11 | -17 |
|  |  |  |  |  | 27 | -7 | -17 |
|  |  |  |  |  | -24 | -4 | -20 |
|  |  |  |  |  | -12 | -73 | -47 |
|  |  |  |  |  | 18 | -76 | -47 |
| Haberling, Corballis &Corballis | 2016 | 10.1016/j.cortex.2016.06.003 | Synonyms > letter strings | 91 | -42 | 29 | -11 |
|  |  |  |  |  | 39 | 35 | -11 |
|  |  |  |  |  | -54 | -40 | -2 |
|  |  |  |  |  | 51 | -37 | 1 |
|  |  |  |  |  | -51 | 26 | 13 |
|  |  |  |  |  | 57 | 26 | 19 |
|  |  |  |  |  | -48 | 17 | -23 |
|  |  |  |  |  | 54 | 11 | -20 |
|  |  |  |  |  | -39 | 8 | 43 |
|  |  |  |  |  | -48 | -4 | -29 |
|  |  |  |  |  | 27 | -82 | -38 |
|  |  |  |  |  | -15 | -88 | -35 |
|  |  |  |  |  | -6 | 23 | 58 |
|  |  |  |  |  | -6 | 41 | 43 |
|  |  |  |  |  | -27 | -13 | -17 |
|  |  |  |  |  | -9 | -94 | 22 |
| Wang, Zhao, Zevin & Yang | 2016 | 10.3389/fpsyg.2016.00947 | Words > nonsense strokes | 16 | -35 | 18 | 34 |
|  |  |  |  |  | -46 | 4 | 22 |
|  |  |  |  |  | -35 | 22 | 1 |
|  |  |  |  |  | 31 | 28 | 2 |
|  |  |  |  |  | -1 | 25 | 51 |
|  |  |  |  |  | -50 | -54 | -15 |
|  |  |  |  |  | -29 | -67 | 34 |
|  |  |  |  |  | -54 | -40 | 41 |
|  |  |  |  |  | -37 | -46 | 43 |
| Kumar | 2016 | 10.1007/s10339-015-0738-1 | Words > pseudowords | 20 | -50 | 11 | 21 |
|  |  |  |  |  | 48 | 19 | 34 |
|  |  |  |  |  | -46 | 30 | 30 |
|  |  |  |  |  | 46 | 30 | 21 |
|  |  |  |  |  | -38 | -84 | -2 |
|  |  |  |  |  | 26 | -98 | 6 |
|  |  |  |  |  | -36 | -68 | -12 |
|  |  |  |  |  | -24 | -56 | 44 |
| Bautista & Wilson | 2016 | 10.1080/23273798.2015.1123281 | Clear > scrambled rotated speech | 12 | -55 | -25 | -5 |
|  |  |  |  |  | 55 | -7 | -14 |
|  |  |  |  |  | 20 | -80 | -34 |
|  |  |  |  |  | 47 | -53 | 22 |
| Higuchi, Moriguchi, Murakami, Katsunuma, Mishima &Uno | 2015 | 10.1002/brb3.413 | Words > checkerboard | 28 | 42 | -66 | -10 |
|  |  |  |  |  | 40 | -82 | -8 |
|  |  |  |  |  | 48 | -52 | -14 |
|  |  |  |  |  | -44 | -58 | -12 |
|  |  |  |  |  | -40 | -78 | -6 |
|  |  |  |  |  | -42 | -40 | -10 |
| Rogalsky, Almeida, Sprouse & Hickok | 2015 | 10.1080/23273798.2015.1066831 | Words > scrambled | 15 | -56 | -7 | -13 |
|  |  |  |  |  | -54 | -37 | -3 |
|  |  |  |  |  | -46 | -63 | 20 |
|  |  |  |  |  | -51 | 24 | 6 |
|  |  |  |  |  | -5 | 13 | 5 |
|  |  |  |  |  | 46 | 14 | -34 |
|  |  |  |  |  | 43 | -69 | -13 |
| Bonhage, Mueller, Friederici & Fiebach | 2015 | 10.1016/j.cortex.2015.04.011 | Words > pseudowords | 18 | -18 | 50 | 16 |
|  |  |  |  |  | 24 | 50 | 19 |
|  |  |  |  |  | 39 | -4 | 43 |
|  |  |  |  |  | -45 | -7 | -5 |
|  |  |  |  |  | -54 | -13 | -29 |
|  |  |  |  |  | -39 | -16 | -8 |
|  |  |  |  |  | -21 | -1 | -14 |
|  |  |  |  |  | -27 | -31 | -2 |
|  |  |  |  |  | 57 | 8 | 1 |
|  |  |  |  |  | 66 | -19 | -20 |
|  |  |  |  |  | 42 | -2 | -32 |
|  |  |  |  |  | 51 | -13 | -29 |
|  |  |  |  |  | 42 | -10 | -5 |
|  |  |  |  |  | 21 | -13 | -2 |
|  |  |  |  |  | 18 | 11 | -2 |
|  |  |  |  |  | 39 | -4 | -26 |
|  |  |  |  |  | 24 | -25 | -11 |
|  |  |  |  |  | -15 | -31 | 37 |
|  |  |  |  |  | 18 | -34 | 1 |
|  |  |  |  |  | 30 | -34 | -11 |
|  |  |  |  |  | 33 | -46 | -11 |
|  |  |  |  |  | 18 | -58 | -5 |
|  |  |  |  |  | 12 | -40 | -8 |
|  |  |  |  |  | -51 | -52 | 31 |
|  |  |  |  |  | 51 | -46 | 28 |
|  |  |  |  |  | -48 | -58 | 1 |
|  |  |  |  |  | 45 | -55 | 7 |
|  |  |  |  |  | -30 | -55 | -8 |
|  |  |  |  |  | 39 | -67 | 16 |
|  |  |  |  |  | -39 | -79 | 16 |
|  |  |  |  |  | -18 | 11 | 4 |
|  |  |  |  |  | -12 | -16 | 34 |
|  |  |  |  |  | 9 | -28 | 40 |
|  |  |  |  |  | 6 | -43 | 43 |
|  |  |  |  |  | -3 | -22 | 58 |
|  |  |  |  |  | 0 | -20 | 5 |
|  |  |  |  |  | 12 | -28 | 7 |
|  |  |  |  |  | -12 | -31 | 4 |
|  |  |  |  |  | -3 | -40 | -20 |
| AbdulSabur, Xu, Liu, Chow, Baxter, Carson & Braun | 2014 | 10.1016/j.cortex.2014.01.017 | Production & semantic > phonological judgement | 18 | -54 | 20 | 15 |
|  |  |  |  |  | -48 | 26 | -3 |
|  |  |  |  |  | -51 | -1 | -21 |
|  |  |  |  |  | -60 | -49 | 0 |
|  |  |  |  |  | -9 | 14 | 63 |
|  |  |  |  |  | -39 | 5 | 51 |
|  |  |  |  |  | -45 | -64 | 18 |
|  |  |  |  |  | -12 | 50 | 33 |
|  |  |  |  |  | -3 | -52 | 39 |
|  |  |  |  |  | -24 | -37 | -18 |
|  |  |  |  |  | -33 | -37 | -21 |
|  |  |  |  |  | 54 | 2 | -21 |
|  |  |  |  |  | 42 | -61 | 18 |
|  |  |  |  |  | 9 | 17 | 63 |
|  |  |  |  |  | 18 | -79 | -45 |
| AbdulSabur, Xu, Liu, Chow, Baxter, Carson & Braun | 2014 | 10.1016/j.cortex.2014.01.017 | Production > phonological decision | 18 | -30 | 29 | -3 |
|  |  |  |  |  | -12 | 35 | 48 |
|  |  |  |  |  | -24 | 47 | 21 |
|  |  |  |  |  | -42 | 20 | 33 |
|  |  |  |  |  | -6 | 14 | 54 |
|  |  |  |  |  | -12 | 29 | 24 |
|  |  |  |  |  | -6 | -58 | 45 |
|  |  |  |  |  | -3 | -61 | 6 |
|  |  |  |  |  | -15 | 8 | 15 |
|  |  |  |  |  | -3 | -16 | 12 |
|  |  |  |  |  | -36 | -61 | -30 |
|  |  |  |  |  | 9 | -57 | 12 |
|  |  |  |  |  | 12 | -55 | 45 |
|  |  |  |  |  | 42 | -76 | 30 |
|  |  |  |  |  | 24 | -55 | 0 |
|  |  |  |  |  | 39 | -58 | -33 |
|  |  |  |  |  | 3 | -76 | -18 |
| AbdulSabur, Xu, Liu, Chow, Baxter, Carson & Braun | 2014 | 10.1016/j.cortex.2014.01.017 | Semantic > phonological decision | 18 | -42 | 2 | -42 |
|  |  |  |  |  | -57 | -4 | -9 |
|  |  |  |  |  | -33 | -46 | 18 |
|  |  |  |  |  | -21 | -4 | -21 |
|  |  |  |  |  | -27 | -16 | -15 |
|  |  |  |  |  | -21 | -76 | -39 |
|  |  |  |  |  | 51 | 26 | -3 |
|  |  |  |  |  | 57 | -10 | -9 |
|  |  |  |  |  | 57 | -40 | -3 |
|  |  |  |  |  | 60 | -58 | 21 |
|  |  |  |  |  | 6 | 44 | 36 |
|  |  |  |  |  | 24 | -4 | -21 |
| Kyong, Scott, Rosen, Howe, Agnew & McGettigan | 2014 | 10.1162/jocn_a_00583 | Intelligibility parameter | 19 | -58 | -14 | -2 |
|  |  |  |  |  | -50 | -42 | 6 |
|  |  |  |  |  | -66 | -22 | 2 |
|  |  |  |  |  | 62 | -8 | -2 |
|  |  |  |  |  | 66 | -22 | 2 |
|  |  |  |  |  | 52 | -34 | 0 |
| Kyong, Scott, Rosen, Howe, Agnew & McGettigan | 2014 | 10.1162/jocn_a_00583 | Intelligibility parameter | 19 | -60 | -10 | -4 |
|  |  |  |  |  | -54 | -42 | 4 |
|  |  |  |  |  | -66 | -24 | 2 |
|  |  |  |  |  | 60 | -8 | -8 |
|  |  |  |  |  | 64 | -16 | -2 |
| Kyong, Scott, Rosen, Howe, Agnew & McGettigan | 2014 | 10.1162/jocn_a_00583 | Intelligible vocoded > inverted vocoded speech | 19 | -54 | -40 | 4 |
|  |  |  |  |  | -62 | -12 | -6 |
|  |  |  |  |  | -58 | -8 | -12 |
| Slioussar, Kireev, Chernigovskaya, Kataeva, Korotkov & Medvedev | 2014 | 10.1016/j.bandl.2014.01.006 | Real verbs > pseudowords | 21 | -51 | -64 | 28 |
|  |  |  |  |  | -30 | 29 | 49 |
|  |  |  |  |  | -9 | -49 | 37 |
|  |  |  |  |  | 51 | -58 | 22 |
| Slioussar, Kireev, Chernigovskaya, Kataeva, Korotkov & Medvedev | 2014 | 10.1016/j.bandl.2014.01.006 | Real nouns > pseudowords | 21 | -6 | -52 | 31 |
|  |  |  |  |  | -48 | -64 | 25 |
|  |  |  |  |  | 51 | -58 | 25 |
| Bruffaerts, Dupont, Peeters, De Deyne, Storms & Vandenberghe | 2013 | 10.1523/JNEUROSCI.1548-13.2013 | Real > scrambled pictures & words | 19 | -36 | -43 | -23 |
|  |  |  |  |  | -36 | -22 | -23 |
|  |  |  |  |  | -9 | -58 | 10 |
|  |  |  |  |  | -36 | -10 | -35 |
|  |  |  |  |  | -30 | -37 | -5 |
|  |  |  |  |  | -45 | -55 | -20 |
|  |  |  |  |  | -15 | -37 | 1 |
|  |  |  |  |  | -30 | -1 | -23 |
|  |  |  |  |  | -3 | -46 | 25 |
|  |  |  |  |  | 9 | -49 | 1 |
|  |  |  |  |  | 27 | -16 | -17 |
|  |  |  |  |  | 15 | -28 | -11 |
|  |  |  |  |  | 36 | -37 | -20 |
|  |  |  |  |  | 6 | 2 | -5 |
|  |  |  |  |  | -39 | 32 | 10 |
|  |  |  |  |  | -30 | 35 | -17 |
|  |  |  |  |  | -45 | 29 | -14 |
|  |  |  |  |  | -9 | 17 | 49 |
|  |  |  |  |  | 0 | 17 | 64 |
|  |  |  |  |  | -48 | -73 | 28 |
|  |  |  |  |  | 39 | 32 | -17 |
|  |  |  |  |  | -9 | 50 | 46 |
|  |  |  |  |  | -12 | 32 | 49 |
| Ludersdorfer, Schurz, Richlan, Kronbichler & Wimmer | 2013 | 10.3389/fnhum.2013.00491 | Words > false fonts | 29 | -9 | -97 | 16 |
|  |  |  |  |  | -21 | -43 | 64 |
|  |  |  |  |  | 15 | -94 | 19 |
|  |  |  |  |  | -9 | -55 | 19 |
|  |  |  |  |  | 15 | -49 | 58 |
|  |  |  |  |  | -6 | -4 | 40 |
|  |  |  |  |  | 51 | -31 | 25 |
|  |  |  |  |  | -48 | -67 | 19 |
|  |  |  |  |  | -60 | -31 | 16 |
| Ludersdorfer, Schurz, Richlan, Kronbichler & Wimmer | 2013 | 10.3389/fnhum.2013.00491 | Words > pseudowords | 29 | -51 | -58 | 22 |
| Ludersdorfer, Schurz, Richlan, Kronbichler & Wimmer | 2013 | 10.3389/fnhum.2013.00491 | Speech > reversed speech | 29 | -45 | -58 | -11 |
| Ludersdorfer, Schurz, Richlan, Kronbichler & Wimmer | 2013 | 10.3389/fnhum.2013.00491 | Words > pseudowords | 29 | -48 | -64 | 28 |
| Simard, Monetta, Nagano-Saito & Monchi | 2013 | 10.1016/j.bandl.2011.08.002 | Semantics > baseline | 14 | -44 | -57 | -3 |
|  |  |  |  |  | -7 | 43 | 23 |
|  |  |  |  |  | 12 | 43 | 22 |
|  |  |  |  |  | -31 | 34 | -3 |
|  |  |  |  |  | 40 | 31 | -8 |
|  |  |  |  |  | -55 | 34 | 24 |
|  |  |  |  |  | 40 | 42 | 17 |
|  |  |  |  |  | -51 | 33 | 15 |
|  |  |  |  |  | -3 | 28 | 49 |
|  |  |  |  |  | -50 | 14 | 44 |
|  |  |  |  |  | -1 | -32 | 28 |
|  |  |  |  |  | -48 | -65 | -4 |
|  |  |  |  |  | -26 | -60 | 49 |
|  |  |  |  |  | 32 | -65 | 60 |
|  |  |  |  |  | -31 | -74 | -8 |
|  |  |  |  |  | 25 | -72 | -7 |
|  |  |  |  |  | -40 | -85 | -5 |
|  |  |  |  |  | 34 | -90 | -1 |
|  |  |  |  |  | -7 | -88 | 9 |
|  |  |  |  |  | 10 | -87 | 13 |
|  |  |  |  |  | -5 | -12 | 8 |
|  |  |  |  |  | -27 | -34 | 6 |
|  |  |  |  |  | 25 | -28 | -2 |
|  |  |  |  |  | 10 | -12 | 8 |
|  |  |  |  |  | -12 | 1 | 13 |
|  |  |  |  |  | 14 | 10 | -4 |
|  |  |  |  |  | -40 | -68 | -29 |
|  |  |  |  |  | 38 | -79 | -18 |
| Simard, Monetta, Nagano-Saito & Monchi | 2013 | 10.1016/j.bandl.2011.08.002 | Semantic > phonological decision | 14 | -60 | 33 | -11 |
|  |  |  |  |  | -62 | 36 | -3 |
|  |  |  |  |  | -57 | 40 | 19 |
|  |  |  |  |  | -31 | -40 | -14 |
|  |  |  |  |  | -40 | -18 | -29 |
|  |  |  |  |  | -14 | -99 | 5 |
|  |  |  |  |  | 18 | 17 | -4 |
| Simard, Monetta, Nagano-Saito & Monchi | 2013 | 10.1016/j.bandl.2011.08.002 | Semantic > phonological decision | 14 | -31 | 28 | 9 |
|  |  |  |  |  | -40 | 32 | -3 |
|  |  |  |  |  | -20 | -50 | -6 |
|  |  |  |  |  | -44 | -33 | -23 |
|  |  |  |  |  | 6 | -74 | 12 |
|  |  |  |  |  | 21 | -97 | 20 |
| Straube, Green, Weis & Kircher | 2012 | 10.1371/journal.pone.0051207 | Known > unknown language | 16 | -64 | -52 | 0 |
|  |  |  |  |  | -44 | 12 | 24 |
|  |  |  |  |  | 60 | -40 | -4 |
|  |  |  |  |  | -8 | 20 | 48 |
| Straube, Green, Weis & Kircher | 2012 | 10.1371/journal.pone.0051207 | Iconic > meaningless gesture | 16 | -4 | -8 | 72 |
|  |  |  |  |  | -32 | 28 | -8 |
|  |  |  |  |  | -56 | -40 | 40 |
|  |  |  |  |  | 36 | 8 | -28 |
|  |  |  |  |  | 48 | 36 | -16 |
|  |  |  |  |  | -32 | -28 | -24 |
|  |  |  |  |  | -8 | -24 | 0 |
|  |  |  |  |  | -8 | 16 | -20 |
|  |  |  |  |  | 28 | -20 | -24 |
|  |  |  |  |  | 56 | -36 | 40 |
|  |  |  |  |  | 40 | 12 | 56 |
|  |  |  |  |  | 20 | 60 | 28 |
|  |  |  |  |  | 28 | -80 | -12 |
| Wende, Straube, Stratmann, Sommer, Kircher & Nagels | 2012 | 10.1016/j.neuroimage.2012.06.003 | Semantic > phonological fluency | 18 | -38 | 14 | 54 |
|  |  |  |  |  | -46 | -66 | 30 |
|  |  |  |  |  | 26 | -80 | -28 |
|  |  |  |  |  | -52 | 26 | 10 |
|  |  |  |  |  | -54 | -10 | -26 |
|  |  |  |  |  | 64 | -12 | 0 |
|  |  |  |  |  | 30 | -22 | 30 |
|  |  |  |  |  | -2 | 50 | -14 |
|  |  |  |  |  | -26 | -16 | 28 |
|  |  |  |  |  | -54 | -20 | 2 |
| Zekveld, Rudner, Johnsrude, Heslenfeld & Ronnberg | 2012 | 10.1016/j.bandl.2012.05.006 | More > less intelligible speech | 18 | -38 | 22 | -4 |
|  |  |  |  |  | -50 | 0 | 48 |
|  |  |  |  |  | -44 | 2 | 36 |
|  |  |  |  |  | 0 | 16 | 54 |
|  |  |  |  |  | -4 | 10 | 66 |
|  |  |  |  |  | 8 | 22 | 46 |
|  |  |  |  |  | 10 | 32 | 26 |
|  |  |  |  |  | 42 | 20 | -6 |
|  |  |  |  |  | 36 | 24 | 0 |
|  |  |  |  |  | 50 | 20 | 4 |
|  |  |  |  |  | 54 | 26 | 24 |
|  |  |  |  |  | 48 | 14 | 28 |
|  |  |  |  |  | 2 | 34 | 40 |
|  |  |  |  |  | -38 | 28 | 26 |
|  |  |  |  |  | -46 | 24 | 30 |
| Zekveld, Rudner, Johnsrude, Heslenfeld & Ronnberg | 2012 | 10.1016/j.bandl.2012.05.006 | More > less intelligible speech based on cue | 18 | -44 | 42 | -8 |
|  |  |  |  |  | -54 | 22 | 20 |
|  |  |  |  |  | -38 | 12 | 50 |
|  |  |  |  |  | -58 | -34 | 0 |
| Zekveld, Rudner, Johnsrude, Heslenfeld & Ronnberg | 2012 | 10.1016/j.bandl.2012.05.006 | More > less intelligible speech based on cue | 18 | -60 | -34 | -4 |
| Visser, Jefferies, Embleton & Lambon Ralph | 2012 | 10.1162/jocn_a_00244 | Semantics > baseline | 15 | -57 | -42 | -3 |
|  |  |  |  |  | 42 | 21 | -33 |
|  |  |  |  |  | -54 | 27 | 6 |
|  |  |  |  |  | -57 | -15 | -24 |
| Visser, Jefferies, Embleton & Lambon Ralph | 2012 | 10.1162/jocn_a_00244 | Pictures > baseline | 15 | -57 | -45 | -3 |
|  |  |  |  |  | 18 | -78 | -30 |
| Visser, Jefferies, Embleton & Lambon Ralph | 2012 | 10.1162/jocn_a_00244 | Words > baseline | 15 | -51 | -30 | -3 |
|  |  |  |  |  | -54 | 33 | 3 |
|  |  |  |  |  | 39 | 21 | -33 |
|  |  |  |  |  | -45 | 9 | -33 |
|  |  |  |  |  | -39 | -39 | -24 |
|  |  |  |  |  | -42 | -69 | -36 |
|  |  |  |  |  | -63 | -21 | -21 |
| Abraham, Pieritz, Thybusch, Rutter, Kroeger, Schweckendiek, Stark, Windmann & Hermann | 2012 | 10.1016/j.neuropsychologia.2012.04.015 | Semantic > working memory task | 19 | -17 | 58 | 7 |
|  |  |  |  |  | -17 | 33 | 43 |
|  |  |  |  |  | -50 | 14 | -12 |
|  |  |  |  |  | -37 | 16 | -26 |
|  |  |  |  |  | -4 | 39 | -25 |
|  |  |  |  |  | -11 | 53 | 24 |
|  |  |  |  |  | 3 | 0 | 32 |
|  |  |  |  |  | 28 | -15 | -24 |
|  |  |  |  |  | -21 | -34 | -14 |
|  |  |  |  |  | 25 | -47 | -14 |
|  |  |  |  |  | -24 | -47 | -16 |
|  |  |  |  |  | -4 | -39 | 33 |
|  |  |  |  |  | 12 | -61 | 8 |
|  |  |  |  |  | -11 | -58 | 5 |
|  |  |  |  |  | -28 | -14 | -40 |
|  |  |  |  |  | -56 | -18 | -19 |
|  |  |  |  |  | -53 | -53 | -9 |
|  |  |  |  |  | -39 | -74 | 40 |
|  |  |  |  |  | 12 | -99 | 12 |
|  |  |  |  |  | -7 | -80 | 4 |
|  |  |  |  |  | -26 | -93 | 42 |
|  |  |  |  |  | -4 | -16 | 4 |
|  |  |  |  |  | -50 | -70 | -20 |
|  |  |  |  |  | 35 | -92 | -23 |
|  |  |  |  |  | -33 | -21 | 22 |
|  |  |  |  |  | -37 | -4 | -34 |
|  |  |  |  |  | -37 | -8 | -47 |
|  |  |  |  |  | 38 | -16 | 3 |
|  |  |  |  |  | 25 | -34 | 19 |
|  |  |  |  |  | 41 | 12 | -40 |
|  |  |  |  |  | 31 | -15 | -54 |
|  |  |  |  |  | 8 | -69 | -48 |
|  |  |  |  |  | -33 | -25 | 49 |
|  |  |  |  |  | -26 | -29 | 73 |
|  |  |  |  |  | 22 | -27 | 58 |
| Zhang, Xiao & Weng | 2012 | 10.1016/j.ijpsycho.2012.02.013 | Words > pseudowords | 14 | 1 | 51 | -12 |
|  |  |  |  |  | -20 | 63 | 2 |
|  |  |  |  |  | -28 | 42 | 26 |
|  |  |  |  |  | -55 | -52 | 69 |
|  |  |  |  |  | 66 | -49 | 27 |
|  |  |  |  |  | -4 | -50 | 33 |
| Szlachta, Bozic, Jelowicka & Marslen-Wilson | 2012 | 10.1016/j.bandl.2012.02.007 | Words > musical rain | 21 | -60 | -8 | -8 |
|  |  |  |  |  | -62 | -34 | 4 |
|  |  |  |  |  | -42 | 20 | -28 |
|  |  |  |  |  | -50 | 10 | -16 |
|  |  |  |  |  | -44 | 32 | -8 |
|  |  |  |  |  | -46 | -40 | 8 |
|  |  |  |  |  | -46 | 34 | 6 |
|  |  |  |  |  | 60 | 6 | -8 |
|  |  |  |  |  | 62 | -16 | -2 |
|  |  |  |  |  | 52 | 18 | -20 |
|  |  |  |  |  | 56 | -26 | -2 |
|  |  |  |  |  | -30 | -34 | -24 |
|  |  |  |  |  | -48 | -62 | -22 |
|  |  |  |  |  | -38 | -46 | -20 |
| Szlachta, Bozic, Jelowicka & Marslen-Wilson | 2012 | 10.1016/j.bandl.2012.02.007 | Nouns > musical rain | 21 | -60 | -8 | -2 |
|  |  |  |  |  | -62 | -32 | 6 |
|  |  |  |  |  | -42 | 20 | -26 |
|  |  |  |  |  | -36 | 34 | -16 |
|  |  |  |  |  | -44 | 34 | 4 |
|  |  |  |  |  | 58 | 6 | -8 |
|  |  |  |  |  | 58 | -14 | -4 |
|  |  |  |  |  | 48 | -32 | 2 |
|  |  |  |  |  | -28 | -34 | -20 |
|  |  |  |  |  | -46 | -62 | -22 |
|  |  |  |  |  | -40 | -48 | -18 |
|  |  |  |  |  | 52 | -62 | 24 |
|  |  |  |  |  | 36 | -54 | 20 |
|  |  |  |  |  | -42 | -70 | 24 |
| Szlachta, Bozic, Jelowicka & Marslen-Wilson | 2012 | 10.1016/j.bandl.2012.02.007 | Inflected nouns > musical rain | 21 | -62 | -6 | -6 |
|  |  |  |  |  | -62 | -36 | 6 |
|  |  |  |  |  | -58 | 4 | -14 |
|  |  |  |  |  | -54 | 10 | -18 |
|  |  |  |  |  | 64 | -14 | -2 |
|  |  |  |  |  | 60 | 2 | -8 |
|  |  |  |  |  | 54 | 16 | -20 |
|  |  |  |  |  | 46 | -32 | 4 |
|  |  |  |  |  | 50 | 18 | -24 |
|  |  |  |  |  | -34 | 30 | -18 |
|  |  |  |  |  | -42 | 30 | -12 |
|  |  |  |  |  | -44 | 34 | 0 |
| Carota, Moseley & Pulvermueller | 2012 | 10.1162/jocn_a_00219 | Words > hashmarks | 18 | -48 | 42 | 6 |
|  |  |  |  |  | -34 | 30 | -16 |
|  |  |  |  |  | -24 | 32 | -16 |
|  |  |  |  |  | -46 | -8 | 46 |
|  |  |  |  |  | -56 | 4 | 36 |
|  |  |  |  |  | -58 | -12 | 22 |
|  |  |  |  |  | -58 | 6 | 20 |
|  |  |  |  |  | -60 | 6 | 28 |
|  |  |  |  |  | -30 | -8 | 54 |
|  |  |  |  |  | 34 | 36 | -12 |
|  |  |  |  |  | 60 | 2 | 38 |
|  |  |  |  |  | 34 | -8 | 56 |
|  |  |  |  |  | 32 | -18 | 54 |
|  |  |  |  |  | -56 | 10 | -10 |
|  |  |  |  |  | -58 | -38 | 2 |
|  |  |  |  |  | -46 | -40 | -16 |
|  |  |  |  |  | -46 | -60 | -10 |
|  |  |  |  |  | -38 | -34 | -20 |
|  |  |  |  |  | -38 | -12 | -26 |
|  |  |  |  |  | -56 | -34 | 40 |
|  |  |  |  |  | -48 | -30 | 38 |
|  |  |  |  |  | -44 | -26 | 50 |
|  |  |  |  |  | -52 | -40 | 26 |
|  |  |  |  |  | -64 | -20 | 34 |
|  |  |  |  |  | 38 | -50 | -30 |
|  |  |  |  |  | 8 | -54 | -6 |
|  |  |  |  |  | 20 | 8 | 2 |
| Carota, Moseley & Pulvermueller | 2012 | 10.1162/jocn_a_00219 | Tool words > hashmarks | 18 | -48 | 38 | 6 |
|  |  |  |  |  | 52 | 0 | 14 |
|  |  |  |  |  | 44 | 2 | 16 |
|  |  |  |  |  | -30 | 30 | 4 |
|  |  |  |  |  | -56 | 4 | 16 |
|  |  |  |  |  | -30 | -8 | 52 |
|  |  |  |  |  | -46 | -8 | 44 |
|  |  |  |  |  | -54 | -2 | 44 |
|  |  |  |  |  | 32 | -8 | 56 |
|  |  |  |  |  | -50 | -28 | 4 |
|  |  |  |  |  | -56 | 10 | -10 |
|  |  |  |  |  | -56 | -36 | 2 |
|  |  |  |  |  | -40 | -32 | -14 |
|  |  |  |  |  | -46 | -40 | -16 |
|  |  |  |  |  | -38 | -44 | -24 |
|  |  |  |  |  | 38 | -48 | -32 |
|  |  |  |  |  | -26 | 12 | 6 |
|  |  |  |  |  | -20 | 4 | 12 |
|  |  |  |  |  | -52 | -38 | 26 |
|  |  |  |  |  | -54 | -26 | 32 |
|  |  |  |  |  | -50 | -32 | 36 |
|  |  |  |  |  | 38 | -48 | -32 |
| Carota, Moseley & Pulvermueller | 2012 | 10.1162/jocn_a_00219 | Animal words > hashmarks | 18 | -26 | 32 | -12 |
|  |  |  |  |  | -46 | 40 | 4 |
|  |  |  |  |  | 36 | 50 | 18 |
|  |  |  |  |  | -58 | -36 | 0 |
|  |  |  |  |  | -64 | -40 | 4 |
|  |  |  |  |  | -42 | 30 | 20 |
|  |  |  |  |  | -46 | -24 | 20 |
|  |  |  |  |  | -38 | -44 | -22 |
|  |  |  |  |  | -38 | -34 | -20 |
|  |  |  |  |  | -34 | -68 | 30 |
|  |  |  |  |  | -30 | -78 | 36 |
|  |  |  |  |  | 18 | -80 | 24 |
| Carota, Moseley & Pulvermueller | 2012 | 10.1162/jocn_a_00219 | Food words > hashmarks | 18 | -34 | 30 | -16 |
|  |  |  |  |  | -34 | 10 | 30 |
|  |  |  |  |  | -44 | 10 | 28 |
|  |  |  |  |  | -48 | 40 | 6 |
|  |  |  |  |  | -50 | -10 | 44 |
|  |  |  |  |  | -56 | -10 | 44 |
|  |  |  |  |  | -38 | -20 | 42 |
|  |  |  |  |  | 58 | 4 | 40 |
|  |  |  |  |  | 34 | -18 | 52 |
|  |  |  |  |  | -56 | -38 | 2 |
|  |  |  |  |  | -42 | -12 | -26 |
|  |  |  |  |  | -44 | -60 | -10 |
|  |  |  |  |  | -46 | -40 | -14 |
|  |  |  |  |  | -38 | -34 | -20 |
| Pulvermueller, Cook & Hauk | 2012 | 10.1016/j.neuroimage.2011.12.020 | Phrases > hashmarks | 23 | 42 | -52 | -18 |
|  |  |  |  |  | -56 | -26 | -2 |
|  |  |  |  |  | -58 | -4 | -14 |
|  |  |  |  |  | -38 | 86 | 16 |
|  |  |  |  |  | -48 | 34 | 8 |
|  |  |  |  |  | -50 | 18 | 24 |
|  |  |  |  |  | -46 | 0 | 50 |
|  |  |  |  |  | -20 | -18 | 60 |
|  |  |  |  |  | -8 | 0 | 56 |
|  |  |  |  |  | 66 | -28 | -4 |
|  |  |  |  |  | 58 | -10 | 46 |
|  |  |  |  |  | 12 | -12 | 60 |
| Pulvermueller, Cook & Hauk | 2012 | 10.1016/j.neuroimage.2011.12.020 | Uninflected words > hashmarks | 23 | -44 | -56 | -18 |
|  |  |  |  |  | -38 | -86 | -16 |
|  |  |  |  |  | -42 | -44 | 0 |
|  |  |  |  |  | -56 | 0 | -14 |
|  |  |  |  |  | 44 | -34 | 0 |
|  |  |  |  |  | 68 | -30 | 0 |
|  |  |  |  |  | -46 | 32 | 10 |
|  |  |  |  |  | -42 | 18 | 24 |
|  |  |  |  |  | -50 | 18 | 20 |
|  |  |  |  |  | -4 | 0 | 56 |
|  |  |  |  |  | -46 | 0 | 50 |
|  |  |  |  |  | -56 | -4 | 40 |
|  |  |  |  |  | -44 | -2 | 36 |
|  |  |  |  |  | -18 | -18 | 60 |
|  |  |  |  |  | 58 | -10 | 46 |
|  |  |  |  |  | 46 | -10 | 52 |
|  |  |  |  |  | -22 | -44 | -32 |
|  |  |  |  |  | -20 | -58 | -34 |
|  |  |  |  |  | -32 | 0 | -22 |
|  |  |  |  |  | -26 | -8 | -18 |
| Pulvermueller, Cook & Hauk | 2012 | 10.1016/j.neuroimage.2011.12.020 | Inflected words > hashmarks | 23 | -56 | -26 | -2 |
|  |  |  |  |  | -50 | -46 | 6 |
|  |  |  |  |  | -44 | 30 | 0 |
|  |  |  |  |  | -48 | 18 | 22 |
|  |  |  |  |  | -42 | -52 | -18 |
|  |  |  |  |  | -8 | 4 | 54 |
| Geranmayeh, Brownsett, Leech, Beckmann, Woodhead & Wise | 2012 | 10.1016/j.bandl.2012.02.005 | Speech > tongue movements | 19 | -8 | 22 | 32 |
|  |  |  |  |  | 10 | 18 | 36 |
|  |  |  |  |  | -4 | 16 | 52 |
|  |  |  |  |  | 6 | 14 | 56 |
|  |  |  |  |  | -50 | 16 | 10 |
|  |  |  |  |  | -34 | 14 | -6 |
|  |  |  |  |  | 34 | 12 | -10 |
|  |  |  |  |  | 28 | 8 | -6 |
|  |  |  |  |  | -52 | -22 | -4 |
|  |  |  |  |  | 48 | -30 | -4 |
| Welcome & Joanisse | 2012 | 10.1016/j.bandl.2011.12.011 | Semantic > phonological/orthographic decision | 20 | -57 | -36 | 1 |
|  |  |  |  |  | -44 | 25 | 24 |
|  |  |  |  |  | -2 | 31 | 50 |
|  |  |  |  |  | 42 | 22 | 31 |
|  |  |  |  |  | 36 | 63 | 6 |
|  |  |  |  |  | -46 | -62 | 50 |
|  |  |  |  |  | 43 | -61 | 45 |
| Hervais-Adelman, Carlyon, Johnsrude & Davis | 2012 | 10.1080/01690965.2012.662280 | Clear > vocoded speech | 15 | 64 | -22 | 0 |
|  |  |  |  |  | 64 | -6 | -2 |
|  |  |  |  |  | 50 | -30 | 2 |
|  |  |  |  |  | -60 | -18 | 0 |
|  |  |  |  |  | -54 | 4 | -12 |
|  |  |  |  |  | -60 | -38 | 4 |
|  |  |  |  |  | -38 | -44 | -14 |
|  |  |  |  |  | -38 | -24 | -16 |
|  |  |  |  |  | -18 | -6 | -12 |
|  |  |  |  |  | -30 | -2 | -18 |
|  |  |  |  |  | -24 | 6 | -14 |
| Hervais-Adelman, Carlyon, Johnsrude & Davis | 2012 | 10.1080/01690965.2012.662280 | More > less intelligible based on cue | 15 | -34 | 0 | 38 |
| Hauk & Pulvermueller | 2011 | 10.3389/fnhum.2011.00149 | Words > hashmarks | 21 | -40 | -38 | -20 |
|  |  |  |  |  | -48 | -42 | -16 |
|  |  |  |  |  | -50 | 30 | 4 |
|  |  |  |  |  | -56 | -66 | 6 |
|  |  |  |  |  | -64 | -30 | 2 |
|  |  |  |  |  | -62 | -38 | 6 |
|  |  |  |  |  | -50 | -8 | 50 |
|  |  |  |  |  | -58 | -18 | 22 |
|  |  |  |  |  | -38 | 32 | -14 |
|  |  |  |  |  | 56 | -30 | 2 |
| Friederici, Kotz, Scott & Obleser | 2010 | 10.1002/hbm.20878 | Intelligible > rotated speech | 17 | -58 | -4 | 4 |
|  |  |  |  |  | 62 | -4 | -14 |
| Birn, Kenworthy, Case, Caravella, Jones, Bandettin & Martin | 2010 | 10.1016/j.neuroimage.2009.07.036 | Fluency > automatic speech | 14 | -41 | 27 | 20 |
|  |  |  |  |  | -48 | -54 | -11 |
|  |  |  |  |  | -6 | 0 | 10 |
|  |  |  |  |  | -2 | 23 | 44 |
|  |  |  |  |  | -35 | 2 | 49 |
|  |  |  |  |  | -32 | -66 | 37 |
|  |  |  |  |  | -4 | -27 | -6 |
| Birn, Kenworthy, Case, Caravella, Jones, Bandettin & Martin | 2010 | 10.1016/j.neuroimage.2009.07.036 | Category > letter fluency | 14 | -8 | -64 | 10 |
|  |  |  |  |  | -21 | 18 | 44 |
|  |  |  |  |  | -50 | -66 | 30 |
|  |  |  |  |  | -15 | -52 | 0 |
|  |  |  |  |  | -38 | -26 | 12 |
| Friese, Rutschmann, Raabe & Schmalhofer | 2008 | 10.1162/jocn.2008.20141 | Words > pseudowords | 40 | -58 | -12 | -14 |
|  |  |  |  |  | -10 | 56 | 32 |
|  |  |  |  |  | 56 | 0 | -18 |
|  |  |  |  |  | 2 | 56 | -14 |
|  |  |  |  |  | -20 | -52 | 14 |
| Seghier, Josse, Leff & Price | 2011 | 10.1093/cercor/bhq203 | Meaningful > meaningless stimuli | 60 | -34 | 32 | -14 |
|  |  |  |  |  | -42 | 26 | -14 |
|  |  |  |  |  | -52 | 30 | -2 |
|  |  |  |  |  | -44 | 48 | -8 |
|  |  |  |  |  | 34 | 34 | -12 |
|  |  |  |  |  | -50 | 16 | 28 |
|  |  |  |  |  | -44 | 28 | 16 |
|  |  |  |  |  | -54 | 28 | 18 |
|  |  |  |  |  | -52 | 14 | 18 |
|  |  |  |  |  | 44 | 20 | 26 |
|  |  |  |  |  | 54 | 22 | 30 |
|  |  |  |  |  | 46 | 32 | 10 |
|  |  |  |  |  | -48 | -48 | -14 |
|  |  |  |  |  | -54 | -38 | 2 |
|  |  |  |  |  | -60 | -44 | -8 |
|  |  |  |  |  | -30 | -66 | 42 |
|  |  |  |  |  | -48 | -68 | 22 |
|  |  |  |  |  | 10 | -82 | -34 |
|  |  |  |  |  | 28 | -74 | -44 |
|  |  |  |  |  | 40 | -72 | -38 |
|  |  |  |  |  | -2 | 14 | 52 |
|  |  |  |  |  | -2 | 26 | 48 |
|  |  |  |  |  | -2 | 36 | 44 |
|  |  |  |  |  | -36 | -40 | -20 |
|  |  |  |  |  | -28 | -32 | -22 |
|  |  |  |  |  | -30 | 26 | 4 |
|  |  |  |  |  | -36 | 28 | 0 |
|  |  |  |  |  | 32 | 24 | -2 |
|  |  |  |  |  | 44 | 24 | -6 |
|  |  |  |  |  | -14 | -14 | 12 |
|  |  |  |  |  | -8 | -16 | 8 |
| Obleser, Meyer & Friederici | 2011 | 10.1016/j.neuroimage.2011.03.035 | Less > more vocoded speech | 14 | 56 | -12 | 4 |
|  |  |  |  |  | -50 | 2 | 4 |
|  |  |  |  |  | -44 | -22 | 12 |
| Hocking, McMahon & de Zubicaray | 2011 | 10.1002/hbm.21040 | Environmental sounds > perceptual baseline | 13 | -6 | -51 | 12 |
|  |  |  |  |  | -27 | -33 | -18 |
|  |  |  |  |  | -36 | 24 | -15 |
|  |  |  |  |  | -15 | 33 | 45 |
|  |  |  |  |  | -45 | -72 | 39 |
|  |  |  |  |  | 39 | -69 | -39 |
|  |  |  |  |  | 6 | -51 | -42 |
|  |  |  |  |  | -60 | -15 | -15 |
| Rapp & Lipka | 2011 | 10.1162/jocn.2010.21507 | Words > checkerboards | 10 | -39 | -47 | -9 |
|  |  |  |  |  | -36 | 20 | 5 |
|  |  |  |  |  | -39 | 1 | 31 |
|  |  |  |  |  | -39 | 26 | 22 |
|  |  |  |  |  | -36 | 20 | 5 |
|  |  |  |  |  | 24 | -18 | 36 |
| Rapp & Lipka | 2011 | 10.1162/jocn.2010.21507 | Words > letter strings | 10 | 55 | -44 | -5 |
|  |  |  |  |  | -42 | -51 | 5 |
|  |  |  |  |  | -24 | 5 | 5 |
|  |  |  |  |  | 58 | -6 | 37 |
| Rodd, Longe, Randall & Tyler | 2010 | 10.1016/j.neuropsychologia.2009.12.035 | Speech > SCN | 14 | -60 | -10 | 0 |
|  |  |  |  |  | -48 | 10 | -22 |
|  |  |  |  |  | -52 | 8 | -20 |
|  |  |  |  |  | -50 | 12 | -16 |
|  |  |  |  |  | -64 | -22 | 0 |
|  |  |  |  |  | -54 | -6 | -14 |
|  |  |  |  |  | -52 | -30 | 2 |
|  |  |  |  |  | 60 | -4 | -10 |
|  |  |  |  |  | 52 | 6 | -22 |
|  |  |  |  |  | 66 | -18 | -2 |
|  |  |  |  |  | 52 | -18 | 2 |
|  |  |  |  |  | -24 | -12 | -20 |
|  |  |  |  |  | 30 | -14 | -20 |
| Khader, Jost, Mertens, Bien & Roesler | 2010 | 10.1016/j.brainres.2009.12.082 | Noun > rhyme generation | 16 | -66 | 51 | -13 |
|  |  |  |  |  | -42 | 62 | 3 |
|  |  |  |  |  | -53 | 73 | 5 |
|  |  |  |  |  | -7 | 63 | 4 |
|  |  |  |  |  | 47 | 66 | -3 |
|  |  |  |  |  | 56 | 60 | -15 |
|  |  |  |  |  | 58 | 53 | -12 |
|  |  |  |  |  | 64 | 31 | 24 |
| Khader, Jost, Mertens, Bien & Roesler | 2010 | 10.1016/j.brainres.2009.12.082 | Verb > rhyme generation | 16 | -42 | -67 | 16 |
|  |  |  |  |  | -54 | -68 | 18 |
| Khader, Jost, Mertens, Bien & Roesler | 2010 | 10.1016/j.brainres.2009.12.082 | Noun generation > letter detection | 16 | 5 | 58 | 12 |
|  |  |  |  |  | 50 | 28 | -3 |
|  |  |  |  |  | 2 | 25 | 23 |
|  |  |  |  |  | 8 | -58 | 13 |
| Khader, Jost, Mertens, Bien & Roesler | 2010 | 10.1016/j.brainres.2009.12.082 | Verb generation > letter detection | 16 | 5 | 59 | 10 |
|  |  |  |  |  | 50 | 25 | -1 |
|  |  |  |  |  | 2 | 23 | 22 |
|  |  |  |  |  | 68 | -51 | 7 |
|  |  |  |  |  | 55 | -77 | 23 |
| Kuchinke, van der Meer & Krueger | 2009 | 10.1016/j.bandc.2008.07.014 | Semantic > syntactic decision | 15 | 19 | -84 | 10 |
|  |  |  |  |  | -31 | 2 | 47 |
|  |  |  |  |  | -46 | 16 | 25 |
|  |  |  |  |  | 36 | 11 | 45 |
|  |  |  |  |  | 25 | -30 | -18 |
|  |  |  |  |  | -53 | -39 | -11 |
|  |  |  |  |  | 30 | -64 | 51 |
| Cao, Peng, Liu, Jin, Fan, Deng, & Booth | 2009 | 10.1002/hbm.20546 | Semantic > perceptual decision | 13 | 27 | -90 | 6 |
|  |  |  |  |  | -3 | -96 | 15 |
|  |  |  |  |  | -24 | -90 | -9 |
|  |  |  |  |  | -45 | 27 | 21 |
|  |  |  |  |  | 6 | -63 | -30 |
|  |  |  |  |  | -60 | -45 | 0 |
|  |  |  |  |  | -3 | 33 | 39 |
|  |  |  |  |  | -36 | -3 | -33 |
|  |  |  |  |  | 30 | -69 | -36 |
|  |  |  |  |  | 33 | -24 | -24 |
|  |  |  |  |  | 39 | 30 | -3 |
| Leff, Schofield, Stephan, Crinion, Friston & Price | 2008 | 10.1523/JNEUROSCI.2903-08.2008 | Speech > reversed speech | 26 | -52 | -48 | 8 |
|  |  |  |  |  | -54 | 10 | -16 |
|  |  |  |  |  | -48 | 28 | -6 |
| Alain, He & Grady | 2008 | 10.1162/jocn.2008.20014 | Category > location judgement | 16 | -40 | 32 | 7 |
|  |  |  |  |  | -1 | 10 | 53 |
|  |  |  |  |  | -42 | -27 | 10 |
|  |  |  |  |  | -56 | -4 | -3 |
|  |  |  |  |  | -56 | -34 | 6 |
|  |  |  |  |  | 46 | -24 | 9 |
|  |  |  |  |  | 49 | -1 | -9 |
|  |  |  |  |  | 6 | -68 | -29 |
| Sabri, Binder, Desai, Medler, Leitl & Liebenthal | 2008 | 10.1016/j.neuroimage.2007.09.052 | Speech > rotated speech | 28 | 21 | -83 | 34 |
|  |  |  |  |  | 52 | -74 | 7 |
|  |  |  |  |  | 19 | -50 | 63 |
|  |  |  |  |  | -22 | 40 | -17 |
| Sabri, Binder, Desai, Medler, Leitl & Liebenthal | 2008 | 10.1016/j.neuroimage.2007.09.052 | Words > pseudowords | 28 | -37 | 21 | 34 |
| Vingerhoets | 2008 | 10.1016/j.neuroimage.2007.12.058 | Familiar > unfamiliar tools | 14 | -54 | -57 | 37 |
|  |  |  |  |  | -63 | -49 | 26 |
|  |  |  |  |  | -7 | -69 | 41 |
|  |  |  |  |  | -57 | -38 | 1 |
| Hakonen, May, Jaaskelainen, Jokinen, Sams & Tiitinen | 2017 | 10.1002/brb3.789 | More > less intelligible based on cue | 20 | 2 | 10 | 48 |
|  |  |  |  |  | -4 | -90 | -12 |
|  |  |  |  |  | -36 | 0 | 40 |
|  |  |  |  |  | 38 | 60 | 18 |
|  |  |  |  |  | 38 | 24 | -14 |
|  |  |  |  |  | -32 | 56 | 4 |
| Hakonen, May, Jaaskelainen, Jokinen, Sams & Tiitinen | 2017 | 10.1002/brb3.789 | More > less intelligible | 20 | -52 | -12 | -20 |
|  |  |  |  |  | 58 | 2 | -24 |
|  |  |  |  |  | -6 | -90 | -8 |
|  |  |  |  |  | 40 | -36 | -8 |
|  |  |  |  |  | -50 | -52 | 16 |
|  |  |  |  |  | -52 | 16 | 26 |
|  |  |  |  |  | 28 | 0 | -30 |
| Hakonen, May, Jaaskelainen, Jokinen, Sams & Tiitinen | 2017 | 10.1002/brb3.789 | More > less intelligible | 20 | -60 | -6 | -18 |
|  |  |  |  |  | 50 | 12 | -26 |
|  |  |  |  |  | -10 | 54 | 42 |
|  |  |  |  |  | -38 | -54 | 18 |
|  |  |  |  |  | 50 | -34 | -2 |
|  |  |  |  |  | -2 | 58 | -18 |
| Garn, Allen & Larsen | 2009 | 10.1016/j.cortex.2008.02.004 | Pictures > scrambled pictures | 26 | 31 | -79 | -15 |
|  |  |  |  |  | -24 | -92 | -6 |
|  |  |  |  |  | 12 | -88 | -6 |
|  |  |  |  |  | -8 | 5 | 54 |
|  |  |  |  |  | -40 | -85 | -15 |
|  |  |  |  |  | -30 | -69 | 55 |
|  |  |  |  |  | -7 | -26 | 10 |
|  |  |  |  |  | 1 | 2 | 33 |
| Boulenger, Hauk & Pulvermuller | 2009 | 10.1093/cercor/bhn217 | Sentences > hashmarks | 18 | -50 | 20 | 16 |
|  |  |  |  |  | -40 | -2 | 48 |
|  |  |  |  |  | -48 | 30 | 2 |
|  |  |  |  |  | -4 | 10 | 54 |
|  |  |  |  |  | -8 | 52 | 32 |
|  |  |  |  |  | -54 | -42 | 2 |
|  |  |  |  |  | -54 | -6 | -16 |
|  |  |  |  |  | -56 | -58 | 12 |
|  |  |  |  |  | -50 | 10 | -20 |
|  |  |  |  |  | -42 | -42 | -18 |
|  |  |  |  |  | -12 | -78 | -34 |
| Boulenger, Hauk & Pulvermuller | 2009 | 10.1093/cercor/bhn217 | Sentences > hashmarks | 18 | -52 | 26 | 2 |
|  |  |  |  |  | -48 | 22 | 24 |
|  |  |  |  |  | -38 | 6 | 48 |
|  |  |  |  |  | -46 | 38 | -6 |
|  |  |  |  |  | -54 | -42 | 2 |
|  |  |  |  |  | -62 | -36 | -2 |
|  |  |  |  |  | -52 | -28 | -4 |
|  |  |  |  |  | -46 | -58 | 18 |
|  |  |  |  |  | -46 | -36 | -12 |
|  |  |  |  |  | -14 | -82 | -30 |
| Raposo, Moss, Stamatakis & Tyler | 2009 | 10.1016/j.neuropsychologia.2008.09.017 | Action sentences > SCN | 22 | -60 | -12 | -4 |
|  |  |  |  |  | 62 | -10 | -4 |
| Lin, Wang, Zhao, Liu, Li & Bi | 2015 | 10.1162/jocn_a_00852 | Words > pseudowords | 20 | 3 | -9 | 39 |
|  |  |  |  |  | 60 | -30 | 36 |
|  |  |  |  |  | -45 | -54 | 27 |
| Ludersdorfer, Wimmer, Richlan, Schurz, Hutzler & Kronbichler | 2016 | 10.1016/j.neuroimage.2015.09.039 | Words > tones | 29 | -48 | 32 | 7 |
|  |  |  |  |  | -42 | 5 | 25 |
|  |  |  |  |  | -48 | 20 | 22 |
|  |  |  |  |  | -60 | -28 | -5 |
|  |  |  |  |  | -45 | -64 | -11 |
|  |  |  |  |  | -27 | -67 | 40 |
|  |  |  |  |  | -12 | -85 | 34 |
|  |  |  |  |  | 60 | -22 | -5 |
| Ludersdorfer, Philipp; Wimmer, Heinz; Richlan, Fabio; Schurz, Matthias; Hutzler, Florian; Kronbichler, Martin | 2016 | 10.1016/j.neuroimage.2015.09.039 | Words > tones | 29 | -60 | -34 | 5 |
|  |  |  |  |  | -48 | 20 | 19 |
|  |  |  |  |  | -33 | -67 | 40 |
|  |  |  |  |  | -12 | -82 | 28 |
|  |  |  |  |  | 36 | 26 | -11 |
|  |  |  |  |  | 60 | -22 | 5 |
|  |  |  |  |  | 9 | -76 | 7 |
| Segal & Petrides | 2012 | 10.1111/j.1460-9568.2011.07937.x | Writing > copying | 90 | -44 | 16 | 2 |
|  |  |  |  |  | -46 | 36 | 8 |
|  |  |  |  |  | -12 | -18 | -6 |
|  |  |  |  |  | -12 | -34 | -6 |
|  |  |  |  |  | -28 | -67 | -12 |
|  |  |  |  |  | -50 | -48 | -14 |
|  |  |  |  |  | -24 | -98 | 12 |
|  |  |  |  |  | -14 | -98 | -2 |
|  |  |  |  |  | 34 | 18 | 0 |
|  |  |  |  |  | 50 | 36 | 10 |
|  |  |  |  |  | 16 | -32 | 2 |
|  |  |  |  |  | 28 | -72 | -14 |
|  |  |  |  |  | 28 | -96 | 18 |
|  |  |  |  |  | 12 | -92 | -8 |
| Segal & Petrides | 2012 | 10.1111/j.1460-9568.2011.07937.x | Words > pseudowords | 90 | -50 | 22 | -8 |
|  |  |  |  |  | -58 | -66 | 26 |
|  |  |  |  |  | -66 | -52 | 24 |
| Carota, Kriegeskorte, Nili & Pulvermuller Friedemann | 2017 | 10.1093/cercor/bhw379 | Words > hashmarks | 23 | 6 | 20 | 44 |
|  |  |  |  |  | -6 | 18 | 48 |
|  |  |  |  |  | 28 | 28 | -8 |
|  |  |  |  |  | -40 | -8 | -28 |
|  |  |  |  |  | -40 | -32 | -20 |
|  |  |  |  |  | -34 | 7 | -30 |
|  |  |  |  |  | 22 | -70 | -44 |
|  |  |  |  |  | 40 | -58 | -32 |
|  |  |  |  |  | 38 | -48 | -32 |
|  |  |  |  |  | -52 | 8 | 18 |
|  |  |  |  |  | -52 | 0 | 28 |
|  |  |  |  |  | -32 | -10 | 50 |
| Schuil, Smits & Zwaan | 2013 | 10.3389/fnhum.2013.00100 | Senteces > pseudowords | 20 | -52 | -38 | 2 |
|  |  |  |  |  | 58 | -8 | 12 |
|  |  |  |  |  | 14 | -88 | 26 |
|  |  |  |  |  | -52 | 26 | 4 |
|  |  |  |  |  | -8 | -56 | 8 |
|  |  |  |  |  | -48 | -2 | 52 |
|  |  |  |  |  | -6 | 6 | 62 |
| Schuil, Smits & Zwaan | 2013 | 10.3389/fnhum.2013.00100 | Verbs > pseudowords | 20 | -48 | 26 | 10 |
|  |  |  |  |  | -48 | 28 | -10 |
|  |  |  |  |  | -4 | 18 | 48 |
|  |  |  |  |  | -12 | 44 | 42 |
|  |  |  |  |  | -10 | 30 | 50 |
|  |  |  |  |  | -50 | 16 | 28 |
| Schuil, Smits & Zwaan | 2013 | 10.3389/fnhum.2013.00100 | Literal sentences > pseudowords | 20 | -48 | 24 | 10 |
|  |  |  |  |  | -4 | 18 | 48 |
|  |  |  |  |  | -10 | 30 | 54 |
|  |  |  |  |  | -10 | 44 | 42 |
|  |  |  |  |  | -10 | 20 | 56 |
|  |  |  |  |  | -48 | 4 | -24 |
| Schuil, Smits & Zwaan | 2013 | 10.3389/fnhum.2013.00100 | Nonliteral sentences > pseudowords | 20 | -48 | 26 | 10 |
|  |  |  |  |  | -50 | 28 | -10 |
|  |  |  |  |  | -12 | 44 | 44 |
|  |  |  |  |  | -6 | 20 | 46 |
| Garbin, Collina & Tabossi | 2012 | 10.1371/journal.pone.0045091 | Object noun > pseudoword | 12 | -60 | -32 | 32 |
| Garbin, Collina & Tabossi | 2012 | 10.1371/journal.pone.0045091 | Event noun > pseudoword | 12 | -28 | 32 | 22 |
|  |  |  |  |  | -44 | 14 | 16 |
|  |  |  |  |  | -62 | -20 | 12 |
|  |  |  |  |  | 60 | -28 | 16 |
|  |  |  |  |  | 56 | -58 | 14 |
| Garbin, Collina & Tabossi | 2012 | 10.1371/journal.pone.0045091 | Verb > pseudoword | 12 | -42 | 18 | 6 |
|  |  |  |  |  | -48 | -32 | 22 |
|  |  |  |  |  | -56 | -6 | 16 |
|  |  |  |  |  | -58 | -4 | 4 |
|  |  |  |  |  | 66 | -52 | 10 |
|  |  |  |  |  | 40 | -54 | 6 |
| Vignali | 2019 | 10.1016/j.neuroimage.2018.08.061 | Words > pseudowords | 21 | -45 | -73 | 25 |
|  |  |  |  |  | -6 | -52 | 10 |
|  |  |  |  |  | 9 | -55 | 10 |
|  |  |  |  |  | -24 | 23 | 49 |
|  |  |  |  |  | -9 | 53 | -11 |
|  |  |  |  |  | -30 | -37 | -20 |
|  |  |  |  |  | -57 | -10 | -20 |
|  |  |  |  |  | -24 | -19 | -23 |
|  |  |  |  |  | -33 | 32 | -17 |
|  |  |  |  |  | 30 | -31 | -20 |
|  |  |  |  |  | 24 | -37 | -20 |
|  |  |  |  |  | 27 | -19 | -23 |
|  |  |  |  |  | 27 | 32 | 40 |
|  |  |  |  |  | -27 | -10 | 7 |
|  |  |  |  |  | -15 | -1 | 19 |
|  |  |  |  |  | 18 | 5 | 19 |
|  |  |  |  |  | 18 | 14 | 13 |
|  |  |  |  |  | 21 | -10 | 19 |
|  |  |  |  |  | 42 | -28 | 58 |
|  |  |  |  |  | 48 | -13 | 52 |
|  |  |  |  |  | 30 | -25 | 58 |
|  |  |  |  |  | -9 | 62 | 22 |
| Wu, Mai, Tang, Ge, Luo & Liu | 2013 | 10.1038/srep02049 | Arm words > checkerboard | 19 | -5 | 14 | 64 |
|  |  |  |  |  | -59 | 9 | 44 |
|  |  |  |  |  | 62 | 10 | 47 |
|  |  |  |  |  | -49 | 4 | 61 |
|  |  |  |  |  | 59 | 4 | 53 |
|  |  |  |  |  | -49 | -12 | 67 |
|  |  |  |  |  | -25 | -60 | 61 |
|  |  |  |  |  | -44 | -19 | -34 |
| Wu, Mai, Tang, Ge, Luo & Liu | 2013 | 10.1038/srep02049 | Leg words > checkerboard | 19 | -1 | 9 | 75 |
|  |  |  |  |  | 5 | 68 | 31 |
|  |  |  |  |  | -59 | 9 | 44 |
|  |  |  |  |  | 62 | 10 | 47 |
|  |  |  |  |  | -49 | 4 | 61 |
|  |  |  |  |  | 50 | -12 | 71 |
|  |  |  |  |  | -28 | -52 | 70 |
|  |  |  |  |  | 29 | -62 | 71 |
|  |  |  |  |  | -52 | -43 | 27 |
|  |  |  |  |  | -59 | -8 | 51 |
|  |  |  |  |  | 59 | -9 | 60 |
|  |  |  |  |  | -46 | -67 | -13 |
|  |  |  |  |  | -50 | -44 | -21 |
|  |  |  |  |  | -50 | 14 | -21 |
|  |  |  |  |  | -63 | 0 | -15 |
|  |  |  |  |  | 49 | -66 | -10 |
| Wu, Mai, Tang, Ge, Luo & Liu | 2013 | 10.1038/srep02049 | Mouth words > checkerboard | 19 | -8 | 6 | 81 |
|  |  |  |  |  | -59 | 9 | 44 |
|  |  |  |  |  | 62 | 10 | 47 |
|  |  |  |  |  | -63 | 27 | 15 |
|  |  |  |  |  | 70 | 14 | 31 |
|  |  |  |  |  | -49 | 4 | 61 |
|  |  |  |  |  | 59 | 7 | 53 |
|  |  |  |  |  | -28 | -54 | 76 |
|  |  |  |  |  | -59 | -37 | 27 |
|  |  |  |  |  | 59 | -16 | 61 |
|  |  |  |  |  | -22 | -60 | 61 |
|  |  |  |  |  | -59 | 8 | -4 |
|  |  |  |  |  | -50 | -44 | -21 |
|  |  |  |  |  | 38 | -45 | -28 |
| Groussard, Viader, Hubert, Landeau, Abbas, Desgranges, Eustache & Platel | 2010 | 10.1016/j.neuroimage.2009.10.039 | Semantics > reference | 11 | 24 | -70 | -30 |
|  |  |  |  |  | -62 | -38 | 0 |
| Kim, Koizumi, Ikuta, Fukumitsu, Kimura, Iwata, Watanabe, Yokoyama, Sato, Horie & Kawashima | 2009 | 10.1016/j.jneuroling.2008.07.005 | Sentences > word lists | 36 | -60 | 31 | -2 |
|  |  |  |  |  | -47 | 28 | -20 |
|  |  |  |  |  | -53 | 24 | 12 |
|  |  |  |  |  | -61 | -41 | 5 |
|  |  |  |  |  | -72 | -48 | 4 |
|  |  |  |  |  | -58 | -1 | -26 |
|  |  |  |  |  | -64 | -2 | -15 |
|  |  |  |  |  | -64 | -28 | 2 |
|  |  |  |  |  | -23 | -15 | -18 |
| Bozic & Marslen-Wilson | 2013 | 10.2298/PSI1304439B | Speech > musical rain | 13 | -66 | -26 | -8 |
|  |  |  |  |  | -30 | -36 | -18 |
|  |  |  |  |  | -42 | -42 | -18 |
|  |  |  |  |  | -46 | -68 | 28 |
|  |  |  |  |  | 62 | -8 | -12 |
|  |  |  |  |  | -38 | 30 | -16 |
|  |  |  |  |  | -36 | 36 | -10 |
|  |  |  |  |  | 54 | -64 | 28 |
|  |  |  |  |  | 4 | 50 | 12 |
|  |  |  |  |  | -8 | 36 | -10 |
|  |  |  |  |  | 8 | 30 | -10 |
|  |  |  |  |  | 34 | -38 | -16 |
|  |  |  |  |  | 32 | -26 | -14 |
|  |  |  |  |  | 32 | -36 | -8 |
| Bagga, Singh, Modi, Kumar, Bhattacharya, Garg & Khushu | 2013 | 10.1007/s12038-013-9387-7 | Semantic > matching judgement | 18 | -39 | 8 | 25 |
|  |  |  |  |  | 21 | -79 | -32 |
|  |  |  |  |  | 48 | 29 | 7 |
|  |  |  |  |  | -51 | -31 | -8 |
|  |  |  |  |  | -12 | 2 | 64 |
| Chan, Tang, Tang, Lee, Lo & Kwong | 2009 | 10.1016/j.neuroimage.2009.06.078 | Semantics > pseudocharacters | 22 | -52 | 24 | -2 |
|  |  |  |  |  | -6 | 23 | 49 |
|  |  |  |  |  | -40 | -45 | 49 |
|  |  |  |  |  | 42 | 25 | -7 |
|  |  |  |  |  | -55 | -34 | -4 |
|  |  |  |  |  | -17 | 10 | 9 |
| Chan, Tang, Tang, Lee, Lo & Kwong | 2009 | 10.1016/j.neuroimage.2009.06.078 | Known > unknown language | 22 | -53 | 26 | 0 |
|  |  |  |  |  | -1 | 33 | 43 |
|  |  |  |  |  | -33 | -58 | 49 |
|  |  |  |  |  | -34 | 25 | -5 |
|  |  |  |  |  | 41 | 26 | -5 |
|  |  |  |  |  | -57 | -40 | -10 |
|  |  |  |  |  | -49 | 58 | -17 |
|  |  |  |  |  | -18 | 9 | 8 |
| Malins, Gumkowski, Buis, Molfese, Rueckl, Frost, Pugh, Morris & Mencl | 2016 | 10.1016/j.neuropsychologia.2016.08.027 | Unrelated words > false font | 18 | -49 | 1 | 20 |
|  |  |  |  |  | 0 | 5 | 59 |
|  |  |  |  |  | -34 | 27 | 3 |
|  |  |  |  |  | 39 | 27 | 2 |
|  |  |  |  |  | -46 | 18 | 18 |
|  |  |  |  |  | -50 | 14 | 2 |
|  |  |  |  |  | -50 | -47 | -17 |
|  |  |  |  |  | -66 | -32 | 6 |
|  |  |  |  |  | 55 | -29 | 1 |
|  |  |  |  |  | -59 | -11 | -4 |
|  |  |  |  |  | -53 | 33 | 0 |
|  |  |  |  |  | -62 | -18 | 25 |
|  |  |  |  |  | 62 | 0 | 38 |
|  |  |  |  |  | 23 | -63 | -20 |
|  |  |  |  |  | 26 | -69 | -50 |
|  |  |  |  |  | 16 | -78 | -42 |
|  |  |  |  |  | -17 | 8 | 8 |
|  |  |  |  |  | 13 | 11 | 1 |
| Malins, Gumkowski, Buis, Molfese, Rueckl, Frost, Pugh, Morris & Mencl | 2016 | 10.1016/j.neuropsychologia.2016.08.027 | Unrelated > pseudowords | 18 | -17 | 8 | 8 |
|  |  |  |  |  | 13 | 11 | 1 |
| Harrington, Farias & Davis | 2009 | 10.1016/j.cortex.2007.10.015 | Real > meaningless objects | 8 | -53 | -31 | 41 |
|  |  |  |  |  | -18 | 12 | 54 |
|  |  |  |  |  | -45 | 3 | 23 |
|  |  |  |  |  | 53 | 14 | 17 |
|  |  |  |  |  | -34 | -3 | 8 |
|  |  |  |  |  | -42 | 29 | 26 |
|  |  |  |  |  | -51 | -59 | -5 |
|  |  |  |  |  | -30 | -39 | -17 |
| Taylor, Arsalidou, Bayless, Morris, Evans & Barbeau | 2009 | 10.1002/hbm.20646 | Own > unfamiliar face | 10 | 4 | 41 | 14 |
|  |  |  |  |  | 0 | 42 | 14 |
|  |  |  |  |  | 9 | 24 | 27 |
|  |  |  |  |  | 0 | 26 | 26 |
|  |  |  |  |  | 21 | 33 | 26 |
|  |  |  |  |  | -15 | 35 | 26 |
|  |  |  |  |  | -40 | 41 | 11 |
| Taylor, Arsalidou, Bayless, Morris, Evans & Barbeau | 2009 | 10.1002/hbm.20646 | Partner's > unfamiliar face | 10 | 7 | 39 | 18 |
|  |  |  |  |  | -2 | 37 | 19 |
|  |  |  |  |  | 7 | 37 | 24 |
|  |  |  |  |  | -1 | 37 | 23 |
|  |  |  |  |  | 8 | 65 | -2 |
|  |  |  |  |  | -6 | 54 | -1 |
|  |  |  |  |  | -23 | 44 | 34 |
|  |  |  |  |  | -23 | 18 | -22 |
|  |  |  |  |  | -52 | -12 | -18 |
|  |  |  |  |  | -29 | 17 | -10 |
|  |  |  |  |  | -26 | -2 | -15 |
|  |  |  |  |  | -20 | -26 | 4 |
|  |  |  |  |  | -14 | -10 | -19 |
|  |  |  |  |  | -5 | -54 | 47 |
| Taylor, Arsalidou, Bayless, Morris, Evans & Barbeau | 2009 | 10.1002/hbm.20646 | Parent's > unfamiliar face | 10 | 7 | 32 | 26 |
|  |  |  |  |  | 0 | 27 | 26 |
|  |  |  |  |  | 8 | 31 | 55 |
| Chanraud-Guillermo, Andoh, Martelli, Artiges, Pallier, Aubin, Martinot & Reynaud | 2009 | 10.1111/j.1530-0277.2009.00918.x | Known > unknown language | 12 | 10 | 45 | 45 |
|  |  |  |  |  | -59 | -32 | 58 |
|  |  |  |  |  | 61 | 23 | -12 |
|  |  |  |  |  | 66 | 23 | 37 |
|  |  |  |  |  | 75 | -15 | -4 |
|  |  |  |  |  | -68 | 6 | 5 |
|  |  |  |  |  | -33 | -13 | 73 |
|  |  |  |  |  | 58 | -53 | 58 |
|  |  |  |  |  | -72 | -46 | 1 |
| Husain, Patkin, Kim, Braun & Horwitz | 2012 | 10.1016/j.brainres.2012.08.029 | Meaningful iconic > meaningless gestures | 16 | -96 | -4 | 26 |
|  |  |  |  |  | -100 | -2 | 16 |
|  |  |  |  |  | -88 | 8 | 12 |
|  |  |  |  |  | 34 | -70 | -22 |
|  |  |  |  |  | 48 | -72 | -12 |
|  |  |  |  |  | 46 | -56 | -22 |
|  |  |  |  |  | -42 | -76 | -18 |
|  |  |  |  |  | -50 | -72 | -16 |
|  |  |  |  |  | -38 | -62 | -22 |
|  |  |  |  |  | -24 | -50 | 42 |
|  |  |  |  |  | -34 | -34 | 34 |
|  |  |  |  |  | -16 | -80 | 50 |
|  |  |  |  |  | 52 | 0 | 22 |
|  |  |  |  |  | 50 | 2 | 34 |
|  |  |  |  |  | 38 | 4 | 16 |
|  |  |  |  |  | -42 | 2 | 20 |
|  |  |  |  |  | -34 | -4 | 30 |
|  |  |  |  |  | -32 | -14 | 34 |
|  |  |  |  |  | 50 | -48 | 52 |
|  |  |  |  |  | 44 | -42 | 36 |
|  |  |  |  |  | 56 | -36 | 46 |
|  |  |  |  |  | -58 | -64 | 14 |
| Yang, Li, Fang, Shu, Liu & Chen | 2016 | 10.1016/j.neuropsychologia.2016.04.029 | Opaque idioms > hashmarks | 20 | -38 | -17 | 54 |
|  |  |  |  |  | -51 | 13 | 34 |
|  |  |  |  |  | -42 | 32 | 22 |
|  |  |  |  |  | -36 | 17 | 0 |
|  |  |  |  |  | -26 | 3 | -39 |
|  |  |  |  |  | -42 | -55 | -13 |
|  |  |  |  |  | -13 | -82 | 9 |
|  |  |  |  |  | -32 | -41 | 2 |
|  |  |  |  |  | -10 | -11 | 9 |
|  |  |  |  |  | -20 | -39 | -42 |
|  |  |  |  |  | 55 | 10 | 26 |
|  |  |  |  |  | 36 | 24 | 8 |
|  |  |  |  |  | 38 | 1 | -40 |
|  |  |  |  |  | 33 | -66 | 44 |
|  |  |  |  |  | 46 | -80 | -5 |
|  |  |  |  |  | 20 | -69 | 11 |
|  |  |  |  |  | 13 | -11 | 9 |
|  |  |  |  |  | 7 | 18 | 43 |
|  |  |  |  |  | 3 | -60 | -30 |
| Yang, Li, Fang, Shu, Liu & Chen | 2016 | 10.1016/j.neuropsychologia.2016.04.029 | Transparent idioms > hashmarks | 20 | -49 | 39 | 1 |
|  |  |  |  |  | -29 | 30 | 5 |
|  |  |  |  |  | -42 | -55 | -13 |
|  |  |  |  |  | -42 | -5 | -28 |
|  |  |  |  |  | -13 | -73 | 8 |
|  |  |  |  |  | 49 | 16 | 26 |
|  |  |  |  |  | 32 | 33 | -6 |
|  |  |  |  |  | 39 | -84 | 18 |
|  |  |  |  |  | 30 | -65 | 54 |
|  |  |  |  |  | 20 | -79 | 8 |
|  |  |  |  |  | 4 | 12 | 50 |
|  |  |  |  |  | 3 | -53 | -27 |
| Yang, Li, Fang, Shu, Liu & Chen | 2016 | 10.1016/j.neuropsychologia.2016.04.029 | Literal phrases > hashmarks | 20 | -35 | 9 | 24 |
|  |  |  |  |  | -45 | 32 | 22 |
|  |  |  |  |  | -45 | 43 | 1 |
|  |  |  |  |  | -29 | 26 | -1 |
|  |  |  |  |  | -48 | -43 | 46 |
|  |  |  |  |  | -36 | -2 | -15 |
|  |  |  |  |  | 46 | 26 | 25 |
|  |  |  |  |  | 36 | 33 | -3 |
|  |  |  |  |  | 39 | -33 | -23 |
|  |  |  |  |  | 30 | -63 | 44 |
|  |  |  |  |  | 17 | -75 | 15 |
|  |  |  |  |  | 17 | 24 | 35 |
|  |  |  |  |  | 4 | 15 | 50 |
| Lindenberg, Uhlig, Scherfeld, Schlaug & Seitz | 2012 | 10.1002/hbm.21258 | Meaningful iconic > meaningless gestures | 20 | -44 | 22 | -23 |
|  |  |  |  |  | 61 | 34 | -6 |
|  |  |  |  |  | -8 | 64 | 14 |
|  |  |  |  |  | -5 | 35 | 29 |
|  |  |  |  |  | -54 | -12 | -18 |
|  |  |  |  |  | -4 | 39 | -26 |
|  |  |  |  |  | -56 | -58 | 33 |
|  |  |  |  |  | 11 | -89 | -22 |
| Mashal, Vishne, Laor & Titone | 2013 | 10.1016/j.bandl.2012.11.012 | Novel metaphor > unrelated words | 14 | 42 | 26 | 1 |
|  |  |  |  |  | -54 | 22 | 22 |
|  |  |  |  |  | 40 | 27 | 2 |
|  |  |  |  |  | -28 | -90 | -6 |
|  |  |  |  |  | 2 | 8 | 62 |
|  |  |  |  |  | -3 | 31 | 43 |
|  |  |  |  |  | -26 | 2 | 1 |
| Mashal, Vishne, Laor & Titone | 2013 | 10.1016/j.bandl.2012.11.012 | Real metaphor > unrelated words | 14 | -40 | 26 | 7 |
|  |  |  |  |  | -45 | 23 | 14 |
|  |  |  |  |  | -40 | 25 | 8 |
|  |  |  |  |  | 0 | 7 | 59 |
|  |  |  |  |  | -16 | 8 | 5 |
|  |  |  |  |  | -11 | 10 | 9 |
| Marques, Canessa & Cappa | 2009 | 10.1016/j.cortex.2008.07.004 | Sentences > crosses | 21 | -38 | -80 | -8 |
|  |  |  |  |  | -22 | -92 | -2 |
|  |  |  |  |  | -34 | -46 | -24 |
|  |  |  |  |  | 40 | -64 | -12 |
|  |  |  |  |  | 36 | -62 | -18 |
|  |  |  |  |  | -56 | -56 | 6 |
|  |  |  |  |  | -52 | -34 | -2 |
|  |  |  |  |  | -28 | -70 | 26 |
|  |  |  |  |  | -24 | -66 | 46 |
|  |  |  |  |  | -28 | -58 | 46 |
|  |  |  |  |  | -44 | -28 | 42 |
|  |  |  |  |  | -6 | 8 | 50 |
|  |  |  |  |  | 8 | -4 | 66 |
|  |  |  |  |  | -48 | 8 | 28 |
|  |  |  |  |  | -54 | 12 | 16 |
|  |  |  |  |  | -54 | 22 | 24 |
|  |  |  |  |  | -50 | 38 | 8 |
|  |  |  |  |  | -44 | 18 | -12 |
|  |  |  |  |  | -30 | 18 | 6 |
|  |  |  |  |  | -50 | 4 | 40 |
|  |  |  |  |  | -28 | -4 | 48 |
|  |  |  |  |  | -22 | 18 | -2 |
|  |  |  |  |  | 42 | 42 | 26 |
|  |  |  |  |  | 40 | 24 | -16 |
|  |  |  |  |  | 32 | 28 | -4 |
|  |  |  |  |  | 52 | 36 | 14 |
|  |  |  |  |  | 52 | 38 | 8 |
|  |  |  |  |  | 50 | 34 | 22 |
|  |  |  |  |  | 36 | 44 | 32 |
|  |  |  |  |  | 22 | 20 | -4 |
|  |  |  |  |  | -22 | -34 | -2 |
|  |  |  |  |  | -8 | 16 | -4 |
|  |  |  |  |  | 6 | 12 | -4 |
|  |  |  |  |  | 36 | 50 | 20 |
| Christensen, Antonucci, Lockwood, Kittleson & Plante | 2008 | 10.1097/WNR.0b013e3283060a9d | Diotic listening > reversed speech | 14 | 6 | 20 | 43 |
|  |  |  |  |  | 35 | 24 | 3 |
|  |  |  |  |  | 35 | 43 | 20 |
|  |  |  |  |  | 38 | 57 | 0 |
|  |  |  |  |  | 32 | 60 | -21 |
|  |  |  |  |  | 32 | -43 | 43 |
|  |  |  |  |  | 19 | 5 | 16 |
|  |  |  |  |  | 16 | -7 | 10 |
|  |  |  |  |  | -5 | 13 | 51 |
|  |  |  |  |  | -38 | 21 | 3 |
|  |  |  |  |  | -44 | 17 | 24 |
|  |  |  |  |  | -34 | -37 | -22 |
|  |  |  |  |  | -46 | 30 | 10 |
|  |  |  |  |  | -60 | -39 | 12 |
|  |  |  |  |  | -11 | 3 | 17 |
|  |  |  |  |  | -24 | 16 | 12 |
|  |  |  |  |  | -13 | -4 | 9 |
| Christensen, Antonucci, Lockwood, Kittleson & Plante | 2008 | 10.1097/WNR.0b013e3283060a9d | Dichotic listening > reversed speech | 14 | 5 | 14 | 51 |
|  |  |  |  |  | 35 | 24 | 0 |
|  |  |  |  |  | 35 | 53 | 16 |
|  |  |  |  |  | 47 | -34 | 43 |
|  |  |  |  |  | 18 | 1 | 16 |
|  |  |  |  |  | 16 | -8 | 13 |
|  |  |  |  |  | -5 | 13 | 51 |
|  |  |  |  |  | -36 | 24 | 4 |
|  |  |  |  |  | -45 | 4 | 32 |
|  |  |  |  |  | -37 | 52 | 24 |
|  |  |  |  |  | -38 | -39 | -19 |
| Saur, Kreher, Schnell, Kummerer, Kellmeyer, Vry, Umarova, Musso, Glauche, Abel, Huber, Rijntjes, Hennig & Weiller | 2008 | 10.1073/pnas.0805234105 | Word > pseudoword sentences | 33 | -48 | -60 | 18 |
|  |  |  |  |  | -51 | 0 | -18 |
|  |  |  |  |  | -30 | -33 | -18 |
|  |  |  |  |  | -36 | -60 | 39 |
|  |  |  |  |  | -48 | 27 | 12 |
|  |  |  |  |  | -45 | 27 | -12 |
|  |  |  |  |  | -9 | 63 | 27 |
|  |  |  |  |  | -3 | 18 | 54 |
|  |  |  |  |  | -39 | 18 | 30 |
|  |  |  |  |  | 51 | -3 | -18 |
|  |  |  |  |  | 42 | -54 | 18 |
|  |  |  |  |  | 33 | -30 | -21 |
|  |  |  |  |  | 48 | 24 | -9 |
|  |  |  |  |  | 42 | -21 | 54 |
|  |  |  |  |  | 51 | 30 | 30 |
| Emmorey, Weisberg, McCullough & Petrich | 2013 | 10.1016/j.bandl.2013.05.001 | Words > false fonts | 14 | -4 | 36 | 50 |
|  |  |  |  |  | 7 | 20 | 48 |
|  |  |  |  |  | 37 | 20 | -4 |
|  |  |  |  |  | -46 | 11 | 34 |
|  |  |  |  |  | -53 | 37 | -15 |
|  |  |  |  |  | -39 | 38 | -4 |
|  |  |  |  |  | -50 | 36 | 12 |
|  |  |  |  |  | -51 | 28 | 1 |
|  |  |  |  |  | -54 | 22 | 26 |
|  |  |  |  |  | -32 | 35 | 4 |
|  |  |  |  |  | -31 | 23 | -3 |
|  |  |  |  |  | -42 | -13 | -23 |
|  |  |  |  |  | -46 | -42 | -9 |
|  |  |  |  |  | -54 | -41 | 4 |
| Emmorey, Weisberg, McCullough & Petrich | 2013 | 10.1016/j.bandl.2013.05.001 | Words > false fonts | 14 | 52 | 34 | 26 |
|  |  |  |  |  | 0 | 17 | 49 |
|  |  |  |  |  | 7 | 17 | 49 |
|  |  |  |  |  | -49 | 11 | 27 |
|  |  |  |  |  | -54 | 46 | 0 |
|  |  |  |  |  | -27 | 31 | 4 |
|  |  |  |  |  | -39 | 20 | 3 |
|  |  |  |  |  | -54 | -53 | -15 |
|  |  |  |  |  | -57 | -32 | 50 |
|  |  |  |  |  | 50 | -58 | 59 |
|  |  |  |  |  | -34 | -48 | 43 |
| Allendorfer, Lindsell, Siegel, Banks, Vannest, Holland & Szaflarski | 2012 | 10.1016/j.cortex.2011.05.014 | Covert verb generation > finger tapping | 40 | 36 | 26 | -4 |
|  |  |  |  |  | -45 | 24 | 20 |
|  |  |  |  |  | 56 | -23 | 0 |
|  |  |  |  |  | -62 | -10 | 1 |
|  |  |  |  |  | 0 | 19 | 60 |
|  |  |  |  |  | -24 | -66 | 60 |
| Allendorfer, Lindsell, Siegel, Banks, Vannest, Holland & Szaflarski | 2012 | 10.1016/j.cortex.2011.05.014 | Overt verb generation > repetition | 40 | 35 | -65 | -31 |
|  |  |  |  |  | 35 | 26 | -13 |
|  |  |  |  |  | -50 | 24 | 20 |
|  |  |  |  |  | -54 | -39 | -1 |
|  |  |  |  |  | 15 | 18 | 1 |
|  |  |  |  |  | -9 | 6 | 3 |
|  |  |  |  |  | 8 | -59 | 8 |
|  |  |  |  |  | -1 | 26 | 41 |
|  |  |  |  |  | -29 | 67 | 33 |
| Matchin | 2019 | 10.1016/j.neuropsychologia.2019.01.019 | Verb phrase > list | 20 | -37 | 17 | -12 |
|  |  |  |  |  | -39 | 27 | -17 |
|  |  |  |  |  | -23 | -16 | -12 |
|  |  |  |  |  | -56 | -4 | -10 |
|  |  |  |  |  | 0 | 41 | -15 |
|  |  |  |  |  | -51 | -39 | 8 |
|  |  |  |  |  | -33 | -17 | 22 |
|  |  |  |  |  | -9 | 59 | 18 |
|  |  |  |  |  | -44 | -61 | 26 |
|  |  |  |  |  | -33 | -34 | -15 |
|  |  |  |  |  | 53 | 22 | -30 |
|  |  |  |  |  | 24 | -81 | -32 |
|  |  |  |  |  | 45 | -16 | 1 |
|  |  |  |  |  | -13 | 14 | 22 |
| Matchin | 2019 | 10.1016/j.neuropsychologia.2019.01.019 | Noun phrase > list | 20 | -1 | 47 | 2 |
|  |  |  |  |  | -37 | 23 | -20 |
|  |  |  |  |  | -45 | -58 | 25 |
|  |  |  |  |  | -57 | -2 | -12 |
|  |  |  |  |  | -6 | -56 | 25 |
|  |  |  |  |  | -23 | -13 | -13 |
|  |  |  |  |  | 50 | 20 | -26 |
|  |  |  |  |  | -32 | -15 | 18 |
|  |  |  |  |  | -49 | -34 | 1 |
|  |  |  |  |  | 58 | 2 | -17 |
|  |  |  |  |  | 32 | 13 | -21 |
| Axmacher, Bialleck, Weber, Helmstaedter, Elger & Fell | 2009 | 10.1002/hbm.20645 | Words > shapes | 32 | -36 | -87 | -18 |
|  |  |  |  |  | 27 | -96 | -9 |
|  |  |  |  |  | -42 | -48 | -30 |
| Zaccarella & Friederici | 2015 | 10.3389/fpsyg.2015.01818 | Words > pseudowords | 22 | -33 | 23 | -2 |
|  |  |  |  |  | 36 | 23 | -2 |
|  |  |  |  |  | -48 | 11 | 7 |
| Jackson, Hoffman, Pobric, Lambon Ralph | 2015 | 10.1093/cercor/bhv003 | Words > letter strings | 24 | 60 | 3 | -3 |
|  |  |  |  |  | 45 | -3 | -15 |
|  |  |  |  |  | 27 | -48 | 9 |
|  |  |  |  |  | -21 | -21 | -21 |
|  |  |  |  |  | -45 | -15 | -27 |
|  |  |  |  |  | -45 | -18 | 63 |
|  |  |  |  |  | -33 | -21 | 72 |
|  |  |  |  |  | 21 | -84 | -36 |
|  |  |  |  |  | -9 | 48 | 39 |
|  |  |  |  |  | -9 | 54 | 30 |
|  |  |  |  |  | -9 | 57 | 18 |
|  |  |  |  |  | 12 | -3 | 45 |
|  |  |  |  |  | 3 | 3 | 39 |
|  |  |  |  |  | 0 | -12 | 48 |
|  |  |  |  |  | -3 | 54 | -12 |
|  |  |  |  |  | -15 | 42 | -3 |
|  |  |  |  |  | 9 | -84 | 27 |
|  |  |  |  |  | -3 | -84 | 27 |
| van Leeuwen, Lamers, Petersson, Gussenhoven, Rietveld, Poser & Hagoort | 2014 | 10.1016/j.neuropsychologia.2014.03.017 | Speech > reversed speech | 16 | -22 | -90 | -16 |
|  |  |  |  |  | 10 | 10 | 48 |
|  |  |  |  |  | -4 | 4 | 48 |
|  |  |  |  |  | 0 | 10 | 40 |
|  |  |  |  |  | -24 | -88 | -2 |
|  |  |  |  |  | -26 | -80 | 2 |
|  |  |  |  |  | -38 | -78 | -2 |
|  |  |  |  |  | -36 | -68 | -26 |
|  |  |  |  |  | -36 | -76 | -30 |
|  |  |  |  |  | -40 | -32 | 44 |
|  |  |  |  |  | 8 | 40 | 14 |
|  |  |  |  |  | 42 | 40 | 24 |
|  |  |  |  |  | -50 | -70 | -6 |
| Zou, Packard, Xia, Liu & Shu | 2016 | 10.3339/fnhum.2015.00714 | Speech > tone | 17 | -45 | 36 | 3 |
|  |  |  |  |  | 33 | 27 | 6 |
|  |  |  |  |  | -3 | 18 | 48 |
|  |  |  |  |  | 12 | -75 | -30 |
|  |  |  |  |  | -45 | 48 | 0 |
|  |  |  |  |  | 60 | -3 | -3 |
|  |  |  |  |  | -60 | -9 | 3 |
|  |  |  |  |  | 12 | -66 | 9 |
|  |  |  |  |  | -6 | -63 | 9 |
|  |  |  |  |  | -5 | -81 | -24 |
|  |  |  |  |  | 30 | -90 | -3 |
|  |  |  |  |  | -30 | -66 | 42 |
|  |  |  |  |  | -36 | -32 | -19 |
|  |  |  |  |  | 0 | -51 | -15 |
|  |  |  |  |  | 33 | 39 | -9 |
|  |  |  |  |  | -66 | -30 | 3 |
|  |  |  |  |  | -6 | 66 | 27 |
|  |  |  |  |  | 0 | 39 | -15 |
|  |  |  |  |  | -33 | -87 | -9 |
|  |  |  |  |  | 45 | 27 | 30 |
| Zou, Packard, Xia, Liu & Shu | 2016 | 10.3339/fnhum.2015.00714 | Speech > tone | 17 | -45 | 36 | 3 |
|  |  |  |  |  | -60 | -12 | 6 |
|  |  |  |  |  | 60 | -3 | -3 |
|  |  |  |  |  | -27 | -9 | -18 |
|  |  |  |  |  | -45 | -51 | -12 |
|  |  |  |  |  | 60 | 0 | 12 |
|  |  |  |  |  | -6 | 21 | 48 |
|  |  |  |  |  | 9 | -81 | -24 |
|  |  |  |  |  | 12 | -84 | -36 |
|  |  |  |  |  | 68 | -15 | 9 |
|  |  |  |  |  | 0 | -48 | -36 |
|  |  |  |  |  | -66 | -30 | 3 |
|  |  |  |  |  | 36 | 39 | -6 |
|  |  |  |  |  | 39 | -27 | 69 |
|  |  |  |  |  | -36 | -32 | -19 |
| Zou, Packard, Xia, Liu & Shu | 2016 | 10.3339/fnhum.2015.00714 | Speech > tone | 17 | -60 | -9 | 6 |
|  |  |  |  |  | -39 | -24 | 12 |
|  |  |  |  |  | -42 | -15 | -27 |
|  |  |  |  |  | -36 | -30 | -18 |
|  |  |  |  |  | 60 | -3 | -3 |
|  |  |  |  |  | 48 | -15 | 12 |
|  |  |  |  |  | -45 | 24 | 24 |
|  |  |  |  |  | -51 | 36 | 9 |
|  |  |  |  |  | -48 | 48 | 0 |
|  |  |  |  |  | 21 | -60 | -24 |
|  |  |  |  |  | -48 | -54 | -12 |
|  |  |  |  |  | -51 | -45 | -3 |
|  |  |  |  |  | -3 | 15 | 57 |
|  |  |  |  |  | -21 | -3 | 9 |
|  |  |  |  |  | -12 | -18 | 9 |
|  |  |  |  |  | 3 | -60 | -36 |
|  |  |  |  |  | 39 | 42 | -9 |
|  |  |  |  |  | -27 | -51 | -30 |
|  |  |  |  |  | -30 | -39 | -24 |
|  |  |  |  |  | 30 | -63 | -48 |
|  |  |  |  |  | -60 | -12 | 6 |
|  |  |  |  |  | -37 | -31 | -18 |
| Zou, Packard, Xia, Liu & Shu | 2016 | 10.3339/fnhum.2015.00714 | Identical speech > tone | 17 | -63 | -9 | 0 |
|  |  |  |  |  | 60 | 0 | 0 |
|  |  |  |  |  | -51 | -15 | 3 |
|  |  |  |  |  | 45 | -18 | 6 |
|  |  |  |  |  | 18 | -30 | 78 |
|  |  |  |  |  | -39 | 30 | -12 |
|  |  |  |  |  | -60 | -5 | 21 |
|  |  |  |  |  | 24 | -24 | 75 |
|  |  |  |  |  | 69 | -27 | 0 |
| Zhang, Liu & Zhang | 2014 | 10.1016/j.cortex.2013.01.015 | Nonliving words > asterisks | 18 | -46 | 12 | 31 |
|  |  |  |  |  | -3 | 15 | 68 |
|  |  |  |  |  | 62 | 39 | 18 |
|  |  |  |  |  | 13 | -101 | 5 |
|  |  |  |  |  | -18 | -44 | -39 |
|  |  |  |  |  | 16 | -2 | -7 |
|  |  |  |  |  | 23 | 9 | 9 |
| Zhang, Liu & Zhang | 2014 | 10.1016/j.cortex.2013.01.015 | Living words > asterisks | 18 | -4 | 18 | 67 |
|  |  |  |  |  | -43 | 9 | 31 |
|  |  |  |  |  | -43 | 19 | -12 |
|  |  |  |  |  | 46 | 0 | 65 |
|  |  |  |  |  | -27 | -104 | 4 |
|  |  |  |  |  | 20 | -94 | 4 |
|  |  |  |  |  | 35 | -8 | -46 |
|  |  |  |  |  | 42 | -14 | -31 |
|  |  |  |  |  | -24 | -27 | -5 |
|  |  |  |  |  | 17 | 0 | 23 |
|  |  |  |  |  | 16 | -2 | -9 |
| Emmorey, Xu, Gannon, Goldin-Meadow & Braun | 2010 | 10.1016/j.neuroimage.2009.08.001 | Meaningful pantomimes > unknown sign language | 14 | -24 | -12 | 66 |
|  |  |  |  |  | -48 | -48 | -24 |
|  |  |  |  |  | 48 | -81 | 15 |
|  |  |  |  |  | -60 | -27 | 48 |
|  |  |  |  |  | -25 | -51 | 71 |
|  |  |  |  |  | 33 | -51 | 69 |
|  |  |  |  |  | -63 | -27 | 42 |
|  |  |  |  |  | 60 | -26 | 43 |
| Smith, Myers, Sethi, Pantazatos, Yanagihara & Hirsch | 2012 | 10.1080/02643294.2012.706218 | Semantics > baseline | 14 | -32 | -36 | -18 |
|  |  |  |  |  | -10 | -56 | 6 |
|  |  |  |  |  | -18 | -52 | 8 |
|  |  |  |  |  | -38 | -70 | 26 |
|  |  |  |  |  | -38 | 32 | -16 |
|  |  |  |  |  | -44 | 14 | 44 |
|  |  |  |  |  | -50 | 28 | 4 |
|  |  |  |  |  | -54 | -64 | 14 |
| Hartung, Hagoort & Willems | 2017 | 10.7554/eLife.25964 | First-person speech > unintelligible reversed speech | 52 | 18 | -30 | 58 |
|  |  |  |  |  | 26 | -32 | 56 |
|  |  |  |  |  | 8 | -30 | 56 |
|  |  |  |  |  | -20 | -28 | 54 |
|  |  |  |  |  | -22 | -26 | 62 |
|  |  |  |  |  | -18 | -36 | 54 |
|  |  |  |  |  | -44 | -74 | -4 |
| Hartung, Hagoort & Willems | 2017 | 10.7554/eLife.25964 | Third-person speech > unintelligible reversed speech | 52 | 18 | -86 | -16 |
|  |  |  |  |  | -40 | -6 | -22 |
|  |  |  |  |  | -34 | -12 | -22 |
| Ryan, Lin, Ketcham & Nadel | 2010 | 10.1002/hipo.20607 | Semantic > letter judgement | 15 | -22 | -12 | -12 |
|  |  |  |  |  | -26 | -30 | -20 |
|  |  |  |  |  | -22 | -8 | -12 |
|  |  |  |  |  | 26 | -34 | 4 |
|  |  |  |  |  | 26 | -38 | -12 |
| Ryan, Lin, Ketcham & Nadel | 2010 | 10.1002/hipo.20607 | Semantic > letter judgement | 15 | -20 | -12 | -14 |
|  |  |  |  |  | -22 | -34 | -2 |
|  |  |  |  |  | -24 | -42 | -10 |
|  |  |  |  |  | -22 | -8 | -12 |
|  |  |  |  |  | 26 | -34 | 4 |
| Ryan, Lin, Ketcham & Nadel | 2010 | 10.1002/hipo.20607 | Semantic > letter judgement | 15 | -14 | -40 | 4 |
|  |  |  |  |  | -20 | -8 | -12 |
|  |  |  |  |  | -22 | -34 | 0 |
|  |  |  |  |  | -20 | -6 | -12 |
|  |  |  |  |  | 38 | -20 | -18 |
|  |  |  |  |  | 22 | -34 | 4 |
| Ryan, Lin, Ketcham & Nadel | 2010 | 10.1002/hipo.20607 | Semantic > letter judgement | 15 | -22 | -12 | -14 |
|  |  |  |  |  | -18 | -6 | -24 |
|  |  |  |  |  | -22 | -8 | -12 |
|  |  |  |  |  | 20 | -12 | -20 |
|  |  |  |  |  | 20 | -26 | -10 |
|  |  |  |  |  | 20 | -26 | -12 |
|  |  |  |  |  | 22 | -12 | -22 |
| Ryan, Lin, Ketcham & Nadel | 2010 | 10.1002/hipo.20607 | Semantic > episodic judgement | 15 | -28 | -8 | -24 |
|  |  |  |  |  | -28 | -10 | -28 |
|  |  |  |  |  | 28 | -6 | -20 |
| Roxbury, McMahon & Copland | 2014 | 10.1186/1744-9081-10-34 | Concrete word > pseudoword | 17 | 7 | -50 | 47 |
|  |  |  |  |  | 32 | -32 | -7 |
|  |  |  |  |  | -32 | 25 | 54 |
|  |  |  |  |  | -54 | -14 | -25 |
| Roxbury, McMahon & Copland | 2014 | 10.1186/1744-9081-10-34 | Abstract word > pseudoword | 17 | -54 | -58 | 18 |
|  |  |  |  |  | 50 | -40 | 32 |
|  |  |  |  |  | 14 | -50 | 32 |
| Hayashi, Okamoto, Yoshimura, Yoshino, Toki, Yamashita, Matsuda & Yamawaki | 2014 | 10.1016/j.neures.2013.10.007 | Concrete word > astersisks | 16 | -6 | 4 | 60 |
|  |  |  |  |  | 14 | 4 | 68 |
|  |  |  |  |  | -40 | 36 | 10 |
|  |  |  |  |  | 56 | 22 | 22 |
|  |  |  |  |  | -28 | 28 | -6 |
|  |  |  |  |  | 36 | 32 | 0 |
|  |  |  |  |  | -20 | 6 | 4 |
|  |  |  |  |  | 30 | 6 | 0 |
|  |  |  |  |  | -40 | -6 | 50 |
|  |  |  |  |  | -36 | -8 | 66 |
|  |  |  |  |  | -22 | -62 | 44 |
|  |  |  |  |  | -50 | -56 | -14 |
|  |  |  |  |  | -24 | -88 | -8 |
| Hayashi, Okamoto, Yoshimura, Yoshino, Toki, Yamashita, Matsuda & Yamawaki | 2014 | 10.1016/j.neures.2013.10.007 | Abstract word > asterisks | 16 | -4 | 16 | 52 |
|  |  |  |  |  | -46 | 20 | 24 |
|  |  |  |  |  | 56 | 28 | 20 |
|  |  |  |  |  | -44 | -2 | 48 |
|  |  |  |  |  | -46 | 42 | -12 |
|  |  |  |  |  | -30 | -60 | 48 |
|  |  |  |  |  | -48 | -54 | -14 |
|  |  |  |  |  | -26 | -86 | -2 |
|  |  |  |  |  | 38 | -82 | -4 |
| Sachs, Weis, Krings, Huber & Kircher | 2008 | 10.1016/j.neuropsychologia.2007.08.015 | Thematic judgement > letters | 14 | -52 | 20 | 28 |
|  |  |  |  |  | -60 | -36 | 0 |
|  |  |  |  |  | -8 | -88 | 20 |
|  |  |  |  |  | -32 | -60 | 36 |
|  |  |  |  |  | -48 | -4 | 48 |
|  |  |  |  |  | -12 | -48 | 0 |
|  |  |  |  |  | 16 | -100 | 4 |
|  |  |  |  |  | 0 | -12 | 68 |
|  |  |  |  |  | 12 | -52 | 100 |
| Sachs, Weis, Krings, Huber & Kircher | 2008 | 10.1016/j.neuropsychologia.2007.08.015 | Taxonomic judgement > letters | 14 | -52 | 20 | 24 |
|  |  |  |  |  | -32 | -60 | 36 |
|  |  |  |  |  | 0 | 4 | 56 |
|  |  |  |  |  | 20 | -100 | 4 |
|  |  |  |  |  | -12 | -48 | 0 |
|  |  |  |  |  | -16 | -96 | -12 |
|  |  |  |  |  | -64 | -44 | 0 |
|  |  |  |  |  | 52 | 20 | 28 |
|  |  |  |  |  | 16 | -52 | 0 |
|  |  |  |  |  | -24 | -28 | -4 |
|  |  |  |  |  | 40 | 16 | 4 |
| Erb, Henry, Eisner & Obleser | 2013 | 10.1523/JNEUROSCI.4596-12.2013 | Speech > vocoded speech | 30 | 6 | 59 | 28 |
|  |  |  |  |  | 54 | -7 | 16 |
|  |  |  |  |  | 27 | -10 | -2 |
|  |  |  |  |  | 63 | -19 | -11 |
|  |  |  |  |  | 51 | -58 | 22 |
|  |  |  |  |  | -54 | -1 | 22 |
|  |  |  |  |  | -27 | -10 | -2 |
|  |  |  |  |  | 63 | -19 | -11 |
|  |  |  |  |  | 51 | -58 | 22 |
|  |  |  |  |  | -54 | -1 | 22 |
|  |  |  |  |  | -27 | -10 | -2 |
|  |  |  |  |  | -54 | -10 | 7 |
|  |  |  |  |  | -51 | -67 | 34 |
|  |  |  |  |  | -60 | -13 | -20 |
|  |  |  |  |  | -6 | -37 | 43 |
| Erb, Henry, Eisner & Obleser | 2013 | 10.1523/JNEUROSCI.4596-12.2013 | Intelligibility parameter | 30 | -3 | 50 | -14 |
|  |  |  |  |  | -51 | 26 | 13 |
|  |  |  |  |  | -57 | -7 | -20 |
|  |  |  |  |  | -54 | -10 | 34 |
|  |  |  |  |  | -57 | -13 | 4 |
|  |  |  |  |  | -30 | -16 | -2 |
|  |  |  |  |  | -51 | -61 | 19 |
|  |  |  |  |  | 57 | -4 | 31 |
|  |  |  |  |  | 57 | -13 | 4 |
|  |  |  |  |  | 30 | -7 | -8 |
|  |  |  |  |  | -12 | -19 | 4 |
|  |  |  |  |  | 12 | -19 | 4 |
|  |  |  |  |  | -3 | -52 | 16 |
|  |  |  |  |  | 15 | -61 | -23 |
|  |  |  |  |  | -12 | -64 | -23 |
| Obleser, Eisner & Kotz | 2008 | 10.1523/JNEUROSCI.1290-08.2008 | Intelligibility parameter | 16 | 62 | -12 | 8 |
|  |  |  |  |  | -50 | -36 | -6 |
|  |  |  |  |  | -48 | 4 | -22 |
|  |  |  |  |  | -62 | -16 | 0 |
| Just, Keller & Cynkar | 2008 | 10.1016/j.brainres.2007.12.075 | Distracted listening > listening to speech | 29 | -56 | -12 | -6 |
|  |  |  |  |  | 50 | -20 | 4 |
|  |  |  |  |  | -44 | 20 | 26 |
|  |  |  |  |  | 2 | 24 | 62 |
| Kinno, Kawamura, Shioda & Sakai | 2008 | 10.1002/hbm.20441 | Canonical sentence > picture & letter strings | 14 | -36 | 12 | 51 |
|  |  |  |  |  | 45 | 15 | 36 |
|  |  |  |  |  | -54 | 18 | 12 |
|  |  |  |  |  | -45 | 39 | 9 |
|  |  |  |  |  | -3 | 24 | 48 |
|  |  |  |  |  | -54 | -51 | 6 |
|  |  |  |  |  | -48 | -60 | 51 |
|  |  |  |  |  | 42 | -63 | 36 |
| Kinno, Kawamura, Shioda & Sakai | 2008 | 10.1002/hbm.20441 | Active sentence > picture & letter strings | 14 | -45 | 6 | 48 |
|  |  |  |  |  | 48 | 21 | 33 |
|  |  |  |  |  | -51 | 18 | 27 |
|  |  |  |  |  | -51 | 27 | 6 |
|  |  |  |  |  | -3 | 18 | 51 |
|  |  |  |  |  | -6 | 3 | 3 |
|  |  |  |  |  | 9 | 3 | 0 |
|  |  |  |  |  | -54 | -54 | 6 |
|  |  |  |  |  | 51 | -54 | 6 |
|  |  |  |  |  | -24 | -78 | 33 |
|  |  |  |  |  | 45 | -78 | 6 |
| Kinno, Kawamura, Shioda & Sakai | 2008 | 10.1002/hbm.20441 | Passive sentence > picture & letter strings | 14 | -39 | -3 | 45 |
|  |  |  |  |  | 45 | 18 | 36 |
|  |  |  |  |  | -51 | 21 | 24 |
|  |  |  |  |  | -54 | 27 | 3 |
|  |  |  |  |  | 3 | 18 | 51 |
|  |  |  |  |  | 9 | 3 | 3 |
|  |  |  |  |  | -54 | -54 | 6 |
|  |  |  |  |  | 51 | -60 | 9 |
|  |  |  |  |  | -39 | -57 | 54 |
|  |  |  |  |  | -30 | -81 | 30 |
|  |  |  |  |  | 42 | -78 | 6 |
| Jensen, Hargreaves, Bass, Pexman, Goodyear & Federico | 2011 | 10.1016/j.eplepsyres.2010.12.003 | Words > pseudowords | 12 | -62 | -54 | -12 |
|  |  |  |  |  | -26 | 34 | 48 |
|  |  |  |  |  | 20 | 42 | 16 |
|  |  |  |  |  | 52 | -66 | 20 |
|  |  |  |  |  | 66 | -44 | 2 |
|  |  |  |  |  | 4 | -34 | 8 |
|  |  |  |  |  | 44 | -4 | 56 |
|  |  |  |  |  | 6 | -62 | 12 |
|  |  |  |  |  | 30 | 26 | 52 |
|  |  |  |  |  | 18 | 0 | 66 |
|  |  |  |  |  | 30 | 48 | 18 |
|  |  |  |  |  | 60 | -42 | 26 |
| Barros-Loscertales, Gonzalez, Pulvermueller, Ventura-Campos, Carlos Bustamante, Costumero, Antonia Parcet & Avila | 2012 | 10.1093/cercor/bhr324 | Words > hashmarks | 59 | -3 | 3 | 60 |
|  |  |  |  |  | -9 | 21 | 48 |
|  |  |  |  |  | -45 | -51 | -18 |
|  |  |  |  |  | -42 | -42 | -18 |
|  |  |  |  |  | -60 | -33 | 3 |
|  |  |  |  |  | -51 | -6 | 45 |
|  |  |  |  |  | -36 | 30 | 9 |
|  |  |  |  |  | -36 | 33 | -9 |
|  |  |  |  |  | 54 | 0 | 42 |
|  |  |  |  |  | -24 | -9 | 6 |
|  |  |  |  |  | -21 | -6 | -3 |
| Europa | 2019 | 10.3389/fnhum.2019.00027 | Speech > reversed speech | 21 | -50 | 28 | 18 |
|  |  |  |  |  | -4 | 8 | 60 |
|  |  |  |  |  | -54 | 8 | -18 |
|  |  |  |  |  | -14 | -2 | 14 |
|  |  |  |  |  | -30 | -4 | 56 |
|  |  |  |  |  | -28 | -46 | 44 |
|  |  |  |  |  | 10 | -68 | -24 |
|  |  |  |  |  | -26 | -70 | 32 |
|  |  |  |  |  | 48 | -76 | 8 |
|  |  |  |  |  | -26 | -100 | -8 |
| Raettig & Kotz | 2008 | 10.1016/j.neuroimage.2007.09.030 | Words > pseudowords | 16 | -4 | -67 | 36 |
|  |  |  |  |  | -49 | -70 | 33 |
|  |  |  |  |  | -8 | 42 | -9 |
|  |  |  |  |  | 67 | -14 | -21 |
| Chow, Kaup, Raabe & Greenlee | 2008 | 10.1016/j.neuroimage.2007.11.044 | Words > pseudowords | 15 | -58 | 20 | 14 |
|  |  |  |  |  | -58 | -34 | 2 |
|  |  |  |  |  | -52 | -56 | 16 |
|  |  |  |  |  | -58 | -14 | -12 |
|  |  |  |  |  | -54 | 2 | -18 |
|  |  |  |  |  | -50 | 24 | 0 |
|  |  |  |  |  | 52 | -36 | 0 |
|  |  |  |  |  | 52 | 10 | -16 |
| Chow, Kaup, Raabe & Greenlee | 2008 | 10.1016/j.neuroimage.2007.11.044 | Words > pseudowords | 15 | -58 | 20 | 14 |
|  |  |  |  |  | -60 | -60 | 20 |
|  |  |  |  |  | -58 | -4 | -14 |
|  |  |  |  |  | -58 | -34 | -2 |
| Chow, Kaup, Raabe & Greenlee | 2008 | 10.1016/j.neuroimage.2007.11.044 | Words > pseudowords | 15 | -54 | 16 | 18 |
|  |  |  |  |  | -58 | -34 | -2 |
|  |  |  |  |  | -50 | 26 | 0 |
|  |  |  |  |  | -50 | -56 | 16 |
|  |  |  |  |  | -58 | -14 | -12 |
|  |  |  |  |  | 52 | -36 | -2 |
|  |  |  |  |  | 46 | 28 | -10 |
| Elman, Klostermann, Marian, Verstaen & Shimamura | 2012 | 10.3758/s13415-012-0096-8 | Semantic > episodic judgement | 19 | 50 | 24 | -22 |
| Bedny, Pascual-Leone, Dodell-Feder, Fedorenko & Saxe | 2011 | 10.1073/pnas.1014818108 | Sentences & lists > pseudoword sentences & lists | 22 | 64 | -42 | 2 |
|  |  |  |  |  | 50 | -14 | -18 |
|  |  |  |  |  | -56 | -4 | -28 |
|  |  |  |  |  | -58 | -46 | 0 |
|  |  |  |  |  | -6 | 50 | 46 |
|  |  |  |  |  | -10 | 38 | 50 |
|  |  |  |  |  | -10 | -52 | 36 |
|  |  |  |  |  | -6 | -52 | 20 |
| Moseley, Carota, Hauk, Mohr & Pulvermueller | 2012 | 10.1093/cercor/bhr238 | Emotional words > hashmarks | 18 | -56 | 4 | 24 |
|  |  |  |  |  | -50 | -10 | 44 |
|  |  |  |  |  | -44 | 26 | -2 |
|  |  |  |  |  | 60 | 2 | 38 |
|  |  |  |  |  | 8 | 22 | 52 |
|  |  |  |  |  | -8 | 22 | 38 |
|  |  |  |  |  | 6 | 40 | 10 |
|  |  |  |  |  | -10 | 54 | 12 |
|  |  |  |  |  | -28 | 34 | -5 |
|  |  |  |  |  | -56 | -34 | 2 |
|  |  |  |  |  | -38 | -44 | -14 |
|  |  |  |  |  | -40 | -62 | -12 |
|  |  |  |  |  | -48 | -34 | 38 |
|  |  |  |  |  | -52 | -38 | 30 |
|  |  |  |  |  | 8 | -62 | 2 |
|  |  |  |  |  | -2 | -70 | 2 |
| Moseley, Carota, Hauk, Mohr & Pulvermueller | 2012 | 10.1093/cercor/bhr238 | Abstract emotional words > hashmarks | 18 | -56 | 4 | 24 |
|  |  |  |  |  | -48 | -12 | 40 |
|  |  |  |  |  | -56 | -8 | 44 |
|  |  |  |  |  | 56 | 0 | 40 |
|  |  |  |  |  | 60 | 2 | 24 |
|  |  |  |  |  | -10 | 56 | 12 |
|  |  |  |  |  | 8 | 52 | 8 |
|  |  |  |  |  | -60 | -34 | 2 |
|  |  |  |  |  | -40 | -40 | -14 |
|  |  |  |  |  | -44 | -74 | -8 |
|  |  |  |  |  | -48 | 16 | -26 |
|  |  |  |  |  | 50 | -32 | 24 |
|  |  |  |  |  | 36 | -64 | 2 |
|  |  |  |  |  | 16 | -50 | 64 |
|  |  |  |  |  | 28 | 24 | 6 |
|  |  |  |  |  | 34 | -16 | 22 |
| Weiss, Katzir & Bitan | 2015 | 10.1016/j.neuroimage.2015.07.029 | Pointed words > asterisks | 18 | -36 | -86 | -2 |
|  |  |  |  |  | -28 | 38 | -8 |
|  |  |  |  |  | -58 | -2 | 20 |
|  |  |  |  |  | 42 | -84 | -12 |
|  |  |  |  |  | -26 | -74 | 24 |
|  |  |  |  |  | 62 | -10 | 8 |
|  |  |  |  |  | 32 | 38 | -8 |
|  |  |  |  |  | -2 | -60 | 20 |
|  |  |  |  |  | 18 | -60 | -26 |
|  |  |  |  |  | -28 | -12 | -6 |
|  |  |  |  |  | -18 | 26 | 42 |
|  |  |  |  |  | -14 | -62 | -22 |
|  |  |  |  |  | -48 | 42 | 0 |
|  |  |  |  |  | 20 | -22 | 0 |
|  |  |  |  |  | 28 | -56 | 58 |
|  |  |  |  |  | 28 | -6 | -8 |
| Weiss, Katzir & Bitan | 2015 | 10.1016/j.neuroimage.2015.07.029 | Unpointed words > asterisks | 18 | -58 | -14 | 24 |
|  |  |  |  |  | -28 | 36 | -10 |
|  |  |  |  |  | -38 | -80 | -6 |
|  |  |  |  |  | 56 | -4 | 24 |
|  |  |  |  |  | -14 | -64 | -20 |
|  |  |  |  |  | -2 | -60 | 20 |
|  |  |  |  |  | -20 | -8 | -16 |
|  |  |  |  |  | -38 | -14 | 16 |
|  |  |  |  |  | -6 | 48 | 0 |
|  |  |  |  |  | 18 | -64 | -24 |
|  |  |  |  |  | -46 | 28 | 10 |
|  |  |  |  |  | -4 | 22 | 32 |
|  |  |  |  |  | -8 | -12 | 14 |
|  |  |  |  |  | -46 | -72 | 32 |
| Chang | 2019 | 10.1002/hbm.24502 | Words > maths | 26 | -58 | 20 | 4 |
|  |  |  |  |  | -58 | -28 | -8 |
|  |  |  |  |  | -46 | 26 | -18 |
|  |  |  |  |  | -60 | -58 | 24 |
|  |  |  |  |  | -46 | 14 | -30 |
|  |  |  |  |  | -60 | -22 | -20 |
|  |  |  |  |  | -36 | 24 | 42 |
|  |  |  |  |  | -12 | 42 | 42 |
|  |  |  |  |  | 6 | 54 | 26 |
|  |  |  |  |  | -8 | 10 | 64 |
|  |  |  |  |  | 52 | -24 | -10 |
|  |  |  |  |  | 48 | 30 | -16 |
|  |  |  |  |  | 60 | -62 | 8 |
|  |  |  |  |  | 24 | -74 | -38 |
|  |  |  |  |  | -24 | -78 | -38 |
| Vitello, Warren, Devlin & Rodd | 2014 | 10.3389/fnhum.2014.00530 | Sentences > SCN | 20 | -54 | -25 | -5 |
|  |  |  |  |  | -57 | -4 | -14 |
|  |  |  |  |  | -60 | -16 | -2 |
|  |  |  |  |  | -57 | -40 | 7 |
|  |  |  |  |  | 60 | -10 | -2 |
|  |  |  |  |  | 60 | -1 | -11 |
|  |  |  |  |  | -48 | -7 | 58 |
| Vagharchakian, Dehaene-Lambertz, Pallier & Dehaene | 2012 | 10.1523/JNEUROSCI.5685-11.2012 | Intelligibility parameter | 16 | -48 | -8 | -16 |
|  |  |  |  |  | -48 | -40 | 0 |
|  |  |  |  |  | -44 | 8 | -32 |
|  |  |  |  |  | -60 | -60 | 24 |
|  |  |  |  |  | -48 | -48 | 28 |
|  |  |  |  |  | -44 | 40 | -16 |
|  |  |  |  |  | -16 | 16 | 48 |
| Vagharchakian, Dehaene-Lambertz, Pallier & Dehaene | 2012 | 10.1523/JNEUROSCI.5685-11.2012 | Intelligible > unintelligble compression rate | 16 | -52 | -8 | -12 |
|  |  |  |  |  | -48 | -48 | 12 |
|  |  |  |  |  | -64 | -24 | -4 |
|  |  |  |  |  | 56 | 0 | -12 |
|  |  |  |  |  | -52 | 32 | 0 |
|  |  |  |  |  | -44 | 16 | 24 |
|  |  |  |  |  | -36 | 32 | -12 |
|  |  |  |  |  | 0 | 56 | -12 |
|  |  |  |  |  | -48 | 0 | 52 |
|  |  |  |  |  | -28 | -92 | 0 |
|  |  |  |  |  | -8 | 8 | 56 |
| Vagharchakian, Dehaene-Lambertz, Pallier & Dehaene | 2012 | 10.1523/JNEUROSCI.5685-11.2012 | Intelligible > unintelligble compression rate | 16 | -52 | -8 | -8 |
|  |  |  |  |  | -48 | -36 | 4 |
|  |  |  |  |  | -64 | -24 | 0 |
|  |  |  |  |  | 48 | -20 | -4 |
|  |  |  |  |  | -52 | 32 | -4 |
|  |  |  |  |  | -48 | 16 | 24 |
|  |  |  |  |  | -32 | 32 | -12 |
|  |  |  |  |  | 0 | 56 | -12 |
| Vagharchakian, Dehaene-Lambertz, Pallier & Dehaene | 2012 | 10.1523/JNEUROSCI.5685-11.2012 | Intelligible > unintelligble compression rate | 16 | -56 | -4 | -16 |
|  |  |  |  |  | -48 | -48 | 12 |
|  |  |  |  |  | 56 | 0 | -16 |
|  |  |  |  |  | -44 | 12 | 24 |
|  |  |  |  |  | -44 | 0 | 52 |
|  |  |  |  |  | -28 | -92 | 0 |
|  |  |  |  |  | 44 | -68 | -4 |
|  |  |  |  |  | 40 | -92 | 0 |
|  |  |  |  |  | -4 | 8 | 60 |
| Evans, McGettigan, Agnew, Rosen & Scott | 2016 | 10.1162/jocn_a_00913 | Intelligibility parameter | 20 | -52 | -10 | -16 |
|  |  |  |  |  | 52 | 2 | -20 |
| Schmitt | 2019 | 10.1080/23273798.2018.1533139 | Known > unknown language | 40 | -54 | -38 | 2 |
|  |  |  |  |  | -56 | -12 | -10 |
|  |  |  |  |  | -56 | -2 | -14 |
|  |  |  |  |  | 54 | 6 | -18 |
|  |  |  |  |  | 52 | -18 | -6 |
|  |  |  |  |  | 46 | 12 | -24 |
|  |  |  |  |  | -52 | 20 | 18 |
|  |  |  |  |  | -54 | 28 | 6 |
|  |  |  |  |  | -42 | 0 | 50 |
|  |  |  |  |  | 20 | -76 | -36 |
|  |  |  |  |  | 18 | -70 | -26 |
| Chouinard, Morrissey, Kohler & Goodale | 2008 | 10.1016/j.neuroimage.2008.02.011 | Meaningful > meaningless objects | 14 | -46 | 14 | 28 |
|  |  |  |  |  | -4 | 20 | 34 |
|  |  |  |  |  | -2 | 12 | 54 |
|  |  |  |  |  | -38 | -42 | -18 |
|  |  |  |  |  | -54 | -58 | -18 |
|  |  |  |  |  | -52 | -62 | -12 |
|  |  |  |  |  | -44 | -80 | -8 |
|  |  |  |  |  | 34 | -68 | -20 |
|  |  |  |  |  | 30 | -74 | -4 |
|  |  |  |  |  | 32 | -86 | 0 |
| Zvyagintsev, Clemens, Chechko, Mathiak, Sack & Mathiak | 2013 | 10.1111/ejn.12140 | Visual imagery > counting | 15 | 52 | 34.3 | -12 |
|  |  |  |  |  | 35 | -81 | -38 |
|  |  |  |  |  | -26 | -31 | -21 |
|  |  |  |  |  | 23 | -13 | 3 |
|  |  |  |  |  | -52 | 43 | -12 |
|  |  |  |  |  | -4 | 56 | -13 |
|  |  |  |  |  | -42 | -66 | 26 |
| Bick, Goelman & Frost | 2008 | 10.1162/jocn.2008.20.3.406 | Semantic > perceptual decision | 14 | -18 | -93 | -2 |
|  |  |  |  |  | -52 | -37 | 3 |
|  |  |  |  |  | -37 | 10 | 33 |
|  |  |  |  |  | -46 | 28 | 15 |
|  |  |  |  |  | -2 | 55 | 37 |
| Bick, Goelman & Frost | 2008 | 10.1162/jocn.2008.20.3.406 | Words > pseudowords | 14 | -17 | -91 | -2 |
|  |  |  |  |  | -40 | -65 | -11 |
|  |  |  |  |  | -33 | -60 | 40 |
|  |  |  |  |  | 10 | -73 | 25 |
|  |  |  |  |  | -53 | -34 | 1 |
|  |  |  |  |  | -44 | 6 | 52 |
|  |  |  |  |  | -44 | 24 | 21 |
|  |  |  |  |  | -15 | 10 | -2 |
| Bick, Goelman & Frost | 2008 | 10.1162/jocn.2008.20.3.406 | Words > pseudowords | 14 | -17 | -91 | -2 |
|  |  |  |  |  | -42 | -63 | -11 |
|  |  |  |  |  | 6 | -72 | 24 |
|  |  |  |  |  | -32 | -65 | 44 |
|  |  |  |  |  | -53 | -31 | -1 |
|  |  |  |  |  | 2 | -28 | 32 |
|  |  |  |  |  | -2 | -11 | 10 |
|  |  |  |  |  | -45 | 26 | 20 |
| Bick, Goelman & Frost | 2008 | 10.1162/jocn.2008.20.3.406 | Words > pseudowords | 14 | -18 | -92 | -3 |
|  |  |  |  |  | -33 | -64 | 43 |
|  |  |  |  |  | -45 | -55 | -12 |
|  |  |  |  |  | -24 | -31 | -1 |
|  |  |  |  |  | -48 | 1 | 50 |
|  |  |  |  |  | -44 | 23 | 21 |
| Matchin & Hickok | 2016 | 10.3389/fpsyg.2016.00241 | Sentences > word lists | 20 | 34 | 26 | 34 |
|  |  |  |  |  | -30 | -91 | -1 |
|  |  |  |  |  | -28 | 30 | 30 |
|  |  |  |  |  | 43 | -8 | 37 |
| Raposo, Frade & Alves | 2016 | 10.1016/j.neuropsychologia.2016.06.036 | Semantic > perceptual decision | 18 | -4 | -54 | 16 |
|  |  |  |  |  | -50 | 28 | -10 |
|  |  |  |  |  | -8 | 52 | 36 |
|  |  |  |  |  | -8 | -84 | -6 |
|  |  |  |  |  | 14 | -96 | 16 |
|  |  |  |  |  | -46 | -68 | 26 |
|  |  |  |  |  | 32 | 34 | -12 |
|  |  |  |  |  | 50 | 16 | -28 |
| Heim, Eickhoff & Amunts | 2008 | 10.1016/j.neuroimage.2008.01.009 | Semantic > phonological fluency | 28 | -60 | -10 | -23 |
|  |  |  |  |  | -28 | -38 | -19 |
| Peelle, Eason, Schmitter, Schwarzbauer & Davis | 2010 | 10.1016/j.neuroimage.2010.05.015 | Sentences > SCN | 6 | -62 | 18 | -20 |
|  |  |  |  |  | -68 | -22 | 10 |
|  |  |  |  |  | -52 | 16 | 18 |
|  |  |  |  |  | -58 | 32 | -6 |
|  |  |  |  |  | 44 | -84 | 28 |
|  |  |  |  |  | 6 | -96 | 16 |
|  |  |  |  |  | 66 | -14 | 6 |
|  |  |  |  |  | 68 | -12 | -6 |
|  |  |  |  |  | 56 | -26 | 6 |
|  |  |  |  |  | -6 | -50 | 10 |
|  |  |  |  |  | 6 | -50 | 10 |
|  |  |  |  |  | 60 | 26 | -18 |
|  |  |  |  |  | 54 | 18 | -18 |
|  |  |  |  |  | -30 | -84 | 34 |
|  |  |  |  |  | -60 | 0 | 50 |
|  |  |  |  |  | -50 | 0 | 50 |
|  |  |  |  |  | -16 | -16 | -12 |
|  |  |  |  |  | -14 | -22 | -4 |
|  |  |  |  |  | -22 | -6 | -4 |
| Gutchess, Hedden, Ketay, Aron & Gabrieli | 2010 | 10.1093/scan/nsp059 | Different > same words | 20 | -46 | -52 | -16 |
|  |  |  |  |  | -4 | -26 | -20 |
|  |  |  |  |  | -46 | 26 | 22 |
|  |  |  |  |  | -18 | -4 | 20 |
|  |  |  |  |  | -18 | 0 | 10 |
|  |  |  |  |  | -30 | -62 | 50 |
|  |  |  |  |  | -48 | 12 | 24 |
|  |  |  |  |  | 38 | 24 | -4 |
|  |  |  |  |  | 4 | -70 | -32 |
|  |  |  |  |  | -28 | -32 | 0 |
| Bhattasali | 2019 | 10.1080/23273798.2018.1518533 | Combinatorial parameter | 51 | 52 | 6 | -20 |
|  |  |  |  |  | 50 | -20 | -10 |
|  |  |  |  |  | 60 | -40 | -10 |
|  |  |  |  |  | -36 | 18 | -14 |
|  |  |  |  |  | -50 | 6 | -26 |
|  |  |  |  |  | -30 | 8 | -4 |
|  |  |  |  |  | 10 | 18 | 62 |
|  |  |  |  |  | 12 | 58 | 32 |
|  |  |  |  |  | -8 | 18 | 66 |
|  |  |  |  |  | -24 | -74 | -30 |
|  |  |  |  |  | 26 | -74 | -36 |
|  |  |  |  |  | 36 | -60 | -32 |
|  |  |  |  |  | -34 | -78 | 12 |
|  |  |  |  |  | -30 | -58 | -10 |
|  |  |  |  |  | -28 | -70 | -14 |
|  |  |  |  |  | 42 | 0 | 48 |
|  |  |  |  |  | -54 | -56 | 30 |
|  |  |  |  |  | -48 | -66 | 50 |
|  |  |  |  |  | -52 | -58 | 50 |
|  |  |  |  |  | 30 | -50 | -10 |
|  |  |  |  |  | -44 | 46 | -12 |
|  |  |  |  |  | -36 | 60 | -6 |
|  |  |  |  |  | -42 | 24 | 44 |
|  |  |  |  |  | -10 | -52 | 38 |
|  |  |  |  |  | 28 | -72 | 22 |
|  |  |  |  |  | -14 | 16 | 10 |
| Marques, Canessa, Siri, Catricala & Cappa | 2008 | 10.1016/j.brainres.2007.11.070 | Semantic features > baseline task | 21 | -28 | -90 | 0 |
|  |  |  |  |  | 32 | -88 | 2 |
|  |  |  |  |  | -56 | -48 | -2 |
|  |  |  |  |  | 0 | 8 | 60 |
|  |  |  |  |  | -4 | -2 | 60 |
|  |  |  |  |  | -52 | 12 | 16 |
|  |  |  |  |  | -36 | 28 | 0 |
|  |  |  |  |  | -42 | 32 | -2 |
| Herve, Razafimandimby, Vigneau, Mazoyer & Tzourio-Mazoyer | 2012 | 10.1016/j.neuroimage.2012.03.073 | Semantic > syntactic decision | 51 | 56 | -2 | -16 |
|  |  |  |  |  | 54 | -16 | -6 |
|  |  |  |  |  | -52 | 2 | -16 |
|  |  |  |  |  | 50 | -34 | 4 |
|  |  |  |  |  | -56 | -4 | -12 |
|  |  |  |  |  | -54 | -16 | -6 |
|  |  |  |  |  | -52 | -40 | 4 |
|  |  |  |  |  | 64 | -34 | 4 |
|  |  |  |  |  | -48 | 32 | -2 |
|  |  |  |  |  | -48 | 24 | 6 |
|  |  |  |  |  | 46 | 28 | -6 |
|  |  |  |  |  | 58 | 30 | 14 |
|  |  |  |  |  | 36 | 26 | -2 |
|  |  |  |  |  | 46 | 20 | 26 |
|  |  |  |  |  | 54 | 26 | 24 |
|  |  |  |  |  | -6 | 56 | 34 |
|  |  |  |  |  | 6 | 54 | 36 |
|  |  |  |  |  | 6 | 58 | 24 |
|  |  |  |  |  | -42 | -60 | 26 |
|  |  |  |  |  | -4 | -50 | 28 |
|  |  |  |  |  | -2 | 46 | -12 |
|  |  |  |  |  | 68 | -46 | 16 |
|  |  |  |  |  | -48 | -56 | 18 |
|  |  |  |  |  | -56 | -54 | 20 |
|  |  |  |  |  | 64 | -56 | 18 |
|  |  |  |  |  | 50 | 16 | -24 |
|  |  |  |  |  | -48 | 12 | -28 |
|  |  |  |  |  | 50 | 24 | -28 |
|  |  |  |  |  | 46 | 22 | -16 |
|  |  |  |  |  | -6 | 32 | 54 |
|  |  |  |  |  | -2 | 42 | 46 |
|  |  |  |  |  | 2 | 14 | 62 |
|  |  |  |  |  | -20 | -8 | -12 |
|  |  |  |  |  | 22 | -6 | -14 |
|  |  |  |  |  | -8 | 48 | 16 |
|  |  |  |  |  | -2 | 38 | -16 |
|  |  |  |  |  | 70 | -34 | 14 |
| Bonhage, Fiebach, Bahlmann & Mueller | 2014 | 10.1162/jocn_a_00566 | Phrases > scrambled phrases | 18 | -9 | 50 | 34 |
|  |  |  |  |  | -6 | 59 | -11 |
|  |  |  |  |  | 30 | 26 | 49 |
|  |  |  |  |  | 6 | 53 | 37 |
|  |  |  |  |  | -3 | 44 | -20 |
|  |  |  |  |  | 0 | 35 | -8 |
|  |  |  |  |  | -48 | -58 | 25 |
|  |  |  |  |  | -6 | -52 | 25 |
|  |  |  |  |  | -54 | -13 | -17 |
|  |  |  |  |  | -39 | 32 | -14 |
|  |  |  |  |  | -27 | -37 | -17 |
|  |  |  |  |  | -21 | -25 | -20 |
|  |  |  |  |  | 54 | -16 | -17 |
|  |  |  |  |  | 57 | -64 | 34 |
|  |  |  |  |  | 27 | -16 | -17 |
|  |  |  |  |  | 54 | -1 | -23 |
|  |  |  |  |  | 66 | -46 | 31 |
|  |  |  |  |  | 36 | -19 | 10 |
|  |  |  |  |  | -21 | -7 | -20 |
|  |  |  |  |  | 54 | -22 | 22 |
|  |  |  |  |  | 24 | -40 | -14 |
|  |  |  |  |  | -39 | -16 | -2 |
|  |  |  |  |  | 33 | -82 | 13 |
|  |  |  |  |  | 24 | -79 | -11 |
|  |  |  |  |  | 15 | -97 | 13 |
|  |  |  |  |  | 48 | -76 | 10 |
|  |  |  |  |  | 45 | -13 | 49 |
|  |  |  |  |  | 27 | -79 | -32 |
| Menz, Blangero, Kunze & Binkofski | 2010 | 10.1016/j.neuroimage.2010.03.050 | Known > unknown objects | 20 | -56 | -29 | 76 |
|  |  |  |  |  | -26 | 28 | 44 |
|  |  |  |  |  | -8 | 34 | -15 |
|  |  |  |  |  | -11 | 49 | 38 |
| Wang, Peelen, Han, Caramazza & Bi | 2016 | 10.1016/j.neuropsychologia.2016.05.007 | Famous faces > face parts | 16 | -3 | -48 | 30 |
|  |  |  |  |  | 60 | -3 | -24 |
|  |  |  |  |  | -54 | -6 | -24 |
|  |  |  |  |  | -6 | 54 | -9 |
|  |  |  |  |  | -54 | -69 | 30 |
|  |  |  |  |  | 21 | -9 | -15 |
|  |  |  |  |  | -33 | 48 | -12 |
|  |  |  |  |  | 54 | -66 | 33 |
|  |  |  |  |  | -15 | -9 | -18 |
|  |  |  |  |  | 39 | 39 | -9 |
| Wang, Peelen, Han, Caramazza & Bi | 2016 | 10.1016/j.neuropsychologia.2016.05.007 | Famous people > scenes | 16 | -3 | -60 | 30 |
|  |  |  |  |  | -9 | 63 | 12 |
|  |  |  |  |  | -48 | -66 | 27 |
|  |  |  |  |  | -21 | 3 | -6 |
|  |  |  |  |  | -63 | -6 | -15 |
|  |  |  |  |  | 18 | 42 | 39 |
| Ryan, Cox, Hayes & Nadel | 2008 | 10.1016/j.neuropsychologia.2008.02.030 | Semantic fluency > crosses | 10 | -23 | -19 | -16 |
|  |  |  |  |  | -19 | -26 | -17 |
|  |  |  |  |  | 21 | -18 | -19 |
|  |  |  |  |  | -4 | 34 | 22 |
|  |  |  |  |  | -29 | 30 | -10 |
|  |  |  |  |  | -26 | 16 | 56 |
|  |  |  |  |  | -16 | -1 | 9 |
|  |  |  |  |  | -6 | -22 | -20 |
|  |  |  |  |  | -4 | -50 | 3 |
|  |  |  |  |  | 7 | -47 | 3 |
|  |  |  |  |  | -30 | -73 | -21 |
|  |  |  |  |  | 21 | -80 | -18 |
|  |  |  |  |  | 33 | 26 | 1 |
|  |  |  |  |  | -45 | 8 | 47 |
|  |  |  |  |  | 14 | 7 | 1 |
|  |  |  |  |  | -5 | -10 | 3 |
| Holle, Gunter, Rueschemeyer, Hennenlotter & Iacoboni | 2008 | 10.1016/j.neuroimage.2007.10.055 | Iconic gesture of dominant meaning > grooming | 17 | -9 | 37 | 40 |
|  |  |  |  |  | 44 | -2 | 37 |
|  |  |  |  |  | -45 | -2 | 33 |
|  |  |  |  |  | 52 | -38 | 29 |
|  |  |  |  |  | -56 | -38 | 29 |
|  |  |  |  |  | 30 | -42 | 39 |
|  |  |  |  |  | 33 | -46 | -5 |
|  |  |  |  |  | -48 | -53 | 12 |
|  |  |  |  |  | -37 | -74 | 18 |
|  |  |  |  |  | -9 | -92 | 4 |
| Holle, Gunter, Rueschemeyer, Hennenlotter & Iacoboni | 2008 | 10.1016/j.neuroimage.2007.10.055 | Iconic gesture of subordinate meaning > grooming | 17 | -45 | 2 | 28 |
|  |  |  |  |  | 38 | -4 | 29 |
|  |  |  |  |  | 49 | -28 | 38 |
|  |  |  |  |  | -53 | -38 | 30 |
|  |  |  |  |  | 36 | -46 | -8 |
|  |  |  |  |  | -29 | -48 | -9 |
|  |  |  |  |  | -45 | -56 | 9 |
|  |  |  |  |  | -50 | -70 | 8 |
| Grindrod, Garnett, Malyutina & den Ouden | 2014 | 10.1016/j.bandl.2014.10.001 | Words > pseudowords | 23 | 45 | 14 | 1 |
|  |  |  |  |  | -63 | -49 | 7 |
|  |  |  |  |  | -39 | 11 | -2 |
|  |  |  |  |  | 48 | -52 | 7 |
|  |  |  |  |  | 15 | 14 | 58 |
|  |  |  |  |  | -30 | 47 | 22 |
|  |  |  |  |  | 0 | -37 | 49 |
|  |  |  |  |  | 3 | 23 | 25 |
|  |  |  |  |  | -33 | 50 | -14 |
|  |  |  |  |  | -12 | 44 | 40 |
|  |  |  |  |  | -21 | -94 | 19 |
| Rodd, Johnsrude & Davis | 2012 | 10.1093/cercor/bhr252 | Speech > SCN | 15 | 62 | -6 | -8 |
|  |  |  |  |  | 50 | 14 | -20 |
|  |  |  |  |  | 66 | -28 | -2 |
|  |  |  |  |  | 56 | -32 | 0 |
|  |  |  |  |  | -62 | -16 | 0 |
|  |  |  |  |  | -50 | 12 | 18 |
|  |  |  |  |  | -60 | -6 | -12 |
|  |  |  |  |  | -60 | -26 | -4 |
|  |  |  |  |  | -58 | -56 | 18 |
|  |  |  |  |  | -44 | 30 | -14 |
|  |  |  |  |  | -54 | 26 | 14 |
|  |  |  |  |  | -42 | -60 | 24 |
|  |  |  |  |  | -26 | -36 | -18 |
|  |  |  |  |  | -40 | -42 | -20 |
|  |  |  |  |  | -30 | -6 | -26 |
|  |  |  |  |  | -8 | 64 | 20 |
| Snijders, Vosse, Kempen, Van Berkum, Petersson & Hagoort | 2009 | 10.1093/cercor/bhn187 | Sentences > word lists | 28 | -54 | 18 | -30 |
|  |  |  |  |  | -56 | -6 | -16 |
|  |  |  |  |  | -62 | -44 | -2 |
|  |  |  |  |  | -52 | 34 | -8 |
|  |  |  |  |  | -58 | -56 | 12 |
|  |  |  |  |  | -58 | 22 | 12 |
|  |  |  |  |  | -44 | -58 | 18 |
|  |  |  |  |  | -26 | -6 | -20 |
|  |  |  |  |  | -44 | -16 | -30 |
|  |  |  |  |  | -26 | -36 | -26 |
|  |  |  |  |  | -18 | 2 | 4 |
|  |  |  |  |  | 54 | 20 | -32 |
|  |  |  |  |  | 56 | 8 | -26 |
|  |  |  |  |  | 52 | -14 | -16 |
|  |  |  |  |  | 62 | -42 | 0 |
|  |  |  |  |  | 56 | 36 | -10 |
|  |  |  |  |  | 60 | 34 | 4 |
|  |  |  |  |  | 62 | 28 | 10 |
|  |  |  |  |  | -4 | 54 | -20 |
|  |  |  |  |  | -8 | 60 | 28 |
|  |  |  |  |  | -38 | -2 | -50 |
|  |  |  |  |  | -6 | -62 | 2 |
|  |  |  |  |  | -12 | -46 | 34 |
|  |  |  |  |  | 30 | -32 | -32 |
| Chiao, Harada, Oby, Li, Parrish & Bridge | 2009 | 10.1016/j.neuropsychologia.2008.09.023 | Uniform status judgement > colour change detection | 12 | 33 | -62 | 42 |
|  |  |  |  |  | 56 | -53 | -6 |
|  |  |  |  |  | -48 | 7 | 33 |
|  |  |  |  |  | -3 | 14 | 52 |
|  |  |  |  |  | -50 | -35 | 49 |
|  |  |  |  |  | 36 | 23 | -4 |
|  |  |  |  |  | -33 | 20 | -1 |
|  |  |  |  |  | -30 | -69 | -44 |
|  |  |  |  |  | -42 | -67 | 1 |
|  |  |  |  |  | 50 | 14 | 38 |
|  |  |  |  |  | 33 | 2 | 50 |
|  |  |  |  |  | 0 | -33 | 32 |
| Chiao, Harada, Oby, Li, Parrish & Bridge | 2009 | 10.1016/j.neuropsychologia.2008.09.023 | Face status judgement > colour change detection | 12 | 45 | -42 | -15 |
|  |  |  |  |  | 33 | -62 | 42 |
|  |  |  |  |  | -3 | 17 | 49 |
|  |  |  |  |  | -30 | -56 | 53 |
|  |  |  |  |  | 36 | 23 | -4 |
|  |  |  |  |  | -36 | 20 | -1 |
|  |  |  |  |  | 53 | 24 | 24 |
|  |  |  |  |  | 3 | -33 | 29 |
| Chiao, Harada, Oby, Li, Parrish & Bridge | 2009 | 10.1016/j.neuropsychologia.2008.09.023 | Car status judgement > colour change detection | 12 | 0 | 31 | 37 |
|  |  |  |  |  | 36 | 23 | -7 |
|  |  |  |  |  | 33 | -62 | 42 |
|  |  |  |  |  | 33 | -59 | 53 |
|  |  |  |  |  | -36 | 20 | -1 |
|  |  |  |  |  | 36 | -39 | -17 |
|  |  |  |  |  | -9 | -77 | -18 |
|  |  |  |  |  | -36 | -50 | -14 |
|  |  |  |  |  | 56 | 25 | 29 |
|  |  |  |  |  | -30 | -61 | 56 |
|  |  |  |  |  | -45 | 52 | 0 |
|  |  |  |  |  | -33 | -50 | 47 |
| Jeon, Lee, Kim & Cho | 2009 | 10.1016/j.neuroimage.2009.06.049 | Synonym generation > pseudowords | 16 | -42 | 22 | 22 |
|  |  |  |  |  | -28 | -62 | 51 |
| Jeon, Lee, Kim & Cho | 2009 | 10.1016/j.neuroimage.2009.06.049 | Antonym generation > pseudowords | 16 | -54 | 30 | 21 |
| Jeon, Lee, Kim & Cho | 2009 | 10.1016/j.neuroimage.2009.06.049 | Words > pseudowords | 16 | -42 | 22 | 18 |
| Jeon, Lee, Kim & Cho | 2009 | 10.1016/j.neuroimage.2009.06.049 | Words > pseudowords | 16 | -42 | 22 | 22 |
|  |  |  |  |  | -44 | 22 | -3 |
|  |  |  |  |  | -28 | -56 | 63 |
| Jeon, Lee, Kim & Cho | 2009 | 10.1016/j.neuroimage.2009.06.049 | Words > pseudowords | 16 | -33 | -53 | 54 |
| Metz-Lutz, Bressan, Heider & Otzenberger | 2010 | 10.3389/fnhum.2010.00059 | Events > not events | 11 | 55 | -29 | -16 |
|  |  |  |  |  | 71 | -54 | 13 |
|  |  |  |  |  | 69 | -40 | 27 |
|  |  |  |  |  | -55 | 36 | -18 |
|  |  |  |  |  | -27 | 25 | -31 |
|  |  |  |  |  | -51 | 23 | -4 |
|  |  |  |  |  | -65 | -45 | 35 |
|  |  |  |  |  | -61 | -69 | 6 |
|  |  |  |  |  | -65 | -56 | 29 |
|  |  |  |  |  | 38 | -16 | -13 |
|  |  |  |  |  | 34 | -19 | -1 |
|  |  |  |  |  | 36 | 6 | -8 |
| Rogalsky & Hickok | 2009 | 10.1093/cercor/bhn126 | Sentences > word lists | 14 | -50 | 17 | -26 |
|  |  |  |  |  | 57 | 19 | -30 |
|  |  |  |  |  | -54 | 13 | -16 |
| Rogalsky & Hickok | 2009 | 10.1093/cercor/bhn126 | Semantic > syntactic error detection | 14 | -50 | -19 | -7 |
|  |  |  |  |  | -43 | 2 | 41 |
|  |  |  |  |  | -30 | 32 | 17 |
|  |  |  |  |  | 62 | 9 | -29 |
|  |  |  |  |  | 51 | 5 | 18 |
| Sun, Xue, Zhang, Zuo, Chen, Wang, Martin, Wang, Chen, He & Wang | 2017 | 10.1016/j.jmr.2016.12.012 | Semantic > orthography judgement | 11 | -58 | -44 | -8 |
|  |  |  |  |  | -64 | -40 | -10 |
|  |  |  |  |  | -60 | -30 | -14 |
| Liu, Deng, Peng, Cao, Ding, Jin, Zeng, Li, Zhu, Fan, Deng, Bolger & Booth | 2009 | 10.1162/jocn.2009.21141 | Words > slashes | 16 | -52 | 34 | 20 |
|  |  |  |  |  | -48 | 26 | 20 |
|  |  |  |  |  | -50 | 16 | 40 |
|  |  |  |  |  | -36 | -48 | -24 |
|  |  |  |  |  | -28 | -100 | -8 |
|  |  |  |  |  | -4 | 20 | 48 |
|  |  |  |  |  | -36 | -12 | -36 |
|  |  |  |  |  | -38 | -58 | -24 |
|  |  |  |  |  | 28 | -92 | -16 |
|  |  |  |  |  | 26 | -96 | -8 |
|  |  |  |  |  | 36 | -64 | -24 |
|  |  |  |  |  | 34 | 24 | 0 |
| Liu, Deng, Peng, Cao, Ding, Jin, Zeng, Li, Zhu, Fan, Deng, Bolger & Booth | 2009 | 10.1162/jocn.2009.21141 | Words > slashes | 16 | -46 | 32 | 16 |
|  |  |  |  |  | -44 | 22 | 8 |
|  |  |  |  |  | -30 | -64 | 36 |
|  |  |  |  |  | -38 | -44 | -28 |
|  |  |  |  |  | -36 | -62 | -28 |
|  |  |  |  |  | 36 | 20 | -8 |
|  |  |  |  |  | 40 | 20 | 8 |
|  |  |  |  |  | 28 | -96 | -4 |
|  |  |  |  |  | 8 | -50 | -32 |
|  |  |  |  |  | 38 | -58 | -28 |
|  |  |  |  |  | 18 | -30 | 20 |
|  |  |  |  |  | 18 | 6 | 20 |
| Liu, Deng, Peng, Cao, Ding, Jin, Zeng, Li, Zhu, Fan, Deng, Bolger & Booth | 2009 | 10.1162/jocn.2009.21141 | Words > tones | 16 | -30 | 22 | -8 |
|  |  |  |  |  | -2 | -84 | 0 |
|  |  |  |  |  | -50 | -4 | -12 |
|  |  |  |  |  | -60 | -24 | 0 |
|  |  |  |  |  | -10 | 0 | 8 |
|  |  |  |  |  | -8 | -18 | 12 |
|  |  |  |  |  | 32 | 24 | -4 |
|  |  |  |  |  | 50 | 34 | 28 |
|  |  |  |  |  | 2 | 24 | 44 |
|  |  |  |  |  | 6 | 32 | 40 |
|  |  |  |  |  | 60 | -12 | -8 |
|  |  |  |  |  | 50 | 12 | -16 |
| Liu, Deng, Peng, Cao, Ding, Jin, Zeng, Li, Zhu, Fan, Deng, Bolger & Booth | 2009 | 10.1162/jocn.2009.21141 | Words > tones | 16 | -46 | 32 | 20 |
|  |  |  |  |  | -46 | 10 | 20 |
|  |  |  |  |  | -52 | -2 | 12 |
|  |  |  |  |  | -62 | -24 | 4 |
|  |  |  |  |  | -48 | 4 | 44 |
|  |  |  |  |  | 2 | 20 | 52 |
|  |  |  |  |  | 42 | 14 | -28 |
|  |  |  |  |  | 0 | -78 | -20 |
| Diaz & McCarthy | 2009 | 10.1016/j.brainres.2009.05.043 | Words > pseudowords | 16 | -54 | -60 | 36 |
|  |  |  |  |  | -66 | -16 | -12 |
|  |  |  |  |  | 66 | -10 | -16 |
|  |  |  |  |  | 74 | -26 | -5 |
|  |  |  |  |  | 66 | -38 | -12 |
|  |  |  |  |  | -40 | 12 | -42 |
|  |  |  |  |  | -47 | 34 | -14 |
|  |  |  |  |  | 34 | 36 | -20 |
|  |  |  |  |  | 24 | -16 | -24 |
|  |  |  |  |  | -11 | 52 | 23 |
|  |  |  |  |  | -10 | 36 | 56 |
| Taminato, Miura, Sugiura & Kawashima | 2014 | 10.1016/j.neures.2014.09.001 | Object recognition > control task | 35 | -34 | -86 | 18 |
|  |  |  |  |  | 42 | -78 | 14 |
|  |  |  |  |  | -16 | -70 | -6 |
|  |  |  |  |  | -46 | -66 | -14 |
|  |  |  |  |  | 44 | -56 | -16 |
|  |  |  |  |  | -34 | -50 | -16 |
|  |  |  |  |  | -32 | -36 | -24 |
|  |  |  |  |  | 34 | -52 | -10 |
|  |  |  |  |  | 32 | -38 | -20 |
|  |  |  |  |  | 32 | -48 | -12 |
|  |  |  |  |  | -40 | 10 | 24 |
| Taminato, Miura, Sugiura & Kawashima | 2014 | 10.1016/j.neures.2014.09.001 | Object recognition > control task | 35 | -32 | -88 | 16 |
|  |  |  |  |  | 40 | -82 | 20 |
|  |  |  |  |  | -8 | -80 | -10 |
|  |  |  |  |  | -46 | -54 | -16 |
|  |  |  |  |  | 40 | -64 | -14 |
|  |  |  |  |  | -32 | -64 | -12 |
|  |  |  |  |  | 32 | -64 | -12 |
|  |  |  |  |  | 32 | -42 | -22 |
|  |  |  |  |  | -38 | 12 | 28 |
|  |  |  |  |  | -6 | 20 | 44 |
| Lane, Kanjlia, Omaki & Bedny | 2015 | 10.1523/JNEUROSCI.1256-15.2015 | Sentences > pseudowords | 18 | -46 | 13 | -30 |
|  |  |  |  |  | -32 | -36 | -15 |
|  |  |  |  |  | -44 | 26 | -14 |
|  |  |  |  |  | -5 | -60 | 33 |
|  |  |  |  |  | -6 | 54 | 31 |
|  |  |  |  |  | -42 | 2 | 50 |
|  |  |  |  |  | -46 | -29 | 45 |
| Assadollahi, Meinzer, Flaisch, Obleser & Rockstroh | 2009 | 10.1186/1471-2202-10-3 | Phrases > words | 20 | -53 | -5 | -15 |
|  |  |  |  |  | 55 | -1 | -18 |
| Wright, Mechelli, Noppeney, Veltman, Rombouts, Glensman, Haynes & Price | 2008 | 10.1002/hbm.20443 | Semantic > perceptual matching | 34 | -42 | -56 | -14 |
|  |  |  |  |  | -50 | -58 | -12 |
| Wright, Mechelli, Noppeney, Veltman, Rombouts, Glensman, Haynes & Price | 2008 | 10.1002/hbm.20443 | Semantic > perceptual matching | 34 | -48 | -58 | -14 |
|  |  |  |  |  | -52 | -52 | -10 |
| Wright, Mechelli, Noppeney, Veltman, Rombouts, Glensman, Haynes & Price | 2008 | 10.1002/hbm.20443 | Semantic > perceptual matching | 34 | -46 | -52 | -10 |
|  |  |  |  |  | -54 | -58 | -10 |
| Leung & Alain | 2011 | 10.1016/j.neuroimage.2010.12.055 | Semantic > location matching | 16 | -5 | 12 | 57 |
|  |  |  |  |  | -47 | 28 | -16 |
|  |  |  |  |  | -27 | 24 | 11 |
|  |  |  |  |  | -51 | -41 | 7 |
|  |  |  |  |  | 51 | 25 | 17 |
|  |  |  |  |  | 12 | -87 | -30 |
| Obleser & Kotz | 2010 | 10.1093/cercor/bhp128 | Intelligibility parameter | 16 | 62 | -6 | -4 |
|  |  |  |  |  | -60 | -8 | -6 |
|  |  |  |  |  | -46 | -64 | 38 |
| McGettigan, Faulkner, Altarelli, Obleser, Baverstock & Scott | 2012 | 10.1016/j.neuropsychologia.2012.01.010 | Intelligibility parameter | 26 | -57 | -36 | 3 |
|  |  |  |  |  | -57 | -12 | -6 |
|  |  |  |  |  | -54 | -51 | 18 |
|  |  |  |  |  | -54 | 0 | -9 |
|  |  |  |  |  | -51 | 9 | -18 |
|  |  |  |  |  | -45 | 15 | -27 |
|  |  |  |  |  | -48 | -18 | -12 |
|  |  |  |  |  | -51 | 30 | 3 |
|  |  |  |  |  | -42 | 15 | 21 |
|  |  |  |  |  | -51 | 15 | 18 |
|  |  |  |  |  | 48 | 6 | -21 |
|  |  |  |  |  | 48 | -36 | 3 |
|  |  |  |  |  | 57 | -6 | -9 |
|  |  |  |  |  | 51 | -18 | -9 |
|  |  |  |  |  | 60 | -27 | 3 |
|  |  |  |  |  | -42 | -39 | -18 |
|  |  |  |  |  | -39 | -30 | -18 |
|  |  |  |  |  | 45 | 30 | -6 |
| McGettigan, Faulkner, Altarelli, Obleser, Baverstock & Scott | 2012 | 10.1016/j.neuropsychologia.2012.01.010 | Auditory degradation parameter | 26 | -57 | -12 | -6 |
|  |  |  |  |  | -51 | -42 | 3 |
|  |  |  |  |  | -60 | -33 | 3 |
|  |  |  |  |  | -57 | -3 | -9 |
|  |  |  |  |  | -51 | 9 | -18 |
|  |  |  |  |  | -60 | -51 | 18 |
|  |  |  |  |  | -42 | -48 | 21 |
|  |  |  |  |  | -48 | 27 | -3 |
|  |  |  |  |  | 51 | 6 | -21 |
|  |  |  |  |  | 51 | -18 | -9 |
|  |  |  |  |  | 54 | -9 | -12 |
|  |  |  |  |  | 36 | -33 | 18 |
|  |  |  |  |  | 42 | -27 | 9 |
|  |  |  |  |  | -6 | -63 | 51 |
|  |  |  |  |  | -45 | 60 | 45 |
|  |  |  |  |  | -36 | -78 | 39 |
|  |  |  |  |  | -24 | -66 | 54 |
|  |  |  |  |  | -12 | -81 | 48 |
|  |  |  |  |  | -3 | -30 | 36 |
|  |  |  |  |  | 6 | -27 | 39 |
|  |  |  |  |  | 9 | -27 | 51 |
|  |  |  |  |  | -27 | 24 | 51 |
|  |  |  |  |  | -30 | 42 | 36 |
|  |  |  |  |  | -42 | 27 | 42 |
|  |  |  |  |  | -39 | 33 | 33 |
|  |  |  |  |  | -39 | 12 | 54 |
|  |  |  |  |  | -36 | 51 | 3 |
|  |  |  |  |  | -27 | 57 | 6 |
|  |  |  |  |  | -33 | 51 | -9 |
|  |  |  |  |  | -24 | 57 | 24 |
|  |  |  |  |  | 57 | -51 | 42 |
|  |  |  |  |  | 27 | 48 | 39 |
|  |  |  |  |  | 24 | 27 | 48 |
|  |  |  |  |  | 33 | 36 | 42 |
|  |  |  |  |  | 42 | 33 | 36 |
|  |  |  |  |  | 27 | 18 | 39 |
|  |  |  |  |  | -3 | -99 | 0 |
|  |  |  |  |  | 33 | -90 | -15 |
|  |  |  |  |  | -42 | 15 | 21 |
|  |  |  |  |  | -36 | -75 | -18 |
|  |  |  |  |  | -36 | -66 | -21 |
|  |  |  |  |  | -45 | -78 | -12 |
|  |  |  |  |  | -33 | -30 | 18 |
|  |  |  |  |  | -36 | -39 | 18 |
|  |  |  |  |  | 60 | -30 | 15 |
|  |  |  |  |  | 54 | 30 | -3 |
|  |  |  |  |  | 57 | 24 | 3 |
|  |  |  |  |  | 45 | 30 | -6 |
|  |  |  |  |  | 42 | -72 | 42 |
|  |  |  |  |  | 42 | -60 | 45 |
|  |  |  |  |  | 39 | -60 | 54 |
| McGettigan, Faulkner, Altarelli, Obleser, Baverstock & Scott | 2012 | 10.1016/j.neuropsychologia.2012.01.010 | More > less visually degraded stimuli | 26 | 27 | -96 | 0 |
|  |  |  |  |  | -21 | -99 | -3 |
|  |  |  |  |  | 39 | -81 | 12 |
|  |  |  |  |  | 48 | -39 | 9 |
|  |  |  |  |  | 57 | -3 | -9 |
|  |  |  |  |  | 54 | -15 | -9 |
|  |  |  |  |  | -36 | -90 | 12 |
|  |  |  |  |  | -30 | -87 | 18 |
|  |  |  |  |  | -39 | -66 | -9 |
|  |  |  |  |  | -48 | -81 | 0 |
|  |  |  |  |  | -51 | -72 | -9 |
|  |  |  |  |  | -9 | 51 | 0 |
|  |  |  |  |  | 45 | -48 | -18 |
|  |  |  |  |  | 39 | -39 | -21 |
|  |  |  |  |  | -51 | -51 | 6 |
|  |  |  |  |  | -54 | -42 | 6 |
|  |  |  |  |  | -60 | -48 | 15 |
|  |  |  |  |  | 36 | -69 | -9 |
|  |  |  |  |  | 27 | -66 | -12 |
|  |  |  |  |  | 48 | -69 | -9 |
|  |  |  |  |  | -27 | 33 | 39 |
|  |  |  |  |  | -45 | -6 | 15 |
|  |  |  |  |  | -36 | -9 | 9 |
|  |  |  |  |  | 36 | 45 | 12 |
|  |  |  |  |  | -39 | -48 | -18 |
|  |  |  |  |  | 42 | -42 | 48 |
|  |  |  |  |  | -21 | -69 | -6 |
|  |  |  |  |  | -6 | -72 | 39 |
|  |  |  |  |  | -15 | -66 | 36 |
|  |  |  |  |  | 18 | -6 | -12 |
| McGettigan, Faulkner, Altarelli, Obleser, Baverstock & Scott | 2012 | 10.1016/j.neuropsychologia.2012.01.010 | More > less visually degraded stimuli | 26 | 27 | -96 | 0 |
|  |  |  |  |  | 15 | -99 | 9 |
|  |  |  |  |  | -12 | -102 | -6 |
|  |  |  |  |  | -24 | -96 | -9 |
|  |  |  |  |  | -24 | -81 | 18 |
|  |  |  |  |  | -30 | -90 | 21 |
|  |  |  |  |  | -54 | -51 | 6 |
|  |  |  |  |  | 36 | -81 | 12 |
|  |  |  |  |  | 39 | -39 | -21 |
|  |  |  |  |  | 51 | -45 | 12 |
|  |  |  |  |  | 51 | -36 | 0 |
|  |  |  |  |  | 54 | -3 | -12 |
| McGettigan, Faulkner, Altarelli, Obleser, Baverstock & Scott | 2012 | 10.1016/j.neuropsychologia.2012.01.010 | Intelligibility parameter | 26 | 27 | -96 | 0 |
|  |  |  |  |  | 12 | -99 | 9 |
|  |  |  |  |  | -21 | -99 | -3 |
|  |  |  |  |  | -12 | -93 | -3 |
|  |  |  |  |  | 51 | -45 | 9 |
|  |  |  |  |  | 60 | -33 | 0 |
|  |  |  |  |  | 63 | -36 | 12 |
|  |  |  |  |  | -51 | 30 | 3 |
|  |  |  |  |  | -54 | 27 | 15 |
|  |  |  |  |  | -42 | -54 | -18 |
|  |  |  |  |  | -39 | -45 | -24 |
|  |  |  |  |  | -33 | -63 | -24 |
|  |  |  |  |  | -6 | -18 | 57 |
|  |  |  |  |  | -3 | -3 | 54 |
|  |  |  |  |  | 42 | -51 | -18 |
|  |  |  |  |  | 33 | -45 | -18 |
|  |  |  |  |  | 57 | -3 | -9 |
|  |  |  |  |  | 57 | -9 | -15 |
|  |  |  |  |  | -51 | -51 | 15 |
| Takeichi, Koyama, Terao, Takeuchi, Toyosawa & Murohashi | 2010 | 10.1016/j.neuroimage.2009.10.063 | Speech > reversed speech | 23 | -64 | -52 | 0 |
|  |  |  |  |  | -52 | -2 | -26 |
|  |  |  |  |  | -40 | -60 | 26 |
|  |  |  |  |  | -46 | 2 | 46 |
|  |  |  |  |  | 62 | -12 | -18 |
| Takeichi, Koyama, Terao, Takeuchi, Toyosawa & Murohashi | 2010 | 10.1016/j.neuroimage.2009.10.063 | Speech > modulated speech | 23 | -64 | -12 | -10 |
|  |  |  |  |  | -60 | -54 | 0 |
|  |  |  |  |  | -38 | -70 | 22 |
|  |  |  |  |  | -20 | 0 | -16 |
|  |  |  |  |  | 0 | -60 | 12 |
|  |  |  |  |  | 0 | 2 | 4 |
|  |  |  |  |  | -2 | -16 | 8 |
|  |  |  |  |  | 58 | 6 | -12 |
| Pallier, Devauchelle & Dehaene | 2011 | 10.1073/pnas.1018711108 | Longer > shorter phrase | 40 | -54 | -12 | -12 |
|  |  |  |  |  | -48 | 15 | -27 |
|  |  |  |  |  | -39 | -57 | 18 |
|  |  |  |  |  | -48 | -45 | 3 |
|  |  |  |  |  | -51 | 30 | 6 |
|  |  |  |  |  | -54 | 21 | 15 |
|  |  |  |  |  | -18 | 6 | 12 |
|  |  |  |  |  | -6 | 54 | 36 |
|  |  |  |  |  | -27 | -3 | -39 |
|  |  |  |  |  | 51 | 0 | -21 |
| Pallier, Devauchelle & Dehaene | 2011 | 10.1073/pnas.1018711108 | Length of real > pseudoword sentences | 40 | -45 | -66 | 24 |
|  |  |  |  |  | -39 | -57 | 21 |
|  |  |  |  |  | -48 | 15 | -30 |
|  |  |  |  |  | -54 | -12 | -12 |
|  |  |  |  |  | -40 | 3 | 15 |
| Mellem, Jasmin, Peng & Martin | 2016 | 10.1016/j.neuropsychologia.2016.06.019 | Longer > shorter phrase | 20 | -48 | 16 | -22 |
|  |  |  |  |  | -56 | 31 | 9 |
|  |  |  |  |  | 47 | 13 | -27 |
|  |  |  |  |  | -51 | 22 | -17 |
| Guediche, Reilly, Santiago, Laurent & Blumstein | 2016 | 10.1016/j.cortex.2016.03.014 | Intelligibility based on cue | 16 | -35 | -91 | 14 |
|  |  |  |  |  | -42 | -86 | 1 |
|  |  |  |  |  | -52 | -64 | 3 |
|  |  |  |  |  | -32 | -76 | 4 |
|  |  |  |  |  | 39 | 41 | 13 |
|  |  |  |  |  | 48 | 4 | 33 |
|  |  |  |  |  | -45 | 24 | 40 |
|  |  |  |  |  | -45 | 4 | 32 |
|  |  |  |  |  | 56 | -46 | 48 |
|  |  |  |  |  | 35 | -47 | 42 |
|  |  |  |  |  | -52 | 42 | -2 |
|  |  |  |  |  | -32 | -57 | 41 |
|  |  |  |  |  | -42 | 19 | -7 |
|  |  |  |  |  | -62 | -42 | -4 |
|  |  |  |  |  | -3 | -22 | -7 |
|  |  |  |  |  | -10 | 11 | 3 |
|  |  |  |  |  | 26 | -76 | -32 |
|  |  |  |  |  | 16 | -49 | -15 |
| Guediche, Reilly, Santiago, Laurent & Blumstein | 2016 | 10.1016/j.cortex.2016.03.014 | Related > repeated sentence | 16 | 68 | -24 | 6 |
|  |  |  |  |  | -42 | -16 | -3 |
|  |  |  |  |  | -52 | 29 | -1 |
|  |  |  |  |  | -62 | -14 | -13 |
|  |  |  |  |  | -45 | 11 | -26 |
| Guediche, Reilly, Santiago, Laurent & Blumstein | 2016 | 10.1016/j.cortex.2016.03.014 | Unrelated > repeated sentence | 16 | 68 | -24 | 6 |
|  |  |  |  |  | -42 | -24 | 14 |
|  |  |  |  |  | 36 | -42 | -6 |
|  |  |  |  |  | 7 | -9 | 2 |
|  |  |  |  |  | -35 | -91 | 14 |
|  |  |  |  |  | -45 | -54 | 41 |
|  |  |  |  |  | -45 | -35 | -5 |
|  |  |  |  |  | -43 | 16 | -37 |
|  |  |  |  |  | -36 | -61 | -36 |
|  |  |  |  |  | 24 | -22 | 26 |
| Sharp, Awad, Warren, Wise, Vigliocco & Scott | 2010 | 10.1002/hbm.20871 | Normal > rotated speech | 12 | -28 | -36 | -8 |
|  |  |  |  |  | -36 | -28 | -24 |
|  |  |  |  |  | -42 | -12 | -36 |
|  |  |  |  |  | -38 | -8 | -36 |
|  |  |  |  |  | -38 | 36 | -16 |
|  |  |  |  |  | -12 | 50 | 50 |
|  |  |  |  |  | -32 | 12 | 46 |
|  |  |  |  |  | -32 | -78 | 48 |
|  |  |  |  |  | -38 | -74 | 24 |
|  |  |  |  |  | 38 | -74 | -40 |
| Thothathiri, Rattinger & Trivedi | 2017 | 10.1080/17588928.2015.1090421 | Sentence > word generation | 14 | -18 | -78 | 46 |
|  |  |  |  |  | -16 | -82 | 50 |
|  |  |  |  |  | 20 | -84 | 42 |
|  |  |  |  |  | 10 | 10 | 50 |
|  |  |  |  |  | 8 | 14 | 36 |
|  |  |  |  |  | 6 | 8 | 48 |
|  |  |  |  |  | -46 | 0 | 44 |
|  |  |  |  |  | -44 | 10 | 26 |
|  |  |  |  |  | -54 | -2 | 42 |
|  |  |  |  |  | 40 | -6 | 52 |
|  |  |  |  |  | 28 | -6 | 56 |
|  |  |  |  |  | 54 | 6 | 40 |
| Soch, Deserno, Assmann, Barman, Walter, Richardson-Klavehn & Schott | 2017 | 10.1093/cercor/bhw206 | Attribute > phonological judgement | 110 | -9 | 50 | 37 |
|  |  |  |  |  | -42 | 23 | -14 |
|  |  |  |  |  | -48 | -67 | 25 |
|  |  |  |  |  | -3 | -52 | 28 |
|  |  |  |  |  | 54 | -64 | 28 |
|  |  |  |  |  | -54 | -7 | -23 |
|  |  |  |  |  | 50 | -7 | -20 |
|  |  |  |  |  | 0 | 11 | -11 |
|  |  |  |  |  | 0 | -19 | 37 |
| Chou, Chen, Wu & Booth | 2009 | 10.1007/s00221-009-1942-y | Related words > false font | 31 | -48 | 30 | -3 |
|  |  |  |  |  | -47 | 24 | 12 |
|  |  |  |  |  | -6 | 36 | 39 |
|  |  |  |  |  | -57 | -46 | -3 |
| Chou, Chen, Wu & Booth | 2009 | 10.1007/s00221-009-1942-y | Unrelated words > false font | 31 | -48 | 30 | -3 |
|  |  |  |  |  | -45 | 24 | 14 |
|  |  |  |  |  | -60 | -45 | 0 |
| Davis, Ford, Kherif & Johnsrude | 2011 | 10.1097/JOM.0b013e31820805d5 | Intelligibility parameter | 12 | -60 | -42 | 0 |
|  |  |  |  |  | -58 | -10 | -8 |
|  |  |  |  |  | -62 | -18 | 4 |
|  |  |  |  |  | -46 | 18 | -26 |
|  |  |  |  |  | -54 | -56 | 14 |
|  |  |  |  |  | -46 | 26 | -6 |
|  |  |  |  |  | -48 | 18 | 16 |
|  |  |  |  |  | -50 | 0 | -24 |
|  |  |  |  |  | 58 | 10 | -14 |
|  |  |  |  |  | 54 | -2 | -22 |
|  |  |  |  |  | 50 | 16 | -20 |
|  |  |  |  |  | 54 | -34 | -2 |
|  |  |  |  |  | 66 | -18 | -8 |
|  |  |  |  |  | 60 | -8 | 0 |
|  |  |  |  |  | 46 | -22 | 10 |
|  |  |  |  |  | 66 | -32 | -4 |
|  |  |  |  |  | 58 | -22 | -16 |
|  |  |  |  |  | -24 | 4 | -6 |
|  |  |  |  |  | -20 | -10 | -14 |
|  |  |  |  |  | 20 | -14 | -16 |
|  |  |  |  |  | 26 | 0 | -12 |
|  |  |  |  |  | -8 | 30 | -2 |
|  |  |  |  |  | -38 | -38 | -22 |
|  |  |  |  |  | -44 | -48 | -20 |
|  |  |  |  |  | -32 | -10 | -24 |
|  |  |  |  |  | -22 | -26 | -6 |
| Moberget, Gullesen, Andersson, Ivry & Endestad | 2014 | 10.1523/JNEUROSCI.2264-13.2014 | Incongruent > scrambled sentence | 32 | 22 | -78 | -29 |
|  |  |  |  |  | -16 | -80 | -29 |
|  |  |  |  |  | 6 | -54 | -41 |
|  |  |  |  |  | -8 | -20 | -7 |
|  |  |  |  |  | -51 | 32 | 8 |
|  |  |  |  |  | -12 | 17 | 5 |
|  |  |  |  |  | -48 | -4 | -19 |
|  |  |  |  |  | -33 | -37 | -19 |
|  |  |  |  |  | -39 | -58 | 23 |
|  |  |  |  |  | 39 | 32 | -7 |
|  |  |  |  |  | 48 | -31 | -7 |
|  |  |  |  |  | 0 | 44 | 50 |
| Moberget, Gullesen, Andersson, Ivry & Endestad | 2014 | 10.1523/JNEUROSCI.2264-13.2014 | Congruent > scrambled sentence | 32 | 32 | -74 | -37 |
|  |  |  |  |  | 8 | -54 | -45 |
|  |  |  |  |  | -20 | -10 | -3 |
|  |  |  |  |  | 6 | -16 | -7 |
|  |  |  |  |  | -9 | 17 | 5 |
|  |  |  |  |  | -30 | 20 | -13 |
|  |  |  |  |  | -57 | -4 | -19 |
|  |  |  |  |  | 33 | 20 | -7 |
|  |  |  |  |  | -60 | -22 | 32 |
|  |  |  |  |  | -9 | 53 | 38 |
|  |  |  |  |  | 57 | -22 | 44 |
| Egidi & Caramazza | 2016 | 10.1162/jocn_a_00982 | Semantic judgement > passive listening | 28 | -26 | -57 | 55 |
|  |  |  |  |  | -21 | -80 | -4 |
|  |  |  |  |  | -65 | -31 | 3 |
|  |  |  |  |  | -50 | 22 | 18 |
|  |  |  |  |  | -42 | 8 | 47 |
|  |  |  |  |  | -12 | 53 | -11 |
|  |  |  |  |  | -1 | -25 | -14 |
|  |  |  |  |  | 33 | -83 | -9 |
|  |  |  |  |  | 40 | 21 | 21 |
|  |  |  |  |  | 39 | -77 | 23 |
|  |  |  |  |  | 50 | -34 | -8 |
|  |  |  |  |  | 42 | -50 | 44 |
|  |  |  |  |  | 52 | 23 | 3 |
|  |  |  |  |  | 11 | -66 | 57 |
|  |  |  |  |  | 11 | 15 | 45 |
| Clos, Langner, Meyer, Oechslin, Zilles & Eickhoff | 2014 | 10.1002/hbm.22151 | Intelligibilty based on cue > unintelligible | 29 | -54 | 20 | 16 |
|  |  |  |  |  | -62 | -26 | -10 |
|  |  |  |  |  | -46 | -62 | 28 |
| Van Ettinger-Veenstra, McAllister, Lundberg, Karlsson & Engstrom | 2016 | 10.3389/fnhum.2016.00110 | Words > symbols | 27 | -44 | -8 | 58 |
|  |  |  |  |  | 46 | 18 | 30 |
|  |  |  |  |  | 38 | 36 | -14 |
|  |  |  |  |  | -58 | -36 | 2 |
|  |  |  |  |  | -42 | -48 | -20 |
|  |  |  |  |  | -12 | -50 | 2 |
|  |  |  |  |  | 6 | 20 | 42 |
| Graves, Binder, Desai, Conant & Seidenberg | 2010 | 10.1016/j.neuroimage.2010.06.055 | Phrases > words | 23 | 58 | -38 | 30 |
| Graves, Binder, Desai, Conant & Seidenberg | 2010 | 10.1016/j.neuroimage.2010.06.055 | Phrases > words | 22 | 20 | 52 | 15 |
|  |  |  |  |  | 0 | 47 | 15 |
|  |  |  |  |  | 1 | 52 | -4 |
|  |  |  |  |  | -9 | 69 | -6 |
|  |  |  |  |  | 26 | 54 | 27 |
|  |  |  |  |  | 8 | 55 | 39 |
|  |  |  |  |  | -6 | -44 | 42 |
|  |  |  |  |  | 62 | -59 | 20 |
|  |  |  |  |  | 61 | -51 | 43 |
|  |  |  |  |  | 7 | 43 | 50 |
|  |  |  |  |  | -1 | 5 | 32 |
|  |  |  |  |  | -27 | -25 | 48 |
| Adank, Davis & Hagoort | 2012 | 10.1016/j.neuropsychologia.2011.10.024 | Speech > noisy speech | 26 | -40 | -22 | 12 |
|  |  |  |  |  | -40 | -20 | 54 |
|  |  |  |  |  | 34 | -22 | 10 |
|  |  |  |  |  | -36 | 22 | -26 |
|  |  |  |  |  | 34 | 42 | -14 |
|  |  |  |  |  | -54 | -44 | -6 |
|  |  |  |  |  | 14 | -50 | -20 |
|  |  |  |  |  | -8 | 50 | 40 |
|  |  |  |  |  | 66 | -42 | -8 |
|  |  |  |  |  | -54 | -44 | -6 |
| Wright, Randall, Marslen-Wilson & Tyler | 2011 | 10.1162/jocn.2010.21450 | Speech > musical rain | 14 | -60 | -3 | -6 |
|  |  |  |  |  | -60 | -30 | 3 |
|  |  |  |  |  | -63 | -18 | -3 |
|  |  |  |  |  | 63 | -18 | -3 |
| Wright, Randall, Marslen-Wilson & Tyler | 2011 | 10.1162/jocn.2010.21450 | Lexical decision > passive listening | 14 | 60 | -6 | -3 |
|  |  |  |  |  | 54 | -27 | 3 |
| Hwang, Palmer, Basho, Zadra & Muller | 2009 | 10.1016/j.neuroimage.2009.06.042 | Fluency generation > production baseline | 13 | -13 | -15 | 14 |
|  |  |  |  |  | -16 | -4 | 11 |
|  |  |  |  |  | -36 | 21 | 13 |
|  |  |  |  |  | -3 | 24 | 37 |
|  |  |  |  |  | -24 | 9 | 53 |
|  |  |  |  |  | -8 | -60 | -1 |

*N.B. Contrasts within the same study conducted using the same sample were combined before they were entered into the meta-analysis.*

Supplementary Table 2. Data included in the phonology meta-analysis.

| **Author(s)** | **Year** | **DOI** | **Contrast included** | **Type** | **N** | **MNI coordinates** | | |
| --- | --- | --- | --- | --- | --- | --- | --- | --- |
|  |  |  |  |  |  | **X** | **Y** | **Z** |
| Baciu et al. | 2005 | 10.1016/j.ejrad.2004.11.004 | Rhyme > visual detection | Decision | 10 | -51 | -62 | -5 |
|  |  |  |  |  |  | -51 | -50 | -12 |
|  |  |  |  |  |  | -44 | 34 | -5 |
|  |  |  |  |  |  | -54 | 17 | 18 |
|  |  |  |  |  |  | -24 | -59 | 42 |
|  |  |  |  |  |  | -24 | -56 | 51 |
|  |  |  |  |  |  | -53 | -32 | 22 |
|  |  |  |  |  |  | -46 | 51 | 9 |
|  |  |  |  |  |  | -37 | -71 | 39 |
| Bohland et al. | 2006 | 10.1016/j.neuroimage.2006.04.173 | Complex > simple syllables | Decision | 13 | 0 | 18 | 46 |
|  |  |  |  |  |  | 0 | 4 | 62 |
|  |  |  |  |  |  | 0 | 0 | 70 |
|  |  |  |  |  |  | 4 | 24 | 38 |
|  |  |  |  |  |  | 50 | 22 | -6 |
|  |  |  |  |  |  | 42 | 20 | -12 |
|  |  |  |  |  |  | 38 | 26 | 0 |
|  |  |  |  |  |  | 38 | 24 | -6 |
|  |  |  |  |  |  | -26 | -62 | 52 |
|  |  |  |  |  |  | -30 | -54 | 52 |
|  |  |  |  |  |  | -48 | -40 | 52 |
|  |  |  |  |  |  | -20 | -66 | 66 |
|  |  |  |  |  |  | -38 | -44 | 44 |
|  |  |  |  |  |  | -34 | 26 | 0 |
|  |  |  |  |  |  | -34 | 22 | 4 |
|  |  |  |  |  |  | -50 | 12 | 0 |
|  |  |  |  |  |  | 22 | -76 | -20 |
|  |  |  |  |  |  | 26 | -62 | -18 |
|  |  |  | Syllable x sequence interaction | Decision | 13 | 0 | 16 | 48 |
|  |  |  |  |  |  | -8 | 8 | 62 |
|  |  |  |  |  |  | 2 | 34 | 36 |
|  |  |  |  |  |  | 8 | 26 | 34 |
|  |  |  |  |  |  | 0 | 16 | 66 |
|  |  |  |  |  |  | 2 | 14 | 32 |
|  |  |  |  |  |  | -6 | 24 | 28 |
|  |  |  |  |  |  | 34 | 22 | -8 |
|  |  |  |  |  |  | 38 | 44 | 24 |
|  |  |  |  |  |  | 52 | 20 | -4 |
|  |  |  |  |  |  | 40 | 20 | 10 |
|  |  |  |  |  |  | 52 | 34 | 26 |
|  |  |  |  |  |  | 58 | 24 | 14 |
|  |  |  |  |  |  | -42 | 30 | 24 |
|  |  |  |  |  |  | -30 | 24 | 6 |
|  |  |  |  |  |  | -42 | 46 | 22 |
|  |  |  |  |  |  | -36 | 16 | -8 |
|  |  |  |  |  |  | -58 | 14 | 18 |
|  |  |  |  |  |  | -52 | 16 | 14 |
|  |  |  |  |  |  | -44 | 14 | 4 |
|  |  |  |  |  |  | -62 | 6 | 28 |
|  |  |  |  |  |  | -40 | 12 | 26 |
|  |  |  |  |  |  | -52 | 10 | 44 |
|  |  |  |  |  |  | -50 | 4 | 36 |
|  |  |  |  |  |  | 42 | -50 | -30 |
|  |  |  |  |  |  | 28 | -52 | -24 |
|  |  |  |  |  |  | 32 | -52 | -28 |
|  |  |  |  |  |  | 36 | -56 | -28 |
|  |  |  |  |  |  | -2 | -72 | -8 |
|  |  |  |  |  |  | 14 | -66 | -12 |
|  |  |  |  |  |  | 42 | -72 | -28 |
|  |  |  |  |  |  | 2 | -56 | -32 |
|  |  |  |  |  |  | 14 | -58 | -20 |
|  |  |  |  |  |  | 14 | -54 | -14 |
|  |  |  |  |  |  | 16 | -6 | 14 |
|  |  |  |  |  |  | -10 | 0 | 10 |
|  |  |  |  |  |  | 10 | -2 | 12 |
|  |  |  |  |  |  | -4 | -10 | 14 |
|  |  |  |  |  |  | 8 | -8 | 2 |
|  |  |  |  |  |  | -30 | -52 | 50 |
|  |  |  |  |  |  | -40 | -44 | 54 |
|  |  |  |  |  |  | -52 | -40 | 56 |
|  |  |  |  |  |  | -36 | -48 | 42 |
|  |  |  |  |  |  | -24 | -72 | 46 |
|  |  |  |  |  |  | -18 | -68 | 64 |
|  |  |  |  |  |  | 34 | 2 | 58 |
|  |  |  |  |  |  | 34 | 2 | 38 |
|  |  |  |  |  |  | 34 | 4 | 44 |
|  |  |  |  |  |  | 44 | 12 | 38 |
|  |  |  |  |  |  | 34 | 0 | 48 |
|  |  |  |  |  |  | -44 | -58 | -16 |
|  |  |  |  |  |  | -32 | 0 | 52 |
|  |  |  |  |  |  | -38 | 0 | 62 |
|  |  |  |  |  |  | -38 | -4 | 42 |
| Booth et al. | 2002 | 10.1002/hbm.10054 | Rhyme > meaning judgement | Decision | 13 | -6 | -36 | 45 |
|  |  |  |  |  |  | -57 | -21 | 39 |
|  |  |  |  |  |  | 0 | -39 | 24 |
|  |  |  |  |  |  | -6 | -63 | 24 |
|  |  |  |  |  |  | -9 | 57 | -12 |
|  |  |  | Phonology > visual perception | Decision | 13 | -3 | 21 | 51 |
|  |  |  |  |  |  | 0 | 39 | 39 |
|  |  |  |  |  |  | -27 | -51 | 36 |
|  |  |  |  |  |  | -45 | 30 | 12 |
|  |  |  |  |  |  | 12 | -63 | 12 |
|  |  |  |  |  |  | -18 | -90 | -3 |
|  |  |  |  |  |  | -30 | -6 | -3 |
|  |  |  |  |  |  | 39 | 27 | -9 |
|  |  |  |  |  |  | 30 | -78 | -9 |
|  |  |  |  |  |  | -45 | -60 | -21 |
|  |  |  |  |  |  | 9 | -75 | -30 |
|  |  |  | Phonology > pitch discrimination | Decision | 13 | -6 | 36 | 36 |
|  |  |  |  |  |  | -51 | 21 | 27 |
|  |  |  |  |  |  | 42 | -27 | 12 |
|  |  |  |  |  |  | 60 | -9 | -3 |
|  |  |  |  |  |  | -54 | -18 | -3 |
|  |  |  |  |  |  | -45 | -60 | -18 |
|  |  |  |  |  |  | 15 | -78 | -24 |
|  |  |  |  |  |  | -18 | -66 | 54 |
|  |  |  |  |  |  | -54 | -27 | 51 |
|  |  |  |  |  |  | -24 | -3 | 48 |
| Cousin et al. | 2007 | 10.1016/j.ejrad.2007.01.030 | Rhyme > visual detection | Decision | 11 | -51 | 31 | 2 |
|  |  |  |  |  |  | -12 | 72 | 3 |
|  |  |  |  |  |  | -15 | 45 | 36 |
|  |  |  |  |  |  | 7 | 40 | -18 |
|  |  |  |  |  |  | -50 | -71 | 44 |
| Demonet et al. | 1992 | 10.1093/brain/115.6.1753 | Phonemes > tones | Decision | 9 | -59 | -11 | 2 |
|  |  |  |  |  |  | -57 | -14 | -2 |
|  |  |  |  |  |  | -57 | -14 | -7 |
|  |  |  |  |  |  | -57 | -12 | -11 |
|  |  |  |  |  |  | -55 | -62 | -11 |
|  |  |  |  |  |  | 53 | -2 | -19 |
|  |  |  |  |  |  | -53 | 22 | 17 |
| Desai et al. | 2006 | 10.1162/jocn.2006.18.2.278 | Irregular > regular verbs | Decision | 125 | 9 | 5 | 1 |
|  |  |  |  |  |  | -33 | -54 | 45 |
|  |  |  |  |  |  | -51 | 9 | 21 |
|  |  |  |  |  |  | 36 | -62 | 45 |
|  |  |  |  |  |  | 39 | 27 | -1 |
|  |  |  |  |  |  | 51 | 7 | 31 |
|  |  |  |  |  |  | -30 | -75 | 26 |
|  |  |  |  |  |  | -45 | -56 | -4 |
|  |  |  |  |  |  | -51 | 33 | -1 |
|  |  |  |  |  |  | -47 | -1 | 41 |
|  |  |  |  |  |  | -12 | -27 | 6 |
| Devlin et al. | 2003 | 10.1162/089892903321107837 | Phonology > semantics | Decision | 12 | -50 | 6 | 24 |
|  |  |  |  |  |  | -42 | 0 | 28 |
|  |  |  |  |  |  | -38 | -2 | 44 |
|  |  |  |  |  |  | 44 | 36 | 26 |
|  |  |  |  |  |  | 46 | 42 | 14 |
|  |  |  |  |  |  | 42 | 2 | 28 |
|  |  |  |  |  |  | 46 | 28 | -16 |
|  |  |  |  |  |  | 36 | 24 | -8 |
|  |  |  |  |  |  | -30 | 20 | -8 |
|  |  |  |  |  |  | -42 | 34 | 12 |
|  |  |  |  |  |  | -38 | 34 | 20 |
|  |  |  |  |  |  | -42 | 36 | 30 |
|  |  |  |  |  |  | 6 | 36 | 42 |
|  |  |  |  |  |  | 4 | 16 | 54 |
|  |  |  |  |  |  | 4 | 28 | 46 |
|  |  |  |  |  |  | 24 | 52 | -8 |
|  |  |  |  |  |  | 46 | -44 | 44 |
|  |  |  |  |  |  | 44 | -56 | 52 |
|  |  |  |  |  |  | 52 | -60 | 48 |
|  |  |  |  |  |  | -42 | -40 | 46 |
|  |  |  |  |  |  | -56 | -44 | 46 |
|  |  |  |  |  |  | -52 | -48 | 54 |
|  |  |  |  |  |  | 28 | -68 | 44 |
|  |  |  |  |  |  | 34 | -64 | 36 |
|  |  |  |  |  |  | -20 | -68 | 46 |
|  |  |  |  |  |  | -24 | -72 | 36 |
|  |  |  |  |  |  | -10 | -70 | 50 |
|  |  |  |  |  |  | -48 | -66 | -20 |
|  |  |  |  |  |  | -50 | -54 | -20 |
|  |  |  |  |  |  | -50 | -66 | -10 |
|  |  |  |  |  |  | 52 | -56 | -16 |
|  |  |  |  |  |  | 52 | -56 | -26 |
|  |  |  |  |  |  | 42 | -58 | -10 |
|  |  |  |  |  |  | -30 | -60 | -44 |
| Ghosh et al. | 2008 | 10.1044/1092-4388(2008/07-0119) | Bisyllable > monosyllable words | Decision | 10 | -52 | -6 | 30 |
|  |  |  |  |  |  | -46 | -8 | 54 |
|  |  |  |  |  |  | 50 | -4 | 38 |
|  |  |  |  |  |  | -56 | 16 | 20 |
|  |  |  |  |  |  | -60 | 16 | 32 |
|  |  |  |  |  |  | -6 | 4 | 68 |
|  |  |  |  |  |  | 8 | 16 | 44 |
|  |  |  |  |  |  | 54 | -2 | 32 |
|  |  |  |  |  |  | -24 | -70 | 42 |
|  |  |  |  |  |  | -20 | -70 | 48 |
|  |  |  |  |  |  | 24 | -68 | 50 |
|  |  |  |  |  |  | -32 | -52 | 50 |
|  |  |  |  |  |  | -62 | -2 | -10 |
|  |  |  |  |  |  | -66 | -18 | 2 |
|  |  |  |  |  |  | 70 | -2 | -6 |
|  |  |  |  |  |  | 70 | -28 | 4 |
|  |  |  |  |  |  | -58 | 6 | -12 |
|  |  |  |  |  |  | -42 | -64 | -10 |
|  |  |  |  |  |  | 18 | -92 | -12 |
|  |  |  |  |  |  | -38 | -50 | -24 |
|  |  |  |  |  |  | -40 | -44 | -24 |
|  |  |  |  |  |  | -22 | 4 | 4 |
|  |  |  |  |  |  | 26 | 4 | 0 |
|  |  |  |  |  |  | 22 | 8 | 4 |
|  |  |  |  |  |  | -22 | 0 | -4 |
|  |  |  |  |  |  | -30 | -84 | -20 |
|  |  |  |  |  |  | -8 | -80 | -32 |
|  |  |  |  |  |  | -30 | -60 | -28 |
|  |  |  |  |  |  | -32 | -56 | -26 |
|  |  |  |  |  |  | 28 | -62 | -26 |
|  |  |  |  |  |  | 2 | -102 | -6 |
|  |  |  |  |  |  | 14 | -66 | 10 |
|  |  |  |  |  |  | -46 | -84 | -2 |
|  |  |  |  |  |  | -28 | -82 | 16 |
|  |  |  |  |  |  | -16 | -92 | -10 |
|  |  |  |  |  |  | 34 | -88 | 20 |
|  |  |  |  |  |  | 34 | -80 | -16 |
| Gitelman et al. | 2005 | 10.1016/j.neuroimage.2005.03.014 | Phonology > semantics | Decision | 14 | -60 | 13 | 22 |
|  |  |  |  |  |  | -51 | 6 | 16 |
|  |  |  |  |  |  | 46 | 35 | -22 |
|  |  |  |  |  |  | 56 | 43 | -10 |
|  |  |  |  |  |  | 56 | 48 | -24 |
|  |  |  |  |  |  | -47 | 3 | 50 |
|  |  |  |  |  |  | 46 | 13 | -27 |
|  |  |  |  |  |  | 30 | 9 | -30 |
|  |  |  |  |  |  | 20 | -17 | -27 |
| Gold & Buckner | 2002 | 10.1016/S0896-6273(02)00800-0 | Phonology > semantics | Decision | 24 | -58 | 3 | 27 |
| Gold et al. | 2005 | 10.1093/cercor/bhi024 | Phonology > semantics | Decision | 32 | -43 | -41 | 38 |
|  |  |  |  |  |  | -58 | 3 | 27 |
| Gourovitch et al. | 2000 | 10.1037/0894-4105.14.3.353 | Phonological fluency > semantic fluency | Decision | 18 | 25 | 38 | 23 |
|  |  |  |  |  |  | 46 | 13 | 2 |
|  |  |  |  |  |  | -57 | 5 | 18 |
|  |  |  |  |  |  | -38 | 8 | -5 |
|  |  |  |  |  |  | 48 | 7 | -15 |
| Heim et al. | 2008 | 10.1016/j.neuroimage.2008.01.009 | Phonological > semantic fluency | Decision | 28 | -50 | 10 | 21 |
|  |  |  |  |  |  | -40 | -40 | 47 |
| Katzir et al. | 2005 | 10.1016/j.neuroimage.2005.04.013 | Phonology > visual perception | Decision | 12 | -39 | -69 | -12 |
|  |  |  |  |  |  | -51 | 15 | 33 |
|  |  |  |  |  |  | 33 | -93 | -3 |
|  |  |  |  |  |  | 33 | 30 | -12 |
|  |  |  |  |  |  | -3 | 36 | 45 |
|  |  |  |  |  |  | -6 | -21 | -6 |
|  |  |  |  |  |  | 30 | -33 | 0 |
|  |  |  |  |  |  | -27 | -60 | 42 |
|  |  |  |  |  |  | 12 | 3 | 0 |
|  |  |  |  |  |  | 45 | 24 | 27 |
|  |  |  |  |  |  | 42 | 3 | 27 |
|  |  |  |  |  |  | 33 | -93 | -3 |
|  |  |  |  |  |  | 12 | 18 | 45 |
|  |  |  |  |  |  | -33 | -84 | -15 |
|  |  |  |  |  |  | -45 | 21 | 24 |
|  |  |  |  |  |  | 33 | 21 | 0 |
|  |  |  |  |  |  | -30 | -60 | 45 |
|  |  |  |  |  |  | -36 | -12 | -33 |
|  |  |  |  |  |  | 27 | -33 | -3 |
|  |  |  |  |  |  | 21 | -6 | 15 |
|  |  |  |  |  |  | 0 | -45 | -21 |
|  |  |  |  |  |  | -60 | -45 | -3 |
|  |  |  |  |  |  | 51 | 36 | 30 |
| Kuo et al. | 2004 | 10.1016/j.neuroimage.2003.12.007 | Phonology > orthography | Decision | 10 | -50 | 6 | 47 |
|  |  |  |  |  |  | -3 | 22 | 52 |
|  |  |  |  |  |  | -46 | 15 | 33 |
|  |  |  |  |  |  | -53 | 29 | 23 |
|  |  |  |  |  |  | -48 | 22 | 14 |
|  |  |  |  |  |  | -44 | 10 | 15 |
|  |  |  |  |  |  | -24 | -60 | 51 |
|  |  |  |  |  |  | -39 | -42 | 59 |
|  |  |  |  |  |  | -57 | -42 | 16 |
|  |  |  |  |  |  | -35 | -88 | 9 |
|  |  |  |  |  |  | -44 | -64 | 5 |
|  |  |  |  |  |  | -33 | -82 | 2 |
|  |  |  |  |  |  | -38 | -57 | -7 |
|  |  |  |  |  |  | 10 | 35 | 32 |
|  |  |  |  |  |  | 47 | 19 | 22 |
|  |  |  |  |  |  | 36 | -87 | 10 |
|  |  |  |  |  |  | -48 | 8 | 45 |
|  |  |  |  |  |  | -3 | 22 | 56 |
|  |  |  |  |  |  | -42 | 19 | 28 |
|  |  |  |  |  |  | -51 | 31 | 18 |
|  |  |  |  |  |  | -48 | 21 | 19 |
|  |  |  |  |  |  | -48 | 8 | 20 |
|  |  |  |  |  |  | -51 | 19 | -3 |
|  |  |  |  |  |  | -44 | -52 | 48 |
|  |  |  |  |  |  | -59 | -44 | 19 |
|  |  |  |  |  |  | -35 | -74 | 12 |
|  |  |  |  |  |  | -31 | -82 | 0 |
|  |  |  |  |  |  | 40 | -65 | -10 |
|  |  |  |  |  |  | 31 | -74 | -13 |
| Liu et al. | 2008 | 10.1162/jocn.2009.21141 | Rhyme > visual perception | Decision | 16 | -46 | 32 | 16 |
|  |  |  |  |  |  | -44 | 22 | 8 |
|  |  |  |  |  |  | -30 | -64 | 36 |
|  |  |  |  |  |  | -38 | -44 | -28 |
|  |  |  |  |  |  | -36 | -62 | -28 |
|  |  |  |  |  |  | 36 | 20 | -8 |
|  |  |  |  |  |  | 40 | 20 | 8 |
|  |  |  |  |  |  | 28 | -96 | -4 |
|  |  |  |  |  |  | 8 | -50 | -32 |
|  |  |  |  |  |  | 38 | -58 | -28 |
|  |  |  |  |  |  | 18 | -30 | 20 |
|  |  |  |  |  |  | 18 | 6 | 20 |
|  |  |  | Rhyme > tone discrimination | Decision | 16 | -46 | 32 | 20 |
|  |  |  |  |  |  | -46 | 10 | 20 |
|  |  |  |  |  |  | -52 | -2 | 12 |
|  |  |  |  |  |  | -62 | -24 | 4 |
|  |  |  |  |  |  | -48 | 4 | 44 |
|  |  |  |  |  |  | 2 | 20 | 52 |
|  |  |  |  |  |  | 42 | 14 | -28 |
|  |  |  |  |  |  | 0 | -78 | -20 |
| Lurito et al. | 2000 | 10.1002/1097-0193(200007)10:3<99::AID-HBM10>3.0.CO;2-Q | Phonological fluency > orthography | Decision | 5 | -61 | -56 | 9 |
|  |  |  |  |  |  | -59 | -32 | 9 |
|  |  |  |  |  |  | -49 | -55 | -20 |
|  |  |  |  |  |  | -45 | -59 | 52 |
|  |  |  |  |  |  | -34 | -59 | 44 |
|  |  |  |  |  |  | -49 | -45 | 48 |
|  |  |  |  |  |  | -4 | 16 | 45 |
|  |  |  |  |  |  | -4 | -1 | 61 |
|  |  |  |  |  |  | -56 | 17 | 1 |
|  |  |  |  |  |  | -28 | -62 | -25 |
|  |  |  |  |  |  | -48 | 38 | 21 |
|  |  |  |  |  |  | -52 | 16 | 31 |
|  |  |  |  |  |  | -58 | 16 | 3 |
|  |  |  |  |  |  | -67 | -29 | 29 |
|  |  |  |  |  |  | -60 | -48 | 2 |
|  |  |  |  |  |  | -47 | -63 | -18 |
|  |  |  |  |  |  | 51 | -47 | 18 |
|  |  |  |  |  |  | 44 | 25 | 5 |
|  |  |  |  |  |  | 39 | 40 | 37 |
|  |  |  |  |  |  | 37 | -64 | -30 |
|  |  |  |  |  |  | 31 | -68 | -59 |
|  |  |  |  |  |  | 61 | 10 | 0 |
| McDermott et al. | 2003 | 10.1016/S0028-3932(02)00162-8 | Rhyme > semantic judgements | Decision | 20 | -58 | 6 | 13 |
|  |  |  |  |  |  | 48 | -35 | 49 |
|  |  |  |  |  |  | -32 | -54 | 55 |
|  |  |  |  |  |  | 35 | -44 | 56 |
|  |  |  |  |  |  | 48 | -63 | -6 |
| Mummery et al. | 1998 | 10.1162/089892998563059 | Syllable number > semantic judgements | Decision | 10 | -59 | -31 | 38 |
|  |  |  |  |  |  | -55 | -3 | 39 |
|  |  |  |  |  |  | 58 | -36 | 50 |
|  |  |  |  |  |  | 61 | -54 | -25 |
| Paulesu et al. | 1997 | 10.1097/00001756-199705260-00042 | Phonological fluency > semantics | Decision | 6 | -38 | 10 | 18 |
| Price et al. | 1997 | 10.1162/jocn.1997.9.6.727 | Syllable number > semantic judgements | Decision | 6 | -42 | -44 | 36 |
|  |  |  |  |  |  | 38 | -48 | 40 |
|  |  |  |  |  |  | 45 | -30 | 42 |
|  |  |  |  |  |  | 32 | -61 | 37 |
|  |  |  |  |  |  | -55 | 2 | 23 |
|  |  |  |  |  |  | -13 | -82 | 26 |
| Roskies et al. | 2001 | 10.1162/08989290152541485 | Rhyme > semantic judgements | Decision | 20 | -26 | 28 | -2 |
|  |  |  |  |  |  | -2 | -8 | 3 |
|  |  |  |  |  |  | -37 | -76 | -19 |
|  |  |  |  |  |  | -39 | -1 | 5 |
|  |  |  |  |  |  | -52 | 3 | 25 |
|  |  |  |  |  |  | -52 | 6 | 14 |
|  |  |  |  |  |  | -58 | -7 | 40 |
|  |  |  |  |  |  | 18 | -6 | 14 |
|  |  |  |  |  |  | 32 | -72 | -20 |
|  |  |  |  |  |  | 34 | -68 | -31 |
|  |  |  |  |  |  | 7 | -70 | 4 |
| Seghier et al. | 2004 | 10.1002/hbm.20053 | Rhyme > visual perception | Decision | 26 | -50 | 19 | 12 |
|  |  |  |  |  |  | -46 | 37 | 23 |
|  |  |  |  |  |  | -47 | 17 | 31 |
|  |  |  |  |  |  | -57 | -35 | 12 |
|  |  |  |  |  |  | -58 | -36 | 3 |
|  |  |  |  |  |  | -39 | -47 | 42 |
|  |  |  |  |  |  | -22 | -71 | 52 |
|  |  |  |  |  |  | -48 | 2 | 50 |
|  |  |  |  |  |  | 0 | 13 | 55 |
| Simon et al. | 2002 | 10.1016/S0896-6273(02)00575-5 | Phonology > orthography | Decision | 10 | -3 | 20 | 56 |
|  |  |  |  |  |  | -59 | 26 | 30 |
|  |  |  |  |  |  | -46 | 20 | 8 |
|  |  |  |  |  |  | 57 | 20 | 1 |
|  |  |  |  |  |  | -54 | 6 | 50 |
|  |  |  |  |  |  | -67 | -33 | 40 |
|  |  |  |  |  |  | -28 | -80 | 49 |
| Simon et al. | 2004 | 10.1016/j.neuroimage.2004.09.023 | Letter detection & calculation > orthography | Decision | 10 | -51 | 23 | -6 |
|  |  |  |  |  |  | -61 | 10 | 18 |
|  |  |  |  |  |  | -44 | 31 | 16 |
|  |  |  |  |  |  | -53 | 23 | 25 |
|  |  |  |  |  |  | -48 | 15 | 44 |
|  |  |  |  |  |  | 0 | -22 | 47 |
|  |  |  |  |  |  | -28 | -71 | 52 |
|  |  |  |  |  |  | 55 | 24 | -3 |
|  |  |  |  |  |  | 8 | 37 | 45 |
| Specht et al. | 2003 | 10.1016/S0304-3940(03)00494-4 | Pseudowords > tones/nonwords | Decision | 15 | -64 | -24 | 4 |
|  |  |  |  |  |  | 60 | -12 | 0 |
|  |  |  |  |  |  | 4 | 40 | 40 |
|  |  |  |  |  |  | 36 | 24 | -8 |
|  |  |  |  |  |  | -28 | 20 | -4 |
|  |  |  |  |  |  | -8 | 36 | 28 |
| Tyler et al. | 2005 | 10.1016/j.neuropsychologia.2005.03.008 | Regular > irregular words | Decision | 18 | -46 | -26 | 8 |
|  |  |  |  |  |  | -56 | -24 | 10 |
|  |  |  |  |  |  | -56 | -48 | 8 |
|  |  |  |  |  |  | 58 | -14 | 2 |
|  |  |  |  |  |  | 58 | -28 | 4 |
|  |  |  |  |  |  | 62 | -6 | 10 |
|  |  |  |  |  |  | 46 | -42 | 34 |
|  |  |  |  |  |  | 38 | -44 | 42 |
|  |  |  |  |  |  | -50 | 12 | 20 |
| Wildgruber et al. | 2005 | 10.1016/j.neuroimage.2004.10.034 | Vowel > emotion identification | Decision | 10 | -54 | 6 | 33 |
|  |  |  |  |  |  | -48 | -39 | 54 |
|  |  |  |  |  |  | 27 | -66 | 39 |
| Xu et al. | 2001 | 10.1093/cercor/11.3.267 | Rhyme > visual perception | Decision | 12 | 32 | -68 | -20 |
|  |  |  |  |  |  | -48 | -54 | -14 |
|  |  |  |  |  |  | -44 | 4 | 28 |
|  |  |  |  |  |  | -34 | -50 | 38 |
|  |  |  | Pseudoword > word rhyming | Decision | 12 | 10 | -72 | -46 |
|  |  |  |  |  |  | -46 | -66 | -10 |
|  |  |  |  |  |  | 32 | -76 | -14 |
|  |  |  |  |  |  | -52 | 10 | 12 |
|  |  |  |  |  |  | -50 | -4 | 46 |
|  |  |  |  |  |  | -38 | -48 | 50 |
| Xu et al. | 2002 | 10.1006/nimg.2002.1215 | Rhyme > orthography | Decision | 6 | -40 | 53 | 18 |
|  |  |  |  |  |  | -2 | 12 | 64 |
|  |  |  |  |  |  | -57 | 34 | 20 |
|  |  |  |  |  |  | -40 | 26 | 11 |
|  |  |  |  |  |  | -55 | 18 | 17 |
|  |  |  |  |  |  | -55 | 10 | 45 |
|  |  |  |  |  |  | -55 | -43 | 28 |
|  |  |  |  |  |  | 0 | 18 | 59 |
|  |  |  |  |  |  | -57 | 36 | 20 |
|  |  |  |  |  |  | -57 | 18 | 17 |
|  |  |  |  |  |  | -44 | 3 | 56 |
|  |  |  |  |  |  | -50 | 12 | 42 |
|  |  |  |  |  |  | -60 | 13 | -12 |
|  |  |  |  |  |  | -59 | -43 | 32 |
|  |  |  |  |  |  | -18 | 58 | 6 |
|  |  |  |  |  |  | 36 | 53 | 14 |
|  |  |  |  |  |  | -48 | 24 | 28 |
|  |  |  |  |  |  | -40 | 31 | -9 |
|  |  |  |  |  |  | -33 | -69 | 45 |
|  |  |  |  |  |  | -46 | -38 | 34 |
|  |  |  |  |  |  | 46 | 59 | 11 |
|  |  |  |  |  |  | 6 | 45 | 49 |
|  |  |  |  |  |  | -48 | 17 | 26 |
|  |  |  |  |  |  | -35 | -62 | 54 |
| Braun et al. | 1997 | 10.1093/brain/120.5.761 | Auto/paced speech > non-lexical speech | Decision | 20 | -20 | 37 | 37 |
|  |  |  |  |  |  | -13 | 23 | 47 |
|  |  |  |  |  |  | -26 | 21 | 48 |
|  |  |  |  |  |  | -13 | -90 | 31 |
|  |  |  |  |  |  | -57 | -33 | -5 |
|  |  |  |  |  |  | -42 | -69 | 30 |
|  |  |  |  |  |  | -14 | 35 | 33 |
| Heim et al. | 2002 | 10.1016/S0304-3940(02)00494-9 | Phoneme production > rest | Passive | 12 | -41 | 4 | 2 |
|  |  |  |  |  |  | -54 | 3 | 3 |
|  |  |  |  |  |  | -38 | 13 | 34 |
|  |  |  |  |  |  | -58 | -15 | 6 |
|  |  |  |  |  |  | -44 | -8 | 36 |
|  |  |  |  |  |  | -41 | -31 | 16 |
| McGuire et al. | 1996 | 10.1017/S0033291700033699 | Inner speech > reading | Passive | 6 | -33 | 24 | 12 |
| Price et al. | 1996 | 10.1093/brain/119.3.919 | Reading aloud > listening | Passive |  | -48 | -8 | 33 |
|  |  |  |  |  |  | -59 | -3 | 24 |
|  |  |  |  |  |  | -59 | 7 | 9 |
|  |  |  |  |  |  | -38 | 8 | 0 |
|  |  |  |  |  |  | -27 | -11 | 6 |
|  |  |  |  |  |  | 8 | -66 | -12 |
|  |  |  |  |  |  | 1 | -9 | 1 |
|  |  |  |  |  |  | -2 | 8 | 62 |
|  |  |  |  |  |  | 53 | -7 | 36 |
|  |  |  | Repetition > rest | Passive | 6 | -57 | -44 | 14 |
|  |  |  |  |  |  | -55 | -36 | 13 |
|  |  |  |  |  |  | -59 | -15 | 7 |
|  |  |  |  |  |  | -59 | -15 | 7 |
|  |  |  |  |  |  | -64 | -2 | 6 |
|  |  |  |  |  |  | -59 | -1 | -3 |
|  |  |  |  |  |  | -50 | -7 | 24 |
|  |  |  |  |  |  | -48 | -8 | 33 |
|  |  |  |  |  |  | -33 | 6 | 0 |
|  |  |  |  |  |  | -23 | -21 | 7 |
|  |  |  |  |  |  | -23 | -17 | 6 |
|  |  |  |  |  |  | 51 | -38 | 7 |
|  |  |  |  |  |  | 55 | -42 | -15 |
|  |  |  |  |  |  | 62 | -22 | -3 |
|  |  |  |  |  |  | 62 | -22 | -3 |
|  |  |  |  |  |  | 62 | -6 | -1 |
|  |  |  |  |  |  | 62 | -6 | -1 |
|  |  |  |  |  |  | 32 | -10 | 32 |
|  |  |  |  |  |  | 29 | 1 | 13 |
|  |  |  |  |  |  | 47 | -12 | 5 |
|  |  |  |  |  |  | 31 | 17 | 2 |
|  |  |  |  |  |  | 5 | -61 | -8 |
|  |  |  |  |  |  | -1 | -55 | -8 |
|  |  |  |  |  |  | -14 | -2 | 9 |
|  |  |  |  |  |  | -8 | -24 | -2 |
|  |  |  |  |  |  | -5 | -17 | 2 |
|  |  |  |  |  |  | -5 | 3 | 62 |
|  |  |  | Reading aloud > rest | Passive | 6 | -61 | -45 | 10 |
|  |  |  |  |  |  | -44 | -8 | 33 |
|  |  |  |  |  |  | -57 | -3 | 24 |
|  |  |  |  |  |  | -62 | 2 | 5 |
|  |  |  |  |  |  | -23 | -21 | 7 |
|  |  |  |  |  |  | 8 | -65 | -7 |
|  |  |  |  |  |  | 19 | -25 | 11 |
|  |  |  |  |  |  | -1 | -15 | 1 |
|  |  |  |  |  |  | 0 | 5 | 58 |
|  |  |  |  |  |  | 55 | -53 | 9 |
|  |  |  |  |  |  | 51 | -38 | 7 |
|  |  |  |  |  |  | 49 | -7 | 36 |
|  |  |  | Repetition > listening | Passive | 6 | -66 | -45 | 10 |
|  |  |  |  |  |  | -55 | -38 | 14 |
|  |  |  |  |  |  | -64 | -26 | 8 |
|  |  |  |  |  |  | -64 | -10 | 11 |
|  |  |  |  |  |  | -57 | -5 | 24 |
|  |  |  |  |  |  | -48 | -8 | 33 |
|  |  |  |  |  |  | -46 | -9 | 24 |
|  |  |  |  |  |  | -59 | -3 | 19 |
|  |  |  |  |  |  | -31 | -10 | 15 |
|  |  |  |  |  |  | -27 | -2 | 1 |
|  |  |  |  |  |  | -33 | 6 | 0 |
|  |  |  |  |  |  | -61 | 11 | 9 |
|  |  |  |  |  |  | -61 | 11 | 9 |
|  |  |  |  |  |  | -25 | -9 | 1 |
|  |  |  |  |  |  | -18 | -15 | 6 |
|  |  |  |  |  |  | 8 | -61 | -12 |
|  |  |  |  |  |  | 5 | -61 | -12 |
|  |  |  |  |  |  | -2 | 3 | 62 |
|  |  |  |  |  |  | -5 | 3 | 62 |
|  |  |  |  |  |  | 55 | -40 | 12 |
|  |  |  |  |  |  | 66 | -21 | 5 |
|  |  |  |  |  |  | 51 | 5 | 3 |
|  |  |  |  |  |  | 53 | -7 | 36 |
|  |  |  |  |  |  | 51 | -8 | 31 |
|  |  |  |  |  |  | 64 | -7 | 13 |
|  |  |  |  |  |  | 62 | -6 | 8 |
|  |  |  |  |  |  | 32 | 3 | 12 |
|  |  |  |  |  |  | 40 | -5 | -5 |
|  |  |  |  |  |  | 29 | -3 | 13 |
| Riecker et al. | 2000 | 10.1006/brln.2000.2356 | Phoneme production > rest | Passive | 10 | 66 | 4 | 21 |
|  |  |  |  |  |  | -57 | -2 | 27 |
|  |  |  |  |  |  | 56 | -3 | 18 |
|  |  |  |  |  |  | -60 | 1 | 27 |
|  |  |  |  |  |  | 44 | -11 | 33 |
|  |  |  | Word production > rest | Passive | 10 | 57 | -9 | 21 |
|  |  |  |  |  |  | -44 | -11 | 34 |
| Warburton et al. | 1996 | 10.1093/brain/119.1.159 | Covert repetition > rest | Passive | 6 | -48 | -34 | 5 |
|  |  |  |  |  |  | -51 | -44 | -9 |
|  |  |  |  |  |  | -51 | -2 | 2 |
|  |  |  |  |  |  | -44 | 7 | -7 |
|  |  |  |  |  |  | -40 | 23 | -4 |
|  |  |  |  |  |  | -51 | 18 | 12 |
|  |  |  |  |  |  | 62 | -32 | 0 |
|  |  |  |  |  |  | 57 | -5 | -5 |
|  |  |  |  |  |  | 38 | 7 | 8 |
|  |  |  | Covert repetition > verb generation | Passive | 6 | -46 | -26 | 4 |
|  |  |  |  |  |  | 60 | -43 | 1 |
|  |  |  |  |  |  | 44 | 49 | -8 |
| Wildgruber et al. | 2001 | 10.1006/nimg.2000.0672 | Covert phoneme production > rest | Passive | 10 | -50 | 2 | 43 |
|  |  |  |  |  |  | 60 | 8 | 31 |
|  |  |  |  |  |  | -28 | -2 | -11 |
|  |  |  |  |  |  | -18 | -68 | -24 |
|  |  |  |  |  |  | 20 | -71 | -21 |
|  |  |  |  |  |  | -60 | -18 | -2 |
|  |  |  |  |  |  | 69 | -15 | -8 |
| Calvert et al. | 1999 | 10.1097/00001756-199908200-00033 | Covert repetition > rest | Passive | 5 | -45 | -72 | 7 |
|  |  |  |  |  |  | -48 | -71 | 12 |
|  |  |  |  |  |  | 54 | -65 | 4 |
|  |  |  |  |  |  | 54 | -57 | 9 |
|  |  |  |  |  |  | 63 | -20 | 11 |
|  |  |  |  |  |  | -48 | -24 | 13 |
|  |  |  |  |  |  | 32 | -5 | -1 |
| Beauregard et al. | 1997 | 10.1162/jocn.1997.9.4.441 | Viewing letters > rest | Passive | 10 | -29 | -91 | -11 |
|  |  |  |  |  |  | -49 | -81 | -13 |
|  |  |  |  |  |  | -45 | -81 | -9 |
|  |  |  |  |  |  | 4 | 51 | -37 |
|  |  |  |  |  |  | 27 | -95 | -5 |
|  |  |  |  |  |  | 23 | -101 | 12 |
|  |  |  |  |  |  | -17 | -100 | 3 |
|  |  |  |  |  |  | -34 | 35 | -13 |
|  |  |  |  |  |  | -37 | 17 | -28 |
|  |  |  |  |  |  | -7 | -93 | -28 |
|  |  |  |  |  |  | -32 | -104 | 22 |
|  |  |  |  |  |  | -62 | -38 | -3 |
|  |  |  |  |  |  | -2 | 35 | 41 |
|  |  |  |  |  |  | -57 | -51 | -18 |
|  |  |  |  |  |  | -16 | 32 | -26 |
|  |  |  |  |  |  | -22 | 28 | -27 |
|  |  |  |  |  |  | -67 | -53 | -18 |
|  |  |  |  |  |  | -63 | -22 | -15 |
| Jancke & Shah | 2002 | 10.1212/WNL.58.5.736 | Dichotic > binaural listening of syllables | Passive | 10 | 60 | -8 | 4 |
|  |  |  |  |  |  | 64 | -32 | -4 |
|  |  |  |  |  |  | 56 | 24 | -16 |
|  |  |  |  |  |  | 40 | 24 | -12 |
|  |  |  |  |  |  | 44 | 16 | -12 |
|  |  |  |  |  |  | -28 | 60 | 12 |
|  |  |  |  |  |  | -28 | 60 | -4 |
|  |  |  |  |  |  | -44 | 24 | -8 |
|  |  |  |  |  |  | 4 | -68 | -8 |
| Jessen et al. | 1999 | 10.1016/S0304-3940(99)00453-X | Letter strings > words | Decision | 12 | -67 | -22 | 19 |
|  |  |  |  |  |  | -51 | 16 | 12 |
|  |  |  |  |  |  | -21 | -58 | 52 |
|  |  |  |  |  |  | -34 | -79 | -3 |
|  |  |  |  |  |  | -50 | 9 | 46 |
|  |  |  |  |  |  | 2 | 11 | 58 |
| Joanisse & Gati | 2003 | 10.1016/S1053-8119(03)00046-6 | Syllables > non-speech tones | Decision | 7 | -59 | -22 | 2 |
|  |  |  |  |  |  | -54 | -14 | -7 |
|  |  |  |  |  |  | 60 | -46 | 9 |
|  |  |  |  |  |  | 57 | -25 | -1 |
|  |  |  |  |  |  | 68 | 2 | -19 |
|  |  |  |  |  |  | 34 | -43 | -18 |
|  |  |  |  |  |  | -15 | 48 | 8 |
|  |  |  |  |  |  | 23 | -69 | -8 |
|  |  |  |  |  |  | 36 | -75 | -9 |
| Paulesu et al. | 2000 | 10.1038/71163 | Nonwords > words | Passive | 78 | -44 | 6 | 45 |
|  |  |  |  |  |  | -44 | 28 | 9 |
|  |  |  |  |  |  | -74 | -31 | 0 |
|  |  |  |  |  |  | -51 | -61 | -4 |
|  |  |  |  |  |  | -55 | -64 | -13 |
|  |  |  |  |  |  | -51 | -72 | -3 |
|  |  |  |  |  |  | -57 | -56 | -20 |
|  |  |  |  |  |  | -49 | -73 | -14 |
|  |  |  |  |  |  | -51 | -47 | -15 |
|  |  |  | Reading > rest | Passive | 78 | -46 | 8 | 18 |
|  |  |  |  |  |  | -40 | 26 | 7 |
|  |  |  |  |  |  | -50 | -4 | 33 |
|  |  |  |  |  |  | -23 | 1 | 14 |
|  |  |  |  |  |  | -40 | -35 | 24 |
|  |  |  |  |  |  | -68 | -44 | 14 |
|  |  |  |  |  |  | -63 | -51 | 11 |
|  |  |  |  |  |  | -53 | -63 | -22 |
|  |  |  |  |  |  | -42 | -56 | -25 |
|  |  |  |  |  |  | -18 | -7 | 21 |
|  |  |  |  |  |  | -7 | -30 | -1 |
| Peoppel et al. | 2004 | 10.1016/j.neuropsychologia.2003.07.010 | Categorising phonemes > discriminating tones | Decision | 10 | -62 | -18 | -29 |
|  |  |  |  |  |  | -64 | -18 | -6 |
|  |  |  |  |  |  | -66 | -37 | 5 |
|  |  |  |  |  |  | -51 | -48 | -3 |
|  |  |  |  |  |  | -46 | -73 | 30 |
|  |  |  |  |  |  | -6 | 45 | -22 |
|  |  |  |  |  |  | -31 | 23 | 43 |
|  |  |  |  |  |  | -45 | 28 | -16 |
|  |  |  |  |  |  | -7 | 42 | 36 |
|  |  |  |  |  |  | -9 | -38 | 31 |
|  |  |  |  |  |  | -27 | -13 | -25 |
|  |  |  | Categorising phonemes > lexical decisions | Decision | 10 | -4 | 49 | -27 |
|  |  |  |  |  |  | -12 | 59 | -6 |
|  |  |  |  |  |  | -32 | 60 | -15 |
|  |  |  |  |  |  | 36 | -11 | 18 |
|  |  |  |  |  |  | -12 | 45 | 0 |
|  |  |  |  |  |  | 49 | -74 | 11 |
|  |  |  |  |  |  | 47 | -49 | 31 |
|  |  |  |  |  |  | 3 | 66 | -7 |
|  |  |  |  |  |  | 34 | 31 | 37 |
|  |  |  |  |  |  | 7 | 25 | -7 |
|  |  |  |  |  |  | 3 | 42 | -9 |
|  |  |  |  |  |  | 18 | -77 | -15 |
| Poeppel et al. | 2004 | 10.1016/j.neuropsychologia.2003.07.010 | Categorising phonemes > rest | Decision | 10 | -40 | -32 | 8 |
|  |  |  |  |  |  | -57 | -3 | -3 |
|  |  |  |  |  |  | -61 | -24 | 3 |
|  |  |  |  |  |  | -51 | -36 | 9 |
|  |  |  |  |  |  | -53 | 29 | 16 |
|  |  |  |  |  |  | -50 | -1 | 41 |
|  |  |  |  |  |  | -3 | 11 | 48 |
|  |  |  |  |  |  | -48 | 6 | 45 |
|  |  |  |  |  |  | -9 | 16 | 34 |
|  |  |  |  |  |  | -31 | -28 | -1 |
|  |  |  |  |  |  | -5 | -84 | -19 |
|  |  |  |  |  |  | -16 | -7 | 19 |
|  |  |  |  |  |  | -5 | 10 | -1 |
|  |  |  |  |  |  | -12 | -2 | 5 |
|  |  |  |  |  |  | -14 | -9 | -3 |
|  |  |  |  |  |  | 53 | -29 | 6 |
|  |  |  |  |  |  | 57 | -3 | -10 |
|  |  |  |  |  |  | 64 | -28 | 2 |
|  |  |  |  |  |  | 60 | -44 | 8 |
|  |  |  |  |  |  | 40 | -52 | -5 |
|  |  |  |  |  |  | 34 | 30 | 19 |
|  |  |  |  |  |  | 40 | 25 | 19 |
|  |  |  |  |  |  | 40 | 21 | -7 |
|  |  |  |  |  |  | 34 | -37 | -2 |
| Sekiyama et al. | 2003 | 10.1016/S0168-0102(03)00214-1 | Phoneme identification > face identification | Decision | 8 | -58 | -20 | -1 |
|  |  |  |  |  |  | -43 | 30 | 8 |
|  |  |  |  |  |  | -45 | 7 | 37 |
|  |  |  |  |  |  | 50 | 31 | 11 |
|  |  |  |  |  |  | 52 | -24 | -2 |
|  |  |  |  |  |  | -58 | -22 | 0 |
|  |  |  |  |  |  | -50 | 20 | 9 |
|  |  |  |  |  |  | 57 | -9 | -25 |
|  |  |  |  |  |  | -48 | -29 | -10 |
|  |  |  |  |  |  | 63 | -11 | -5 |
| Zatorre et al. | 1992 | 10.1126/science.256.5058.846 | Phonemes > speech | Decision | 10 | -51 | 7 | 22 |
|  |  |  |  |  |  | 9 | -29 | 22 |
|  |  |  |  |  |  | 17 | -99 | -7 |
|  |  |  |  |  |  | -29 | -54 | 52 |
|  |  |  |  |  |  | 0 | 18 | 24 |
|  |  |  |  |  |  | -66 | -38 | -20 |
| Belin & Zatorre | 2003 | 10.1097/00001756-200311140-00019 | Syllable > speaker adaptation (passive listening) | Passive | 14 | 64 | 3 | -15 |
|  |  |  | Syllable listening > rest | Passive |  | -42 | -31 | 17 |
|  |  |  |  |  |  | 53 | -16 | 7 |
| Hugdahl et al. | 2003 | 10.1016/S0093-934X(02)00500-X | Attend vowels > rest | Decision | 13 | -64 | -12 | -7 |
|  |  |  |  |  |  | 66 | -20 | -4 |
|  |  |  |  |  |  | -55 | -65 | -20 |
|  |  |  |  |  |  | -59 | -67 | 3 |
|  |  |  | Attend pseudowords > rest |  | 13 | 66 | -20 | -4 |
|  |  |  |  |  |  | -68 | -17 | -11 |
|  |  |  |  |  |  | -59 | -27 | 13 |
| Jancke et al. | 2002 | 10.1006/nimg.2001.1027 | Syllables > tones (passive listening) | Passive | 6 | -64 | -12 | -8 |
|  |  |  |  |  |  | -56 | -16 | -12 |
|  |  |  |  |  |  | -60 | -24 | 8 |
|  |  |  |  |  |  | 44 | -24 | 8 |
|  |  |  |  |  |  | 52 | -28 | 8 |
|  |  |  | Syllables > vowels (passive listening) | Passive | 6 | 60 | -8 | -4 |
|  |  |  |  |  |  | 56 | -28 | 4 |
|  |  |  |  |  |  | 44 | -28 | 12 |
|  |  |  |  |  |  | -48 | -32 | 12 |
|  |  |  |  |  |  | -40 | -36 | 16 |
|  |  |  |  |  |  | -56 | -24 | 8 |
|  |  |  | Voiceless syllables > voiced syllables (passive listening) | Passive | 6 | -44 | -28 | 12 |
|  |  |  |  |  |  | -44 | -24 | 0 |
| Herbster et al. | 1997 | 10.1002/(SICI)1097-0193(1997)5:2<84::AID-HBM2>3.0.CO;2-I | Irregular words > say “hiya” to letter strings | Passive | 10 | -40 | -44 | -26 |
|  |  |  |  |  |  | -64 | -18 | -2 |
|  |  |  |  |  |  | -42 | 14 | -10 |
|  |  |  |  |  |  | -12 | 1 | -5 |
|  |  |  |  |  |  | -8 | -68 | -11 |
|  |  |  |  |  |  | -16 | -36 | -14 |
|  |  |  |  |  |  | -64 | -22 | 3 |
|  |  |  |  |  |  | -32 | 18 | -33 |
|  |  |  |  |  |  | -51 | 8 | -4 |
|  |  |  |  |  |  | -5 | 10 | 39 |
|  |  |  |  |  |  | -33 | -96 | -12 |
|  |  |  |  |  |  | -27 | -59 | -30 |
|  |  |  |  |  |  | 46 | -77 | -25 |
|  |  |  |  |  |  | 18 | -62 | -21 |
| Kotz et al. | 2002 | 10.1006/nimg.2002.1316 | Pseudowords > words | Decision | 13 | -35 | 9 | 28 |
|  |  |  |  |  |  | -59 | 3 | -4 |
|  |  |  |  |  |  | -59 | -21 | 9 |
|  |  |  |  |  |  | 32 | 11 | 23 |
|  |  |  |  |  |  | 28 | 29 | 0 |
|  |  |  |  |  |  | 44 | 31 | 13 |
| Mechelli et al. | 2000 | 10.1162/089892900564000 | Pseudowords > words (silent reading) | Passive | 6 | -35 | -98 | -6 |
|  |  |  |  |  |  | 51 | -72 | -14 |
|  |  |  |  |  |  | -26 | -63 | 63 |
|  |  |  |  |  |  | -62 | -14 | -11 |
|  |  |  |  |  |  | -52 | 0 | 48 |
|  |  |  |  |  |  | -46 | 6 | 27 |
|  |  |  |  |  |  | -42 | 35 | -16 |
| Meyer et al. | 2002 | 10.1002/hbm.10042 | Pseudospeech > normal speech (tense judgement) | Decision | 14 | -36 | 30 | -1 |
|  |  |  |  |  |  | -53 | -5 | -1 |
|  |  |  |  |  |  | 41 | 16 | 19 |
|  |  |  |  |  |  | 40 | 36 | 8 |
|  |  |  |  |  |  | 47 | 10 | -9 |
|  |  |  |  |  |  | 49 | -24 | 9 |
|  |  |  | Pseudospeech > prosodic speech (tense judgement) | Decision | 14 | -52 | 31 | 15 |
|  |  |  |  |  |  | -47 | -19 | 5 |
|  |  |  |  |  |  | -27 | -6 | -5 |
|  |  |  |  |  |  | -5 | -13 | 6 |
|  |  |  |  |  |  | -14 | -29 | 2 |
|  |  |  |  |  |  | 53 | -9 | -1 |
|  |  |  |  |  |  | 13 | -7 | -9 |
| Poldrack et al. | 1999 | 10.1006/nimg.1999.0441 | Syllable number > orthography | Decision | 8 | 36 | 41 | 15 |
|  |  |  |  |  |  | -2 | 14 | 34 |
|  |  |  |  |  |  | 42 | 26 | 29 |
|  |  |  |  |  |  | -50 | 33 | 11 |
|  |  |  |  |  |  | 7 | 26 | 26 |
|  |  |  |  |  |  | 66 | 13 | 23 |
|  |  |  |  |  |  | -48 | 4 | 23 |
|  |  |  | Syllable number > semantics | Decision | 8 | -59 | 12 | 21 |
|  |  |  |  |  |  | 43 | 33 | 9 |
|  |  |  |  |  |  | 2 | 8 | 43 |
|  |  |  |  |  |  | 2 | 27 | 40 |
|  |  |  |  |  |  | 47 | 4 | 22 |
|  |  |  |  |  |  | -51 | 25 | 23 |
|  |  |  |  |  |  | 48 | 45 | 4 |
|  |  |  |  |  |  | 53 | 21 | 19 |
|  |  |  |  |  |  | -53 | 40 | 6 |
|  |  |  |  |  |  | 14 | 3 | 11 |
|  |  |  |  |  |  | -17 | 11 | 3 |
|  |  |  |  |  |  | 2 | 43 | 43 |
|  |  |  |  |  |  | 45 | 61 | -6 |
|  |  |  |  |  |  | -17 | 31 | -5 |
|  |  |  |  |  |  | -44 | 41 | 21 |
|  |  |  |  |  |  | 54 | 3 | 12 |
|  |  |  |  |  |  | 17 | 39 | -5 |
|  |  |  |  |  |  | 37 | 27 | 38 |
|  |  |  |  |  |  | -50 | 3 | 11 |
|  |  |  |  |  |  | 53 | 40 | -2 |
|  |  |  |  |  |  | 52 | 14 | 35 |
|  |  |  |  |  |  | 0 | 52 | -39 |
|  |  |  |  |  |  | 12 | 55 | -6 |
|  |  |  |  |  |  | 45 | 55 | -9 |
|  |  |  |  |  |  | 30 | 66 | -6 |
|  |  |  |  |  |  | 1 | 6 | 47 |
| Binder et al. | 2000 | 10.1093/cercor/10.5.512 | Pseudowords > tones | Decision | 28 | -56 | -44 | 7 |
|  |  |  |  |  |  | -60 | -33 | 4 |
|  |  |  |  |  |  | -57 | -26 | -1 |
|  |  |  |  |  |  | -57 | -13 | -2 |
|  |  |  |  |  |  | -59 | -5 | -4 |
|  |  |  |  |  |  | 62 | -30 | 2 |
|  |  |  | Reversed syllables > tones | Decision | 28 | -65 | -33 | 4 |
|  |  |  |  |  |  | -59 | -24 | 0 |
|  |  |  |  |  |  | -56 | -12 | -2 |
|  |  |  |  |  |  | -59 | -5 | -4 |
|  |  |  |  |  |  | 64 | -30 | 2 |
|  |  |  |  |  |  | 65 | -11 | -2 |
|  |  |  |  |  |  | 66 | -4 | -11 |
| Cappa et al. | 1998 | 10.1006/nimg.1998.0368 | Pseudowords (detect phonemes) > rest | Decision | 13 | -20 | -102 | -8 |
|  |  |  |  |  |  | -16 | -104 | -3 |
|  |  |  |  |  |  | -35 | -27 | 57 |
|  |  |  |  |  |  | -49 | -56 | -21 |
|  |  |  |  |  |  | -22 | -78 | 17 |
|  |  |  |  |  |  | 4 | -77 | 3 |
|  |  |  |  |  |  | 19 | -104 | -4 |
|  |  |  |  |  |  | 51 | -56 | -22 |
|  |  |  |  |  |  | 32 | -69 | 46 |
|  |  |  |  |  |  | 36 | -58 | 50 |
|  |  |  |  |  |  | 5 | -67 | -29 |
|  |  |  |  |  |  | 23 | -56 | -22 |
| Fiez et al. | 1996 | 10.1093/cercor/6.1.1 | Words (passive listening) > rest | Passive | 3 | -54 | -55 | 22 |
|  |  |  | Pseudowords (passive listening) > rest | Passive | 3 | -54 | -55 | 24 |
| Hickok et al. | 2003 | 10.1162/jocn.2003.15.5.673 | Pseudowords > rest | Passive | 9 | -47 | -56 | 6 |
|  |  |  |  |  |  | -63 | -31 | 0 |
|  |  |  |  |  |  | -47 | -57 | 6 |
|  |  |  |  |  |  | -63 | -32 | 4 |
| Heim et al. | 2003 | 10.1016/S0926-6410(02)00284-7 | Phoneme judgement > semantics | Decision | 8 | -52 | 23 | 20 |
|  |  |  |  |  |  | -39 | 38 | 8 |
|  |  |  |  |  |  | -45 | 23 | 26 |
|  |  |  |  |  |  | -25 | 6 | 48 |
|  |  |  |  |  |  | -48 | 10 | 21 |
|  |  |  |  |  |  | -45 | 5 | 38 |
|  |  |  |  |  |  | -51 | -52 | 34 |
|  |  |  |  |  |  | -52 | -46 | -1 |
|  |  |  | Vowel judgement > semantics | Decision | 8 | -55 | 19 | 17 |
|  |  |  |  |  |  | -42 | 27 | 29 |
|  |  |  |  |  |  | -4 | 49 | 33 |
|  |  |  |  |  |  | -32 | 3 | 32 |
|  |  |  |  |  |  | -14 | -41 | 9 |
|  |  |  |  |  |  | -55 | -43 | 23 |
|  |  |  |  |  |  | -35 | -41 | 9 |
| Zatorre et al. | 1996 | 10.1093/cercor/6.1.21 | Phoneme discrimination > passive word listening | Decision | 10 | 0 | -22 | -19 |
|  |  |  |  |  |  | -37 | 25 | 17 |
|  |  |  |  |  |  | -20 | -16 | 31 |
|  |  |  | Phoneme monitoring > passive world listening | Decision | 10 | -32 | -48 | 43 |
|  |  |  |  |  |  | -12 | -78 | -19 |
|  |  |  |  |  |  | -16 | -75 | 13 |
|  |  |  |  |  |  | 28 | -82 | 27 |
|  |  |  |  |  |  | 0 | -79 | 9 |
|  |  |  |  |  |  | -36 | -60 | -8 |
|  |  |  |  |  |  | -46 | 13 | 25 |
|  |  |  |  |  |  | -40 | -4 | 44 |
|  |  |  | Phoneme discrimination > pitch discrimination | Decision | 10 | -16 | -60 | -5 |
|  |  |  |  |  |  | 42 | -73 | -7 |
|  |  |  |  |  |  | -59 | 11 | 28 |
|  |  |  |  |  |  | 18 | -104 | 2 |
|  |  |  |  |  |  | -39 | -70 | 27 |
|  |  |  | Phoneme discrimination > noise | Decision | 10 | -45 | 9 | 26 |
|  |  |  |  |  |  | -62 | -13 | -6 |
|  |  |  |  |  |  | -60 | 22 | -11 |
|  |  |  |  |  |  | 61 | 35 | -24 |
|  |  |  |  |  |  | 52 | -81 | -21 |
|  |  |  |  |  |  | 65 | -15 | -4 |
|  |  |  | Phoneme discrimination > passive speech listening | Decision | 10 | -51 | 7 | 22 |
|  |  |  |  |  |  | 9 | -29 | 22 |
|  |  |  |  |  |  | 18 | -96 | 17 |
|  |  |  |  |  |  | 0 | 18 | 24 |
|  |  |  |  |  |  | -29 | -57 | 52 |
|  |  |  |  |  |  | -66 | -38 | -20 |
| Belin et al. | 2000 | 10.1038/35002078 | Vocal > non-vocal sounds (passive listening) | Passive | 8 | 64 | 7 | -18 |
|  |  |  |  |  |  | 66 | 0 | -10 |
|  |  |  |  |  |  | 69 | -12 | -6 |
|  |  |  |  |  |  | 57 | -19 | -5 |
|  |  |  |  |  |  | 62 | -30 | 4 |
|  |  |  |  |  |  | 51 | -45 | 6 |
|  |  |  |  |  |  | 6 | -51 | 34 |
|  |  |  |  |  |  | -64 | -2 | -13 |
|  |  |  |  |  |  | -66 | -14 | -2 |
|  |  |  |  |  |  | -42 | -37 | 14 |
|  |  |  |  |  |  | -66 | -40 | 12 |
| Belin et al. | 2002 | 10.1016/S0926-6410(01)00084-2 | Vocal > non-vocal sounds (passive listening) | Passive | 8 | -66 | -14 | -2 |
|  |  |  |  |  |  | 69 | -12 | -6 |
| Bookheimer et al. | 2000 | 10.1212/WNL.55.8.1151 | Repeating word sequence > rest | Passive | 8 | -55 | -15 | 11 |
|  |  |  |  |  |  | 53 | -17 | 5 |
|  |  |  |  |  |  | 53 | -11 | 0 |
|  |  |  |  |  |  | -50 | -3 | 41 |
|  |  |  |  |  |  | -57 | -1 | 23 |
|  |  |  |  |  |  | 51 | -1 | 17 |
|  |  |  |  |  |  | 45 | 1 | 35 |
|  |  |  |  |  |  | -3 | 5 | 53 |
|  |  |  |  |  |  | -12 | -65 | -25 |
|  |  |  |  |  |  | -23 | -14 | 11 |
|  |  |  | Repeating phoneme sequence > rest | Passive | 8 | -53 | -35 | 18 |
|  |  |  |  |  |  | -57 | -13 | 7 |
|  |  |  |  |  |  | 49 | -18 | -4 |
|  |  |  |  |  |  | 55 | -9 | -5 |
|  |  |  |  |  |  | -50 | -1 | 41 |
|  |  |  |  |  |  | 51 | -1 | 17 |
|  |  |  |  |  |  | 45 | 1 | 35 |
|  |  |  |  |  |  | -3 | 6 | 49 |
|  |  |  |  |  |  | -14 | -62 | -12 |
|  |  |  |  |  |  | -7 | -19 | 11 |

*N.B. Contrasts within the same study conducted using the same sample were combined before they were entered into the meta-analysis.*

Supplementary Table 3. Data included in the semantic control meta-analysis.

| **Author(s)** | **Year** | **DOI** | **Contrast included** | **N** | **MNI coordinates** | | |
| --- | --- | --- | --- | --- | --- | --- | --- |
|  |  |  |  |  | **X** | **Y** | **Z** |
| Abraham | 2018 | 10.1016/j.neuropsychologia.2018.05.004 | Alternative uses task > name typical objects for location | 34 | -52 | 9 | 6 |
|  |  |  |  |  | -45 | 22 | 11 |
|  |  |  |  |  | -36 | -8 | 0 |
|  |  |  |  |  | -13 | 6 | 5 |
|  |  |  |  |  | -13 | -8 | -7 |
|  |  |  |  |  | 16 | -8 | -7 |
|  |  |  |  |  | -3 | -17 | 7 |
|  |  |  |  |  | -19 | -31 | -1 |
|  |  |  |  |  | 55 | 31 | -2 |
|  |  |  |  |  | -6 | 28 | 43 |
|  |  |  |  |  | -3 | 20 | 54 |
|  |  |  |  |  | -13 | 49 | 24 |
|  |  |  |  |  | 0 | -1 | 36 |
|  |  |  |  |  | -32 | 18 | 41 |
|  |  |  |  |  | 16 | 6 | 8 |
|  |  |  |  |  | -58 | -30 | 43 |
|  |  |  |  |  | -48 | -73 | -3 |
|  |  |  |  |  | -26 | -105 | -1 |
|  |  |  |  |  | 16 | -8 | -7 |
|  |  |  |  |  | 26 | -78 | -24 |
|  |  |  |  |  | 42 | -66 | -39 |
|  |  |  |  |  | 13 | -90 | 14 |
|  |  |  |  |  | -6 | -93 | 15 |
| Bitan, Kaftory, Meiri-Leib, Eviatar, Peleg | 2017 | 10.1037/neu0000357 | Subordinate>dominant meanings | 23 | -46 | 20 | 28 |
|  |  |  |  |  | -48 | -32 | -4 |
|  |  |  |  |  | -40 | 4 | 28 |
|  |  |  |  |  | -44 | 42 | -6 |
| Canini, Della Rosa, Catricala, Strijkers, Branzi, Costa, Abutalebi | 2016 | 10.1002/hbm.23304 | Interference time between exemplars of same category | 24 | -28 | 42 | 10 |
|  |  |  |  |  | -14 | 6 | 18 |
| Krieger-Redwood, Teige, Davey, Hymers, Jefferies | 2015 | 10.1016/j.neuropsychologia.2015.02.030 | Weak>strong associations between words | 22 | -4 | 28 | 32 |
|  |  |  |  |  | -4 | 26 | 36 |
|  |  |  |  |  | 0 | 18 | 50 |
|  |  |  |  |  | 4 | 24 | 52 |
|  |  |  |  |  | 0 | 30 | 46 |
|  |  |  |  |  | -6 | 28 | 22 |
|  |  |  |  |  | 2 | -60 | 50 |
|  |  |  |  |  | 16 | -62 | 24 |
|  |  |  |  |  | -8 | -68 | 34 |
|  |  |  |  |  | -4 | -70 | 34 |
|  |  |  |  |  | -6 | -64 | 50 |
|  |  |  |  |  | -6 | -58 | 48 |
|  |  |  |  |  | -34 | 22 | -10 |
|  |  |  |  |  | -46 | 14 | -10 |
|  |  |  |  |  | -14 | 0 | -4 |
|  |  |  |  |  | -52 | 20 | -12 |
|  |  |  |  |  | -48 | 20 | 2 |
|  |  |  |  |  | -16 | 18 | -6 |
|  |  |  |  |  | -32 | 48 | 6 |
|  |  |  |  |  | -22 | 48 | 4 |
|  |  |  |  |  | -26 | 62 | -10 |
|  |  |  |  |  | -28 | 62 | -6 |
|  |  |  |  |  | -28 | 58 | 2 |
|  |  |  |  |  | -16 | 58 | -14 |
|  |  |  |  |  | 32 | 20 | -4 |
|  |  |  |  |  | 40 | 18 | -12 |
|  |  |  |  |  | 30 | 16 | -18 |
|  |  |  |  |  | 42 | 20 | -6 |
|  |  |  |  |  | 22 | 8 | -24 |
|  |  |  |  |  | 22 | 8 | -20 |
|  |  |  |  |  | 54 | 24 | 18 |
|  |  |  |  |  | 54 | 22 | 26 |
|  |  |  |  |  | 48 | 26 | 30 |
|  |  |  |  |  | 60 | 24 | 12 |
|  |  |  |  |  | 36 | 20 | 16 |
| Krieger-Redwood, Teige, Davey, Hymers, Jefferies | 2015 | 10.1016/j.neuropsychologia.2015.02.030 | Weak>strong associations between pictures | 22 | -30 | 50 | 4 |
|  |  |  |  |  | 4 | 34 | 28 |
|  |  |  |  |  | 2 | 34 | 32 |
|  |  |  |  |  | 0 | 20 | 48 |
|  |  |  |  |  | 32 | 50 | 8 |
|  |  |  |  |  | -52 | 14 | 10 |
|  |  |  |  |  | -4 | -48 | 40 |
|  |  |  |  |  | -6 | -64 | 44 |
|  |  |  |  |  | -4 | -54 | 42 |
|  |  |  |  |  | -12 | -50 | 34 |
|  |  |  |  |  | -12 | -64 | 26 |
|  |  |  |  |  | -4 | -68 | 38 |
|  |  |  |  |  | -56 | -60 | 26 |
|  |  |  |  |  | -56 | -44 | 6 |
|  |  |  |  |  | -34 | -80 | 40 |
|  |  |  |  |  | -62 | -40 | -4 |
|  |  |  |  |  | -58 | -40 | -8 |
|  |  |  |  |  | -62 | -50 | 34 |
| Abraham, Pieritz, Thybusch, Rutter, Kroeger, Schweckendiek, Stark, Windmann, Hermann | 2012 | 10.1016/j.neuropsychologia.2012.04.015 | High > low flexibility | 19 | -46 | 18 | 21 |
|  |  |  |  |  | -50 | 22 | 1 |
|  |  |  |  |  | -44 | 37 | -14 |
|  |  |  |  |  | -33 | 36 | 6 |
|  |  |  |  |  | -53 | 10 | -28 |
|  |  |  |  |  | -1 | 24 | 23 |
|  |  |  |  |  | -4 | 20 | 47 |
|  |  |  |  |  | -36 | 10 | 39 |
|  |  |  |  |  | -59 | -32 | 40 |
|  |  |  |  |  | -43 | -25 | 49 |
|  |  |  |  |  | -33 | -10 | -3 |
|  |  |  |  |  | -50 | -59 | -8 |
|  |  |  |  |  | 34 | -81 | -34 |
|  |  |  |  |  | 18 | -92 | -16 |
|  |  |  |  |  | 25 | -100 | -5 |
| Snyder, Banich, Munakata | 2011 | 10.1162/jocn_a_00023 | High > low competition | 18 | -6 | 16 | 60 |
|  |  |  |  |  | -44 | 20 | 14 |
|  |  |  |  |  | -30 | 26 | -16 |
|  |  |  |  |  | 36 | 28 | -12 |
|  |  |  |  |  | 10 | 56 | 12 |
|  |  |  |  |  | -66 | -28 | -6 |
|  |  |  |  |  | 56 | 10 | -36 |
|  |  |  |  |  | 16 | -92 | -8 |
|  |  |  |  |  | -14 | -92 | -14 |
| Snyder, Banich, Munakata | 2011 | 10.1162/jocn_a_00023 | High > low competition with low association strength | 18 | -42 | 20 | 14 |
|  |  |  |  |  | -6 | 14 | 60 |
|  |  |  |  |  | 42 | 26 | -24 |
|  |  |  |  |  | -56 | 28 | -6 |
|  |  |  |  |  | 16 | -32 | -16 |
|  |  |  |  |  | -52 | -52 | -4 |
|  |  |  |  |  | -68 | -30 | -4 |
|  |  |  |  |  | -42 | 24 | -26 |
|  |  |  |  |  | 30 | -74 | -30 |
|  |  |  |  |  | -26 | -58 | -34 |
| Snyder, Banich, Munakata | 2011 | 10.1162/jocn_a_00023 | High > low competition with high association strength | 18 | 10 | 54 | 18 |
| Snyder, Banich, Munakata | 2011 | 10.1162/jocn_a_00023 | Low > high association strength | 18 | -6 | 14 | 60 |
|  |  |  |  |  | 4 | 4 | 64 |
|  |  |  |  |  | -4 | 24 | 54 |
|  |  |  |  |  | -2 | 38 | 26 |
|  |  |  |  |  | 10 | 34 | 22 |
|  |  |  |  |  | -8 | 18 | 38 |
|  |  |  |  |  | 12 | 22 | 38 |
|  |  |  |  |  | -50 | 24 | 18 |
|  |  |  |  |  | -46 | 26 | -4 |
|  |  |  |  |  | -46 | 4 | 46 |
|  |  |  |  |  | -44 | 10 | 38 |
|  |  |  |  |  | -44 | 48 | -12 |
|  |  |  |  |  | -2 | 12 | 52 |
|  |  |  |  |  | -4 | 48 | 48 |
|  |  |  |  |  | -48 | 12 | 2 |
|  |  |  |  |  | 60 | 20 | 6 |
|  |  |  |  |  | 52 | 26 | -6 |
|  |  |  |  |  | 40 | 40 | -20 |
|  |  |  |  |  | -26 | 46 | 24 |
|  |  |  |  |  | 30 | 52 | 32 |
|  |  |  |  |  | 50 | 6 | 44 |
|  |  |  |  |  | 44 | 50 | -18 |
|  |  |  |  |  | 26 | 62 | -12 |
|  |  |  |  |  | 44 | 26 | 42 |
|  |  |  |  |  | 20 | -76 | 2 |
|  |  |  |  |  | -16 | -94 | -12 |
|  |  |  |  |  | -2 | -96 | -18 |
|  |  |  |  |  | 14 | -88 | -16 |
|  |  |  |  |  | -26 | -84 | 18 |
|  |  |  |  |  | 30 | -84 | 10 |
|  |  |  |  |  | -56 | -48 | -2 |
|  |  |  |  |  | -66 | -60 | 12 |
|  |  |  |  |  | 30 | -76 | 14 |
|  |  |  |  |  | -48 | -66 | -22 |
|  |  |  |  |  | 34 | -56 | 16 |
|  |  |  |  |  | 52 | -26 | -10 |
|  |  |  |  |  | 4 | -84 | 44 |
|  |  |  |  |  | 36 | -52 | -26 |
| Balthasar, Huber, Weis | 2011 | 10.1016/j.brainres.2011.06.054 | Homonyms > identification | 18 | -51 | 43 | 2 |
|  |  |  |  |  | 69 | 11 | 5 |
|  |  |  |  |  | -33 | -62 | -6 |
|  |  |  |  |  | 53 | 48 | -8 |
|  |  |  |  |  | 19 | -36 | 55 |
|  |  |  |  |  | 14 | 46 | 9 |
|  |  |  |  |  | 45 | 6 | 56 |
|  |  |  |  |  | 6 | -5 | 5 |
|  |  |  |  |  | -16 | -63 | -14 |
|  |  |  |  |  | -38 | 2 | 0 |
|  |  |  |  |  | -3 | 7 | 36 |
|  |  |  |  |  | -42 | 53 | 9 |
|  |  |  |  |  | -58 | -15 | 31 |
|  |  |  |  |  | 70 | -14 | 34 |
|  |  |  |  |  | 28 | 6 | 56 |
|  |  |  |  |  | -58 | -19 | 29 |
|  |  |  |  |  | 57 | -11 | 21 |
|  |  |  |  |  | -34 | 7 | -28 |
|  |  |  |  |  | -38 | 2 | 4 |
|  |  |  |  |  | -46 | -66 | 27 |
|  |  |  |  |  | 10 | -18 | 3 |
|  |  |  |  |  | 40 | -68 | 9 |
|  |  |  |  |  | 10 | 50 | 42 |
|  |  |  |  |  | -20 | -58 | -6 |
|  |  |  |  |  | -66 | -27 | 34 |
|  |  |  |  |  | 5 | 46 | 13 |
| Balthasar, Huber, Weis | 2011 | 10.1016/j.brainres.2011.06.054 | Homonyms > identification | 18 | 27 | -61 | -29 |
|  |  |  |  |  | -20 | 67 | 15 |
|  |  |  |  |  | 1 | -27 | 4 |
|  |  |  |  |  | -12 | 16 | -18 |
|  |  |  |  |  | 56 | 11 | 6 |
|  |  |  |  |  | 1 | -91 | -6 |
|  |  |  |  |  | 27 | 48 | -4 |
|  |  |  |  |  | 1 | -32 | 4 |
|  |  |  |  |  | 24 | -39 | 66 |
|  |  |  |  |  | -12 | 8 | -13 |
|  |  |  |  |  | 23 | -75 | -12 |
|  |  |  |  |  | 23 | -26 | 37 |
|  |  |  |  |  | -12 | -57 | -30 |
|  |  |  |  |  | 10 | 50 | 13 |
|  |  |  |  |  | 1 | -5 | 5 |
|  |  |  |  |  | 35 | 20 | -23 |
|  |  |  |  |  | 32 | -80 | 12 |
|  |  |  |  |  | -50 | -79 | 16 |
|  |  |  |  |  | 49 | 23 | 25 |
|  |  |  |  |  | 1 | -25 | -12 |
|  |  |  |  |  | -50 | -56 | 38 |
|  |  |  |  |  | 5 | 58 | 12 |
|  |  |  |  |  | -4 | 30 | -21 |
|  |  |  |  |  | 35 | 29 | -20 |
|  |  |  |  |  | -43 | 20 | -26 |
|  |  |  |  |  | 49 | -79 | 19 |
|  |  |  |  |  | -38 | -38 | 14 |
|  |  |  |  |  | 11 | -55 | 66 |
|  |  |  |  |  | 27 | -39 | -26 |
|  |  |  |  |  | -46 | -41 | -12 |
|  |  |  |  |  | -3 | -32 | 29 |
|  |  |  |  |  | 32 | -52 | 31 |
|  |  |  |  |  | 5 | -81 | -22 |
| Hsu, Kraemer, Oliver, Schlichting, Thompson-Schill | 2011 | 10.1162/jocn.2011.21619 | Low > high distance to foil | 12 | -38 | -28 | -8 |
|  |  |  |  |  | -31 | 55 | 51 |
|  |  |  |  |  | 56 | 40 | 13 |
|  |  |  |  |  | -38 | -50 | -14 |
|  |  |  |  |  | -15 | -89 | 1 |
|  |  |  |  |  | 1 | 33 | 34 |
|  |  |  |  |  | 37 | -45 | 45 |
|  |  |  |  |  | -47 | 35 | 10 |
|  |  |  |  |  | -41 | 16 | 31 |
|  |  |  |  |  | 63 | 44 | -5 |
|  |  |  |  |  | -43 | -40 | 62 |
|  |  |  |  |  | 47 | 26 | 47 |
|  |  |  |  |  | 11 | 9 | 19 |
| Hsu, Kraemer, Oliver, Schlichting, Thompson-Schill | 2011 | 10.1162/jocn.2011.21619 | Low > high distance to foil | 12 | -2 | 18 | 41 |
|  |  |  |  |  | -40 | 5 | 49 |
|  |  |  |  |  | -72 | -26 | 6 |
|  |  |  |  |  | -35 | 18 | -1 |
|  |  |  |  |  | -18 | -26 | -3 |
|  |  |  |  |  | 11 | -71 | 15 |
|  |  |  |  |  | -18 | -86 | -2 |
|  |  |  |  |  | 47 | -26 | -2 |
|  |  |  |  |  | 33 | 20 | -8 |
|  |  |  |  |  | -40 | -36 | 50 |
|  |  |  |  |  | -47 | 21 | 21 |
|  |  |  |  |  | 65 | 15 | 26 |
|  |  |  |  |  | -44 | -56 | -12 |
| Rodd, Longe, Randall, Tyler | 2010 | 10.1016/j.neuropsychologia.2009.12.035 | Semantic ambiguity > no ambiguity | 14 | -52 | 10 | 12 |
| Rodd, Longe, Randall, Tyler | 2011 | 10.1016/j.neuropsychologia.2009.12.036 | Semantic ambiguity > syntactic ambiguity | 14 | -52 | 22 | 4 |
| Schnur, Schwartz, Kimberg, Hirshorn, Coslett, Thompson-Schill | 2009 | 10.1073/pnas.0805874106 | More > less interference | 16 | -48 | 21 | 24 |
|  |  |  |  |  | -33 | 15 | -9 |
|  |  |  |  |  | -51 | 9 | 39 |
|  |  |  |  |  | -45 | -66 | 30 |
|  |  |  |  |  | 21 | -9 | 3 |
|  |  |  |  |  | 60 | -51 | 15 |
|  |  |  |  |  | 48 | -54 | 18 |
| Sun, Chen, Zhang, Li, Li, Wei, Yang, Qiu | 2016 | 10.1002/hbm.23246 | Alternative uses task > name typical objects for location | 28 | 12 | -87 | 12 |
| Grindrod, Bilenko, Myers, Blumstein | 2008 | 10.1016/j.brainres.2008.07.017 | Ambiguous incongruent meanings>single meaning | 15 | 1 | -67 | 28 |
| Grindrod, Bilenko, Myers, Blumstein | 2008 | 10.1016/j.brainres.2008.07.017 | Ambiguous congruent meanings>unrelated words | 15 | -49 | 32 | -11 |
|  |  |  |  |  | 7 | 50 | 43 |
|  |  |  |  |  | -58 | -38 | 6 |
| Jackson, Hoffman, Pobric, Lambon Ralph | 2015 | 10.1093/cercor/bhv003 | High>low association strength | 24 | -51 | 15 | 27 |
|  |  |  |  |  | -9 | -96 | -9 |
|  |  |  |  |  | 48 | 18 | 27 |
|  |  |  |  |  | 36 | 21 | 54 |
|  |  |  |  |  | -39 | -21 | -24 |
|  |  |  |  |  | 30 | 24 | -6 |
|  |  |  |  |  | 18 | -93 | -3 |
|  |  |  |  |  | -30 | -69 | 45 |
| Madore | 2019 | 10.1093/cercor/bhx312 | Alternate uses > associations after control induction | 32 | -48 | -31 | 36 |
|  |  |  |  |  | -51 | -62 | -2 |
|  |  |  |  |  | 22 | -68 | -24 |
|  |  |  |  |  | 58 | -58 | -6 |
|  |  |  |  |  | -38 | -12 | -2 |
|  |  |  |  |  | 0 | 5 | 36 |
|  |  |  |  |  | 60 | -24 | 42 |
|  |  |  |  |  | -26 | -44 | 64 |
|  |  |  |  |  | 30 | -54 | 70 |
|  |  |  |  |  | 27 | -7 | 56 |
|  |  |  |  |  | -50 | 5 | 24 |
|  |  |  |  |  | -26 | -10 | 62 |
|  |  |  |  |  | 32 | -38 | 44 |
|  |  |  |  |  | -20 | -1 | 70 |
|  |  |  |  |  | -44 | -38 | 53 |
|  |  |  |  |  | -26 | 32 | -16 |
|  |  |  |  |  | -6 | -56 | 62 |
|  |  |  |  |  | 57 | 38 | 4 |
|  |  |  |  |  | -8 | -49 | 62 |
|  |  |  |  |  | 22 | -67 | -46 |
|  |  |  |  |  | -34 | -48 | 64 |
|  |  |  |  |  | 24 | -48 | 68 |
|  |  |  |  |  | -46 | -40 | -18 |
| Madore | 2019 | 10.1093/cercor/bhx312 | Alternate uses > associations after specificity induction | 32 | -58 | -31 | 42 |
|  |  |  |  |  | 0 | 2 | 38 |
|  |  |  |  |  | 56 | -66 | -1 |
|  |  |  |  |  | 30 | -74 | -24 |
|  |  |  |  |  | -52 | -67 | -1 |
|  |  |  |  |  | -22 | -2 | 68 |
|  |  |  |  |  | 46 | -26 | 42 |
|  |  |  |  |  | 46 | 26 | -10 |
|  |  |  |  |  | 33 | -34 | 41 |
|  |  |  |  |  | -39 | -12 | 0 |
|  |  |  |  |  | 46 | -43 | -36 |
|  |  |  |  |  | 46 | -79 | 17 |
|  |  |  |  |  | -39 | -7 | 10 |
|  |  |  |  |  | 32 | -49 | 64 |
|  |  |  |  |  | 26 | -6 | 66 |
|  |  |  |  |  | -18 | -73 | -22 |
|  |  |  |  |  | 15 | -74 | -49 |
|  |  |  |  |  | -33 | 38 | -13 |
|  |  |  |  |  | -44 | 2 | 22 |
|  |  |  |  |  | 9 | -78 | 2 |
|  |  |  |  |  | -48 | -49 | -20 |
|  |  |  |  |  | -26 | -2 | -25 |
|  |  |  |  |  | 48 | -37 | -20 |
|  |  |  |  |  | 14 | 42 | 47 |
|  |  |  |  |  | 32 | -12 | -14 |
|  |  |  |  |  | -12 | -78 | -48 |
|  |  |  |  |  | 18 | -88 | 23 |
|  |  |  |  |  | -28 | -44 | 70 |
|  |  |  |  |  | -15 | 6 | 68 |
|  |  |  |  |  | -46 | 34 | 11 |
|  |  |  |  |  | -44 | -80 | 10 |
|  |  |  |  |  | 12 | -74 | -4 |
|  |  |  |  |  | 33 | -2 | -18 |
|  |  |  |  |  | -9 | -55 | 65 |
|  |  |  |  |  | 6 | -84 | 30 |
|  |  |  |  |  | 52 | 38 | 6 |
| Musz, Thompson-Schill | 2017 | 10.1016/j.bandl.2016.11.002 | Subordinate > dominant homonym meaning | 13 | -49 | 40 | 5 |
| Musz, Thompson-Schill | 2017 | 10.1016/j.bandl.2016.11.002 | Delayed > immediate subordinate homonoym meaning | 13 | -49 | 30 | 2 |
| Vitello, Warren, Devlin, Rodd | 2014 | 10.3389/fnhum.2014.00530 | Ambiguous word in sentence > not | 20 | -45 | 32 | 4 |
|  |  |  |  |  | -45 | -55 | -11 |
|  |  |  |  |  | -48 | -58 | -8 |
| Hallam, Whitney, Hymers, Gouws, Jefferies | 2016 | 10.1016/j.neuropsychologia.2016.09.012 | Weak > strong association | 18 | -52 | 24 | 20 |
|  |  |  |  |  | -4 | 16 | 56 |
|  |  |  |  |  | 36 | 28 | -8 |
|  |  |  |  |  | 12 | -80 | -32 |
|  |  |  |  |  | 44 | 24 | 24 |
|  |  |  |  |  | 12 | 12 | 48 |
|  |  |  |  |  | -52 | -40 | 0 |
|  |  |  |  |  | 20 | -80 | -48 |
| Jeon | 2012 | 10.1097/WNR.0b013e32835a19ae | Ambiguous homonym > unambiguous word | 14 | 9 | 34 | -14 |
|  |  |  |  |  | 47 | -46 | -10 |
|  |  |  |  |  | -50 | 21 | 47 |
|  |  |  |  |  | -25 | 32 | -21 |
| Mestres-Misse, Trampel, Turner, Kotz | 2016 | 10.1007/s00429-014-0966-7 | Ambiguous homonym > unambiguous word | 23 | 35 | 24 | -6 |
|  |  |  |  |  | -42 | 21 | -3 |
|  |  |  |  |  | -50 | 35 | 13 |
|  |  |  |  |  | -35 | 25 | -5 |
|  |  |  |  |  | -51 | 32 | 0 |
|  |  |  |  |  | -51 | 13 | 19 |
| Hoenig, Scheef | 2009 | 10.1016/j.neuroimage.2008.12.044 | Ambiguous homonym > unambiguous word | 22 | -27 | 7 | -34 |
|  |  |  |  |  | -5 | 54 | 33 |
|  |  |  |  |  | -48 | -61 | 45 |
|  |  |  |  |  | -13 | 28 | 53 |
|  |  |  |  |  | -7 | -64 | 27 |
|  |  |  |  |  | -37 | 17 | 44 |
|  |  |  |  |  | 23 | 47 | 40 |
|  |  |  |  |  | 60 | -62 | 45 |
|  |  |  |  |  | 31 | -17 | -42 |
|  |  |  |  |  | -1 | 46 | -16 |
|  |  |  |  |  | 35 | 22 | -37 |
|  |  |  |  |  | -47 | 8 | -49 |
|  |  |  |  |  | -6 | 53 | -1 |
|  |  |  |  |  | -48 | -83 | 34 |
|  |  |  |  |  | 30 | 27 | 64 |
| Grindrod, Garnett, Malyutina, den Ouden | 2014 | 10.1016/j.bandl.2014.10.001 | Ambiguous homonym > unambiguous word | 23 | 18 | 29 | 4 |
|  |  |  |  |  | -54 | 38 | 4 |
| Grindrod, Garnett, Malyutina, den Ouden | 2014 | 10.1016/j.bandl.2014.10.001 | Balanced ambiguous > unambiguous nouns | 23 | 18 | 29 | 4 |
| Grindrod, Garnett, Malyutina, den Ouden | 2014 | 10.1016/j.bandl.2014.10.001 | Balanced ambiguous > unambiguous nouns and verbs | 23 | -6 | 14 | 4 |
|  |  |  |  |  | 18 | 29 | 4 |
|  |  |  |  |  | -18 | 20 | -5 |
|  |  |  |  |  | -51 | 14 | -17 |
|  |  |  |  |  | 24 | 2 | -5 |
|  |  |  |  |  | -54 | 38 | 4 |
| Grindrod, Garnett, Malyutina, den Ouden | 2014 | 10.1016/j.bandl.2014.10.001 | Unbalanced ambiguous > unambiguous noun and verbs | 23 | 18 | 29 | 4 |
| Rodd, Johnsrude, Davis | 2012 | 10.1093/cercor/bhr252 | Ambiguity across time | 15 | -52 | 22 | 18 |
|  |  |  |  |  | -52 | 36 | 2 |
|  |  |  |  |  | -44 | 20 | 26 |
|  |  |  |  |  | -44 | 12 | 32 |
|  |  |  |  |  | -48 | 38 | -10 |
|  |  |  |  |  | -48 | 20 | -8 |
|  |  |  |  |  | -44 | -50 | -18 |
|  |  |  |  |  | -60 | -48 | 8 |
|  |  |  |  |  | -48 | -56 | -12 |
|  |  |  |  |  | -54 | -44 | -8 |
|  |  |  |  |  | -52 | -38 | -2 |
|  |  |  |  |  | -56 | -58 | 0 |
|  |  |  |  |  | -50 | -44 | 6 |
|  |  |  |  |  | -62 | -50 | 10 |
|  |  |  |  |  | -32 | -38 | -18 |
|  |  |  |  |  | -56 | -26 | -6 |
| Tahmasebi, Davis, Wild, Rodd, Hakyemez, Abolmaesumi, Johnsrude | 2012 | 10.1093/cercor/bhr205 | Ambiguous homonym > unambiguous word | 24 | -49 | -67 | -4 |
|  |  |  |  |  | -53 | 10 | 25 |
| Hargreaves, Pexman, Pittman, Goodyear | 2011 | 10.1027/1618-3169/a000062 | Ambiguous > not | 20 | -40 | 8 | 24 |
|  |  |  |  |  | -59 | 24 | 12 |
| Bekinschtein, Davis, Rodd, Owen | 2011 | 10.1523/JNEUROSCI.5058-10.2011 | Ambiguous homonym > unambiguous word | 12 | -42 | -62 | -16 |
|  |  |  |  |  | -44 | -48 | -20 |
| Satpute, Badre, Ochsner | 2014 | 10.1093/cercor/bhs408 | Weakly-related > strongly-related target | 33 | -39 | 27 | 18 |
|  |  |  |  |  | 51 | 33 | 24 |
|  |  |  |  |  | 9 | 15 | 0 |
|  |  |  |  |  | -33 | -21 | 57 |
|  |  |  |  |  | 12 | -78 | 12 |
|  |  |  |  |  | -18 | -84 | -9 |
| Tylen, Christensen, Roepstorff, Lund, Ostergaard, Donald | 2015 | 10.1016/j.neuroimage.2015.07.047 | Incoherent > coherent story | 24 | 4 | 28 | 34 |
|  |  |  |  |  | 34 | 48 | 28 |
|  |  |  |  |  | 52 | -44 | 50 |
|  |  |  |  |  | -56 | -42 | 52 |
|  |  |  |  |  | 46 | 40 | 30 |
|  |  |  |  |  | -32 | 42 | 28 |
|  |  |  |  |  | 58 | -36 | -16 |
|  |  |  |  |  | 36 | 16 | 0 |
|  |  |  |  |  | -36 | 14 | 2 |
|  |  |  |  |  | 26 | 8 | 64 |
|  |  |  |  |  | -16 | 6 | 64 |
|  |  |  |  |  | -58 | -58 | -10 |
| Li, Jiang, Yu, Zhou | 2014 | 10.1093/scan/nst091 | Incongruent > congruent sentences | 24 | 4 | 44 | 18 |
|  |  |  |  |  | 2 | 40 | 34 |
| Mestres-Misse, Bazin, Trampel, Turner, Kotz | 2014 | 10.1016/j.neuroimage.2014.05.002 | Subordinate > dominant homonym meaning | 23 | 10 | 13 | 3 |
| Mestres-Misse, Bazin, Trampel, Turner, Kotz | 2014 | 10.1016/j.neuroimage.2014.05.002 | Ambiguous > unambiguous word in sentences | 23 | -45 | 26 | 18 |
|  |  |  |  |  | -51 | 23 | 23 |
|  |  |  |  |  | -48 | 23 | 21 |
|  |  |  |  |  | -53 | 35 | 10 |
|  |  |  |  |  | -49 | 27 | 6 |
|  |  |  |  |  | -56 | 18 | 17 |
|  |  |  |  |  | -43 | 23 | -2 |
|  |  |  |  |  | -29 | 24 | -1 |
|  |  |  |  |  | -52 | 32 | 0 |
|  |  |  |  |  | 30 | 24 | -2 |
|  |  |  |  |  | -14 | 3 | 12 |
|  |  |  |  |  | 44 | 29 | 14 |
|  |  |  |  |  | 51 | 26 | 23 |
|  |  |  |  |  | -63 | -39 | -1 |
|  |  |  |  |  | -60 | -41 | 5 |
| Mestres-Misse, Bazin, Trampel, Turner, Kotz | 2014 | 10.1016/j.neuroimage.2014.05.002 | Incongruent>congruent sentence | 23 | -49 | 30 | 9 |
|  |  |  |  |  | -60 | -37 | 8 |
| Deen, McCarthy | 2010 | 10.1016/j.neuropsychologia.2010.01.028 | Incongruent>congruent story end | 15 | -2 | 24 | 20 |
|  |  |  |  |  | 40 | 10 | -2 |
|  |  |  |  |  | -30 | 22 | -4 |
| Smirnov, Glerean, Lahnakoski, Salmi, Jaaskelainen, Sams, Nummenmaa | 2014 | 10.1016/j.neuropsychologia.2014.09.007 | Miscue > cue | 20 | -43 | 18 | 18 |
| Newman, Ikuta, Burns | 2010 | 10.1016/j.bandl.2010.02.001 | Unrelated > related sentence | 20 | -28 | 18 | -4 |
|  |  |  |  |  | -44 | 24 | 4 |
|  |  |  |  |  | -8 | 6 | 56 |
|  |  |  |  |  | -44 | -2 | 40 |
|  |  |  |  |  | -26 | -58 | 42 |
|  |  |  |  |  | -2 | -58 | -20 |
|  |  |  |  |  | -6 | -16 | -12 |
|  |  |  |  |  | 34 | 22 | -2 |
|  |  |  |  |  | 42 | 22 | 24 |
|  |  |  |  |  | 30 | -58 | 42 |
|  |  |  |  |  | 40 | -16 | 62 |
| Mano, Harada, Sugiura, Saito, Sadato | 2009 | 10.1016/j.neuropsychologia.2008.12.011 | Less > more coherent texts | 18 | 2 | -68 | 26 |
|  |  |  |  |  | 46 | -72 | 28 |
| Tune, Schlesewsky, Nagels, Small, Bornkessel-Schlesewsky | 2016 | 10.1016/j.neuroimage.2016.05.020 | Easy anomalous > normal sentences | 18 | -61 | -32 | 26 |
|  |  |  |  |  | -50 | -15 | 2 |
|  |  |  |  |  | -47 | -43 | -18 |
|  |  |  |  |  | -7 | -82 | 31 |
|  |  |  |  |  | -4 | -16 | 33 |
|  |  |  |  |  | -22 | -37 | 1 |
|  |  |  |  |  | 17 | 54 | 23 |
|  |  |  |  |  | 58 | -29 | 35 |
|  |  |  |  |  | 33 | 5 | -16 |
|  |  |  |  |  | 47 | -58 | 38 |
|  |  |  |  |  | 27 | 15 | 43 |
|  |  |  |  |  | 65 | -22 | 1 |
| Tune, Schlesewsky, Nagels, Small, Bornkessel-Schlesewsky | 2016 | 10.1016/j.neuroimage.2016.05.020 | Hard anomalous > normal sentences | 18 | -49 | -45 | 45 |
|  |  |  |  |  | -7 | 40 | 4 |
|  |  |  |  |  | -2 | -9 | 12 |
|  |  |  |  |  | -59 | -34 | 39 |
|  |  |  |  |  | 63 | -47 | -2 |
|  |  |  |  |  | 18 | -4 | -14 |
|  |  |  |  |  | 51 | 33 | -1 |
|  |  |  |  |  | 2 | -7 | 9 |
|  |  |  |  |  | 63 | -32 | -14 |
| Peelle, Troiani, Grossman | 2009 | 10.1016/j.neuropsychologia.2008.10.027 | Inconsistent > consistent description for nominal kinds | 25 | -44 | -66 | 30 |
|  |  |  |  |  | -50 | -18 | -14 |
|  |  |  |  |  | 0 | -40 | 40 |
|  |  |  |  |  | 44 | -68 | 34 |
|  |  |  |  |  | 60 | -2 | -18 |
|  |  |  |  |  | -32 | 58 | 6 |
|  |  |  |  |  | 0 | 48 | -6 |
|  |  |  |  |  | -48 | 30 | 18 |
|  |  |  |  |  | -48 | 38 | -14 |
|  |  |  |  |  | 30 | 58 | 2 |
|  |  |  |  |  | -10 | -26 | -24 |
|  |  |  |  |  | 4 | -54 | 20 |
|  |  |  |  |  | 4 | -16 | 14 |
| Peelle, Troiani, Grossman | 2009 | 10.1016/j.neuropsychologia.2008.10.027 | Inconsistent > consistent description for nouns | 25 | 30 | 10 | 44 |
|  |  |  |  |  | 38 | 14 | 48 |
| Zhu, Hagoort, Zhang, Feng, Chen, Bastiaansen, Wang | 2012 | 10.1016/j.neuroimage.2012.02.036 | Incongruence (cloze probability) in reading task | 27 | -2 | 20 | 54 |
|  |  |  |  |  | -48 | 26 | 24 |
|  |  |  |  |  | 50 | 20 | 30 |
|  |  |  |  |  | -56 | -48 | -4 |
|  |  |  |  |  | -30 | -60 | 58 |
|  |  |  |  |  | 32 | -66 | 56 |
| Zhu, Hagoort, Zhang, Feng, Chen, Bastiaansen, Wang | 2012 | 10.1016/j.neuroimage.2012.02.036 | Incongruence (cloze probability) in semantic task | 27 | -42 | 40 | 34 |
|  |  |  |  |  | -2 | 22 | 54 |
|  |  |  |  |  | 32 | 56 | 12 |
|  |  |  |  |  | -54 | -46 | 40 |
| Zhu, Hagoort, Zhang, Feng, Chen, Bastiaansen, Wang | 2012 | 10.1016/j.neuroimage.2012.02.036 | Incongruence (cloze probability) in font task | 27 | -42 | 48 | -2 |
|  |  |  |  |  | -18 | -84 | -26 |
|  |  |  |  |  | 32 | -62 | 46 |
|  |  |  |  |  | 10 | -84 | -24 |
| Willems, Frank, Nijhof, Hagoort, van den Bosch | 2016 | 10.1093/cercor/bhv075 | Word context surprisal | 24 | -46 | -46 | -16 |
|  |  |  |  |  | -64 | -36 | 12 |
|  |  |  |  |  | -58 | -50 | 10 |
|  |  |  |  |  | -54 | -58 | 6 |
|  |  |  |  |  | -52 | 0 | -4 |
|  |  |  |  |  | -56 | 8 | -6 |
|  |  |  |  |  | 56 | 8 | -6 |
|  |  |  |  |  | 30 | -2 | -11 |
|  |  |  |  |  | 12 | -12 | -8 |
|  |  |  |  |  | 32 | -6 | -12 |
|  |  |  |  |  | 50 | 12 | -4 |
|  |  |  |  |  | 64 | -26 | 10 |
| Carter, Foster, Muncy, Luke | 2019 | 10.1016/j.neuroimage.2019.01.018 | Word context surprisal | 41 | 2 | 44 | -5 |
|  |  |  |  |  | 2 | -59 | 62 |
|  |  |  |  |  | 62 | -44 | 26 |
|  |  |  |  |  | 32 | -86 | 32 |
|  |  |  |  |  | -41 | -17 | 5 |
|  |  |  |  |  | 59 | -65 | 14 |
| Carter, Foster, Muncy, Luke | 2019 | 10.1016/j.neuroimage.2019.01.018 | Semantic unpredictability | 41 | -2 | -74 | 53 |
|  |  |  |  |  | -2 | 53 | -5 |
|  |  |  |  |  | 53 | -71 | -29 |
|  |  |  |  |  | -26 | 20 | 44 |
| Obleser, Kotz | 2010 | 10.1093/cercor/bhp128 | Low > high cloze probability | 16 | -60 | 12 | 16 |
|  |  |  |  |  | -50 | -42 | 2 |
|  |  |  |  |  | 40 | -32 | 8 |
| Obleser, Kotz | 2010 | 10.1093/cercor/bhp128 | Low > high cloze probability with high intelligibility | 16 | -60 | 14 | 14 |
|  |  |  |  |  | -52 | -8 | -16 |
|  |  |  |  |  | 46 | -18 | -6 |
| Scharinger, Bendixen, Herrmann, Henry, Mildner, Obleser | 2016 | 10.1002/hbm.23060 | Unpredictable > predictable word for incomplete > complete sentence | 22 | 60 | -4 | -2 |
|  |  |  |  |  | -57 | -16 | -2 |
|  |  |  |  |  | -42 | -49 | 49 |
|  |  |  |  |  | -42 | -49 | 34 |
| Huang, Zhu, Zhang, Wu, Chen, Wang | 2012 | 10.1016/j.brainres.2011.11.060 | Unexpected > expected word | 23 | -36 | 26 | -4 |
|  |  |  |  |  | 0 | 24 | 50 |
|  |  |  |  |  | -52 | -80 | 4 |
|  |  |  |  |  | -58 | -39 | 30 |
|  |  |  |  |  | 35 | 42 | -3 |
| Kambara, Tsukiura, Yokoyama, Takahashi, Shigemune, Miyamoto, Takahashi, Sato, Kawashima | 2013 | 10.1016/j.langsci.2012.07.003 | Semantic violations > normal sentences | 38 | -38 | 30 | -23 |
|  |  |  |  |  | -18 | 63 | 12 |
|  |  |  |  |  | 41 | 48 | -22 |
|  |  |  |  |  | 33 | -5 | -9 |
| Rothermich, Kotz | 2013 | 10.1016/j.neuroimage.2012.12.013 | Unpredictable > predictable | 16 | -54 | 29 | 16 |
| Rothermich, Kotz | 2013 | 10.1016/j.neuroimage.2012.12.013 | Unpredictable > predictable with normal intonation | 16 | -54 | 29 | 16 |
| Rothermich, Kotz | 2013 | 10.1016/j.neuroimage.2012.12.013 | Unpredictable > predictable with abnormal intonation | 16 | -54 | 29 | 16 |
|  |  |  |  |  | -6 | 17 | 46 |
|  |  |  |  |  | -6 | 5 | 58 |
| Nieuwland, Martin, Carreiras | 2012 | 10.1002/hbm.21377 | Semantic anomalies > normal sentences | 24 | -50 | 34 | -16 |
|  |  |  |  |  | -36 | 26 | -24 |
|  |  |  |  |  | -32 | 20 | -20 |
|  |  |  |  |  | 46 | 38 | -18 |
|  |  |  |  |  | 44 | 28 | -14 |
|  |  |  |  |  | 28 | 16 | -18 |
|  |  |  |  |  | -4 | 42 | 44 |
|  |  |  |  |  | -6 | 48 | 22 |
|  |  |  |  |  | 6 | 38 | 56 |
| Ye, Donamayor, Muente | 2014 | 10.1002/hbm.22182 | Incongruent > congruent sentences | 20 | 6 | 16 | 58 |
|  |  |  |  |  | -44 | 26 | -2 |
|  |  |  |  |  | 56 | 19 | 1 |
| van de Meerendonk, Rueschemeyer, Kolk | 2013 | 10.1016/j.bandl.2013.07.004 | Less congruent > more congruent words | 24 | 18 | 4 | 18 |
|  |  |  |  |  | 38 | 20 | 22 |
|  |  |  |  |  | 36 | 36 | 8 |
|  |  |  |  |  | -40 | 40 | -2 |
|  |  |  |  |  | -40 | 40 | -14 |
|  |  |  |  |  | -52 | 32 | 10 |
|  |  |  |  |  | 42 | -8 | 40 |
|  |  |  |  |  | 52 | -4 | 44 |
|  |  |  |  |  | 54 | -8 | 22 |
|  |  |  |  |  | -28 | -8 | 40 |
|  |  |  |  |  | -46 | -12 | 44 |
|  |  |  |  |  | -36 | -10 | 44 |
|  |  |  |  |  | 12 | -44 | -38 |
|  |  |  |  |  | 20 | -40 | -36 |
| van de Meerendonk, Rueschemeyer, Kolk | 2013 | 10.1016/j.bandl.2013.07.004 | Incongruent > congruent words | 24 | 38 | 20 | 20 |
|  |  |  |  |  | 50 | 30 | 10 |
|  |  |  |  |  | 32 | 28 | -12 |
|  |  |  |  |  | 50 | 0 | -22 |
|  |  |  |  |  | 42 | -2 | -22 |
|  |  |  |  |  | 50 | 8 | -30 |
| Willems, Ozyurek, Hagoort | 2008 | 10.1162/jocn.2008.20085 | Incongruent > congruent | 19 | -45 | 22 | 20 |
|  |  |  |  |  | -53 | -30 | -7 |
|  |  |  |  |  | -40 | -61 | 18 |
|  |  |  |  |  | 19 | -42 | -31 |
| Willems, Ozyurek, Hagoort | 2008 | 10.1162/jocn.2008.20085 | Incongruent > congruent words and sentences | 19 | -47 | 20 | 21 |
|  |  |  |  |  | -55 | -35 | -1 |
| Willems, Ozyurek, Hagoort | 2008 | 10.1162/jocn.2008.20085 | Incongruent > congruent pictures and sentences | 19 | -40 | 13 | 31 |
| Raposo, Mendes, Marques | 2012 | 10.1016/j.neuroimage.2011.08.072 | Rarer > prototypical features | 17 | -60 | 20 | 12 |
| Raposo, Mendes, Marques | 2012 | 10.1016/j.neuroimage.2011.08.072 | Rarer > prototypical features of basic item | 17 | -54 | 24 | 8 |
|  |  |  |  |  | 4 | 30 | 10 |
|  |  |  |  |  | -44 | 24 | -12 |
| Kroeger, Rutter, Stark, Windmann, Hermann & Abraham | 2012 | 10.1016/j.brainres.2011.10.031 | Appropriate and unusual > inappropriate and usual uses | 19 | -36 | 35 | 4 |
|  |  |  |  |  | -45 | 11 | 16 |
|  |  |  |  |  | 51 | 38 | 4 |
|  |  |  |  |  | 27 | 29 | -11 |
|  |  |  |  |  | -27 | 23 | -14 |
|  |  |  |  |  | 51 | 26 | 1 |
|  |  |  |  |  | -9 | 23 | 46 |
|  |  |  |  |  | -21 | 14 | 52 |
|  |  |  |  |  | -9 | 26 | 22 |
|  |  |  |  |  | -9 | 29 | 28 |
|  |  |  |  |  | -6 | 32 | 31 |
|  |  |  |  |  | 9 | -13 | -8 |
|  |  |  |  |  | -9 | -5 | -1 |
|  |  |  |  |  | -6 | -10 | -8 |
| Rutter, Kroeger, Stark, Schweckendiek,Windmann, Hermann, Abraham | 2012 | 10.1016/j.bandc.2011.11.002 | Appropriate and unusual > inappropriate and usual word choices | 18 | -48 | 17 | 4 |
|  |  |  |  |  | 60 | 17 | 4 |
|  |  |  |  |  | -42 | 20 | 1 |
|  |  |  |  |  | -36 | 29 | -8 |
|  |  |  |  |  | -33 | 50 | 13 |
|  |  |  |  |  | -30 | 53 | 19 |
|  |  |  |  |  | -48 | 20 | -14 |
| Zhu, Feng, Zhang, Li, Li, Wang | 2013 | 10.1016/j.neuroimage.2013.02.060 | Violation > low cloze > high cloze parametric modulator in orthographic task | 26 | -40 | 20 | 24 |
|  |  |  |  |  | -64 | -46 | -4 |
|  |  |  |  |  | -46 | -10 | -20 |
|  |  |  |  |  | -4 | 22 | 52 |
| Zhu, Feng, Zhang, Li, Li, Wang | 2013 | 10.1016/j.neuroimage.2013.02.060 | Violation > low cloze > high cloze parametric modulator in semantic task | 26 | -46 | 22 | 34 |
|  |  |  |  |  | -8 | -70 | 38 |
|  |  |  |  |  | -36 | -58 | 42 |
|  |  |  |  |  | 0 | 26 | 38 |
|  |  |  |  |  | 46 | 26 | 30 |
|  |  |  |  |  | 24 | 50 | -12 |
|  |  |  |  |  | 4 | -26 | 26 |
| Sitnikova, Rosen, Lord, West | 2014 | 10.1016/j.neuroimage.2014.09.012 | Novel > typical object use | 16 | -42 | 0 | 45 |
|  |  |  |  |  | -46 | 6 | 14 |
|  |  |  |  |  | -31 | 24 | 40 |
| Moberget, Gullesen, Andersson, Ivry, Endestad | 2014 | 10.1523/JNEUROSCI.2264-13.2014 | Incongruent > congruent sentences | 32 | -10 | -84 | -31 |
|  |  |  |  |  | 18 | -74 | -29 |
|  |  |  |  |  | -10 | -84 | -29 |
|  |  |  |  |  | 18 | -74 | -29 |
|  |  |  |  |  | 9 | 29 | 41 |
|  |  |  |  |  | -54 | -40 | 8 |
| Clos, Langner, Meyer, Oechslin, Zilles, Eickhoff | 2014 | 10.1002/hbm.22151 | Cue different > same sentence | 29 | -52 | 18 | 18 |
| Van Ettinger-Veenstra, McAllister, Lundberg, Karlsson, Engstrom | 2016 | 10.3389/fnhum.2016.00110 | Incongruent > congruent sentences | 27 | -50 | 28 | 6 |
|  |  |  |  |  | 52 | 26 | 24 |
| Tobia, Michael J.; Madan, Christopher R. | 2017 | 10.14814/phy2.13078 | Subordinate > prototypical uses | 16 | -47 | -35 | 38 |
|  |  |  |  |  | -25 | -62 | 35 |
|  |  |  |  |  | -42 | 12 | 28 |
|  |  |  |  |  | -9 | 24 | 48 |
| Ferstl et al | 2005 | 10.1162/0898929053747658 | Incongruent > congruent | 20 | 44 | 1 | -19 |
| Allen et al. |  | 10.1162/jocn.2008.20107 | Inappropriate > appropriate | 15 | -44 | 30 | 30 |
|  |  |  |  |  | -42 | 38 | 32 |
|  |  |  |  |  | 8 | -70 | 50 |
|  |  |  |  |  | -4 | -70 | 54 |
|  |  |  |  |  | -24 | 32 | -18 |
| Badre et al. | 2005 | 10.1016/j.neuron.2005.07.023 | Weak > strong association | 22 | -51 | 27 | -3 |
|  |  |  |  |  | -48 | 30 | -12 |
|  |  |  |  |  | -45 | 9 | 51 |
|  |  |  |  |  | -51 | 9 | 33 |
|  |  |  |  |  | -39 | 12 | 48 |
|  |  |  |  |  | -42 | 12 | 18 |
|  |  |  |  |  | -48 | 15 | 24 |
|  |  |  |  |  | -51 | 21 | 21 |
|  |  |  |  |  | -9 | 21 | 42 |
|  |  |  |  |  | -45 | 27 | 15 |
|  |  |  |  |  | -33 | 27 | -6 |
|  |  |  |  |  | -48 | 30 | 12 |
|  |  |  |  |  | -36 | 30 | -3 |
|  |  |  |  |  | -45 | 42 | -9 |
|  |  |  |  |  | 33 | 0 | 57 |
|  |  |  |  |  | 42 | 12 | 27 |
|  |  |  |  |  | 6 | 21 | 51 |
|  |  |  |  |  | 9 | 30 | 12 |
|  |  |  |  |  | 36 | 30 | -9 |
| Badre et al. | 2005 | 10.1016/j.neuron.2005.07.023 | Incongruent > congruent relations | 22 | -39 | 3 | 27 |
|  |  |  |  |  | -42 | 9 | 21 |
|  |  |  |  |  | -54 | 12 | 18 |
|  |  |  |  |  | -21 | 15 | 60 |
|  |  |  |  |  | -48 | 18 | 18 |
|  |  |  |  |  | -30 | 27 | -18 |
|  |  |  |  |  | -54 | 30 | 12 |
|  |  |  |  |  | -3 | 33 | 48 |
|  |  |  |  |  | -45 | 39 | 3 |
|  |  |  |  |  | 30 | 24 | -15 |
| Bedny et al | 2008 | 10.1093/cercor/bhn018 | Ambiguous > not | 20 | -54 | 9 | 39 |
|  |  |  |  |  | -3 | 33 | 42 |
|  |  |  |  |  | 36 | -69 | -39 |
|  |  |  |  |  | -24 | 57 | 24 |
|  |  |  |  |  | -12 | -6 | 18 |
|  |  |  |  |  | -45 | -57 | 54 |
|  |  |  |  |  | -57 | -42 | -3 |
| Bedny et al | 2008 | 10.1093/cercor/bhn018 | Ambiguous conjunction | 20 | 0 | 27 | 42 |
|  |  |  |  |  | 39 | -66 | -39 |
|  |  |  |  |  | -48 | 15 | 9 |
|  |  |  |  |  | -51 | 6 | 45 |
| Bunge | 2005 | https://academic.oup.com/cercor/article/15/3/239/375113 | Low > high relation between probe and target | 20 | -57 | 24 | 6 |
| Chan et al. | 2004 | 10.1016/j.neuroimage.2004.02.034 | Semantic ambiguity > non-ambiguous words | 8 | -37 | 59 | 26 |
|  |  |  |  |  | -10 | 68 | 18 |
|  |  |  |  |  | -47 | 48 | 21 |
|  |  |  |  |  | -39 | -17 | 67 |
|  |  |  |  |  | 24 | 2 | 64 |
|  |  |  |  |  | 49 | 34 | 15 |
|  |  |  |  |  | 12 | 67 | 19 |
|  |  |  |  |  | 6 | 39 | -22 |
|  |  |  |  |  | 37 | -17 | 46 |
|  |  |  |  |  | 49 | -59 | 45 |
|  |  |  |  |  | -14 | -92 | 18 |
|  |  |  |  |  | -52 | -74 | -5 |
|  |  |  |  |  | 14 | -93 | 45 |
|  |  |  |  |  | 10 | -75 | 14 |
|  |  |  |  |  | 7 | -82 | -6 |
|  |  |  |  |  | 7 | 22 | -17 |
|  |  |  |  |  | 6 | -30 | 26 |
| de Zubicaray et al. | 2000 | 10.1016/S0028-3932(00)00026-9 | Inappropriate > appropriate responses | 8 | 1 | 51 | -17 |
|  |  |  |  |  | 2 | 36 | 39 |
|  |  |  |  |  | 4 | 47 | -24 |
|  |  |  |  |  | -17 | -56 | 48 |
|  |  |  |  |  | 17 | -74 | 44 |
|  |  |  |  |  | -2 | -69 | 12 |
|  |  |  |  |  | -2 | -71 | -12 |
|  |  |  |  |  | 36 | -74 | -18 |
|  |  |  |  |  | -12 | 44 | 33 |
|  |  |  |  |  | -33 | -54 | 48 |
|  |  |  |  |  | 17 | -79 | 32 |
|  |  |  |  |  | -2 | -75 | 7 |
|  |  |  |  |  | 1 | -79 | 1 |
|  |  |  |  |  | -2 | -71 | -12 |
| Gennari et al | 2007 | 10.1016/j.neuroimage.2007.01.015 | Ambiguous > unambiguous | 17 | -52 | -61 | 5 |
|  |  |  |  |  | -44 | 21 | -4 |
|  |  |  |  |  | -35 | -43 | 48 |
|  |  |  |  |  | -50 | 4 | 13 |
| Gurd et al. | 2002 | 10.1093/brain/awf093 | Switching > not switching | 11 | -15 | -65 | 69 |
|  |  |  |  |  | 32 | -60 | 66 |
| Hirshorn & Thompson Schill | 2006 | 10.1016/j.neuropsychologia.2006.03.035 | Switching > free generation | 10 | -42 | 3 | 66 |
|  |  |  |  |  | -18 | 48 | -6 |
|  |  |  |  |  | -3 | -57 | 63 |
|  |  |  |  |  | -57 | -57 | 54 |
|  |  |  |  |  | -3 | -66 | 60 |
|  |  |  |  |  | -3 | -72 | 0 |
|  |  |  |  |  | 48 | 24 | 54 |
|  |  |  |  |  | 48 | 24 | 42 |
|  |  |  |  |  | 48 | 42 | 30 |
|  |  |  |  |  | 3 | 51 | -15 |
|  |  |  |  |  | 39 | 57 | 3 |
|  |  |  |  |  | 36 | 42 | -9 |
|  |  |  |  |  | 12 | -27 | -27 |
|  |  |  |  |  | 42 | -51 | 45 |
|  |  |  |  |  | -39 | 24 | 18 |
| Hirshorn & Thompson Schill | 2006 | 10.1016/j.neuropsychologia.2006.03.035 | Switching > clustering | 10 | -12 | 21 | 63 |
|  |  |  |  |  | -30 | 54 | 18 |
|  |  |  |  |  | -51 | 45 | 9 |
|  |  |  |  |  | -54 | 21 | 39 |
|  |  |  |  |  | -33 | 18 | 51 |
|  |  |  |  |  | -39 | -3 | 51 |
|  |  |  |  |  | 15 | 54 | -3 |
|  |  |  |  |  | -33 | 27 | 6 |
|  |  |  |  |  | -39 | -24 | 33 |
|  |  |  |  |  | -33 | -63 | 21 |
|  |  |  |  |  | -18 | -21 | 15 |
|  |  |  |  |  | -39 | -39 | 36 |
|  |  |  |  |  | -39 | -66 | 39 |
|  |  |  |  |  | -12 | -66 | 51 |
|  |  |  |  |  | 39 | 45 | 39 |
|  |  |  |  |  | 36 | 51 | 15 |
|  |  |  |  |  | 15 | 33 | 48 |
|  |  |  |  |  | 66 | -33 | 21 |
|  |  |  |  |  | 51 | -30 | -18 |
|  |  |  |  |  | 36 | -54 | 42 |
|  |  |  |  |  | 48 | -60 | -33 |
| Ketteler et al. | 2008 | 10.1016/j.neuroimage.2007.10.023 | Weak > strong target relation and strong > weak distractor relation | 12 | -12 | 48 | 38 |
|  |  |  |  |  | 21 | 45 | 16 |
|  |  |  |  |  | -48 | 39 | -27 |
|  |  |  |  |  | 53 | 19 | 4 |
|  |  |  |  |  | 43 | 39 | -28 |
|  |  |  |  |  | 50 | -51 | 43 |
|  |  |  |  |  | -52 | -37 | 58 |
|  |  |  |  |  | -31 | -16 | -16 |
|  |  |  |  |  | -21 | 41 | 49 |
|  |  |  |  |  | -2 | -15 | 10 |
|  |  |  |  |  | -47 | 36 | 22 |
|  |  |  |  |  | -66 | -43 | -6 |
|  |  |  |  |  | 50 | 33 | 18 |
|  |  |  |  |  | -2 | -39 | 2 |
|  |  |  |  |  | 41 | -60 | 44 |
|  |  |  |  |  | -2 | -29 | 6 |
| Nagel et al | 2008 | 10.1016/j.neuroimage.2008.07.017 | High > low selection | 14 | -51 | 26 | -4 |
|  |  |  |  |  | -46 | -39 | 46 |
|  |  |  |  |  | -44 | 36 | 18 |
| Nelson et al | 2009 | 10.1016/j.brainres.2008.12.001 | Many > few associates | 17 | -8 | 18 | 40 |
|  |  |  |  |  | 0 | 11 | 50 |
|  |  |  |  |  | -45 | 0 | 50 |
|  |  |  |  |  | -52 | 23 | 20 |
|  |  |  |  |  | -56 | 11 | 20 |
| Noppeney et al | 2004 | 10.1016/j.neuropsychologia.2003.12.014 | High > low semantic competition | 15 | -44 | 32 | 10 |
|  |  |  |  |  | 37 | -46 | -28 |
|  |  |  |  |  | -44 | -64 | -18 |
|  |  |  |  |  | -41 | -22 | -15 |
| Noppeney et al. | 2004 | 10.1016/j.neuroimage.2003.12.010 | Difficult > easy judgements | 15 | -32 | 20 | -13 |
|  |  |  |  |  | -5 | 32 | 46 |
|  |  |  |  |  | -9 | -5 | -1 |
|  |  |  |  |  | -44 | 14 | 25 |
|  |  |  |  |  | 1 | -64 | -49 |
| Persson et al. | 2004 | 10.1016/j.neuroimage.2004.08.004 | High > low selection | 22 | -49 | 26 | 15 |
|  |  |  |  |  | 41 | 15 | 5 |
|  |  |  |  |  | -52 | -52 | -5 |
|  |  |  |  |  | -4 | 8 | 60 |
| Race et al. | 2009 | 10.1162/jocn.2009.21132 | Different > same attribute | 26 | -51 | 36 | 12 |
|  |  |  |  |  | -48 | 33 | 0 |
|  |  |  |  |  | -45 | 15 | 24 |
|  |  |  |  |  | -51 | 12 | 15 |
| Race et al. | 2009 | 10.1162/jocn.2009.21132 | Reversed > repeated decision | 26 | -24 | 0 | 66 |
|  |  |  |  |  | -39 | -6 | 54 |
|  |  |  |  |  | -12 | 18 | 30 |
| Race et al. | 2009 | 10.1162/jocn.2009.21132 | Novel > repeated decision | 26 | -42 | 33 | -3 |
|  |  |  |  |  | -45 | 42 | -9 |
|  |  |  |  |  | -45 | 9 | 21 |
|  |  |  |  |  | -54 | -42 | 3 |
|  |  |  |  |  | -45 | -63 | -24 |
|  |  |  |  |  | -39 | -48 | -27 |
|  |  |  |  |  | -33 | -33 | -21 |
|  |  |  |  |  | -33 | -33 | -30 |
|  |  |  |  |  | -45 | -57 | -12 |
|  |  |  |  |  | -45 | -51 | -15 |
| Roskies et al. | 2001 | 10.1162/08989290152541485 | Low > high category typicality | 20 | -54 | 23 | -8 |
|  |  |  |  |  | 17 | -92 | -25 |
| Snyder et al. | 2007 | 10.1162/jocn.2007.19.5.761 | Specific attribute related > globally semantically related | 14 | -24 | -102 | -12 |
|  |  |  |  |  | -51 | -57 | -18 |
|  |  |  |  |  | -57 | -48 | -3 |
|  |  |  |  |  | -27 | -69 | 48 |
|  |  |  |  |  | -39 | -45 | 51 |
|  |  |  |  |  | 33 | -48 | 51 |
|  |  |  |  |  | -54 | 27 | 24 |
|  |  |  |  |  | -36 | 42 | 0 |
|  |  |  |  |  | -57 | 15 | 6 |
|  |  |  |  |  | 54 | 27 | 24 |
|  |  |  |  |  | -60 | 9 | 39 |
|  |  |  |  |  | 42 | 18 | 48 |
|  |  |  |  |  | 0 | 45 | 57 |
| Spalek et al. | 2008 | 10.1016/j.bandl.2008.05.005 | Semantically related > unrelated distractors | 21 | -35 | -47 | -16 |
|  |  |  |  |  | 17 | -86 | -25 |
| Thompson schill et al. | 1997 | 10.1073/pnas.94.26.14792 | High > low selection | 6 | -52 | 13 | 29 |
| Thompson schill et al. | 1997 | 10.1073/pnas.94.26.14792 | High > low selection of pictures | 6 | -3 | 18 | 44 |
|  |  |  |  |  | -40 | 20 | 28 |
|  |  |  |  |  | -40 | 13 | 33 |
|  |  |  |  |  | 46 | 21 | 31 |
|  |  |  |  |  | -3 | 19 | 53 |
|  |  |  |  |  | -52 | -55 | 2 |
|  |  |  |  |  | -40 | -64 | -14 |
|  |  |  |  |  | -31 | -71 | 41 |
| Thompson schill et al. | 1997 | 10.1073/pnas.94.26.14792 | High > low selection of words | 6 | -47 | 9 | 29 |
|  |  |  |  |  | 42 | 20 | 18 |
|  |  |  |  |  | -43 | 34 | 2 |
|  |  |  |  |  | -3 | 45 | 29 |
|  |  |  |  |  | -3 | 19 | 53 |
|  |  |  |  |  | -52 | -59 | -7 |
|  |  |  |  |  | -35 | -66 | 53 |
| Thompson Schill et al. | 1999 | 10.1016/S0896-6273(00)80804-1 | Different > same attribute | 8 | -46 | 20 | 19 |
| Wagner et al. | 2001 | 10.1016/S0896-6273(01)00359-2 | Weak > strong association | 14 | -45 | 27 | -12 |
|  |  |  |  |  | -51 | 18 | 27 |
|  |  |  |  |  | -51 | 21 | -3 |
|  |  |  |  |  | 6 | 18 | 39 |
|  |  |  |  |  | 9 | 27 | 36 |
|  |  |  |  |  | 0 | 9 | 57 |
|  |  |  |  |  | 45 | 21 | 6 |
|  |  |  |  |  | 30 | 24 | -6 |
|  |  |  |  |  | 39 | 27 | -9 |
|  |  |  |  |  | 54 | 24 | 27 |
| Wagner et al. | 2001 | 10.1016/S0896-6273(01)00359-2 | More > less foils | 14 | -36 | 21 | 27 |
|  |  |  |  |  | -39 | 6 | 24 |
|  |  |  |  |  | -45 | 27 | 9 |
|  |  |  |  |  | 3 | 30 | 36 |
|  |  |  |  |  | 3 | 15 | 42 |
|  |  |  |  |  | -51 | 21 | -12 |
| Whitney et al. | 2009 | 10.1093/cercor/bhp007 | Ambiguous homonym > unambiguous of doubly-related words | 18 | -44 | 20 | 28 |
|  |  |  |  |  | -48 | 24 | 32 |
| Whitney et al. | 2009 | 10.1093/cercor/bhp007 | Ambiguous homonym > unambiguous of single-related words | 18 | -48 | 28 | 16 |
|  |  |  |  |  | -44 | 12 | 24 |
| Whitney et al. | 2009 | 10.1093/cercor/bhp007 | Subordinate > dominant word ambiguity | 18 | -52 | 32 | -4 |
|  |  |  |  |  | -52 | 20 | 24 |
|  |  |  |  |  | -48 | 8 | 40 |
| Wig et al. | 2009 | 10.1152/jn.91213.2008 | Reversed > different decision | 27 | -44 | 11 | 23 |
|  |  |  |  |  | 55 | 9 | 24 |
| Zhang et al. | 2004 | 10.1016/j.neuroimage.2004.07.008 | High conflict > low conflict words | 14 | -25 | 23 | -21 |
|  |  |  |  |  | 39 | 20 | -22 |
|  |  |  |  |  | 6 | 37 | 17 |
| Zhang et al. | 2004 | 10.1016/j.neuroimage.2004.07.008 | High conflict > neutral words | 14 | -41 | 23 | -14 |
|  |  |  |  |  | -45 | 15 | 18 |
|  |  |  |  |  | 1 | 25 | 33 |
|  |  |  |  |  | 39 | 20 | -18 |
| Collette et al. | 2001 | 10.1006/nimg.2001.0846 | Inhibit>initiate | 12 | -33 | 21 | 27 |
|  |  |  |  |  | -40 | 48 | -13 |
|  |  |  |  |  | -55 | 24 | 14 |
|  |  |  |  |  | -46 | 25 | 18 |
|  |  |  |  |  | -33 | 26 | 29 |
|  |  |  |  |  | -46 | 27 | 18 |
|  |  |  |  |  | -55 | 27 | 23 |
|  |  |  |  |  | -57 | 36 | 22 |
|  |  |  |  |  | -42 | 44 | 12 |
|  |  |  |  |  | -49 | 45 | 3 |
|  |  |  |  |  | -45 | 50 | -13 |
|  |  |  |  |  | -38 | 56 | 2 |
|  |  |  |  |  | -36 | 63 | -6 |
|  |  |  |  |  | -32 | 70 | 0 |
|  |  |  |  |  | 42 | 72 | -15 |
|  |  |  |  |  | -47 | 27 | -4 |
|  |  |  |  |  | -53 | 38 | -3 |
|  |  |  |  |  | -38 | 28 | -16 |
|  |  |  |  |  | -53 | 33 | -9 |
|  |  |  |  |  | -47 | 45 | -17 |
|  |  |  |  |  | 66 | 39 | -21 |
|  |  |  |  |  | -30 | 55 | -32 |
| Mason & Just | 2007 | 10.1016/j.brainres.2007.02.076 | Ambiguous > unambiguous | 12 | -28 | 12 | 16 |
|  |  |  |  |  | 24 | 16 | 12 |
|  |  |  |  |  | 26 | 46 | 16 |
|  |  |  |  |  | -8 | 18 | 6 |
|  |  |  |  |  | 12 | 18 | 8 |
|  |  |  |  |  | -20 | 54 | 18 |
|  |  |  |  |  | 34 | 30 | 6 |
|  |  |  |  |  | 24 | 46 | 16 |
|  |  |  |  |  | -16 | 46 | 14 |
|  |  |  |  |  | -52 | 26 | 12 |
|  |  |  |  |  | 24 | 12 | 10 |
|  |  |  |  |  | -12 | 20 | 12 |
|  |  |  |  |  | -24 | 22 | 2 |
|  |  |  |  |  | -22 | -28 | 0 |
|  |  |  |  |  | 46 | 18 | 10 |
|  |  |  |  |  | -12 | -12 | 10 |
|  |  |  |  |  | -42 | 32 | 10 |
|  |  |  |  |  | -46 | 24 | 2 |
|  |  |  |  |  | -8 | 4 | 8 |
|  |  |  |  |  | -56 | 26 | 16 |
|  |  |  |  |  | -16 | 8 | 14 |
| Zempleni et al. | 2007 | 10.1016/j.neuroimage.2006.09.048 | Ambiguous > unambiguous | 16 | -48 | 26 | 20 |
|  |  |  |  |  | -52 | 16 | 26 |
|  |  |  |  |  | 34 | 20 | -10 |
|  |  |  |  |  | -50 | -48 | -12 |
|  |  |  |  |  | 56 | -34 | -16 |
| Liu et al. | 2009 | 10.1162/jocn.2009.21141 | Weak > strong association | 16 | -58 | 26 | 4 |
|  |  |  |  |  | -54 | 20 | 32 |
| Liu et al. | 2009 | 10.1162/jocn.2009.21141 | Weak > strong association | 16 | -58 | 20 | 0 |
|  |  |  |  |  | -52 | 18 | 28 |

*N.B. Contrasts within the same study conducted using the same sample were combined before they were entered into the meta-analysis.*

Supplementary Table 4. Data included in the n-back meta-analysis.

| **Author(s)** | **Year** | **DOI** | **Contrast included** | **Stimuli** | **Spatial/**  **Identity** | **N** | **MNI coordinates** | | |
| --- | --- | --- | --- | --- | --- | --- | --- | --- | --- |
|  |  |  |  |  |  |  | **X** | **Y** | **Z** |
| Garrett et al. | 2011 | 10.1176/appi.ajp.2010.09121718 | 1 > 0 back | Letters | Identity | 19 | 63 | 37 | 5 |
|  |  |  |  |  |  |  | 61 | 37 | 8 |
|  |  |  |  |  |  |  | 41 | 15 | 48 |
|  |  |  |  |  |  |  | -44 | -42 | 46 |
|  |  |  |  |  |  |  | -44 | -41 | 42 |
|  |  |  |  |  |  |  | 57 | -31 | 56 |
|  |  |  |  |  |  |  | 47 | -55 | 55 |
|  |  |  |  |  |  |  | 56 | -27 | 54 |
|  |  |  |  |  |  |  | -31 | 22 | 55 |
|  |  |  |  |  |  |  | -60 | 22 | 28 |
|  |  |  |  |  |  |  | 63 | 37 | 5 |
|  |  |  |  |  |  |  | -33 | 22 | 51 |
|  |  |  |  |  |  |  | -60 | 22 | 27 |
|  |  |  | 2 >0 back | Letters | Identity | 19 | 49 | 52 | 35 |
|  |  |  |  |  |  |  | 47 | 49 | 31 |
|  |  |  |  |  |  |  | 41 | 13 | 45 |
|  |  |  |  |  |  |  | 57 | 19 | 28 |
|  |  |  |  |  |  |  | 4 | 26 | 50 |
|  |  |  |  |  |  |  | 4 | 24 | 52 |
|  |  |  |  |  |  |  | 16 | 87 | -1 |
|  |  |  |  |  |  |  | 29 | 70 | -4 |
|  |  |  |  |  |  |  | 63 | 57 | 4 |
|  |  |  |  |  |  |  | 58 | 58 | 7 |
|  |  |  |  |  |  |  | 60 | 72 | -2 |
| Yan et al. | 2011 | 10.1016/j.bandc.2011.06.002 | 2 > 0 back | Shapes | Spatial | 28 (A) | 32 | 69 | 38 |
|  |  |  |  |  |  |  | -35 | 69 | 39 |
|  |  |  |  |  |  |  | -1 | 62 | 16 |
|  |  |  |  |  |  |  | -41 | 7 | 44 |
|  |  |  |  |  |  |  | 3 | -51 | 20 |
|  |  |  |  |  |  |  | 30 | 14 | 56 |
|  |  |  |  |  |  |  | -2 | -2 | 51 |
|  |  |  |  |  |  |  | -19 | 12 | 5 |
|  |  |  |  |  |  |  | 49 | 4 | 35 |
|  |  |  |  |  |  |  | 27 | 64 | -40 |
|  |  |  | 2 > 0 back | Shapes | Spatial | 28 (B) | -31 | 70 | 40 |
|  |  |  |  |  |  |  | 4 | -49 | 18 |
|  |  |  |  |  |  |  | -46 | 2 | 34 |
|  |  |  |  |  |  |  | 5 | 64 | 13 |
|  |  |  |  |  |  |  | 38 | 59 | 36 |
|  |  |  |  |  |  |  | 17 | 80 | 36 |
|  |  |  |  |  |  |  | 34 | 17 | 50 |
|  |  |  |  |  |  |  | -5 | -1 | 54 |
| Gropman et al. | 2013 | 10.1002/hbm.21470 | 2 > 1 back | Letters | Identity | 21 | 42 | -44 | 46 |
|  |  |  |  |  |  |  | -14 | -64 | 52 |
|  |  |  |  |  |  |  | -36 | -46 | 42 |
|  |  |  |  |  |  |  | 14 | -61 | 56 |
|  |  |  |  |  |  |  | -48 | -42 | 46 |
|  |  |  |  |  |  |  | -28 | -66 | 32 |
|  |  |  |  |  |  |  | -30 | 18 | 0 |
|  |  |  |  |  |  |  | 2 | 20 | 44 |
|  |  |  |  |  |  |  | -22 | 6 | 56 |
|  |  |  |  |  |  |  | -46 | 24 | 34 |
|  |  |  |  |  |  |  | -28 | 3 | 54 |
|  |  |  |  |  |  |  | -18 | 4 | 4 |
|  |  |  |  |  |  |  | -40 | 2 | 36 |
|  |  |  |  |  |  |  | -40 | 4 | 32 |
|  |  |  |  |  |  |  | -51 | 10 | 20 |
|  |  |  |  |  |  |  | -4 | 6 | 58 |
|  |  |  |  |  |  |  | 26 | 10 | 58 |
|  |  |  |  |  |  |  | 48 | 30 | 27 |
|  |  |  |  |  |  |  | 38 | 36 | 32 |
|  |  |  |  |  |  |  | 46 | 26 | 36 |
|  |  |  |  |  |  |  | 42 | 6 | 45 |
|  |  |  |  |  |  |  | 52 | 8 | 34 |
|  |  |  |  |  |  |  | 42 | 48 | 18 |
|  |  |  |  |  |  |  | -28 | 46 | -14 |
|  |  |  |  |  |  |  | -36 | 52 | -2 |
|  |  |  |  |  |  |  | 30 | 22 | -1 |
|  |  |  |  |  |  |  | 20 | 6 | 6 |
|  |  |  |  |  |  |  | -50 | -58 | -8 |
|  |  |  |  |  |  |  | 35 | -65 | -51 |
|  |  |  |  |  |  |  | 33 | -68 | -22 |
|  |  |  |  |  |  |  | 42 | -62 | -29 |
|  |  |  |  |  |  |  | -28 | -64 | -33 |
|  |  |  |  |  |  |  | -30 | -66 | -42 |
|  |  |  |  |  |  |  | 62 | -48 | -10 |
|  |  |  |  |  |  |  | 16 | 6 | 6 |
|  |  |  |  |  |  |  | 10 | 5 | -1 |
|  |  |  |  |  |  |  | 16 | 13 | 2 |
|  |  |  |  |  |  |  | -16 | 2 | 15 |
|  |  |  |  |  |  |  | -16 | -2 | 16 |
|  |  |  |  |  |  |  | -12 | 8 | 4 |
|  |  |  |  |  |  |  | -16 | 6 | 12 |
|  |  |  |  |  |  |  | -12 | 10 | 0 |
|  |  |  |  |  |  |  | -12 | 0 | 10 |
| Fernandez-Corcuera et al. | 2013 | 10.1016/j.jad.2012.04.009 | 2 > 0 back | Letters | Identity | 41 | -42 | 14 | 28 |
|  |  |  |  |  |  |  | 46 | -44 | 44 |
| Derrfuss et al. | 2004 | 10.1016/j.neuroimage.2004.06.007 | 2 > 0 back | Letters | Identity | 19 | -40 | 7 | 31 |
|  |  |  |  |  |  |  | 52 | 13 | 27 |
|  |  |  |  |  |  |  | -3 | 18 | 50 |
|  |  |  |  |  |  |  | -43 | 28 | 26 |
|  |  |  |  |  |  |  | -28 | 6 | 61 |
|  |  |  |  |  |  |  | -32 | 23 | 0 |
|  |  |  |  |  |  |  | 43 | -41 | 47 |
|  |  |  |  |  |  |  | -2 | -59 | 54 |
|  |  |  |  |  |  |  | -11 | -24 | 15 |
| Kim et al. | 2006 | 10.1016/j.neuroimage.2005.11.035 | 2 back > rest | Letters | Identity | 12 | -52 | -30 | 44 |
|  |  |  |  |  |  |  | -34 | -48 | 47 |
|  |  |  |  |  |  |  | -53 | 9 | 29 |
|  |  |  |  |  |  |  | -41 | 28 | 24 |
|  |  |  |  |  |  |  | -5 | 24 | 41 |
|  |  |  |  |  |  |  | 11 | 20 | 36 |
|  |  |  |  |  |  |  | -1 | 10 | 56 |
|  |  |  |  |  |  |  | -51 | -68 | -5 |
| Honey et al. | 1999 | 10.1073/pnas.96.23.13432 | 2 > 0 back | Letters | Identity | 10 | -39 | -51 | 43 |
|  |  |  |  |  |  |  | 36 | -63 | 42 |
|  |  |  |  |  |  |  | -45 | 8 | 37 |
|  |  |  |  |  |  |  | 39 | 50 | 7 |
|  |  |  |  |  |  |  | -37 | 22 | 4 |
|  |  |  |  |  |  |  | -2 | 15 | 48 |
|  |  |  |  |  |  |  | -45 | 9 | 42 |
|  |  |  |  |  |  |  | -41 | -9 | 51 |
|  |  |  |  |  |  |  | 36 | 1 | 54 |
|  |  |  |  |  |  |  | -42 | -73 | -11 |
|  |  |  |  |  |  |  | 20 | -80 | -12 |
|  |  |  |  |  |  |  | -34 | -72 | -17 |
| Rodriguez-Jimenez et al. | 2009 | 10.1016/j.bbr.2009.08.022 | 2 > 0 back | Letters (audio and visual) | Identity | 13 | -1 | 13 | 55 |
|  |  |  |  |  |  |  | -1 | 8 | 62 |
|  |  |  |  |  |  |  | 8 | 35 | 39 |
|  |  |  |  |  |  |  | -54 | 8 | 36 |
|  |  |  |  |  |  |  | -47 | 33 | 27 |
|  |  |  |  |  |  |  | -44 | 11 | 29 |
|  |  |  |  |  |  |  | -27 | 4 | 59 |
|  |  |  |  |  |  |  | -24 | 0 | 50 |
|  |  |  |  |  |  |  | -44 | -43 | 44 |
|  |  |  |  |  |  |  | -27 | -68 | 46 |
|  |  |  |  |  |  |  | -27 | -52 | 45 |
|  |  |  |  |  |  |  | 1 | -54 | -13 |
|  |  |  |  |  |  |  | 43 | 43 | 24 |
|  |  |  |  |  |  |  | 63 | 21 | 29 |
|  |  |  |  |  |  |  | 50 | 27 | 22 |
|  |  |  |  |  |  |  | 37 | 10 | 54 |
|  |  |  |  |  |  |  | 28 | 7 | 55 |
|  |  |  |  |  |  |  | 18 | -63 | 59 |
|  |  |  |  |  |  |  | 47 | -41 | 56 |
|  |  |  |  |  |  |  | 50 | -33 | 41 |
| Carlson et al. | 1998 | 10.1093/cercor/8.8.743 | 2 > 0 back | Shapes | Spatial | 7 | 42 | 29 | 27 |
|  |  |  |  |  |  |  | -39 | 39 | 28 |
|  |  |  |  |  |  |  | 21 | 11 | 59 |
|  |  |  |  |  |  |  | -23 | 22 | 51 |
|  |  |  |  |  |  |  | 4 | 18 | 54 |
|  |  |  |  |  |  |  | -8 | 18 | 55 |
|  |  |  |  |  |  |  | 41 | 20 | 25 |
|  |  |  |  |  |  |  | -45 | 22 | 24 |
|  |  |  |  |  |  |  | 38 | 59 | 0 |
|  |  |  |  |  |  |  | -27 | 55 | 3 |
|  |  |  |  |  |  |  | 42 | 0 | 50 |
|  |  |  |  |  |  |  | -46 | 1 | 41 |
|  |  |  |  |  |  |  | 5 | -66 | 53 |
|  |  |  |  |  |  |  | -2 | -62 | 52 |
|  |  |  |  |  |  |  | 11 | -53 | 73 |
|  |  |  |  |  |  |  | -12 | -60 | 65 |
|  |  |  |  |  |  |  | 33 | -57 | 51 |
|  |  |  |  |  |  |  | -36 | -56 | 52 |
|  |  |  |  |  |  |  | 46 | -49 | 36 |
|  |  |  |  |  |  |  | -47 | -52 | 46 |
|  |  |  |  |  |  |  | 29 | -74 | 33 |
|  |  |  |  |  |  |  | -22 | -76 | 28 |
|  |  |  |  |  |  |  | 4 | 21 | 38 |
|  |  |  |  |  |  |  | -2 | 18 | 43 |
|  |  |  |  |  |  |  | 8 | -50 | 33 |
|  |  |  |  |  |  |  | -4 | -49 | 26 |
|  |  |  |  |  |  |  | 38 | 20 | -4 |
|  |  |  |  |  |  |  | -38 | 14 | 1 |
|  |  |  | 1 > 0 back | Shapes | Spatial | 7 | 17 | 12 | 57 |
|  |  |  |  |  |  |  | -33 | 4 | 65 |
|  |  |  |  |  |  |  | -7 | 2 | 57 |
|  |  |  |  |  |  |  | -40 | 37 | -15 |
|  |  |  |  |  |  |  | 14 | 54 | -19 |
|  |  |  |  |  |  |  | -27 | 59 | -7 |
|  |  |  |  |  |  |  | 4 | -48 | 61 |
|  |  |  |  |  |  |  | -6 | -54 | 61 |
|  |  |  |  |  |  |  | 22 | -63 | 64 |
|  |  |  |  |  |  |  | -14 | -61 | 49 |
|  |  |  |  |  |  |  | -32 | -57 | 50 |
|  |  |  |  |  |  |  | -57 | -40 | 58 |
|  |  |  |  |  |  |  | 31 | -86 | 1 |
|  |  |  |  |  |  |  | -14 | -81 | 16 |
|  |  |  |  |  |  |  | 9 | 35 | -13 |
|  |  |  |  |  |  |  | -8 | 57 | -12 |
|  |  |  |  |  |  |  | -7 | -62 | 33 |
|  |  |  |  |  |  |  | 44 | 40 | 14 |
|  |  |  |  |  |  |  | -35 | 47 | 27 |
|  |  |  |  |  |  |  | 28 | 20 | 49 |
|  |  |  |  |  |  |  | -26 | 14 | 52 |
|  |  |  |  |  |  |  | 14 | 18 | 56 |
|  |  |  |  |  |  |  | -10 | 31 | 52 |
|  |  |  |  |  |  |  | 58 | 28 | -3 |
|  |  |  |  |  |  |  | -53 | 23 | 21 |
|  |  |  |  |  |  |  | 14 | 59 | -10 |
|  |  |  |  |  |  |  | -13 | 63 | 8 |
|  |  |  |  |  |  |  | 56 | 6 | 45 |
|  |  |  |  |  |  |  | -37 | -2 | 52 |
|  |  |  |  |  |  |  | 11 | -59 | 51 |
|  |  |  |  |  |  |  | -7 | -56 | 60 |
|  |  |  |  |  |  |  | 28 | -56 | 58 |
|  |  |  |  |  |  |  | -18 | -58 | 63 |
|  |  |  |  |  |  |  | 33 | -55 | 41 |
|  |  |  |  |  |  |  | -38 | -59 | 44 |
|  |  |  |  |  |  |  | 47 | -46 | 41 |
|  |  |  |  |  |  |  | -45 | -54 | 40 |
|  |  |  |  |  |  |  | 38 | -79 | 16 |
|  |  |  |  |  |  |  | -27 | -80 | 30 |
|  |  |  |  |  |  |  | 7 | 35 | 20 |
|  |  |  |  |  |  |  | -3 | 11 | 43 |
|  |  |  |  |  |  |  | 31 | 15 | 5 |
|  |  |  |  |  |  |  | -47 | 15 | -4 |
| Dima et al. | 2014 | 10.1002/hbm.22382 | 1 > 0 back | Letters | Identity | 40 | 46 | -46 | 44 |
|  |  |  |  |  |  |  | -44 | -42 | 38 |
|  |  |  |  |  |  |  | -46 | 32 | 30 |
|  |  |  |  |  |  |  | 52 | 36 | 30 |
|  |  |  |  |  |  |  | 32 | 6 | 62 |
|  |  |  | 2 > 0 back | Letters | Identity | 40 | 40 | -48 | 44 |
|  |  |  |  |  |  |  | -36 | -52 | 46 |
|  |  |  |  |  |  |  | 46 | 32 | 28 |
|  |  |  |  |  |  |  | -42 | 8 | 28 |
|  |  |  |  |  |  |  | 8 | 18 | 48 |
|  |  |  |  |  |  |  | -4 | 10 | 58 |
|  |  |  |  |  |  |  | 34 | 24 | -2 |
|  |  |  | 3 > 0 back | Letters | Identity | 40 | 30 | 8 | 58 |
|  |  |  |  |  |  |  | 50 | -42 | 42 |
|  |  |  |  |  |  |  | -48 | -48 | 48 |
|  |  |  |  |  |  |  | -48 | 26 | 30 |
|  |  |  |  |  |  |  | 48 | 40 | 30 |
|  |  |  |  |  |  |  | -10 | 26 | 30 |
|  |  |  |  |  |  |  | 8 | 20 | 28 |
|  |  |  |  |  |  |  | -34 | 22 | 0 |
|  |  |  |  |  |  |  | -4 | 10 | 58 |
|  |  |  |  |  |  |  | -12 | -10 | 6 |
|  |  |  |  |  |  |  | -44 | 46 | 2 |
|  |  |  |  |  |  |  | 34 | 20 | 0 |
| Mendrek et al. | 2005 | 10.1017/s0033291704003228 | 2 > 0 back | Letters | Identity | 12 | -46 | 1 | 36 |
|  |  |  |  |  |  |  | 49 | 34 | 18 |
|  |  |  |  |  |  |  | 36 | 4 | 62 |
|  |  |  |  |  |  |  | -42 | 54 | 13 |
|  |  |  |  |  |  |  | 45 | -58 | 5 |
|  |  |  |  |  |  |  | 41 | -67 | 46 |
|  |  |  |  |  |  |  | -33 | -71 | 52 |
|  |  |  |  |  |  |  | -41 | -49 | 50 |
|  |  |  |  |  |  |  | -38 | -67 | -42 |
|  |  |  |  |  |  |  | 44 | -70 | -39 |
|  |  |  |  |  |  |  | 6 | -14 | 10 |
|  |  |  |  |  |  |  | -7 | -19 | 11 |
| Barnes et al. | 2009 | 10.1371/journal.pone.0006626 | 1 and 2 > 0 back | Numbers | Identity and Spatial | 14 | -16 | 12 | -2 |
|  |  |  |  |  |  |  | 18 | 16 | 2 |
|  |  |  |  |  |  |  | 36 | -42 | 42 |
|  |  |  |  |  |  |  | -46 | -46 | 16 |
|  |  |  |  |  |  |  | 56 | 50 | 20 |
|  |  |  |  |  |  |  | -32 | -46 | 38 |
|  |  |  |  |  |  |  | 36 | -42 | 42 |
|  |  |  |  |  |  |  | 28 | 6 | 56 |
|  |  |  |  |  |  |  | -44 | 30 | 26 |
|  |  |  |  |  |  |  | 42 | 42 | 24 |
|  |  |  |  |  |  |  | 0 | 14 | 46 |
| Townsend et al. | 2010 | 10.1016/j.pscychresns.2009.11.010 | 2 > 0 back | Letters | Identity | 14 | 47 | 39 | 24 |
|  |  |  |  |  |  |  | 53 | 18 | 29 |
|  |  |  |  |  |  |  | -44 | 33 | 18 |
|  |  |  |  |  |  |  | -57 | 21 | 23 |
|  |  |  |  |  |  |  | 38 | 27 | -12 |
|  |  |  |  |  |  |  | -51 | 14 | 20 |
|  |  |  |  |  |  |  | -35 | 13 | 26 |
|  |  |  |  |  |  |  | 62 | -51 | 31 |
|  |  |  |  |  |  |  | 34 | -56 | 45 |
|  |  |  |  |  |  |  | 43 | -43 | 42 |
|  |  |  |  |  |  |  | -59 | -55 | 33 |
|  |  |  |  |  |  |  | 49 | -48 | 42 |
|  |  |  |  |  |  |  | -61 | -58 | 27 |
|  |  |  |  |  |  |  | 28 | -73 | 45 |
|  |  |  |  |  |  |  | -29 | -75 | 30 |
| Lythe et al. | 2012 | 10.1016/j.bbr.2012.04.050 | 1,2,3 back > 0 back | Letters | Identity | 20 | 42 | 30 | 36 |
|  |  |  |  |  |  |  | 32 | 2 | 62 |
|  |  |  |  |  |  |  | -30 | -2 | 44 |
|  |  |  |  |  |  |  | 32 | -56 | 38 |
|  |  |  |  |  |  |  | -28 | -58 | 44 |
|  |  |  | 3 > 2 > 1 > 0 back | Letters | Identity | 20 | 30 | 2 | 58 |
|  |  |  |  |  |  |  | 36 | -50 | 42 |
| Ravizza et al. | 2004 | 10.1016/j.neuroimage.2004.01.039 | 3 > 0 back | Letters | Identity | 20 | 8 | 24 | 44 |
|  |  |  |  |  |  |  | -23 | 4 | 51 |
|  |  |  |  |  |  |  | 31 | 5 | 45 |
|  |  |  |  |  |  |  | 52 | 3 | 36 |
|  |  |  |  |  |  |  | -31 | 53 | 10 |
|  |  |  |  |  |  |  | 48 | 26 | 37 |
|  |  |  |  |  |  |  | 38 | 38 | 16 |
|  |  |  |  |  |  |  | -43 | 23 | 34 |
|  |  |  |  |  |  |  | 47 | -69 | 50 |
|  |  |  |  |  |  |  | -49 | -53 | 51 |
|  |  |  |  |  |  |  | 52 | -45 | 46 |
|  |  |  |  |  |  |  | -24 | -63 | -30 |
|  |  |  |  |  |  |  | -33 | -45 | -34 |
|  |  |  |  |  |  |  | 33 | -66 | -30 |
| Lamp et al. | 2006 | 10.3389/fnbeh.2016.00087 | 1 > 0 back | Shapes | Identity | 16 | 34 | -64 | 50 |
|  |  |  |  |  |  |  | 38 | -44 | 40 |
|  |  |  |  |  |  |  | 34 | -56 | 44 |
|  |  |  |  |  |  |  | 32 | -48 | 42 |
|  |  |  |  |  |  |  | 38 | -52 | 42 |
|  |  |  |  |  |  |  | 12 | 14 | 46 |
|  |  |  |  |  |  |  | 14 | 4 | 52 |
|  |  |  |  |  |  |  | -4 | 6 | 58 |
|  |  |  |  |  |  |  | -12 | 18 | 42 |
|  |  |  |  |  |  |  | -2 | 16 | 54 |
|  |  |  |  |  |  |  | -8 | 16 | 44 |
|  |  |  |  |  |  |  | 26 | -8 | 58 |
|  |  |  |  |  |  |  | 22 | -8 | 52 |
|  |  |  |  |  |  |  | 24 | -10 | 54 |
|  |  |  |  |  |  |  | 28 | -10 | 48 |
|  |  |  |  |  |  |  | 28 | -18 | 48 |
|  |  |  |  |  |  |  | 32 | -16 | 54 |
| Schmidt et al. | 2015 | 10.3389/fneur.2015.00199 | 3 > 0 back | Letters | Identity | 27 | -48 | 10 | 32 |
|  |  |  |  |  |  |  | 44 | 35 | 24 |
|  |  |  |  |  |  |  | 32 | 5 | 52 |
|  |  |  |  |  |  |  | -36 | 52 | 8 |
|  |  |  |  |  |  |  | -24 | 46 | 6 |
|  |  |  |  |  |  |  | -5 | 24 | 46 |
|  |  |  |  |  |  |  | 52 | -48 | 50 |
|  |  |  |  |  |  |  | -30 | -58 | 50 |
|  |  |  |  |  |  |  | 34 | -58 | 48 |
|  |  |  |  |  |  |  | 10 | -72 | 54 |
|  |  |  |  |  |  |  | 32 | 24 | -4 |
|  |  |  |  |  |  |  | -28 | 22 | 2 |
|  |  |  |  |  |  |  | 34 | -62 | -36 |
|  |  |  |  |  |  |  | -28 | -64 | -36 |
|  |  |  |  |  |  |  | 10 | -18 | 10 |
|  |  |  |  |  |  |  | 16 | 0 | 12 |
|  |  |  |  |  |  |  | -10 | -16 | 10 |
|  |  |  | 2 > 0 back | Letters | Identity | 27 | -44 | 6 | 32 |
|  |  |  |  |  |  |  | 30 | 8 | 54 |
|  |  |  |  |  |  |  | -34 | 54 | 22 |
|  |  |  |  |  |  |  | -8 | 20 | 50 |
|  |  |  |  |  |  |  | -30 | -58 | 50 |
|  |  |  |  |  |  |  | 32 | 24 | -4 |
|  |  |  |  |  |  |  | -30 | 22 | 4 |
|  |  |  |  |  |  |  | 58 | -50 | -14 |
|  |  |  |  |  |  |  | 44 | -66 | -32 |
|  |  |  |  |  |  |  | -28 | -64 | -36 |
|  |  |  |  |  |  |  | -10 | -16 | 12 |
|  |  |  |  |  |  |  | 10 | -16 | 10 |
|  |  |  | 3 > 2 back | Letters | Identity |  | 42 | 32 | 38 |
| Belayachi et al. | 2015 | 10.1016/j.bbr.2015.07.042 | 3 > 0 back | Letters | Identity | 18 | -2 | 12 | 48 |
|  |  |  |  |  |  |  | -36 | 0 | 34 |
|  |  |  |  |  |  |  | -24 | -2 | 50 |
|  |  |  |  |  |  |  | -24 | 24 | 0 |
|  |  |  |  |  |  |  | 24 | 0 | 48 |
|  |  |  |  |  |  |  | 30 | 6 | 62 |
|  |  |  |  |  |  |  | 36 | 34 | 22 |
|  |  |  |  |  |  |  | -38 | -38 | 42 |
|  |  |  |  |  |  |  | -24 | -50 | 42 |
|  |  |  |  |  |  |  | 36 | -34 | 38 |
| El-Hage et al. | 2013 | 10.1038/mp.2011.145 | 3 > 2 > 1 > 0 back | Letters | Identity | 90 | 0 | 18 | 43 |
|  |  |  |  |  |  |  | 40 | -40 | 37 |
|  |  |  |  |  |  |  | 25 | 11 | 52 |
|  |  |  |  |  |  |  | -14 | 4 | 10 |
|  |  |  |  |  |  |  | 10 | -4 | 10 |
| Nagel et al. | 2011 | 10.1162/jocn.2010.21560 | 3 > 1 back | Letters | Identity | 30 | 8 | 22 | 42 |
|  |  |  |  |  |  |  | 28 | -68 | 52 |
|  |  |  |  |  |  |  | 36 | 24 | -2 |
|  |  |  |  |  |  |  | 4 | -82 | -18 |
|  |  |  |  |  |  |  | -56 | -50 | -10 |
| Savini et al. | 2012 | 10.1016/j.ijpsycho.2012.09.007 | 0,1,2 back > rest | Shapes (tactile) | Identity | 12 | -41 | -33 | 55 |
|  |  |  |  |  |  |  | -58 | -16 | 18 |
|  |  |  |  |  |  |  | 57 | -17 | 27 |
|  |  |  |  |  |  |  | 43 | -24 | 53 |
|  |  |  |  |  |  |  | 6 | 27 | 34 |
|  |  |  |  |  |  |  | 0 | 4 | 61 |
|  |  |  |  |  |  |  | -47 | 9 | 29 |
|  |  |  |  |  |  |  | 48 | 7 | 34 |
|  |  |  |  |  |  |  | 47 | 34 | 27 |
|  |  |  |  |  |  |  | 27 | 39 | 25 |
|  |  |  |  |  |  |  | -30 | 48 | 25 |
|  |  |  |  |  |  |  | 53 | -31 | -7 |
|  |  |  |  |  |  |  | 56 | -37 | 28 |
|  |  |  |  |  |  |  | -31 | -5 | 56 |
|  |  |  |  |  |  |  | 30 | 17 | 49 |
|  |  |  |  |  |  |  | 35 | -5 | 56 |
|  |  |  |  |  |  |  | -42 | -2 | 9 |
|  |  |  |  |  |  |  | -37 | 22 | -1 |
|  |  |  |  |  |  |  | 39 | 20 | -3 |
|  |  |  |  |  |  |  | -44 | 19 | 3 |
|  |  |  |  |  |  |  | 45 | -35 | 49 |
|  |  |  |  |  |  |  | -44 | -34 | 48 |
|  |  |  |  |  |  |  | 9 | -61 | 66 |
|  |  |  |  |  |  |  | -28 | -61 | 56 |
| Sabri et al. | 2014 | 10.1016/j.neuropsychologia.2014.06.009 | 2 > 1 back | Tones (auditory) | Identity | 13 | 42 | -46 | 45 |
|  |  |  |  |  |  |  | -5 | 23 | 43 |
|  |  |  |  |  |  |  | -29 | 24 | -3 |
|  |  |  |  |  |  |  | 70 | -42 | -12 |
|  |  |  |  |  |  |  | -24 | -59 | 40 |
|  |  |  |  |  |  |  | -30 | 59 | 9 |
|  |  |  |  |  |  |  | 34 | -66 | 51 |
|  |  |  |  |  |  |  | -10 | -68 | 54 |
|  |  |  |  |  |  |  | -46 | -39 | 47 |
|  |  |  |  |  |  |  | -41 | 16 | 28 |
|  |  |  |  |  |  |  | 11 | -68 | 46 |
|  |  |  |  |  |  |  | 51 | 22 | 36 |
|  |  |  |  |  |  |  | 26 | 12 | 58 |
|  |  |  |  |  |  |  | 41 | 46 | 17 |
|  |  |  |  |  |  |  | 33 | -64 | 46 |
|  |  |  |  |  |  |  | -36 | 53 | 10 |
| Valera et al. | 2005 | 10.1016/j.biopsych.2004.11.034 | 2 > 0 back | Letters | Identity | 20 | -33 | 27 | -6 |
|  |  |  |  |  |  |  | -27 | -69 | 42 |
|  |  |  |  |  |  |  | 9 | -75 | -24 |
|  |  |  |  |  |  |  | 15 | -81 | 12 |
|  |  |  |  |  |  |  | 57 | -54 | -18 |
| Schneiders et al. | 2011 | 10.1093/cercor/bhr037 | 2 > 0 back | Patterns | Identity | 48 | -40 | -43 | 45 |
|  |  |  |  |  |  |  | 44 | -40 | 43 |
|  |  |  |  |  |  |  | -34 | -57 | 43 |
|  |  |  |  |  |  |  | 31 | -55 | 45 |
|  |  |  |  |  |  |  | -24 | -66 | 29 |
|  |  |  |  |  |  |  | 31 | -61 | 40 |
|  |  |  |  |  |  |  | -11 | -71 | 61 |
|  |  |  |  |  |  |  | 28 | 4 | 63 |
|  |  |  |  |  |  |  | 47 | 32 | 26 |
|  |  |  |  |  |  |  | 47 | 45 | 22 |
|  |  |  |  |  |  |  | 53 | 15 | 41 |
|  |  |  |  |  |  |  | 37 | 20 | 1 |
|  |  |  |  |  |  |  | -27 | 0 | 61 |
|  |  |  |  |  |  |  | -2 | 15 | 49 |
|  |  |  |  |  |  |  | -47 | 7 | 34 |
|  |  |  |  |  |  |  | -41 | 50 | 12 |
|  |  |  |  |  |  |  | -35 | 17 | 2 |
|  |  |  |  |  |  |  | -41 | -64 | -33 |
|  |  |  |  |  |  |  | 30 | -63 | -31 |
|  |  |  |  |  |  |  | 11 | -2 | 3 |
|  |  |  |  |  |  |  | -9 | 7 | 3 |
|  |  |  |  |  |  |  | -51 | -62 | -10 |
| Seo et al. | 2012 | 10.1371/journal.pone.0037808 | 2 > 0 back | Letters | Identity | 22 | -42 | 9 | 30 |
|  |  |  |  |  |  |  | 30 | 9 | 57 |
|  |  |  |  |  |  |  | -33 | 24 | -6 |
|  |  |  |  |  |  |  | 33 | 26 | -6 |
|  |  |  |  |  |  |  | -6 | 24 | 45 |
|  |  |  |  |  |  |  | 9 | 27 | 45 |
|  |  |  |  |  |  |  | -3 | 15 | 54 |
|  |  |  |  |  |  |  | 6 | 18 | 54 |
|  |  |  |  |  |  |  | -15 | 3 | 0 |
|  |  |  |  |  |  |  | 15 | 3 | 0 |
|  |  |  |  |  |  |  | -12 | 9 | 0 |
|  |  |  |  |  |  |  | 15 | 9 | 6 |
|  |  |  |  |  |  |  | -54 | -51 | -15 |
|  |  |  |  |  |  |  | -27 | -66 | 45 |
|  |  |  |  |  |  |  | 39 | -66 | 51 |
|  |  |  |  |  |  |  | -39 | -60 | 48 |
|  |  |  |  |  |  |  | 42 | -45 | 42 |
|  |  |  |  |  |  |  | -9 | -78 | -34 |
|  |  |  |  |  |  |  | 35 | -63 | -39 |
| Caldu et al. | 2007 | 10.1016/j.neuroimage.2007.06.021 | 2 > 0 back | Numbers | Identity | 75 | 42 | -44 | 46 |
|  |  |  |  |  |  |  | 32 | -64 | 50 |
|  |  |  |  |  |  |  | -38 | -52 | 50 |
|  |  |  |  |  |  |  | 30 | -2 | 56 |
|  |  |  |  |  |  |  | -24 | -2 | 60 |
|  |  |  |  |  |  |  | 42 | 32 | 32 |
|  |  |  |  |  |  |  | -46 | 24 | 36 |
|  |  |  |  |  |  |  | 36 | 52 | 22 |
|  |  |  |  |  |  |  | -36 | 50 | 14 |
|  |  |  |  |  |  |  | 6 | 22 | 48 |
|  |  |  |  |  |  |  | 34 | 24 | 0 |
|  |  |  |  |  |  |  | -32 | 22 | 0 |
|  |  |  |  |  |  |  | 52 | 10 | 30 |
|  |  |  |  |  |  |  | -2 | 12 | 54 |
|  |  |  |  |  |  |  | 58 | -38 | -16 |
|  |  |  |  |  |  |  | 58 | -44 | -18 |
|  |  |  |  |  |  |  | -54 | -46 | -18 |
|  |  |  |  |  |  |  | 30 | -64 | -34 |
|  |  |  |  |  |  |  | -10 | -74 | -28 |
|  |  |  |  |  |  |  | 6 | -28 | 12 |
|  |  |  |  |  |  |  | -12 | -10 | 6 |
|  |  |  |  |  |  |  | 14 | -4 | 6 |
|  |  |  |  |  |  |  | -14 | 2 | 6 |
|  |  |  |  |  |  |  | 16 | 4 | 20 |
|  |  |  |  |  |  |  | -18 | 2 | 22 |
| Drobyshevsky et al. | 2006 | 10.1016/j.neuroimage.2005.12.016 | 2 > 0 back | Letters | Identity | 22 | -42 | 27 | 28 |
|  |  |  |  |  |  |  | 47 | 35 | 30 |
|  |  |  |  |  |  |  | -49 | 15 | 9 |
|  |  |  |  |  |  |  | 52 | 17 | 6 |
|  |  |  |  |  |  |  | -33 | 3 | 51 |
|  |  |  |  |  |  |  | 39 | 6 | 54 |
|  |  |  |  |  |  |  | -30 | 58 | 8 |
|  |  |  |  |  |  |  | 38 | 64 | 4 |
|  |  |  |  |  |  |  | 3 | 12 | 52 |
|  |  |  |  |  |  |  | -24 | -61 | 49 |
|  |  |  |  |  |  |  | 22 | -66 | 51 |
|  |  |  |  |  |  |  | -38 | -52 | 43 |
|  |  |  |  |  |  |  | 44 | -50 | 44 |
| Cohen et al. | 1997 | 10.1038/386604a0 | 3 > 2 > 1 > 0 back | Letters | Identity | 10 | 41 | 39 | 24 |
|  |  |  |  |  |  |  | 9 | 36 | 33 |
|  |  |  |  |  |  |  | -43 | 11 | 13 |
|  |  |  |  |  |  |  | 50 | 18 | 22 |
|  |  |  |  |  |  |  | -36 | 8 | 40 |
|  |  |  |  |  |  |  | 36 | 14 | 58 |
|  |  |  |  |  |  |  | 15 | -58 | 60 |
|  |  |  |  |  |  |  | -41 | -46 | 45 |
|  |  |  |  |  |  |  | 49 | -52 | 48 |
| Braver et al. | 1997 | 10.1006/nimg.1996.0247 | 3 > 2 > 1 > 0 back | Letters | Identity | 8 | 16 | 16 | 54 |
|  |  |  |  |  |  |  | -44 | 30 | 37 |
|  |  |  |  |  |  |  | 41 | 44 | 27 |
|  |  |  |  |  |  |  | -50 | 9 | 12 |
|  |  |  |  |  |  |  | 48 | 11 | 28 |
|  |  |  |  |  |  |  | -34 | 23 | 3 |
|  |  |  |  |  |  |  | -41 | 3 | 28 |
|  |  |  |  |  |  |  | -49 | 4 | 44 |
|  |  |  |  |  |  |  | -31 | 1 | 60 |
|  |  |  |  |  |  |  | -26 | -58 | 52 |
|  |  |  |  |  |  |  | 36 | -54 | 48 |
|  |  |  |  |  |  |  | 21 | 5 | 20 |
| Callicott et al. | 1999 | 10.1093/cercor/9.1.20 | 3 > 2 > 1 > 0 back | Letters | Identity | 9 | -44 | 31 | 16 |
|  |  |  |  |  |  |  | -38 | 35 | 11 |
|  |  |  |  |  |  |  | -32 | 35 | -8 |
|  |  |  |  |  |  |  | -29 | 21 | 48 |
|  |  |  |  |  |  |  | -20 | 16 | 57 |
|  |  |  |  |  |  |  | -33 | 11 | 53 |
|  |  |  |  |  |  |  | 56 | 16 | 33 |
|  |  |  |  |  |  |  | 53 | 29 | 14 |
|  |  |  |  |  |  |  | 36 | 7 | 57 |
|  |  |  |  |  |  |  | 11 | 6 | 66 |
|  |  |  |  |  |  |  | 6 | 0 | 72 |
|  |  |  |  |  |  |  | 2 | 8 | 66 |
|  |  |  |  |  |  |  | -16 | -2 | 9 |
|  |  |  |  |  |  |  | -1 | -16 | 11 |
|  |  |  |  |  |  |  | -8 | 4 | 4 |
|  |  |  |  |  |  |  | -46 | -25 | 35 |
|  |  |  |  |  |  |  | -2 | -56 | 68 |
|  |  |  |  |  |  |  | 41 | -55 | 58 |
| LaBar et al. | 1999 | 10.1006/nimg.1999.0503 | 2 > 0 back | Letters | Identity | 11 | 28 | -56 | 68 |
|  |  |  |  |  |  |  | 51 | -41 | 56 |
|  |  |  |  |  |  |  | -21 | -57 | 55 |
|  |  |  |  |  |  |  | -44 | -45 | 51 |
|  |  |  |  |  |  |  | 60 | 18 | 26 |
|  |  |  |  |  |  |  | -57 | 20 | 25 |
|  |  |  |  |  |  |  | -44 | 8 | 32 |
|  |  |  |  |  |  |  | 2 | 19 | 44 |
|  |  |  |  |  |  |  | -27 | 3 | 46 |
|  |  |  |  |  |  |  | 41 | 13 | 47 |
|  |  |  |  |  |  |  | 57 | 9 | 41 |
|  |  |  |  |  |  |  | -12 | -7 | 13 |
|  |  |  |  |  |  |  | -9 | -1 | 6 |
|  |  |  |  |  |  |  | 17 | 0 | 12 |
|  |  |  |  |  |  |  | -32 | -73 | -41 |
|  |  |  |  |  |  |  | -41 | -63 | -41 |
|  |  |  |  |  |  |  | 53 | -62 | -36 |
|  |  |  |  |  |  |  | 40 | -72 | -38 |
|  |  |  |  |  |  |  | 40 | -82 | -38 |
|  |  |  |  |  |  |  | 4 | -55 | -26 |
|  |  |  |  |  |  |  | -51 | -66 | -4 |
|  |  |  |  |  |  |  | -48 | -57 | -12 |
|  |  |  |  |  |  |  | -64 | -43 | 7 |
|  |  |  |  |  |  |  | -57 | -60 | -4 |
|  |  |  |  |  |  |  | 46 | 23 | -14 |
|  |  |  |  |  |  |  | -57 | 17 | -12 |
|  |  |  |  |  |  |  | 37 | 23 | 46 |
|  |  |  |  |  |  |  | 34 | 14 | 54 |
|  |  |  |  |  |  |  | -27 | 17 | 55 |
|  |  |  |  |  |  |  | -21 | 26 | 54 |
|  |  |  |  |  |  |  | -41 | 20 | 24 |
|  |  |  |  |  |  |  | 5 | -61 | 52 |
|  |  |  |  |  |  |  | 50 | -59 | 37 |
|  |  |  |  |  |  |  | 43 | -67 | -49 |
| Honey et al. | 2000 | 10.1006/nimg.2000.0624 | 2 > 0 back | Letters | Identity | 20 | -36 | -51 | 43 |
|  |  |  |  |  |  |  | -45 | 22 | 23 |
|  |  |  |  |  |  |  | 39 | 53 | 6 |
|  |  |  |  |  |  |  | 2 | 15 | 48 |
|  |  |  |  |  |  |  | -45 | -1 | 43 |
|  |  |  |  |  |  |  | 36 | -1 | 54 |
|  |  |  |  |  |  |  | -20 | -8 | 56 |
|  |  |  |  |  |  |  | -37 | -77 | -11 |
|  |  |  |  |  |  |  | -37 | -68 | -12 |
|  |  |  |  |  |  |  | 29 | -74 | -18 |
| Martinkauppi et al. | 2000 | 10.1093/cercor/10.9.889 | 3 > 1 back | Tones (auditory) | Spatial | 10 | 6 | 23 | 50 |
|  |  |  |  |  |  |  | -3 | 22 | 58 |
|  |  |  |  |  |  |  | 31 | 14 | 51 |
|  |  |  |  |  |  |  | -23 | 15 | 55 |
|  |  |  |  |  |  |  | 45 | 39 | 24 |
|  |  |  |  |  |  |  | -43 | 42 | 31 |
|  |  |  |  |  |  |  | 43 | 21 | 49 |
|  |  |  |  |  |  |  | -39 | 22 | 45 |
|  |  |  |  |  |  |  | 50 | 23 | 14 |
|  |  |  |  |  |  |  | -46 | 23 | 14 |
|  |  |  |  |  |  |  | 33 | 56 | -6 |
|  |  |  |  |  |  |  | -36 | 58 | 3 |
|  |  |  |  |  |  |  | 46 | 4 | 35 |
|  |  |  |  |  |  |  | -43 | 10 | 41 |
|  |  |  |  |  |  |  | 27 | -65 | 57 |
|  |  |  |  |  |  |  | -23 | -65 | 54 |
|  |  |  |  |  |  |  | 40 | -50 | 58 |
|  |  |  |  |  |  |  | -36 | -47 | 50 |
|  |  |  |  |  |  |  | 47 | -46 | 44 |
|  |  |  |  |  |  |  | -49 | -47 | 47 |
|  |  |  |  |  |  |  | 8 | -60 | 51 |
|  |  |  |  |  |  |  | -4 | -63 | 59 |
|  |  |  |  |  |  |  | 56 | -46 | 0 |
|  |  |  |  |  |  |  | -53 | -52 | 9 |
|  |  |  |  |  |  |  | 47 | -66 | -4 |
|  |  |  |  |  |  |  | -51 | -58 | -7 |
|  |  |  |  |  |  |  | 9 | 24 | 25 |
|  |  |  |  |  |  |  | -3 | 22 | 33 |
|  |  |  |  |  |  |  | 7 | -40 | 36 |
|  |  |  |  |  |  |  | -1 | -43 | 28 |
|  |  |  |  |  |  |  | 38 | 21 | 5 |
|  |  |  |  |  |  |  | -35 | 20 | 2 |
|  |  |  | 2 > 1 back | Tones (auditory) | Spatial | 10 | 6 | 27 | 48 |
|  |  |  |  |  |  |  | -2 | 30 | 56 |
|  |  |  |  |  |  |  | 26 | 10 | 58 |
|  |  |  |  |  |  |  | -22 | 13 | 55 |
|  |  |  |  |  |  |  | 44 | 38 | 28 |
|  |  |  |  |  |  |  | -43 | 31 | 30 |
|  |  |  |  |  |  |  | 39 | 18 | 46 |
|  |  |  |  |  |  |  | -38 | 22 | 45 |
|  |  |  |  |  |  |  | 44 | 26 | 9 |
|  |  |  |  |  |  |  | -46 | 25 | 13 |
|  |  |  |  |  |  |  | 24 | 57 | -6 |
|  |  |  |  |  |  |  | -25 | 57 | 6 |
|  |  |  |  |  |  |  | 46 | -1 | 40 |
|  |  |  |  |  |  |  | -41 | 10 | 40 |
|  |  |  |  |  |  |  | 23 | -53 | 61 |
|  |  |  |  |  |  |  | -21 | -62 | 57 |
|  |  |  |  |  |  |  | 35 | -48 | 48 |
|  |  |  |  |  |  |  | -37 | -48 | 50 |
|  |  |  |  |  |  |  | 45 | -48 | 45 |
|  |  |  |  |  |  |  | -40 | -54 | 49 |
|  |  |  |  |  |  |  | 7 | -56 | 56 |
|  |  |  |  |  |  |  | -5 | -62 | 55 |
|  |  |  |  |  |  |  | 54 | -42 | 2 |
|  |  |  |  |  |  |  | -51 | -54 | 7 |
|  |  |  |  |  |  |  | -49 | -58 | -10 |
|  |  |  |  |  |  |  | 10 | 36 | 17 |
|  |  |  |  |  |  |  | -4 | 27 | 26 |
|  |  |  |  |  |  |  | 10 | -42 | 37 |
|  |  |  |  |  |  |  | 41 | 6 | -1 |
|  |  |  |  |  |  |  | -38 | 11 | 1 |
| Ragland et al. | 2002 | 10.1037/0894-4105.16.3.370 | 1 > 0 back | Letters | Identity | 11 | 40 | 27 | 10 |
|  |  |  |  |  |  |  | 40 | 24 | 24 |
|  |  |  |  |  |  |  | 32 | -1 | 35 |
|  |  |  |  |  |  |  | 41 | -39 | 39 |
|  |  |  |  |  |  |  | -33 | -48 | 46 |
|  |  |  |  |  |  |  | -46 | -35 | 44 |
|  |  |  | 2 > 0 back | Letters | Identity | 11 | 40 | 21 | -7 |
|  |  |  |  |  |  |  | 53 | -51 | 4 |
|  |  |  |  |  |  |  | 40 | 59 | -7 |
|  |  |  |  |  |  |  | 49 | 41 | 22 |
|  |  |  |  |  |  |  | -51 | 17 | 4 |
|  |  |  |  |  |  |  | 49 | -39 | 43 |
|  |  |  |  |  |  |  | -54 | -39 | 45 |
|  |  |  | 2 > 1 back | Letters | Identity | 11 | 31 | 30 | 1 |
|  |  |  |  |  |  |  | -51 | 17 | 4 |
|  |  |  |  |  |  |  | 40 | 53 | 12 |
|  |  |  |  |  |  |  | -38 | 31 | 20 |
|  |  |  |  |  |  |  | -46 | 11 | 35 |
|  |  |  |  |  |  |  | 45 | 12 | 38 |
|  |  |  |  |  |  |  | -12 | -8 | 14 |
|  |  |  |  |  |  |  | 6 | 29 | 33 |
|  |  |  |  |  |  |  | 49 | -43 | 44 |
|  |  |  |  |  |  |  | -24 | -56 | 46 |
|  |  |  | 2 > 0 back | Fractals | Identity | 11 | -25 | 25 | -2 |
|  |  |  |  |  |  |  | 36 | 25 | -3 |
|  |  |  |  |  |  |  | -34 | 47 | 0 |
|  |  |  |  |  |  |  | 44 | 36 | 18 |
|  |  |  |  |  |  |  | -42 | 31 | 20 |
|  |  |  |  |  |  |  | -46 | 11 | 31 |
|  |  |  |  |  |  |  | 1 | 33 | 37 |
|  |  |  |  |  |  |  | 45 | -47 | 44 |
|  |  |  |  |  |  |  | -28 | -52 | 46 |
|  |  |  | 1 > 0 back | Fractals | Identity | 11 | -25 | -56 | -3 |
|  |  |  |  |  |  |  | -42 | 31 | 20 |
|  |  |  |  |  |  |  | -29 | -1 | 41 |
|  |  |  |  |  |  |  | 41 | -1 | 40 |
|  |  |  |  |  |  |  | -37 | -56 | 51 |
|  |  |  | 2 > 1 back | Fractals | Identity | 11 | -34 | 47 | 5 |
|  |  |  |  |  |  |  | 49 | 36 | 18 |
|  |  |  |  |  |  |  | 45 | 29 | 32 |
|  |  |  |  |  |  |  | -46 | 7 | 31 |
|  |  |  |  |  |  |  | -3 | 33 | 33 |
|  |  |  |  |  |  |  | -33 | -52 | 46 |
| Kumari et al. | 2003 | 10.1016/S1053-8119(03)00110-1 | 3,2,1 > 0 back | Numbers | Identity and Spatial | 11 | -5 | 15 | 52 |
|  |  |  |  |  |  |  | -31 | 2 | 54 |
|  |  |  |  |  |  |  | -39 | -26 | 50 |
|  |  |  |  |  |  |  | -33 | -51 | 55 |
|  |  |  |  |  |  |  | 40 | 47 | 17 |
|  |  |  |  |  |  |  | 39 | -10 | -56 |
|  |  |  |  |  |  |  | 41 | -41 | 44 |
|  |  |  |  |  |  |  | 25 | -63 | -30 |
| Perlstein et al. | 2003 | 10.1016/S0006-3223(02)01675-X | 2 > 1 > 0 back | Letters | Identity | 15 | 37 | 8 | 52 |
|  |  |  |  |  |  |  | 7 | 21 | 38 |
|  |  |  |  |  |  |  | 44 | 42 | 15 |
|  |  |  |  |  |  |  | -39 | 43 | 16 |
|  |  |  |  |  |  |  | 49 | 19 | 8 |
|  |  |  |  |  |  |  | -41 | 19 | 12 |
|  |  |  |  |  |  |  | 48 | -53 | 40 |
|  |  |  |  |  |  |  | -36 | -50 | 44 |
|  |  |  |  |  |  |  | 2 | 17 | 7 |
| Veltman et al. | 2003 | 10.1016/S1053-8119(02)00049-6 | 3 > 2 > 1 > 0 back | Letters | Identity | 22 | -38 | 55 | 14 |
|  |  |  |  |  |  |  | -51 | 30 | 27 |
|  |  |  |  |  |  |  | -47 | 42 | 22 |
|  |  |  |  |  |  |  | -54 | 15 | 5 |
|  |  |  |  |  |  |  | 60 | 21 | 33 |
|  |  |  |  |  |  |  | 50 | 44 | 31 |
|  |  |  |  |  |  |  | 60 | 16 | 13 |
|  |  |  |  |  |  |  | -47 | -35 | 57 |
|  |  |  |  |  |  |  | -48 | -58 | -15 |
|  |  |  |  |  |  |  | 12 | 21 | 64 |
|  |  |  |  |  |  |  | 43 | -63 | -46 |
| Perlstein et al. | 2004 | 10.1017/S1355617704105110 | 3 > 2 > 1 > 0 back | Letters | Identity | 26 | -28 | 41 | 23 |
|  |  |  |  |  |  |  | -36 | 32 | 28 |
|  |  |  |  |  |  |  | 37 | 52 | 22 |
|  |  |  |  |  |  |  | 35 | 17 | 10 |
|  |  |  |  |  |  |  | -37 | 17 | 17 |
|  |  |  |  |  |  |  | 49 | 0 | 37 |
|  |  |  |  |  |  |  | 3 | -15 | -1 |
|  |  |  |  |  |  |  | 29 | -31 | -7 |
|  |  |  |  |  |  |  | -20 | -35 | -2 |
|  |  |  |  |  |  |  | 32 | 45 | -6 |
| Knops et al. | 2006 | 10.1016/j.neuroimage.2005.07.009 | 2 > 1 back | Letters/numbers | Identity | 13 | -25 | 49 | -27 |
|  |  |  |  |  |  |  | -7 | 29 | 41 |
|  |  |  |  |  |  |  | 1 | 26 | 47 |
|  |  |  |  |  |  |  | -46 | 30 | 21 |
|  |  |  |  |  |  |  | -41 | 14 | 56 |
|  |  |  |  |  |  |  | -47 | 48 | -3 |
|  |  |  |  |  |  |  | 56 | 42 | 13 |
|  |  |  |  |  |  |  | 49 | 37 | 10 |
|  |  |  |  |  |  |  | 32 | 18 | 50 |
|  |  |  |  |  |  |  | 36 | 23 | -4 |
|  |  |  |  |  |  |  | 41 | -46 | 43 |
|  |  |  |  |  |  |  | -37 | -42 | 44 |
|  |  |  |  |  |  |  | 45 | -57 | 57 |
|  |  |  |  |  |  |  | -33 | -81 | 9 |
|  |  |  |  |  |  |  | 1 | -51 | -20 |
|  |  |  |  |  |  |  | -16 | -80 | -19 |
| Cerasa et al. | 2008 | 10.1016/j.brainres.2008.01.048 | 2 > 0 back | Shapes | Spatial | 30 | 56 | -36 | 50 |
|  |  |  |  |  |  |  | 36 | -53 | 53 |
|  |  |  |  |  |  |  | -46 | -47 | 57 |
|  |  |  |  |  |  |  | -33 | -54 | 48 |
|  |  |  |  |  |  |  | -42 | 9 | 37 |
|  |  |  |  |  |  |  | -1 | 30 | 45 |
|  |  |  |  |  |  |  | 44 | 51 | 17 |
|  |  |  |  |  |  |  | 42 | 21 | -14 |
|  |  |  |  |  |  |  | -42 | 55 | 6 |
|  |  |  |  |  |  |  | -34 | 26 | -1 |
|  |  |  |  |  |  |  | -10 | -5 | 8 |
|  |  |  |  |  |  |  | 10 | 19 | 9 |
|  |  |  |  |  |  |  | -16 | -4 | 21 |
|  |  |  |  |  |  |  | 10 | -8 | 22 |
|  |  |  |  |  |  |  | -10 | -83 | -20 |
|  |  |  |  |  |  |  | 35 | -68 | -17 |
| Ricciardi et al. | 2006 | 10.1016/j.neuroscience.2005.08.045 | 1 back > rest | 3D shape (tactile) | Identity | 6 | 24 | 59 | -5 |
|  |  |  |  |  |  |  | 14 | 80 | -6 |
|  |  |  |  |  |  |  | -22 | 54 | -5 |
|  |  |  |  |  |  |  | 3 | 33 | 43 |
|  |  |  |  |  |  |  | 2 | 11 | 53 |
|  |  |  |  |  |  |  | -25 | 1 | 69 |
|  |  |  |  |  |  |  | 0 | -8 | 63 |
|  |  |  |  |  |  |  | 38 | 5 | 51 |
|  |  |  |  |  |  |  | 34 | 45 | 32 |
|  |  |  |  |  |  |  | 41 | 50 | 11 |
|  |  |  |  |  |  |  | 45 | 61 | -20 |
|  |  |  |  |  |  |  | -47 | 42 | 23 |
|  |  |  |  |  |  |  | -26 | 71 | 12 |
|  |  |  |  |  |  |  | -2 | 21 | 16 |
|  |  |  |  |  |  |  | 54 | 9 | 15 |
|  |  |  |  |  |  |  | 41 | 37 | -18 |
|  |  |  |  |  |  |  | -55 | 14 | 27 |
|  |  |  |  |  |  |  | -45 | 55 | -7 |
|  |  |  |  |  |  |  | 45 | 19 | -6 |
|  |  |  |  |  |  |  | -36 | 28 | 8 |
|  |  |  |  |  |  |  | -40 | 4 | 8 |
|  |  |  |  |  |  |  | 40 | -13 | 63 |
|  |  |  |  |  |  |  | -37 | -18 | 66 |
|  |  |  |  |  |  |  | 59 | -18 | 32 |
|  |  |  |  |  |  |  | -15 | -29 | 71 |
|  |  |  |  |  |  |  | -55 | -17 | 40 |
|  |  |  |  |  |  |  | 10 | -29 | 80 |
|  |  |  |  |  |  |  | 58 | -40 | 38 |
|  |  |  |  |  |  |  | -58 | -22 | 16 |
|  |  |  |  |  |  |  | 19 | -65 | 52 |
|  |  |  |  |  |  |  | 24 | -72 | 23 |
|  |  |  |  |  |  |  | 6 | -48 | 66 |
|  |  |  |  |  |  |  | -21 | -81 | 36 |
|  |  |  |  |  |  |  | -8 | -82 | 56 |
|  |  |  |  |  |  |  | 28 | -48 | 67 |
|  |  |  |  |  |  |  | -25 | -63 | 57 |
|  |  |  |  |  |  |  | -44 | -21 | -15 |
|  |  |  |  |  |  |  | 51 | -64 | 1 |
|  |  |  |  |  |  |  | 47 | -12 | -27 |
|  |  |  |  |  |  |  | -55 | -60 | -4 |
|  |  |  |  |  |  |  | -43 | -77 | 22 |
|  |  |  |  |  |  |  | 65 | -9 | 0 |
|  |  |  |  |  |  |  | 42 | 19 | -30 |
|  |  |  |  |  |  |  | -51 | 19 | -15 |
|  |  |  |  |  |  |  | 51 | -46 | -14 |
|  |  |  |  |  |  |  | -42 | -45 | -15 |
|  |  |  |  |  |  |  | 21 | -105 | 0 |
|  |  |  |  |  |  |  | -13 | -89 | -7 |
|  |  |  |  |  |  |  | -22 | -72 | 6 |
|  |  |  |  |  |  |  | -29 | -105 | 4 |
|  |  |  |  |  |  |  | 1 | -93 | 11 |
|  |  |  |  |  |  |  | 1 | -79 | 29 |
|  |  |  |  |  |  |  | -43 | -77 | 22 |
|  |  |  |  |  |  |  | 16 | -22 | 6 |
|  |  |  |  |  |  |  | -10 | -22 | 5 |
|  |  |  |  |  |  |  | 11 | 2 | 12 |
|  |  |  |  |  |  |  | -13 | -1 | 14 |
|  |  |  |  |  |  |  | 17 | -51 | -45 |
|  |  |  |  |  |  |  | 38 | -67 | -17 |
|  |  |  |  |  |  |  | 21 | -55 | -12 |
|  |  |  |  |  |  |  | -21 | -45 | -19 |
|  |  |  |  |  |  |  | -38 | -58 | -37 |
|  |  |  | 1 back > rest | 3D shape (visual) | Identity | 6 | 8 | 14 | 72 |
|  |  |  |  |  |  |  | 1 | 16 | 50 |
|  |  |  |  |  |  |  | 33 | 1 | 58 |
|  |  |  |  |  |  |  | 47 | 35 | 25 |
|  |  |  |  |  |  |  | 53 | 11 | 33 |
|  |  |  |  |  |  |  | 38 | 55 | -5 |
|  |  |  |  |  |  |  | -47 | 55 | -5 |
|  |  |  |  |  |  |  | -36 | -3 | 47 |
|  |  |  |  |  |  |  | -54 | 12 | 34 |
|  |  |  |  |  |  |  | -36 | 46 | 14 |
|  |  |  |  |  |  |  | 10 | 39 | -4 |
|  |  |  |  |  |  |  | -9 | 30 | 7 |
|  |  |  |  |  |  |  | -16 | 53 | -12 |
|  |  |  |  |  |  |  | 7 | 30 | 29 |
|  |  |  |  |  |  |  | 57 | 14 | 30 |
|  |  |  |  |  |  |  | 44 | 36 | -18 |
|  |  |  |  |  |  |  | -53 | 20 | -13 |
|  |  |  |  |  |  |  | -58 | 15 | 27 |
|  |  |  |  |  |  |  | -30 | 23 | 7 |
|  |  |  |  |  |  |  | 59 | 12 | -1 |
|  |  |  |  |  |  |  | -51 | 13 | 4 |
|  |  |  |  |  |  |  | 6 | -21 | 66 |
|  |  |  |  |  |  |  | -59 | -20 | 17 |
|  |  |  |  |  |  |  | 17 | -74 | 61 |
|  |  |  |  |  |  |  | 31 | -72 | 24 |
|  |  |  |  |  |  |  | 3 | -56 | 66 |
|  |  |  |  |  |  |  | -6 | -82 | 51 |
|  |  |  |  |  |  |  | -23 | -63 | 57 |
|  |  |  |  |  |  |  | 35 | -42 | 52 |
|  |  |  |  |  |  |  | 57 | -27 | 33 |
|  |  |  |  |  |  |  | -39 | -44 | 53 |
|  |  |  |  |  |  |  | 43 | -74 | 1 |
|  |  |  |  |  |  |  | -47 | -67 | 3 |
|  |  |  |  |  |  |  | -35 | -86 | 5 |
|  |  |  |  |  |  |  | 50 | -49 | 11 |
|  |  |  |  |  |  |  | 31 | -44 | -17 |
|  |  |  |  |  |  |  | -35 | -61 | -18 |
|  |  |  |  |  |  |  | 14 | -81 | -11 |
|  |  |  |  |  |  |  | 7 | -102 | 18 |
|  |  |  |  |  |  |  | 17 | -82 | 11 |
|  |  |  |  |  |  |  | -5 | -77 | 20 |
|  |  |  |  |  |  |  | 14 | 9 | 11 |
|  |  |  |  |  |  |  | 36 | -83 | -24 |
| Nebel et al. | 2005 | 10.1016/j.cogbrainres.2005.09.011 | 1 back > rest | Letters/shapes | Identity | 19 | 34 | 49 | 15 |
|  |  |  |  |  |  |  | 31 | 28 | -15 |
|  |  |  |  |  |  |  | -32 | 21 | -11 |
|  |  |  |  |  |  |  | -11 | 8 | 52 |
|  |  |  |  |  |  |  | 12 | 23 | 41 |
|  |  |  |  |  |  |  | 10 | 27 | 31 |
|  |  |  |  |  |  |  | -9 | 15 | 46 |
|  |  |  |  |  |  |  | -9 | 23 | 34 |
|  |  |  |  |  |  |  | 37 | -55 | 53 |
|  |  |  |  |  |  |  | 49 | -34 | 42 |
|  |  |  | 2 back > rest | Letters/shapes | Identity | 19 | 47 | 42 | 25 |
|  |  |  |  |  |  |  | 38 | 55 | 12 |
|  |  |  |  |  |  |  | 49 | 47 | 19 |
|  |  |  |  |  |  |  | 32 | 18 | 50 |
|  |  |  |  |  |  |  | 32 | 15 | 61 |
|  |  |  |  |  |  |  | 49 | 15 | 50 |
|  |  |  |  |  |  |  | -26 | 9 | 51 |
|  |  |  |  |  |  |  | 38 | 32 | -12 |
|  |  |  |  |  |  |  | 44 | 23 | -14 |
|  |  |  |  |  |  |  | -48 | 7 | 18 |
|  |  |  |  |  |  |  | -36 | 21 | -11 |
|  |  |  |  |  |  |  | 38 | 24 | -4 |
|  |  |  |  |  |  |  | 12 | 36 | 28 |
|  |  |  |  |  |  |  | -7 | 15 | 51 |
|  |  |  |  |  |  |  | -7 | 9 | 61 |
|  |  |  |  |  |  |  | 14 | 21 | 36 |
|  |  |  |  |  |  |  | -33 | -43 | 53 |
|  |  |  |  |  |  |  | -17 | -57 | 66 |
|  |  |  |  |  |  |  | 15 | -66 | 61 |
|  |  |  |  |  |  |  | 49 | -34 | 52 |
|  |  |  |  |  |  |  | 49 | -42 | 55 |
|  |  |  |  |  |  |  | -48 | -29 | 46 |
|  |  |  |  |  |  |  | -10 | -83 | -13 |
|  |  |  |  |  |  |  | -14 | 6 | 15 |
|  |  |  |  |  |  |  | -20 | 7 | 7 |
|  |  |  |  |  |  |  | 31 | -63 | -23 |
|  |  |  |  |  |  |  | 8 | -80 | -20 |
|  |  |  |  |  |  |  | 3 | -58 | -12 |
|  |  |  |  |  |  |  | -31 | -70 | -22 |
|  |  |  |  |  |  |  | -38 | -63 | -28 |
| Matsuo et al. | 2007 | 10.1038/sj.mp.4001894 | 1 > 0 back | Numbers | Identity and Spatial | 15 | -31 | 6 | 47 |
|  |  |  |  |  |  |  | -46 | 23 | 16 |
|  |  |  |  |  |  |  | -24 | 17 | 43 |
|  |  |  |  |  |  |  | 48 | 60 | -5 |
|  |  |  | 2 > 0 back | Numbers | Identity and Spatial | 15 | 49 | 40 | 13 |
|  |  |  |  |  |  |  | -26 | 6 | 45 |
| Schmidt et al. | 2009 | 10.1002/hbm.20783 | 3,2,1 > 0 back | Letters | Identity | 25 | 30 | 20 | 53 |
|  |  |  |  |  |  |  | -5 | 30 | 44 |
|  |  |  |  |  |  |  | 47 | 43 | 26 |
|  |  |  |  |  |  |  | 35 | 70 | -3 |
|  |  |  |  |  |  |  | -36 | 24 | -11 |
|  |  |  |  |  |  |  | 17 | -65 | 60 |
|  |  |  |  |  |  |  | -28 | 7 | 60 |
|  |  |  |  |  |  |  | -50 | -45 | 57 |
| Schmidt et al. | 2009 | 10.1002/hbm.20783 | 3,2,1 > 0 back | Letters | Identity | 21 | -3 | 22 | 54 |
|  |  |  |  |  |  |  | 60 | 28 | 25 |
|  |  |  |  |  |  |  | -59 | 26 | 30 |
|  |  |  |  |  |  |  | -44 | 35 | -16 |
|  |  |  |  |  |  |  | 50 | -50 | 64 |
| Qin et al. | 2009 | 10.1016/j.biopsych.2009.03.006 | 2 > 0 back | Numbers | Identity | 27 | 28 | 4 | 58 |
|  |  |  |  |  |  |  | -30 | 2 | 58 |
|  |  |  |  |  |  |  | 32 | 24 | 0 |
|  |  |  |  |  |  |  | -28 | 24 | 4 |
|  |  |  |  |  |  |  | 36 | 48 | 18 |
|  |  |  |  |  |  |  | -34 | 52 | 14 |
|  |  |  |  |  |  |  | 18 | -4 | 0 |
|  |  |  |  |  |  |  | -16 | -4 | 0 |
|  |  |  |  |  |  |  | 40 | -44 | 46 |
|  |  |  |  |  |  |  | -44 | -42 | 50 |
|  |  |  |  |  |  |  | 18 | -66 | 60 |
|  |  |  |  |  |  |  | -14 | -68 | 58 |
|  |  |  |  |  |  |  | 30 | -58 | -32 |
|  |  |  |  |  |  |  | -30 | -58 | -34 |
| Scheuerecker et al. | 2008 | 10.1016/j.jpsychires.2007.04.001 | 2 > 0 back | Letters | Identity | 23 | -32 | 20 | -4 |
|  |  |  |  |  |  |  | 0 | -24 | 8 |
|  |  |  |  |  |  |  | -54 | -44 | 40 |
|  |  |  |  |  |  |  | 34 | -64 | 48 |
|  |  |  |  |  |  |  | 50 | 20 | 22 |
|  |  |  |  |  |  |  | 32 | 22 | -4 |
|  |  |  |  |  |  |  | -36 | 54 | 4 |
|  |  |  |  |  |  |  | 36 | -62 | -22 |
| Choo et al. | 2005 | 10.1016/j.neuroimage.2004.11.029 | 2 > 1 back | Letters | Identity | 12 | -48 | 14 | 22 |
|  |  |  |  |  |  |  | 39 | 38 | 29 |
|  |  |  |  |  |  |  | -1 | 29 | 42 |
|  |  |  |  |  |  |  | -33 | -48 | 40 |
|  |  |  |  |  |  |  | 35 | -62 | 41 |
|  |  |  |  |  |  |  | 6 | -66 | 56 |
|  |  |  |  |  |  |  | -16 | -18 | 18 |
| Koppelstaetter et al. | 2008 | 10.1016/j.neuroimage.2007.08.037 | 2 > 0 back | Letters | Identity | 15 | -3 | 25 | 38 |
|  |  |  |  |  |  |  | -42 | 37 | 12 |
|  |  |  |  |  |  |  | -29 | 13 | 48 |
|  |  |  |  |  |  |  | -28 | -58 | 51 |
|  |  |  |  |  |  |  | -50 | -33 | 51 |
|  |  |  |  |  |  |  | -7 | -58 | 54 |
|  |  |  |  |  |  |  | -16 | -97 | 11 |
|  |  |  |  |  |  |  | -34 | -68 | -24 |
|  |  |  |  |  |  |  | 6 | 25 | 38 |
|  |  |  |  |  |  |  | 53 | 33 | 6 |
|  |  |  |  |  |  |  | 32 | 8 | 43 |
|  |  |  |  |  |  |  | 28 | -58 | 46 |
|  |  |  |  |  |  |  | 54 | -38 | 42 |
|  |  |  |  |  |  |  | 24 | -62 | 54 |
|  |  |  |  |  |  |  | 23 | -97 | 10 |
|  |  |  |  |  |  |  | 31 | -70 | -34 |
| Harvey et al. | 2005 | 10.1016/j.neuroimage.2005.02.048 | 3,2,1 > 0 back | Letters | Identity | 10 | -34 | -42 | 47 |
|  |  |  |  |  |  |  | -11 | -67 | 56 |
|  |  |  |  |  |  |  | 18 | -58 | 48 |
|  |  |  |  |  |  |  | 57 | -27 | 37 |
|  |  |  |  |  |  |  | 47 | -62 | 41 |
|  |  |  |  |  |  |  | 11 | 46 | 28 |
|  |  |  |  |  |  |  | -47 | 23 | 18 |
|  |  |  |  |  |  |  | -41 | 50 | -2 |
|  |  |  |  |  |  |  | 43 | 50 | 0 |
|  |  |  |  |  |  |  | 53 | 46 | 17 |
| Forn & Barros-Loscertales | 2007 | 10.1002/hbm.20284 | 2 > 0 back | Letters (auditory) | Identity |  | 41 | 44 | 23 |
|  |  |  |  |  |  |  | 43 | 26 | 25 |
|  |  |  |  |  |  |  | 32 | 48 | -10 |
|  |  |  |  |  |  |  | 6 | 15 | 50 |
|  |  |  |  |  |  |  | -40 | 30 | 26 |
|  |  |  |  |  |  |  | -25 | 62 | -9 |
|  |  |  |  |  |  |  | -42 | 7 | 37 |
|  |  |  |  |  |  |  | 33 | -57 | 49 |
|  |  |  |  |  |  |  | -27 | -56 | 50 |
|  |  |  |  |  |  |  | -36 | -57 | 50 |
| Loughead et al. | 2009 | 10.1038/mp.2008.132 | 3 > 2 > 1 > 0 back | Fractals | Identity |  | -3 | 31 | 28 |
|  |  |  |  |  |  |  | 43 | -45 | 47 |
|  |  |  |  |  |  |  | -33 | -45 | 42 |
|  |  |  |  |  |  |  | 47 | 38 | 12 |
|  |  |  |  |  |  |  | 23 | 15 | 46 |
|  |  |  |  |  |  |  | 44 | 19 | -14 |
|  |  |  |  |  |  |  | -36 | 19 | -13 |
|  |  |  |  |  |  |  | -31 | 10 | 52 |
|  |  |  |  |  |  |  | -38 | -67 | -28 |
|  |  |  |  |  |  |  | -44 | 33 | 16 |
|  |  |  |  |  |  |  | 38 | -68 | -25 |
|  |  |  |  |  |  |  | -42 | 9 | 23 |
|  |  |  |  |  |  |  | 53 | 18 | 19 |
| Allen et al. | 2006 | 10.1007/s00213-006-0444-x | 2 > 0 back | Letters | Identity | 10 | 48 | -46 | 41 |
|  |  |  |  |  |  |  | -42 | 36 | 22 |
|  |  |  |  |  |  |  | 44 | 36 | 20 |
|  |  |  |  |  |  |  | -25 | -66 | 49 |
|  |  |  |  |  |  |  | 1 | 18 | 47 |
|  |  |  |  |  |  |  | 6 | 26 | 34 |
| Meisenzhal et al. | 2006 | 10.1007/s00406-006-0687-x | 2 > 0 back | Letters | Identity | 12 | -52 | 23 | -16 |
|  |  |  |  |  |  |  | -48 | -39 | 43 |
|  |  |  |  |  |  |  | -43 | -41 | 54 |
|  |  |  |  |  |  |  | 53 | 16 | 19 |
|  |  |  |  |  |  |  | 56 | 27 | 9 |
|  |  |  |  |  |  |  | -12 | -9 | 8 |
|  |  |  |  |  |  |  | -10 | 4 | 5 |
|  |  |  |  |  |  |  | 8 | -18 | 0 |
|  |  |  |  |  |  |  | -42 | 5 | 28 |
|  |  |  |  |  |  |  | 37 | 22 | -25 |
|  |  |  |  |  |  |  | -48 | 28 | 23 |
|  |  |  |  |  |  |  | 51 | -38 | 44 |
|  |  |  |  |  |  |  | -67 | -39 | 27 |
|  |  |  |  |  |  |  | 37 | -57 | 55 |
|  |  |  |  |  |  |  | -38 | 52 | -1 |
|  |  |  |  |  |  |  | -36 | 48 | 8 |
|  |  |  |  |  |  |  | -44 | 16 | 39 |
|  |  |  |  |  |  |  | -1 | 42 | 11 |
|  |  |  |  |  |  |  | -27 | 23 | -13 |
|  |  |  |  |  |  |  | 3 | 38 | 23 |
| Frangou et al. | 2008 | 10.1016/j.eurpsy.2007.05.002 | 3,2,1 > 0 back | Letters | Identity | 7 | -55 | 12 | 40 |
|  |  |  |  |  |  |  | -28 | -63 | 65 |
|  |  |  |  |  |  |  | -30 | -64 | 52 |
|  |  |  |  |  |  |  | -57 | -46 | 14 |
|  |  |  |  |  |  |  | -14 | -1 | 14 |
|  |  |  |  |  |  |  | 6 | 26 | 53 |
|  |  |  |  |  |  |  | 58 | 19 | 37 |
|  |  |  |  |  |  |  | 53 | 45 | 17 |
|  |  |  |  |  |  |  | 43 | -59 | 59 |
|  |  |  |  |  |  |  | 47 | -48 | 42 |
|  |  |  |  |  |  |  | 10 | 1 | 9 |
|  |  |  | 3 > 2 > 1 > 0 back | Letters | Identity | 7 | -52 | 8 | 43 |
|  |  |  |  |  |  |  | 41 | 28 | 52 |
|  |  |  |  |  |  |  | 41 | 20 | 55 |
|  |  |  |  |  |  |  | 10 | 47 | 26 |
|  |  |  |  |  |  |  | 40 | 67 | -14 |
| Deckersbach et al. | 2008 | 10.1111/j.1399-5618.2008.00633.x | 2 back > fixation | Letters | Identity | 17 | -8 | 4 | -56 |
|  |  |  |  |  |  |  | -58 | 6 | 22 |
|  |  |  |  |  |  |  | 36 | 18 | -6 |
|  |  |  |  |  |  |  | 26 | -8 | 46 |
|  |  |  |  |  |  |  | 12 | -6 | 12 |
|  |  |  |  |  |  |  | -46 | -4 | 42 |
|  |  |  |  |  |  |  | -56 | -16 | 28 |
|  |  |  |  |  |  |  | 24 | -10 | -2 |
|  |  |  |  |  |  |  | -8 | -20 | -12 |
|  |  |  |  |  |  |  | 68 | -12 | -6 |
|  |  |  |  |  |  |  | -58 | -10 | -8 |
|  |  |  |  |  |  |  | -32 | 18 | 2 |
| Sanchez-Carrion et al. | 2008 | 10.1089/neu.2007.0417 | 2 > 0 back | Numbers | Identity | 14 | 24 | 4 | 62 |
|  |  |  |  |  |  |  | -4 | 6 | 60 |
|  |  |  |  |  |  |  | -52 | 12 | 26 |
|  |  |  |  |  |  |  | -34 | 24 | 2 |
|  |  |  |  |  |  |  | -36 | 44 | 24 |
|  |  |  |  |  |  |  | -40 | 50 | 8 |
|  |  |  |  |  |  |  | 34 | 22 | 4 |
|  |  |  |  |  |  |  | -42 | -46 | 46 |
|  |  |  |  |  |  |  | -22 | -66 | 42 |
|  |  |  |  |  |  |  | -32 | -56 | 48 |
|  |  |  |  |  |  |  | 40 | -54 | 52 |
|  |  |  |  |  |  |  | 44 | -46 | 44 |
|  |  |  |  |  |  |  | 16 | -68 | 56 |
|  |  |  |  |  |  |  | 8 | -76 | -26 |
|  |  |  |  |  |  |  | 32 | -56 | -38 |
|  |  |  |  |  |  |  | -34 | -54 | -38 |
|  |  |  | 3 > 0 back | Numbers | Identity | 14 | -52 | 10 | 24 |
|  |  |  |  |  |  |  | -30 | -2 | 46 |
|  |  |  |  |  |  |  | -32 | -2 | 56 |
|  |  |  |  |  |  |  | 24 | 2 | 62 |
|  |  |  |  |  |  |  | 4 | 20 | 44 |
|  |  |  |  |  |  |  | -34 | 26 | 6 |
|  |  |  |  |  |  |  | 42 | 32 | 34 |
|  |  |  |  |  |  |  | 40 | 46 | 20 |
|  |  |  |  |  |  |  | 52 | 10 | 30 |
|  |  |  |  |  |  |  | -42 | 50 | 10 |
|  |  |  |  |  |  |  | -34 | 42 | 20 |
|  |  |  |  |  |  |  | 34 | 22 | 4 |
|  |  |  |  |  |  |  | -20 | -68 | 40 |
|  |  |  |  |  |  |  | -38 | -48 | 44 |
|  |  |  |  |  |  |  | 44 | -46 | 46 |
|  |  |  |  |  |  |  | 34 | -74 | 36 |
|  |  |  |  |  |  |  | 14 | -82 | -26 |
|  |  |  |  |  |  |  | 38 | -66 | -32 |
| Caseras et al. | 2006 | 10.1097/01.psy.0000242770.50979.5f | 3 > 2 > 1 > 0 back | Letters | Identity | 12 | -2 | -69 | 47 |
|  |  |  |  |  |  |  | 44 | 43 | 16 |
|  |  |  |  |  |  |  | 9 | -86 | -14 |
|  |  |  |  |  |  |  | 48 | 29 | 32 |
|  |  |  |  |  |  |  | 2 | -74 | 40 |
|  |  |  |  |  |  |  | 33 | 15 | 59 |
|  |  |  |  |  |  |  | 36 | 63 | 11 |
|  |  |  |  |  |  |  | 6 | 29 | 37 |
|  |  |  |  |  |  |  | 45 | -50 | 37 |
|  |  |  |  |  |  |  | 56 | 31 | 14 |
| Cader et al. | 2006 | 10.1093/brain/awh670 | 2 > 1 back | Letters | Identity | 16 | 40 | 40 | 28 |
|  |  |  |  |  |  |  | -28 | -12 | 44 |
|  |  |  |  |  |  |  | -30 | 18 | -14 |
|  |  |  |  |  |  |  | -32 | 22 | -6 |
|  |  |  |  |  |  |  | 34 | -56 | 50 |
|  |  |  |  |  |  |  | -44 | -40 | 50 |
|  |  |  |  |  |  |  | 8 | 14 | 44 |
|  |  |  |  |  |  |  | 6 | 32 | 30 |
|  |  |  | 3 > 2 > 1 back | Letters | Identity | 16 | 40 | 34 | 28 |
|  |  |  |  |  |  |  | -48 | -2 | 36 |
|  |  |  |  |  |  |  | -34 | 20 | -10 |
|  |  |  |  |  |  |  | -34 | 22 | -8 |
|  |  |  |  |  |  |  | 34 | -58 | 50 |
|  |  |  |  |  |  |  | -44 | -40 | 48 |
|  |  |  |  |  |  |  | 8 | 10 | 56 |
|  |  |  |  |  |  |  | 4 | 16 | 34 |
| Kumari et al. | 2006 | 10.1016/j.schres.2006.02.017 | 1 > 0 back | Dots | Spatial | 13 | -48 | 2 | 46 |
|  |  |  |  |  |  |  | -28 | 12 | 48 |
|  |  |  |  |  |  |  | -34 | 5 | 52 |
|  |  |  |  |  |  |  | -38 | 40 | 26 |
|  |  |  |  |  |  |  | -44 | 34 | 22 |
|  |  |  |  |  |  |  | -48 | 40 | 18 |
|  |  |  |  |  |  |  | -44 | -48 | 46 |
|  |  |  |  |  |  |  | 36 | 8 | 46 |
|  |  |  |  |  |  |  | 26 | 15 | 52 |
|  |  |  |  |  |  |  | 48 | 12 | 42 |
|  |  |  |  |  |  |  | 36 | 30 | 36 |
|  |  |  |  |  |  |  | 44 | 42 | 24 |
|  |  |  |  |  |  |  | 36 | 24 | -6 |
|  |  |  |  |  |  |  | 48 | 20 | -8 |
|  |  |  |  |  |  |  | 0 | 10 | 46 |
|  |  |  |  |  |  |  | 4 | 24 | 36 |
|  |  |  |  |  |  |  | -12 | 18 | 48 |
|  |  |  |  |  |  |  | 14 | -70 | 58 |
|  |  |  |  |  |  |  | 18 | -74 | 50 |
|  |  |  |  |  |  |  | 58 | -42 | 40 |
|  |  |  |  |  |  |  | 52 | -46 | 6 |
|  |  |  |  |  |  |  | 44 | -50 | 48 |
|  |  |  | 2 > 0 back | Dots | Spatial | 13 | -42 | 46 | 2 |
|  |  |  |  |  |  |  | -48 | 18 | 20 |
|  |  |  |  |  |  |  | -4 | 24 | 36 |
|  |  |  |  |  |  |  | -16 | -4 | 12 |
|  |  |  |  |  |  |  | -12 | -55 | 46 |
|  |  |  |  |  |  |  | -42 | -50 | 46 |
|  |  |  |  |  |  |  | -4 | -76 | -34 |
|  |  |  |  |  |  |  | -28 | -74 | -32 |
|  |  |  |  |  |  |  | -30 | -52 | -35 |
|  |  |  |  |  |  |  | 36 | 60 | 8 |
|  |  |  |  |  |  |  | 34 | 34 | 40 |
|  |  |  |  |  |  |  | 34 | 8 | 48 |
|  |  |  |  |  |  |  | 48 | 14 | 28 |
|  |  |  |  |  |  |  | 38 | 26 | -4 |
|  |  |  |  |  |  |  | 4 | 26 | 36 |
|  |  |  |  |  |  |  | 2 | 18 | 48 |
|  |  |  |  |  |  |  | 8 | -62 | 46 |
|  |  |  |  |  |  |  | 40 | -64 | -36 |
|  |  |  | 0 back > rest | Dots | Spatial | 13 | -42 | 0 | 54 |
|  |  |  |  |  |  |  | -35 | -22 | 42 |
|  |  |  |  |  |  |  | -52 | -24 | 50 |
|  |  |  |  |  |  |  | -46 | 0 | 6 |
|  |  |  |  |  |  |  | -58 | 8 | 10 |
|  |  |  |  |  |  |  | -50 | 8 | 28 |
|  |  |  |  |  |  |  | -2 | -10 | 48 |
|  |  |  |  |  |  |  | -8 | 6 | 52 |
|  |  |  |  |  |  |  | -58 | -40 | 16 |
|  |  |  |  |  |  |  | 44 | 0 | 48 |
|  |  |  |  |  |  |  | 30 | -2 | 52 |
|  |  |  |  |  |  |  | 6 | 4 | 50 |
|  |  |  |  |  |  |  | 54 | -66 | -10 |
|  |  |  |  |  |  |  | 8 | -58 | -28 |
|  |  |  |  |  |  |  | 22 | -56 | -32 |
|  |  |  |  |  |  |  | 4 | -70 | -20 |
| Takeuchi et al. | 2012 | 10.1002/hbm.22167 | 2 > 0 back | Letters | Identity | 248 | -30 | -54 | 30 |
|  |  |  |  |  |  |  | 24 | 3 | 42 |
|  |  |  |  |  |  |  | -30 | 27 | 21 |
|  |  |  |  |  |  |  | 33 | -48 | 33 |
|  |  |  |  |  |  |  | -24 | 3 | 45 |
|  |  |  |  |  |  |  | 30 | -63 | 27 |
|  |  |  |  |  |  |  | -6 | 9 | 48 |
|  |  |  |  |  |  |  | 45 | -57 | 42 |
|  |  |  |  |  |  |  | -24 | 42 | 12 |
|  |  |  |  |  |  |  | -36 | 0 | 27 |
|  |  |  |  |  |  |  | -42 | -60 | 42 |
| Kim et al. | 2003 | 10.1176/appi.ajp.160.5.919 | 2 > 0 back | Shapes | Identity | 12 | 43 | 28 | 48 |
|  |  |  |  |  |  |  | -30 | 67 | -15 |
|  |  |  |  |  |  |  | 8 | 39 | 32 |
|  |  |  |  |  |  |  | -24 | -57 | 60 |
|  |  |  |  |  |  |  | 67 | -34 | 47 |
|  |  |  |  |  |  |  | -54 | -60 | 56 |
|  |  |  |  |  |  |  | -23 | 12 | -10 |
|  |  |  |  |  |  |  | -49 | -90 | -37 |
| Honey et al. | 2003 | 10.1017/s0033291703007864 | 2 > 0 back | Letters | Identity | 27 | -44 | -53 | 48 |
|  |  |  |  |  |  |  | -45 | 8 | 37 |
|  |  |  |  |  |  |  | 42 | 48 | 7 |
|  |  |  |  |  |  |  | -34 | 22 | -2 |
|  |  |  |  |  |  |  | 57 | 19 | 15 |
|  |  |  |  |  |  |  | 5 | 15 | 48 |
|  |  |  |  |  |  |  | -45 | 2 | 43 |
|  |  |  |  |  |  |  | -40 | -73 | -11 |
|  |  |  |  |  |  |  | 36 | -68 | -13 |
|  |  |  |  |  |  |  | -12 | -83 | -16 |
|  |  |  |  |  |  |  | 32 | -68 | -19 |
| Binder et al. | 2006 | 10.1148/radiol.2381041622 | 2 > 0 back | Letters | Identity | 12 | -24 | -62 | 47 |
|  |  |  |  |  |  |  | -32 | -46 | 43 |
|  |  |  |  |  |  |  | -40 | -38 | 46 |
|  |  |  |  |  |  |  | -42 | 5 | 26 |
|  |  |  |  |  |  |  | -44 | 24 | 23 |
|  |  |  |  |  |  |  | -40 | 4 | 35 |
|  |  |  |  |  |  |  | -4 | 21 | 41 |
|  |  |  |  |  |  |  | 8 | 21 | 41 |
|  |  |  |  |  |  |  | 36 | 25 | -10 |
|  |  |  |  |  |  |  | 38 | 21 | -1 |
|  |  |  |  |  |  |  | 30 | -56 | 40 |
|  |  |  |  |  |  |  | 32 | -60 | 51 |
|  |  |  |  |  |  |  | 26 | -71 | 51 |
|  |  |  |  |  |  |  | -36 | 50 | 20 |
|  |  |  |  |  |  |  | -30 | -62 | 1 |
|  |  |  |  |  |  |  | 48 | -38 | 48 |
|  |  |  |  |  |  |  | -20 | -87 | 10 |
|  |  |  |  |  |  |  | -18 | 1 | 15 |
|  |  |  |  |  |  |  | -55 | -42 | 9 |
|  |  |  | 2 > 0 back | Texture patterns | Identity | 12 | -48 | 23 | 25 |
|  |  |  |  |  |  |  | -42 | 19 | -1 |
|  |  |  |  |  |  |  | -34 | 5 | 29 |
|  |  |  |  |  |  |  | 38 | -49 | 37 |
|  |  |  |  |  |  |  | 10 | 31 | 28 |
|  |  |  |  |  |  |  | 48 | 32 | 22 |
|  |  |  |  |  |  |  | 50 | 21 | 25 |
|  |  |  |  |  |  |  | -26 | -43 | 41 |
|  |  |  |  |  |  |  | -30 | -66 | 35 |
|  |  |  |  |  |  |  | -38 | -45 | 39 |
|  |  |  |  |  |  |  | 36 | 23 | -3 |
|  |  |  |  |  |  |  | 26 | -72 | 7 |
|  |  |  |  |  |  |  | -14 | -38 | 20 |
|  |  |  |  |  |  |  | 38 | 11 | 25 |
|  |  |  |  |  |  |  | 38 | 3 | 27 |
|  |  |  |  |  |  |  | -6 | 27 | 35 |
|  |  |  |  |  |  |  | 20 | 7 | 14 |

*N.B. Contrasts within the same study conducted using the same sample were combined before they were entered into the meta-analysis.*
